# Evaluating the effectiveness of various small RNA alignment techniques in transcriptomic analysis by examining different sources of variability through a multi-alignment approach

**DOI:** 10.1101/2025.02.18.638849

**Authors:** Xinwei Zhao, Eberhard Korsching

## Abstract

DNA and RNA nucleotide sequences are ubiquitous in all biological cells, serving as both a comprehensive library of capabilities for the cells and as an impressive regulatory system to control cellular function. The multi-alignment framework (MAF) provided in this study offers a user-friendly platform for sequence alignment and quantification. It is adaptable to various research needs and can incorporate different tools and parameters for in-depth analysis, especially in low read rate scenarios. This framework can be used to compare results from different alignment programs and algorithms on the same dataset, allowing for a comprehensive analysis of subtle to significant differences. This concept is demonstrated in a small RNA case study. MAF is specifically designed for the Linux platform, commonly used in bioinformatics. Its script structure streamlines processing steps, saving time when repeating procedures with various data sets. While the focus is on microRNA analysis, the templates provided can be adapted for all transcriptomic and genomic analyses. The template structure allows for flexible integration of pre- and post-processing steps. MicroRNA analysis indicates that STAR and Bowtie2 alignment programs are more effective than BBMap. Combining STAR with the Salmon quantifier, or with some limitations, the Samtools quantification, appears to be the most reliable approach. This method is ideal for scientists who want to thoroughly analyze their alignment results to ensure quality. The detailed microRNA analysis demonstrates the quality of three alignment and two quantification methods, offering guidance on assessing result quality and reducing false positives.

## 1. Introduction

Sequence aligners have been in development since the 1970s and are continuously evolving with ongoing adjustments to algorithms and concepts [1] . By 1982, de novo sequence assembly approaches and programs had also been introduced to complement alignment methods [2].

Alignment primarily aims to identify near-perfect matches between sequences of interest and also to uncover conserved sequence regions across diverse biological species. Established algorithms like BLAST (basic local alignment search tool) [3], BLAT [4, 5], and FASTA (sequence comparison) algorithm [6, 7] are widely used for comparing sequences and detecting local or global similarities. These tools can analyze both protein and DNA sequences and facilitate translation between the two types.

In addition to the core alignment tools, there are numerous other alignment programs available. A recent review focused on short read alignment programs, summarizing current developments in sequence alignment tools and sequencing technologies [8] . Musich et al. [9] critically examined the differences between alignment programs, discussing various alignment and assembly algorithms in terms of their strengths and weaknesses, which can result in slightly to significantly different outcomes, requiring a more thorough analysis. Finally, the treatment of ’gaps’ also impacts alignment quality and may differ among alignment programs [10, 11].

The multi-alignment framework (MAF) presented here considers the differences in alignment tools and aims to efficiently utilize multiple algorithms simultaneously in various scenarios. Different data sets may require different alignment approaches, so it is recommended to compare alignment programs for each specific situation. As programs evolve, their behavior may change, making it important to have a flexible framework like MAF that can adapt to different applications and user requirements over time. The concept is not limited to a specific sequencing technology or procedure, such as ’bulk’ or single-cell sequencing. Instead, it primarily focuses on the alignment procedure itself.

The MAF is built on Linux and uses Bash commands to integrate alignment and post-processing programs into a unified workflow. The main Bash scripts provided are 30_se_mrna.sh, 30_pe_mrna.sh, and 30_se_mir.sh for single-end mRNA analysis, paired-end mRNA analysis, and small RNA analysis, respectively.

All the programs which are called from within these scripts are already well established like the alignment programs STAR and Bowtie2 as well as all the other programs summarized in the readme.txt file in the supplementary information file MAF.zip. In this file also all the links to further program resources are provided. The workflow is based throughout on the sequence data format fastq and BAM. Major steps are including quality control, trim adapters, optionally adjust the reads for other sequence features, optionally deduplicate based on read sequence similarities, align to genomic or transcriptomic references, and finally optionally deduplicate BAM file hosted reads by UMI barcode information. The system is open source based and detailed instructions are provided below.

The intermediate and final workflow results are organized in a folder structure sorted chronologically by an increasing number, each with a specific folder name. This allows for separate analysis of each major workflow result.

So far, each script file is a complete application that uses custom human- and program-specific references for alignment and quality control of microRNA and mRNA data sets. However, any additional references may be utilized if they exist or are generated.

The three Bash scripts are designed to handle data sets of different sizes and quantities, depending on the computing resources available. Our in-house experiments processed fastq files up to 10 GB in size and up to 200 samples on a server with 24 cores and 256 GB of memory.

In addition to the pre-built script and resources, the modular design of the scripts enables easy customization and modification of Bash code, as it includes all essential functions for creating your own application modules.

There has been debate about using containerized solutions like Docker or Singularity for this program framework. However, these may not be ideal for maintaining flexibility with program versions or resolving errors. A stepwise installation process can offer better control and understanding for biologists with some computer science knowledge. The solution should be user-friendly for biologists with basic IT skills, rather than requiring expertise in advanced technology. Another reason is that Docker is not ideal for handling large datasets and parallel computing.

The simple Bash script setup, which assigns a specific task to a specific script, was preferred over a universal Bash script with configuration files and a lot more script code due to the complexity of the latter approach.

The approach has been used in three published studies [12–14] and is currently being used in three ongoing studies. Table 1 explains basic terms to help less experienced scientists understand the technology more easily.

**Table 1.**
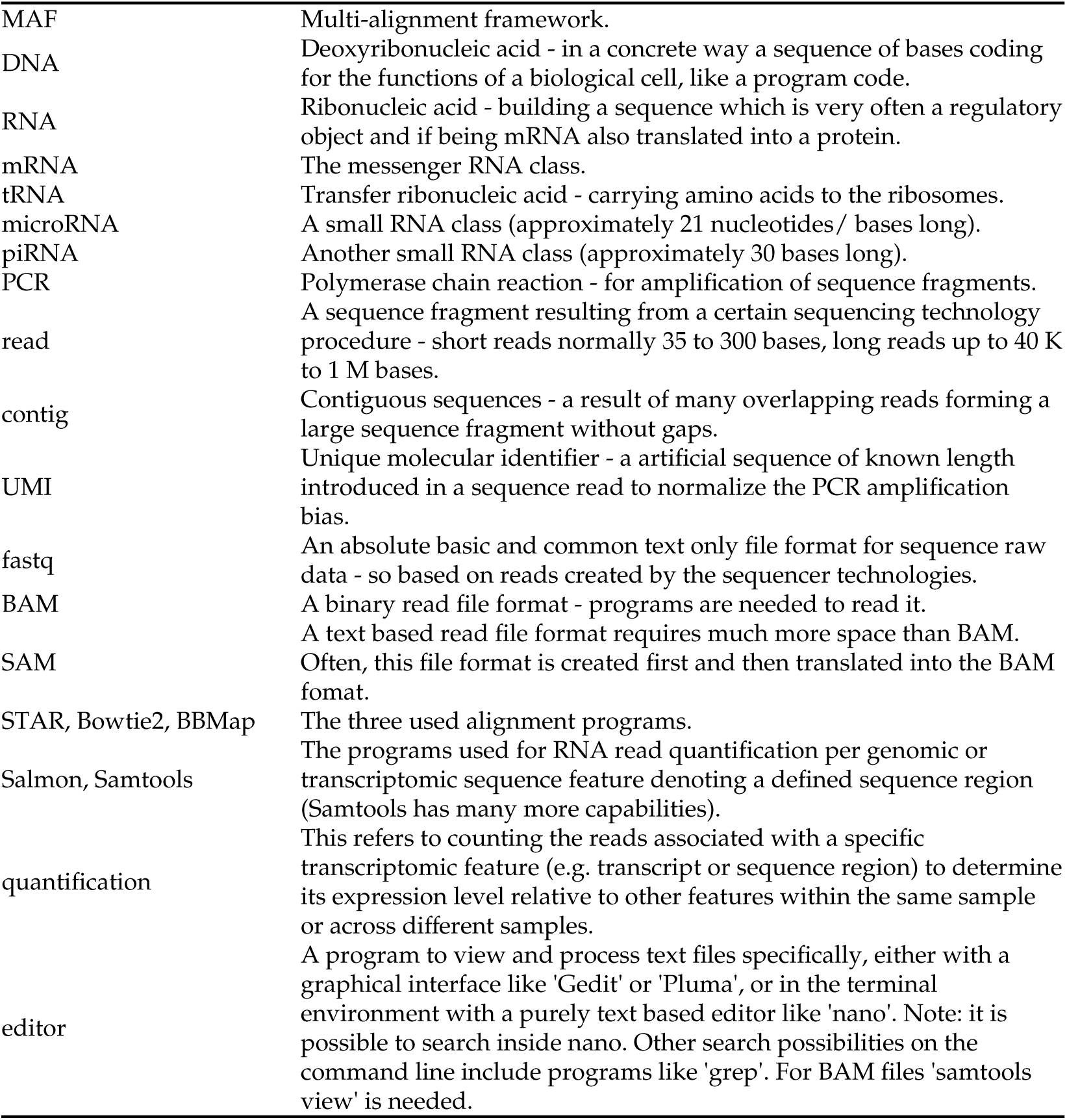
Terms and abbreviations placed in context. MAF Multi-alignment framework.

## 2. Implementation

### 2.1. Overview

The MAF analysis framework (Figure 1) aligns transcriptomic read data (mRNA or microRNA) and quantifies them in various ways. It serves as a versatile tool for sequence-related tasks.

**Figure 1.**
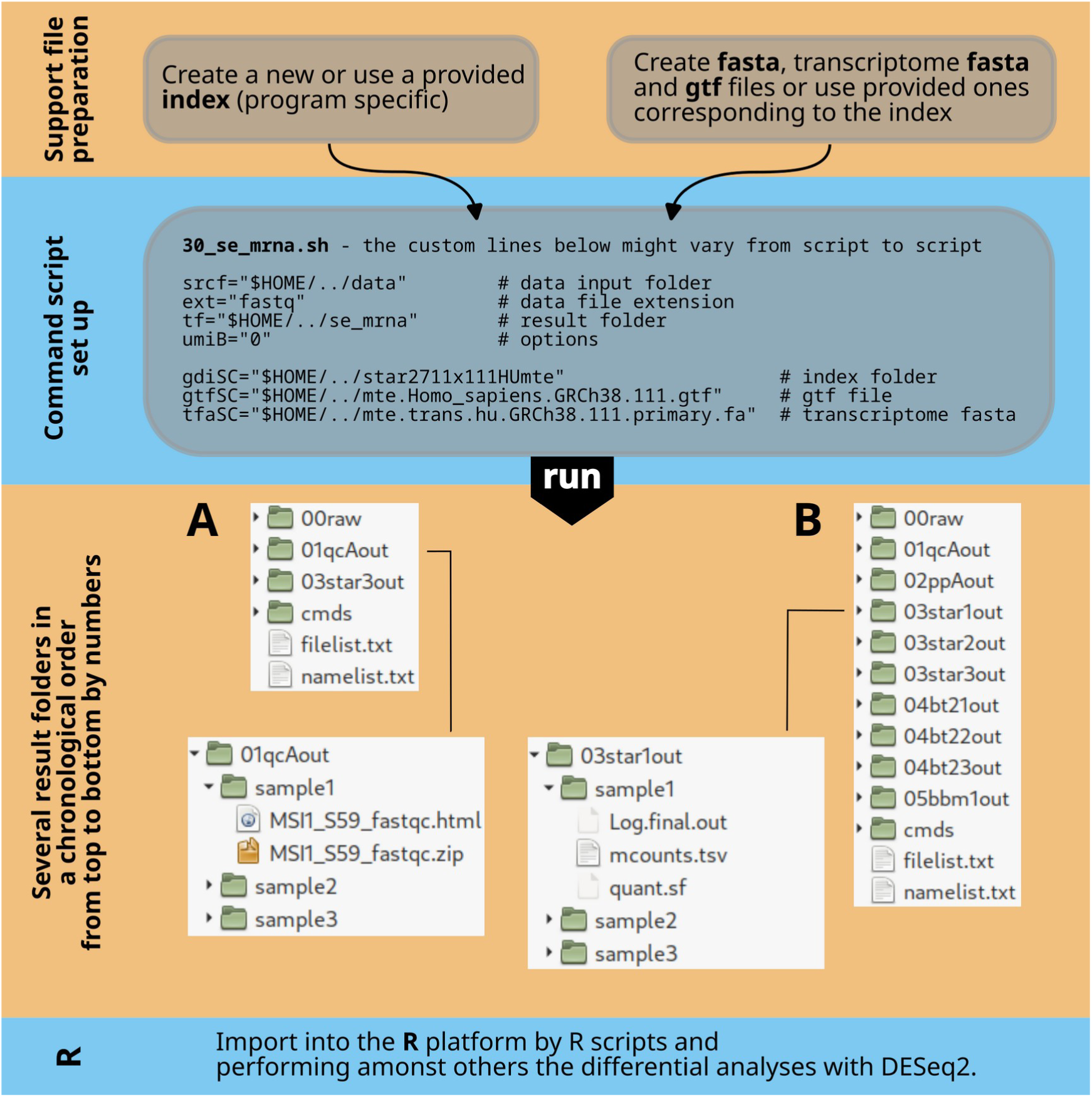
Overall workflow. The first step (top) involves organizing or creating all necessary files for alignment and preprocessing. The second step focuses on customizing a Bash script that combines different programs by adjusting path information and program options. The third step explains the output folder structure created by the Bash script, with result folders numbered in processing order and named after the tools used. For example, (A) shows this for scripts 30_se_mrna.sh and 30_pe_mrna.sh, while (B) does so for 30_se_mir.sh. The fourth step includes data analysis in R.

### 2.2. Prerequisites

The MAF scripts must run on a Linux operating system. Our examples have been tested on Debian 12, but they should work on any Linux version from 2024 onwards that supports Bash and Linux standard libraries. While program versions may vary slightly, this should not significantly affect the parameters of the Bash script command file.

The example provided (30_se_mrna.sh) demonstrates the analysis of ’single end’ mRNA data using one aligner, quality control, and UMI normalization. It serves as a basic introduction to the framework and its practical application, making it valuable for everyday use.

The same setup is available for transcriptomic ’paired end’ mRNA data (30_pe_mrna.sh) including a modified module for this purpose (icreatePE, instead of icreate in 30_se_mrna.sh).

The most advanced example for analyzing transcriptomic microRNA or small RNA data (30_se_mir.sh, single end) includes three aligners and seven pre- and post-processing tools, as well as additional utilities for a multiperspective view of data alignment and analysis results. All three scripts can be found in the /bin folder of the unpacked MAF.zip file (Figure 2).

**Figure 2.**
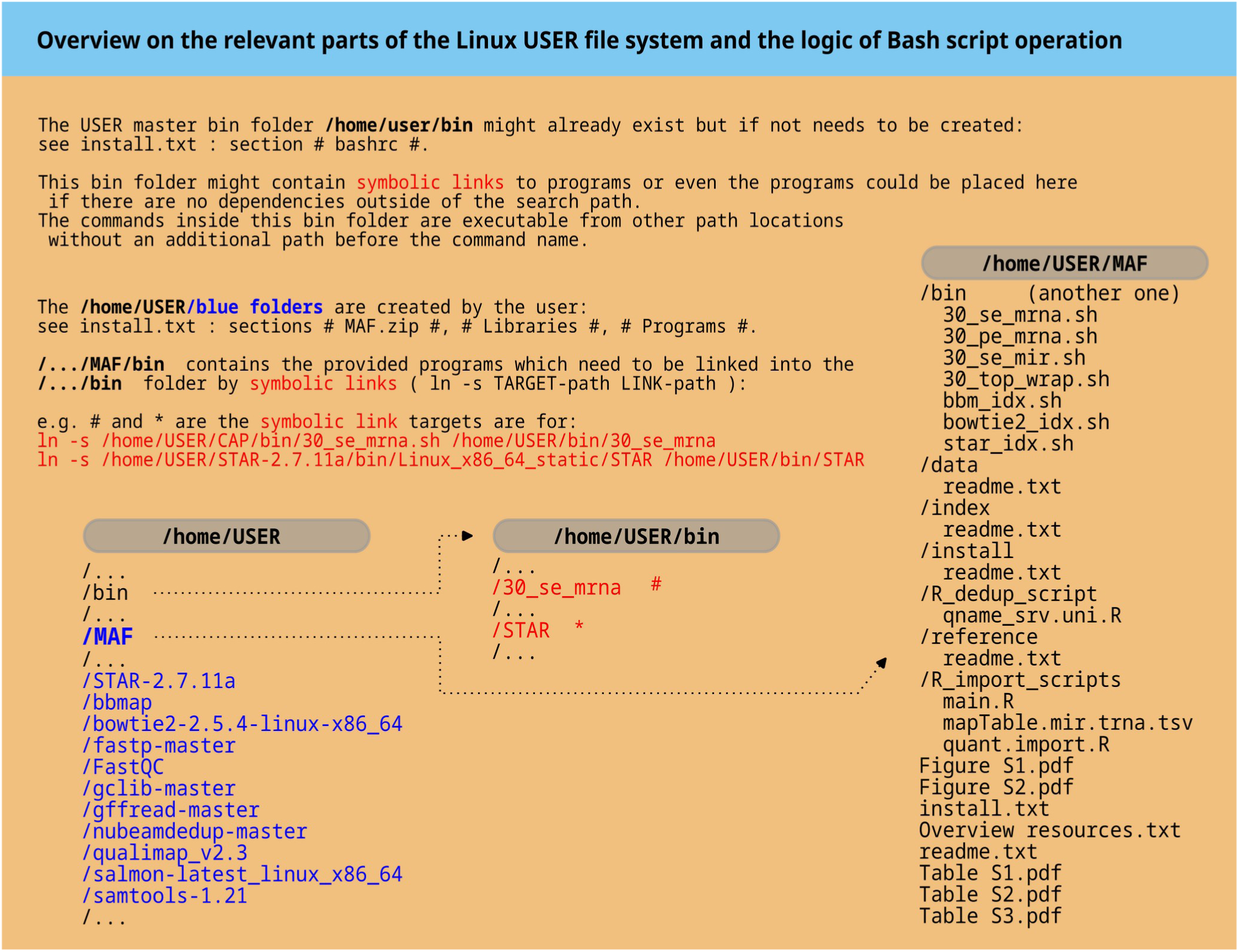
The master plan for folders. The left side of the diagram displays system folders such as ’home/USER’ and ’home/USER/bin’ that should be present after a standard Linux installation. The USER placeholder should be replaced with the actual username. The blue color highlights newly created folders after unpacking the MAF.zip file and installing all programs. The ’home/USER/bin’ path includes symbolic links to Bash command scripts in MAF/bin and installed programs like STAR (red). On the far right, the structure of the unpacked MAF.zip file is shown, including additional information files such as Figure S1.

All three approaches include minimal test data, index, and reference files to expedite the setup process. These files are available for download. Please be aware that the human reference files 111HUmt.tar.gz, 111HUmte.tar.gz, 111HUpmm.tar.gz are large (27 GB each).

Each command script requires customization of the parameter section for user-specific run parameters. After extracting the MAF.zip supplementary file to the user’s home folder, ensure all additional files are downloaded, unpacked, and placed in the appropriate subfolders of the MAF.zip folder structure (refer to Figure 2 and readme.txt files). All the relevant programs must be accessible and executable by the script commands. Detailed instructions for these initial procedures are provided at the top level of the MAF.zip folder structure in ’install.txt’ a step by step guide. If different program versions are needed, download locations are provided in the ’readme.txt’ file.

The R script functions to import the results into R [15] and for analyzing quantification count data can be accessed in the /R_import_scripts folder. A naive deduplication script is also available in the /R_dedup_script folder for the framework option barcodeB ($umiB="1") if needed.

In the folders /data, /index, /reference, there are ’readme.txt’ files that provide instructions on populating the empty folders (Figure 2). These instructions are essential for utilizing the provided resources, but alternative resources can also be utilized if desired.

### 2.3. Linux libraries

Several essential computer libraries need to be installed to establish a functional system. The details are included in the supplementary file ’install.txt’ in MAF.zip at section: # Libraries #. In the section # Programs # the installation of the utilized programs is described. It is good practice to follow all instructions in ’install.txt’ from top to bottom. The following section provides an overview of the programs used.

### 2.4. Sequence tools

Current versions of the used sequence tools can be found on the download page for installation. Simply place the compressed tool files in the /install folder of the unpacked MAF file. Additional installation instructions are provided in the ’install.txt’ file within the # Programs # section.

The list below briefly introduces tools and their purposes.

Samtools - an established program family for processing high-throughput sequencing data. The program collection is based on three basic elements : Samtools, BCFtools and HTSlib. Here we are only using Samtools [16, 17] with the sub programs: index, view, sort, markdup and idxstats (for quantification).

STAR - sequence aligner. The program is well described in publications [10, 11, 18] . This tool is used by the ENCODE project [19] and handles gapped situations very well.

Bowtie2 - sequence aligner. The recommended publications are [20, 21] . This program is widely used in the genomics community, although it can also be used for transcriptome analysis and can address gaps in version 2.

BBMap - sequence aligner. The tool collection is only partly published [22] and more detailed information can be found via links in the main ’readme.txt’ file. BBMap performs less perfect in the presented small RNA situation.

FastQC - quality control for fastq read files. Before starting any sequence alignment, it is essential to perform quality control on the sample(s) to asses their homogeneity and specific characteristics. FastQC [23] is a good choice for this purpose and is scriptable.

fastp - preprocessing of fastq read files, quality control, and deduplication. The fastp tool [24, 25] is specifically used for preprocessing purpose in the Bash scripts: deletion of adapters, cut and move UMI codes to the ’qname’ variable of the SAM format, and removing reads with a length smaller than 16 bases. In a second run, the tool trims CA artifacts at the start of the read and again removes reads with a length smaller than 16 bases.

nubeam-dedup - deduplicates sequencing reads without a reference genome. This approach is an option if no UMI codes are used and no perfect reference is available. Nubeam-dedup [26] is a pure C++ tool, so it is fast.

umi_tools - improves quantification accuracy by UMI code based deduplication. The umi_tools [27] is a Python tool set with 6 major program features, but only the "dedup" part is used here. Since it is Python, we recommend installing it via pipx as a self executable program, as described in detail in the supplementary file ’install.txt’.

Salmon - transcript quantification. The Salmon tool [28] is a very fast quantification program. The tool is used here in alignment mode, which is recommended. This tool also has a single cell extension called Alevin. There is a similar tool from a different group named Kallisto [29].

QualiMap - many sequence analysis functions. Only the transcript quantification feature is used in the context of this program [30] developed with biology in mind.

GffRead - not only performing a genome- to transcriptome fasta modification. GffRead together with GffCompare has a broad functional scope [31] . The tool is used here to create an adjusted fasta file necessary for quantification with the Salmon tool together with the transcriptomic BAM files of STAR (option: --quantMode TranscriptomeSAM).

## 3. In-depth guide to Bash command scripts

The central component that unifies various programs and data formats is a Bash script. Specifically, there are three command files named ’30_se_mrna.sh’, ’30_pe_mrna.sh’, and ’30_se_mir.sh’ in the supplementary file MAF.zip, located in the

/bin folder. A wrapper shell script ’30_top_wrap.sh’ is included to streamline multiple tasks and execute them from a top-level file, potentially spanning several days. Index creation scripts for alignment program-specific indices are also available for use with alternative references. Further details can be found in the help files of the alignment programs.

The wrapper script and other Bash files may require customization to suit the user’s computing environment. This customization is mainly done at the beginning of the scripts. If adjustments to the program parameters are needed, identify and modify the corresponding lines. The script names include version numbers for reference.

These script files can be viewed using a text editor, which is designed for pure text-based files like Bash scripts or small fastq sequence files. Using an editor with text coloring for Bash scripts can help in reading the commands more efficiently. For example, in a command terminal, you can use "nano [path/]filename," and in a graphical environment, you might use editors like Gedit or Pluma.

### 3.1. Single end and paired end Bash scripts

In the following paragraphs the ’30_se_mrna.sh’ and ’30_pe_mrna.sh’ files will be discussed as examples of a single-end respectively a paired-end mRNA analysis using the STAR alignment program, along with postprocessing support for deduplication.

The code for the single end alignment script ’30_se_mrna.sh’ begins with comment lines denoted by a hash character ’#’ followed by explanatory text. The hash character is indicating that the the following characters are not executed. These lines typically provide helpful information for the user or briefly explain the purpose of the following command line(s).

The ’parameters’ section at the start of the script file is essential for script execution.

It may need customization in a user-specific computing environment:

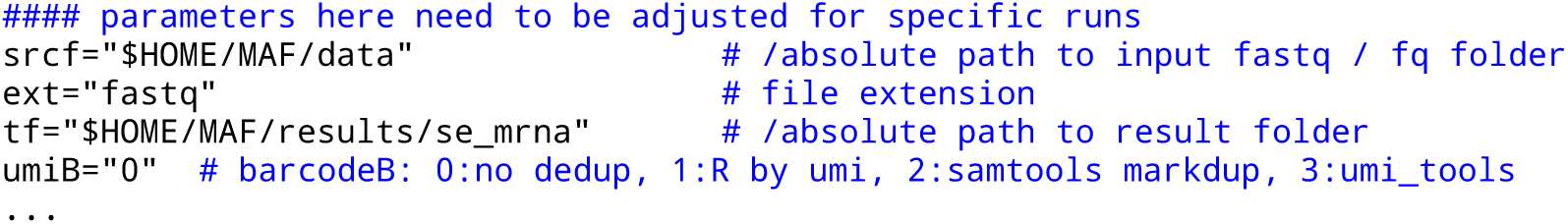

The srcf parameter or variable receives a quoted path to the fastq data folder. "$HOME" is a system variable representing the user’s home path, which can also be written literally. ext is the variable holding the file extension of the data files without the dot, typically ’fastq’ but occasionally also written ’fq’. The crucial aspect is that the file is itself in fastq format.

Before using a fastq file, it is recommended to check its format by opening a terminal and running the command: **`head -n 10 filename`**. This step ensures that the file is correct. For fasta files, some adjustments to the script may be needed, but for fastq files, everything should work smoothly. Compressed files with the **`.fastq.gz`** extension may require additional steps for compatibility with programs. It is advisable to start with an uncompressed fastq file for initial experiments (use command: **`gunzip name.fastq.gz`**).

The tf parameter specifies the path to a result folder. The folder may or may not exist. It is deleted and recreated in each script call, so be careful.

The umiB parameter has four options represented by numbers in quotes. ’0’ means no deduplication is performed, ’1’ uses a simple R script to unify UMI codes, ’2’ utilizes the Samtools markdup feature, and ’4’ (recommended) employs the umi_tools program to remove a PCR amplification bias. The last option is applied post-alignment and pre-quantification. For the R script option, a standard R installation is required.

The parameters gdiSC, gtfSC and tfaSC specify the paths for the STAR aligner index folder, the corresponding gtf file used in STAR index creation, and the transcriptome fasta file generated by GffRead from the fasta file also used in STAR index creation. For custom files, refer to **`MAF/bin/star_idx.sh`**.

At this stage, in a standard situation, all necessary steps have been completed, and the script is ready to run. The ’echo’ commands in the Bash script will display progress updates in the terminal window.

The following paragraphs describe the remaining program sections.

The term ’module’ with its corresponding number refers to a specific Bash function that combines multiple Bash command lines to perform a certain task. The sequence of Bash function calls at the end of the script determines the processing flow. Figure 3A illustrates all the modules in the command scripts for 30_se_mrna.sh and 30_pe_mrna.sh, while Figure 3B shows the modules for 30_se_mir.sh.

**Figure 3.**
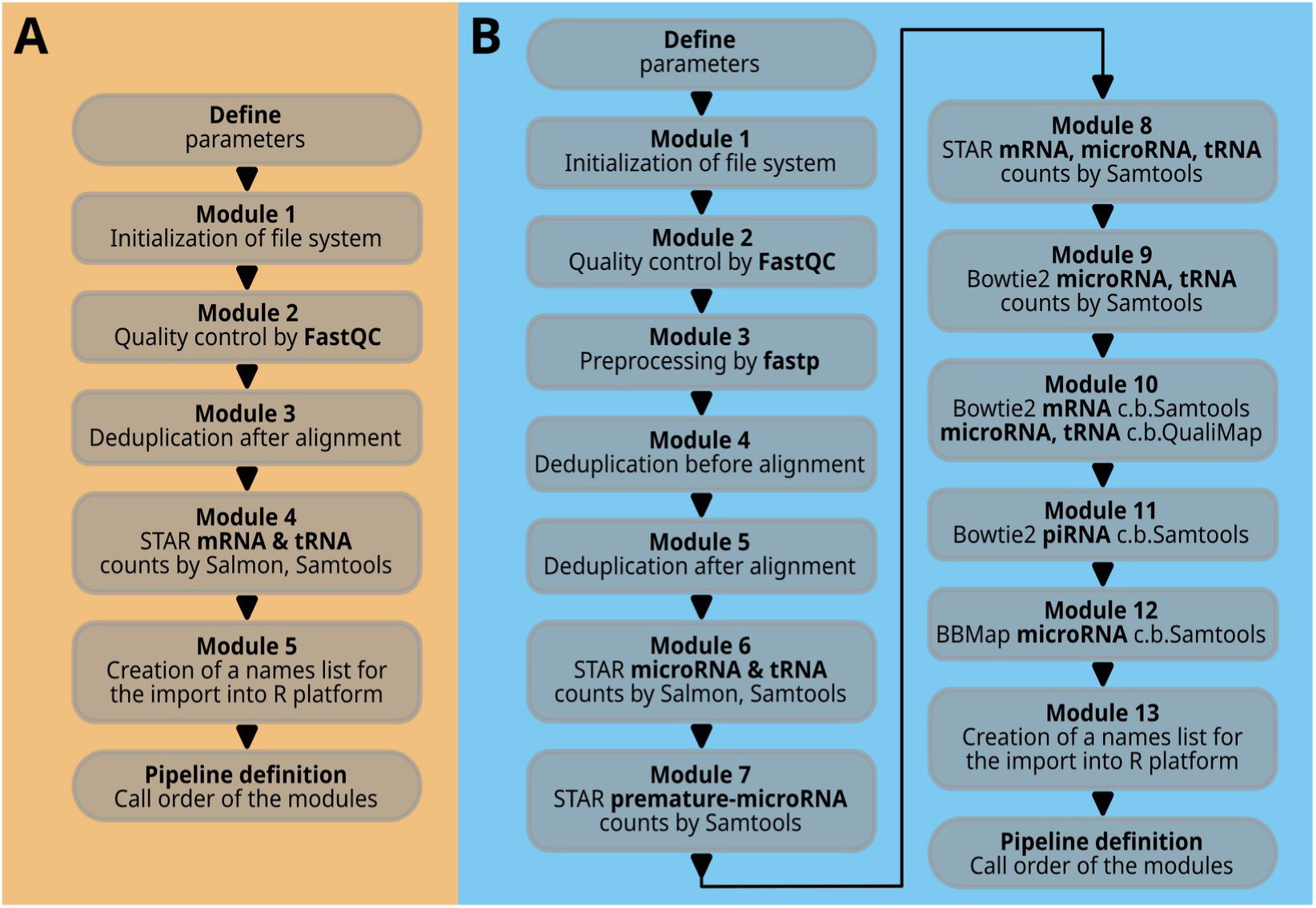
Code overview. In part (A), the modules of the single end or paired end workflow are listed. In part (B), the comprehensive microRNA workflow is presented, which is composed of many more modules. A more detailed description of all the modules is given in the text.

Module 1, internally named ’icreate’, creates the result file folder or deletes and recreates it if it already exists. It also creates symbolic links to the raw data files in the source folder within the ’00raw’ folder. Note that the two mRNA scripts 30_se_mrna.sh and 30_pe_mrna.sh have different functions (icreate, icreatePE) for single-end and paired-end situations with one or two fastq files. Interleaved files are not considered in this context.

Module 2, named ’qualcon’, and based on the FastQC program, generates an html file and a corresponding zip file for each raw data file. These files offer a detailed analysis of the data quality. The quality control files are stored in a folder named ’sample’ followed by a number, maintaining the order of sample names in the file filelist.txt.

The file structure is as follows: /result folder /01qcAout /sample1, with file names sample1_fastqc.html and sample1_fastqc.zip. This structure is repeated for each module and task. The file namelist.txt is used by R import functions to create a data structure in R with the original names. Changes in the bash code may require corresponding adjustments in the R code.

The next module is Module 3, referred to as ’barcodeB’. It is already described in the ’parameter’ section above. Each deduplication option is denoted by a code element comprising a few to several lines of code.

In all Bash functions, the code lines trigger the creation of a command line file, e.g. in this case called ’sub1.sh’, which is finally executed by ’cmds/sub1.sh’. The entire script generates several command line files, which are all collected in the /cmds folder. If there are issues with the script, examining these files may offer some clarity.

Module 4, STAR alignment (staroutC), integrates multiple tasks including running the STAR command line with various parameters, performing Samtools actions, counting with **`samtools idxstats`** and Salmon, and removing intermediary files. These tasks are organized in command files (e.g. mapstC.sh, sc3C.sh, cu1c.sh, sc3C.sh, sal3.sh, cu1c.sh) and executed step by step. It is important to mention that the STAR command line differs slightly for single-end and paired-end data.

The xargs command is often used in the scripts. The ls command generates a list of sample files, which is then passed to xargs. Xargs creates an output command line for each input element based on a template command line in quotes after the xargs command. This output command line is saved in a Bash script file using the ’echo’ command. ’>’ is used to save and overwrite, while ’>>’ appends to the file.

Module 5, named ’fnlA’, extracts the original sample names from the file filelist.txt. This list is essential for translating the ’sample1’ nomenclature back to the actual sample name when the results are imported in the R platform. The cut and tr commands are utilized to extract a specific name element from the input text file.

The final part of the Bash command script runs all the individual Bash functions (modules) in a defined sequence. ’echo’ statements provide feedback in the terminal about the successful execution of each module. The ’date’ command reports the program’s runtime.

### 3.2. Small RNA Bash script

The script for microRNAs and tRNAs, or small RNAs in general (30_se_mir.sh), is depicted in Figure 3B and is more complex. Upon closer examination, it is evident that the basic structure of the modules closely resembles those described earlier. Therefore, only selected important code elements are discussed here.

The microRNA script begins with a longer list of parameters required for its execution. These parameters are organized in the order of the script, starting with general elements such as the source path, followed by sections for the STAR aligner, Bowtie2, and BBMap. All of these file or folder links must be accurate and relevant to the alignment task at hand.

Modules 1 and 2 are similar to the previously mentioned files 30_se_mrna.sh and 30_pe_mrna.sh. A new addition is Module 3, named ’preprocess’, which utilizes the program fastp to remove adapter sequences, relocate the UMI identifier to the ’qname’, trim the CA prefix, and filter out reads smaller than 16 bases. This module is designed for single end reads and generates additional quality control files (fp1- and fp2.html and .json) that can be accessed through a web browser.

Another new module is Module 4 (’seqcodeA’). This module is designed for deduplicating sequences using sequence similarity rather than barcoding methods.

Module 5 (Figure 3B) corresponds to Module 3 in Figure 3A, which is based on barcoding concepts.

Modules 6, 7, and 8 (’staroutA’, ’staroutB’, ’staroutC’) are STAR aligner-based modules. The first module analyzes microRNAs and tRNAs, the second focuses on premature microRNAs, and the third includes microRNAs, tRNAs, and mRNAs. The latter can be useful for identifying sequence similarities between mRNA sequences and small RNAs, potentially resulting in ambiguous counts.

Modules 9, 10, and 11 (’bt2outA’, ’bt2outB’, ’bt2outC’) use Bowtie2 and provide a comparative third perspective on micro- and tRNAs. The QualiMap tool on top of Bowtie2 provides an independent fourth perspective on tRNA and microRNA quantification. Chromosomal and consolidated contig counts offer an overview of transcriptome feature distribution across large genomic elements. Module 11 focuses on piRNAs, a small RNA domain with minimal overlap with microRNAs. Overlaps were identified using blastn, with microRNAs given priority in custom reference creation.

Module 12 (’bbmoutA’) utilizes the BBMap aligner to detect microRNAs, making it the fifth approach in this category.

The last two modules closely resemble the script approaches shown in Figure 3A.

### 3.3. Formats - understand the role of fastq, fasta, GTF and transcriptome fasta files

As previously stated, the input file format follows the fastq format. Here are two examples records:

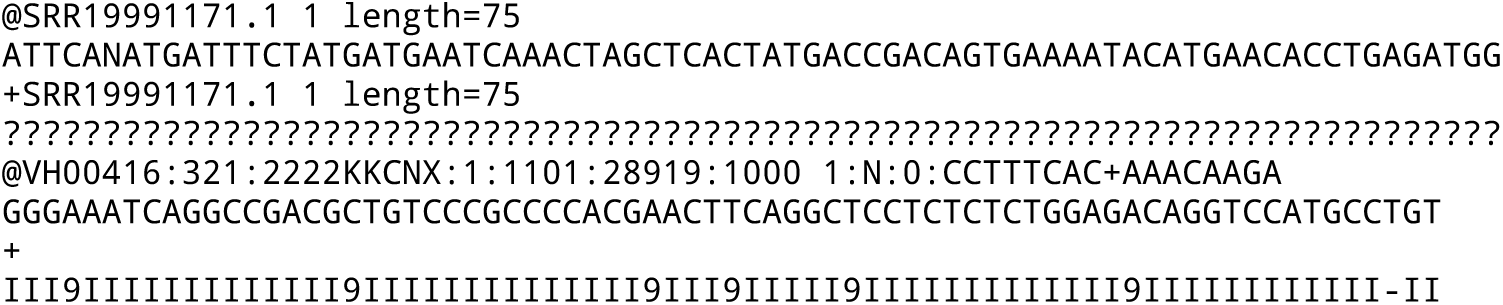

Each record consists of four lines: the first line starts with a unique identifier beginning with ’@’, possibly followed by split characters ’:’ and barcode sequences. The second line contains the sequence, the third line has a ’+’ with or without (standard) an identifier, and the fourth line contains the quality sequence. The quality sequence uses Phred scores, representing the probability of an incorrect base call from 1 to 0.0001, denoted by symbols !"#$%&’()*+,-./0123456789:;<=>?@ABCDEFGHI.

The fastq file may be compressed with a ’.gz’ algorithm or other compression formats. It is advisable to decompress the files before running the scripts. While the script can be modified to handle gz compressed files, not all tools may support this.

Starting with uncompressed files is more general as many programs have their own decompression mechanisms.

If the quality information is missing the ’fasta’ file format is used. It begins with the ’>’ character followed by a sequence name, and the sequence itself on the next line. Some institutions prefer a line break every 80 characters, but there is no strict standard. The sequence is gap-free, and the next continuous sequence starts with a name line, a line break, and the sequence:

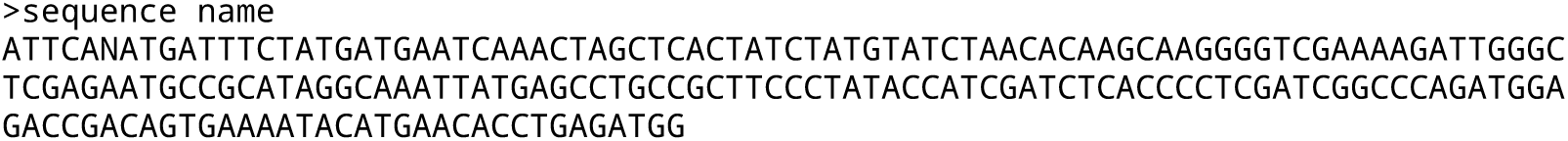

A genome fasta file is required to create index data structures for STAR, Bowtie2, and BBMap, each comprising multiple files. Scripts like star_idx.sh, bowtie2_idx.sh, and bbm_idx.sh are available to help generate new index files.

The human genome fasta file has many lines until the next chromosome name appears. The nano editor in the terminal might be able to handle such large files, at least for viewing and searching, but the computer’s main memory may be a limiting factor.

The STAR star_idx.sh script includes a call to the gffread tool to create a transcriptome fasta file from the primary genome fasta file, which is needed for Salmon quantification. Samtools idxstats quantification does not need this reference. It is important to understand that Samtools counts data without normalization for paired-end situations, unlike Salmon. In cases of strong overlapping paired-end data, Samtools counts may be approximately twice of those in Salmon.

The GTF file describes features in a fasta sequence file, with key information like start and end positions (columns 4 and 5) and annotations. The chromosome name in column 1 should match (!) the genome fasta chromosome name (e.g., ’1’ or ’chr1’). Not all columns are displayed below:

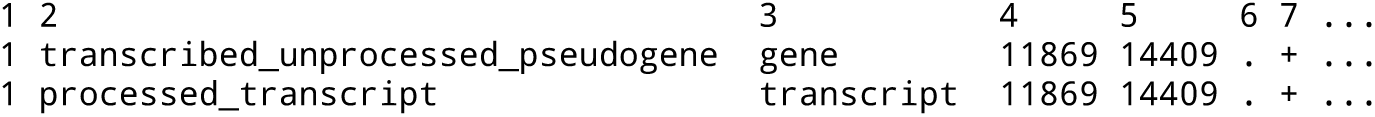

It is possible to convert a GFF file to a GTF file (e.g. **`gffread f.gff3 -T -o f.gtf`**).

Standard GTF files can be used for standard feature information. In the described microRNA workflow, microRNA and tRNA information from the respective databases is integrated into the standard GTF files at the appropriate place. The same is also done for the piRNAs. This integration is also carried out for piRNAs, as Ensembl and other main databases lack these features. Hence, the reference files provided hold significance for small RNA projects.

Reducing the features to focus on specific targets can greatly decrease run time. However, it’s important to note that viewing the entire genome can uncover signal crosstalk between features or transcripts. The microRNA script offers various approaches, including whole transcriptome views, to address this.

For detailed information on alignment procedures, parameters, and files, refer to Table S1 in the supplementary MAF.zip file.

## 4. Results

As described in the opening, this concept has been utilized in published and ongoing studies. This section delves into the rigorousness of the algorithmic outcomes to supplement existing information.

The study is conducted using Ensemble version 111 with an extension for microRNA and tRNA, and the sequence data is provided in the ’Data Availability Statement’ section at the end.

The microRNA script (30_se_mir.sh) was used on two human samples sets: a) proliferation features of eutopic endometrium from women with adenomyosis, GSE207522, here termed DE-1|divided into two conditions g1, g2 and b) microRNA expression in endometriosis and suppression of cell migration, GSE275002, here termed DE-2|also divided into two conditions g3, g4. These two different studies are analyzed each with the 30_se_mir.sh script in 5 different technical conditions concerning microRNAs and tRNAs and in a one technical condition for premature microRNAs. Further results, such as piRNAs and chromosomal views, are not explicitly considered or described here, but they behave technically similarly.

The study aims to analyze general and specific count distributions and count proportions to detect any trend in expression characteristics and the associated result qualities. It also seeks to understand how significant candidates establish their significance, which may lead to rejection upon further examination. These analyses shed light on the data complexity and the challenges of identifying robust and well-annotated candidates. In addition to the results section, more detailed information and core R scripts for personal evaluations and handling complex data sets can be found in the supplementary information, specifically Statistic S1, S2.

The microRNA script results are imported into R using further R scripts found in /MAF/R_import_scripts. Two differential analyses are performed using the DESeq2 R package, and R graphs are generated to visualize the results. It is important to be aware that the DESeq2 procedures may slightly modify the raw data counts due to variance smoothing ([32] and the Bioconductor page). Therefore, comparing the DESeq2 counts from the differential analysis with the imported raw data counts may be necessary.

Figure 4 shows the raw data after import, indicating that the majority of counts correspond to RNA transcripts that are not part of the microRNA family. The small RNA library kit focus on fragments smaller than 100 bases, suggesting that other RNA transcript families may be present as full molecules or fragments that can be detected through alignment procedures.

**Figure 4.**
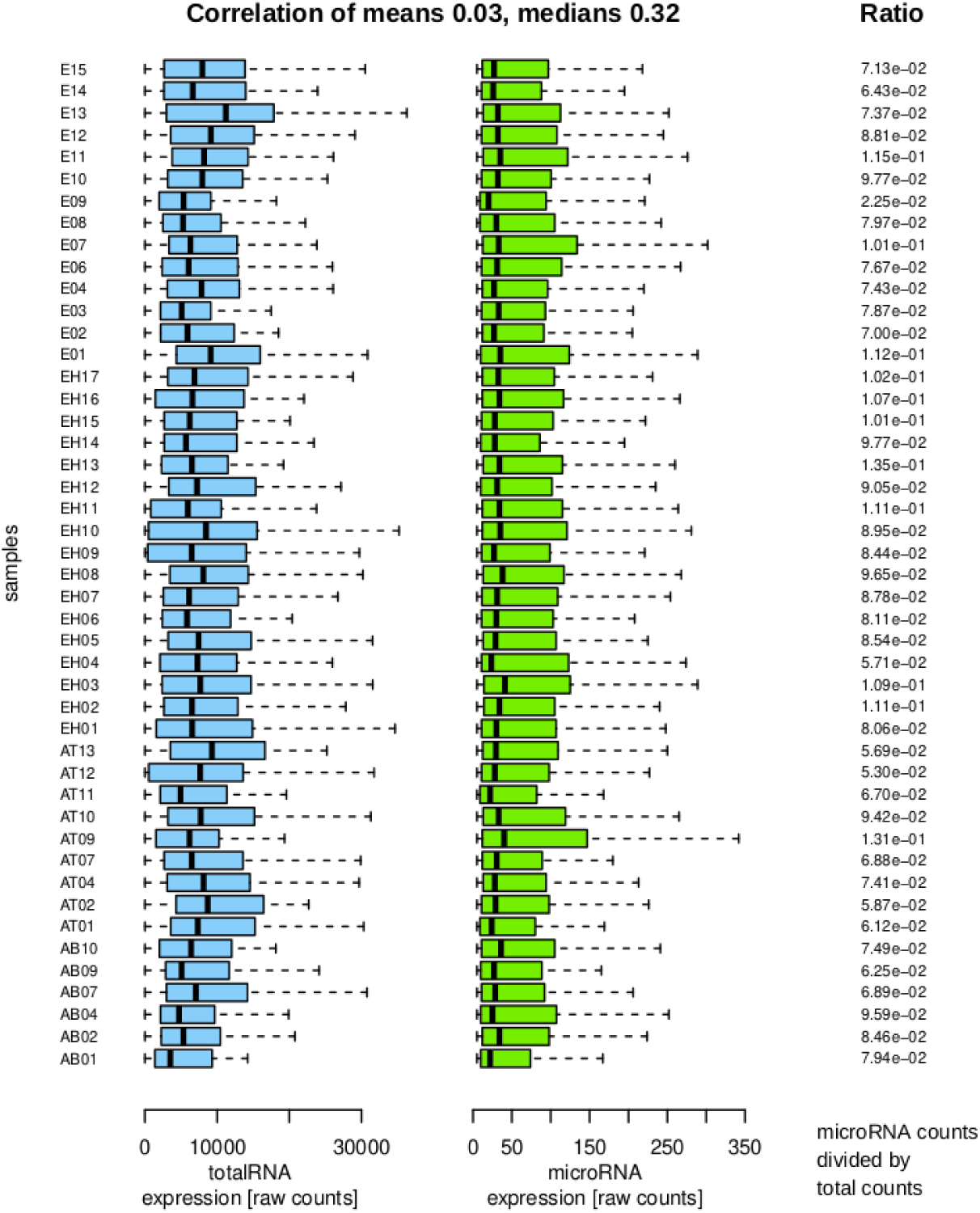
Sample distributions and properties. The left side shows sample acronyms with blue boxplots for total RNA counts and green boxplots for microRNA expression counts, indicating lower counts in the microRNA fraction. The right side displays ratios of total RNA to microRNA counts per sample. The correlation coefficient between mean and median values of total RNA (blue) and microRNA (green) groups is calculated at the top.

It is decisive for the differential analysis to assess whether the total counts and microRNA counts per sample exhibit a consistent trend, as this could invalidate the differential results. The mean values per sample show a weak positive correlation, suggesting a lack of strong association. The median correlation, while higher, provides less insight into the distribution characteristics. The features display significant heterogeneity, diminishing the positive correlation. Analysis of the ratios indicates that the samples do not display extreme variability within or between groups, supporting their suitability for the final differential analysis.

The differential analysis mainly relies on the group variance, the size of each group, and the absolute count numbers per feature. Statistical analysis at this level is recommended but not explicitly shown here but the following graphs revisit this topic from a different perspective. Table 2 offers more insight into the final significant candidate numbers from the pool of known microRNAs.

**Table 2.** Number of significant results for the differential analyses with DESeq2. In this study, DE-1 and DE-2 refer to two differential DESeq2 analyses with Benjamini-Hochberg correction for multiple testing. ’of’ indicates the input number of non-negative expression features. ’mt’ is an abbreviation for microRNA and tRNA. ’pmm’ stands for pre-mature microRNA. Further details on the five mature microRNA strategies can be found in Figure 5 Venn diagrams and supplementary Table S2. Details on procedures can be found in Statistic S1.

| A |  |  |  |
| --- | --- | --- | --- |
| MicroRNA | DE-1 | DE-2 |  |
| of 1074 | 46 | 14 | STAR mt index, Salmon quantification whole transcriptome (sts). |
| of 1076 | 46 | 12 | STAR mt index, Samtools quantification idxstats (stm). |
| of 1359 | 22 | 11 | Bowtie2 mt index, end-to-end, Samtools quantification (btq). |
| of 731 | 45 | 14 | Bowtie2 genome index, Qualimap quantification, mt reference (btm). |
| of 2016 | 0 | 2 | BBMap mt index, Samtools quantification (bbm). |
| of 994 | 49 | 17 | STAR pmm index, Samtools quantification (pmm). |
| B |  |  |  |
| tRNA | DE-1 | DE-2 |  |
| of 141 | 6 | 24 | STAR mt index, Salmon quantification whole transcriptome (sts). |
| of 142 | 6 | 24 | STAR mt index, Samtools quantification idxstats (stm). |
| of 257 | 35 | 15 | Bowtie2 mt index, end-to-end, Samtools quantification (btq). |
| of 235 | 18 | 17 | Bowtie2 genome index, Qualimap quantification, mt reference (btm). |
| of 256 | 8 | 8 | BBMap mt index, Samtools quantification (bbm). |

In addition to a single differential analysis alone, implementing a sampling procedure at the input data level could serve as a good control for the robustness and number of candidates in the result set. This could be achieved by sampling the expression values of samples.

Following the perspective on the differential analysis reveals a strong coherence between the STAR and Bowtie2 aligners, with some differences observed. However, BBMap (line five) encounters challenges in handling microRNAs in this dataset.

Figure 5 and supplementary Table S2 illustrate the overlap of established candidates using various technical approaches and alignment procedures. Differences in results can be observed with regard to the associated p values when using two different quantification algorithms based on the same STAR alignment. More pronounced discrepancies are seen between the STAR and Bowtie2 approaches, as well as between the Bowtie2 approach and QualiMap-based quantification, where significant candidates may disappear or new ones may emerge in the result lists.

**Figure 5.**
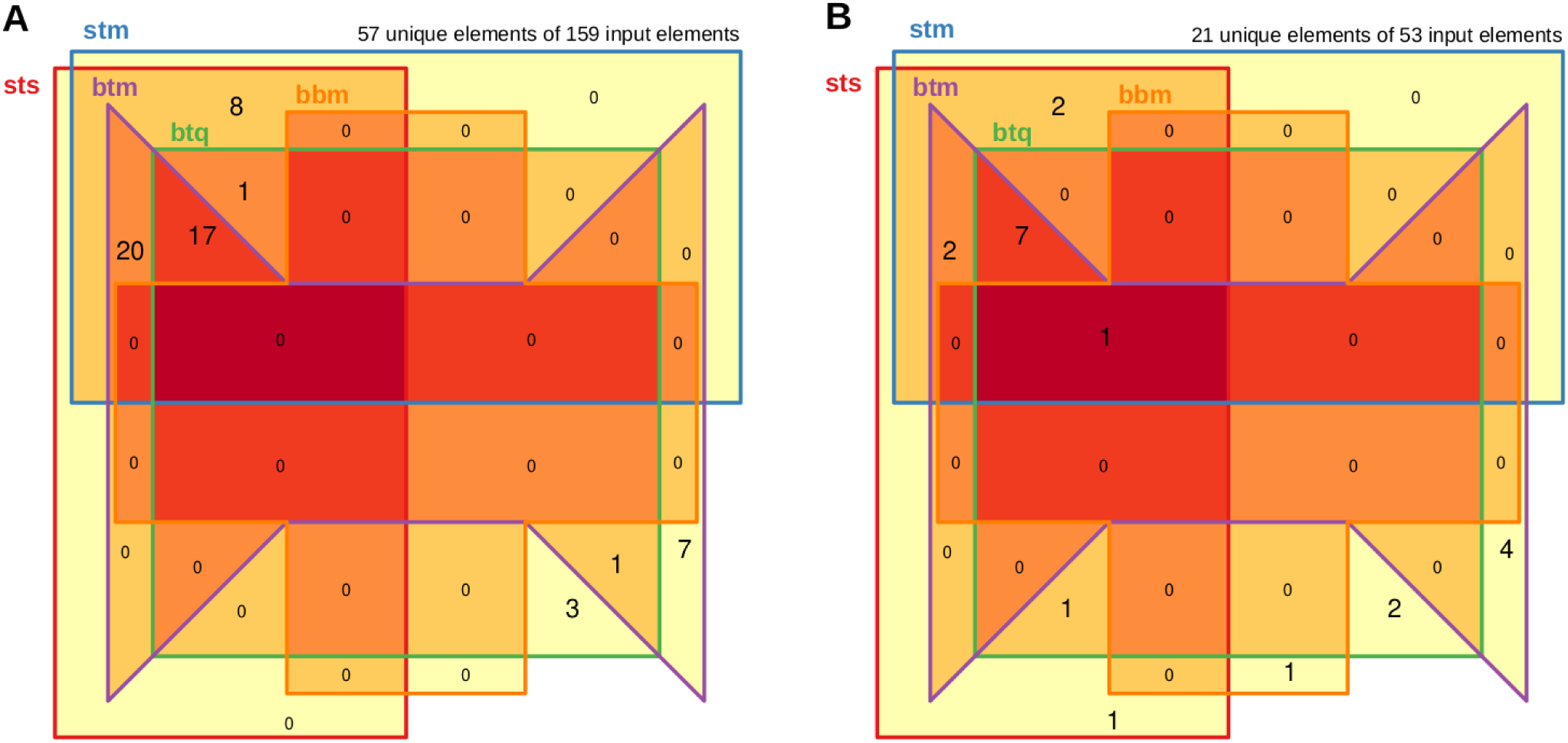
Overlap of significant candidates between the 5 algorithmic approaches. Panel A displays the comparison of the DE-1 set, while Panel B shows the comparison of the DE-2 set. Each set consists of five mature microRNA approaches, illustrating their shared significant candidates. The darkest red color indicates overlap across all sets, with yellow highlighting unique features in each group. The nomenclature of the five analyses, from ’sts’ to ’bbm’, is detailed in Table 2. The total number of unique features compared to all features of the five groups is shown at the top, forming the basis of this Venn diagram. Unique features may vary in distribution among the groups.

A statistical perspective on the overlaps is provided in supplementary information Statistic S2. Here, four approaches based on two models rank the framework alternatives STS to PMM based on Table 2A. Beyond the direct comparison of overlaps, the statistical approach provides a test-based ranking between the approaches and emphasizes the first exploratory analysis.

BBMap is not suitable for analyzing microRNA data as it fails to detect certain microRNAs that are identified by other tools and vice versa. In one analysis, it misses detecting any microRNAs, and in another analysis, it only confirms one candidate while introducing a new one that other tools do not find. Detailed examination reveals that BBMap generates counts for a specific microRNA that other tools report as zero counts.

This microRNA’s sequence, with a pattern of one A, three Gs, and a 10-base stretch of A in the mature proportion of the sequence, highlights a different performance of the BBMap’s algorithm in handling such sequences. The prominent poly A region might be one of the sequence motifs here which trigger the difference. The sequence details can be seen in Supplementary Table S3, based on the miRBase database and the analysis of aligned BAM files of each sample. It is important to note that the variability in the sequence itself is low, but the count variability between the samples is high, even when the selected groups were taken into consideration. Nevertheless, regarding poly A, not only BBMap might be the problem, but the other aligner might also misinterpret the situation by assuming the setup of an mRNA molecule class here.

The premature sequences aligned with STAR shown in line number six of Table 2A indicate that the overlap of significant premature microRNAs with mature microRNAs using STAR or Bowtie2 ranges from 24% to 33% for the first differential set (DE-1) and from 34% to 41% in the second differential set (DE-2, see supplementary Figure S1). Although belonging to different biological states, these two molecule classes share a common biological pathway and have overlapping sequences.

This sequence overlap can result in crosstalk between premature and mature forms due to alignment algorithms that struggle to differentiate between them effectively. However, the findings of premature microRNA analysis are consistent with those of mature microRNA analysis, and vice versa, even though regulatory conclusions may be less definitive. A further approach would be to define control sequence in the premature part of the sequence absent in the mature part of the microRNA and to compare those two result sets.

Table 2B displays results related to tRNAs, which are significantly longer and serve another biological function compared to microRNAs. The distribution pattern of differential candidates in this table differs notably from that of microRNA results, indicating that we are identifying molecule class-specific outcomes rather than just echoing expression strength trends. Additionally, it is noteworthy that the BBMap results exhibit a better performance with this particular molecule class.

Figure 6 highlights the robustness of individual count distributions for each molecular candidate. This approach allows for filtering out significant candidates that exhibit a skewed distribution. Absolute count numbers are also considered to ensure alignment with basic biological assumptions.

**Figure 6.**
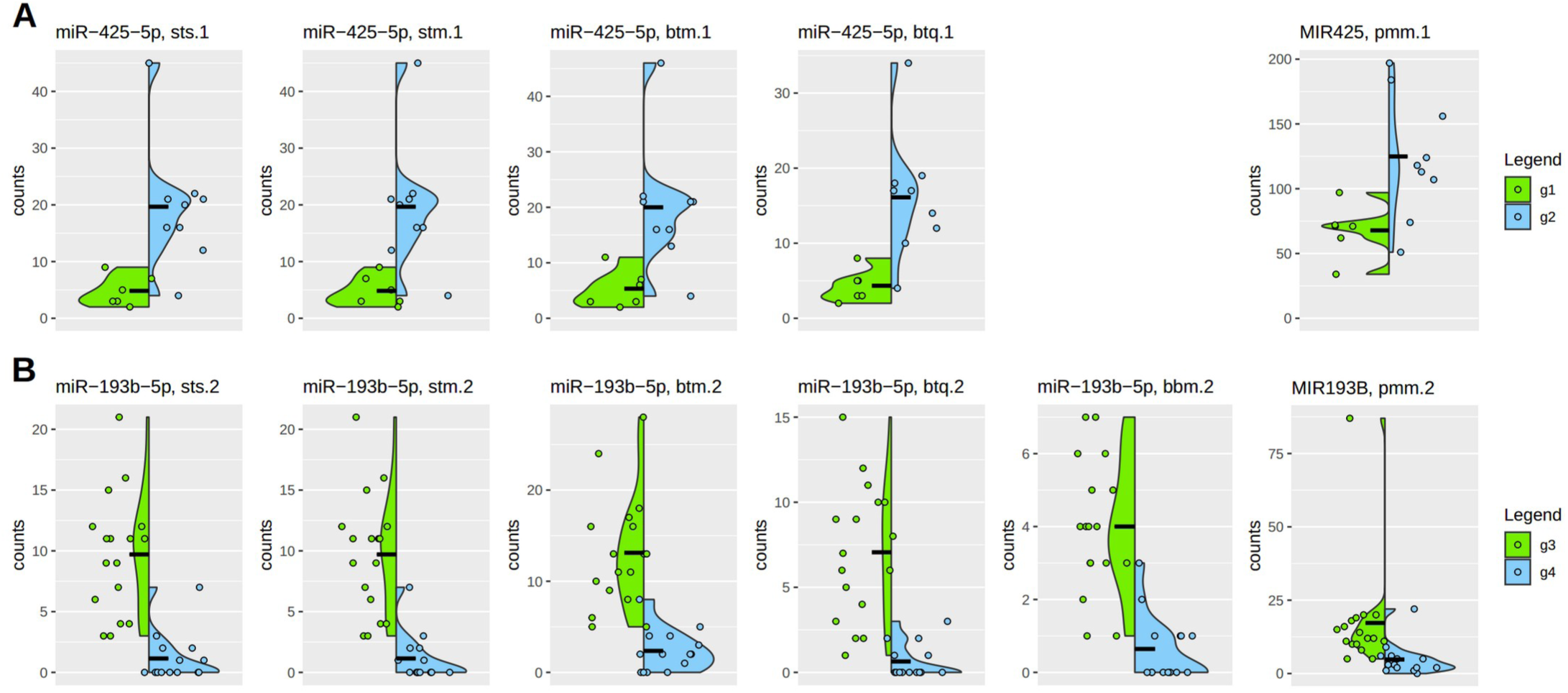
Individual count distributions in highly significant microRNAs. The results of the two differential analyses are shown as green and blue vertical density curves as a violin plot. Sample sizes are as follows: g1 (6), g2 (9), g3 (17), g4 (14), so including 3 unpublished samples in g2. Individual counts are represented by colored dots. Because many count values in a low count situation have the same value these values are scattered horizontally. Vertical differences indicate scale variations. The left side of the plot shows mature microRNA results, while the right side displays premature microRNA results. The naming scheme indicates the difference between mature and premature states. Acronyms in the header lines are explained in supplementary Table S1 and represent different algorithmic approaches. The number following the acronym indicates the differential comparison: one for DE-1 (g1-g2) and two for DE-2 (g3-g4).

Under this perspective, the variation between different algorithmic procedures resulting in significant candidates is small. Premature microRNA forms are typically more abundant than mature forms, as expected. However, the higher count numbers do not significantly impact the strength of the differential effect. Again it can be seen that the BBMap algorithm in the case, which is significant and coherent with the other algorithmic procedures, behaves a bit different and even better. But this is as discussed above not the typical performance of the BBMap algorithm in the case of microRNAs as can be seen in DE-1 where zero of 57 candidates are supported and in the case of DE-2 of 21 candidates only this one is supported.

## 5. Discussion

The fundamental idea behind the presented method is to establish a workbench or framework for analyzing alignment and preprocessing variabilities. The main objective is not to simply benchmark average performance on datasets but rather to highlight the significance of data variability and pinpoint its root causes. In addition to this conceptual goal, specific objectives can also be accomplished.

The global distribution of count variations [33, 34] shows variability per sample. The main concern is whether these variations distort subsequent differential gene expression analysis. As a surrogate marker, the coherence between total sample counts and molecule class-specific sample counts was investigated. Pearson correlation was used on sample mean values and sample median values, as well as the ratios between total and molecule class specific sample counts, providing initial insights into variability. The results show consistent trends, although the quantitative conclusion suggests two expression strength groups. Overall, the data behavior is still acceptable to proceed to more molecule class-based perspectives.

In addition to exploratory data analysis, the differential analysis [35–37] involves ranking and filtering group-based count pools using statistical tests. Not only are statistical tests applied at this level, but in this case, a normalization concept called ’variance smoothing’ is also implemented as one of several customization steps in the DESeq algorithm. The framework generates six microRNA result sets for an overlap analysis to aid in decision-making on the most effective approach based on the data. The suboptimal outcome of BBMap is evident in the remaining counts post-differential analysis and in the statistical analysis in Statistic S2. However, a more important aspect may be the examination of sequence properties presented in Table S3, which indicate sequence-specific alignment issues, although the extremely low read count itself may also suggest artifacts.

The individual analysis of count number distributions of single significant candidate microRNAs, superimposed by its group-based density curves, is a final filter on the data results by discarding those microRNAs that show outliers or uneven distributions.

The framework is therefore useful to support several important aspects of sequence alignment and associated data analysis to yield robust results.

The library preparation procedures [12, 13] in this context limit the size of the detected RNAs to smaller than 100 bases. Therefore, only small non-coding regulatory RNAs like microRNAs can be detected. Differences in the detected microRNAs have a direct impact on the identified downstream regulatory pathways of the cellular system. But in many cases, a few false positives or false negative factors might be compensated for by other factors in a downstream GO analysis [38]. Even on a broader scale, the identified differential pathways may not alter the outcome phenotype due to the compensatory and robust nature of cellular biology. However, this depends on the specifc case.

If the variations between different methodological procedures increase, this results in a completely different identified functional context. For instance, a minor discrepancy in DE-1 between STAR and Bowtie2 (9, 37, 4) or more significant disparities with QualiMap (28, 18, 4) may still show consistent trends in a GO analysis. However, in a scenario like STAR versus BBMap (46, 0, 0), where the differences are extreme, the results from STAR alone dictate the outcome of a GO analysis, but this is beneficial only if STAR performs effectively.

The functional context [39] is important because small RNAs, specifically microRNAs, have a one-to-many relationship with their interaction partners (see target prediction at https://mirdb.org/). On one hand, this compensates for a total failure in one potentially crucial molecule, but on the other hand, it spreads the effect to many molecules. This complex nature makes it difficult to identify the major related pathways. In this context, differences in the alignment procedure are likely to lead to small to large variations in the identified biological pathways.

When proceeding with validating a subset of primary candidates through additional wetlab procedures, it is likely that some validation candidates will again either fail or succeed. However, misinterpretation can occur due to incorrect primary guiding hypotheses, leading to a loss of conceptual context and additional potentially incorrect regulatory considerations based on the validation results. This kind of error propagation leads to a loss of systemic insight.

## 6. Conclusions

The MAF framework and protocol aid researchers in understanding how technology and software tools impact their results, helping them to choose robust candidates. The time investment is reasonable as a significant amount of work has already been put into developing a basic framework, customized indices, and corresponding references, especially for the microRNA, tRNA, and piRNA situation.

For microRNAs, STAR and Bowtie2 outperform BBMap in alignment. Overall, STAR appears to be the most stable procedure. However, each procedure has its strengths, justifying the need for comparison of different algorithms with each dataset under investigation. Differential analysis reveals that significance alone may not indicate relevance, with count distributions providing additional insight.

The MAF approach enhances researchers’ bioinformatics skills, offering flexibility and open-source principles to explore computational methods’ strengths and limitations from multiple perspectives.

## Supplementary Materials

MAF.zip is also available via SOURCEFORGE

Home page: CCSR

GEO: GSE207522, GSE275002.

## Supporting information

Supplemetary MAF file

