## Supplementary material for "Evaluating the effectiveness of various small RNA alignment techniques in transcriptomic analysis by examining different sources of variability through a multi-alignment approach": Supplemetary MAF file: Figure S1.pdf

### Figure S1. Venn diagrams of the differential sets in DE-1 and DE-2.

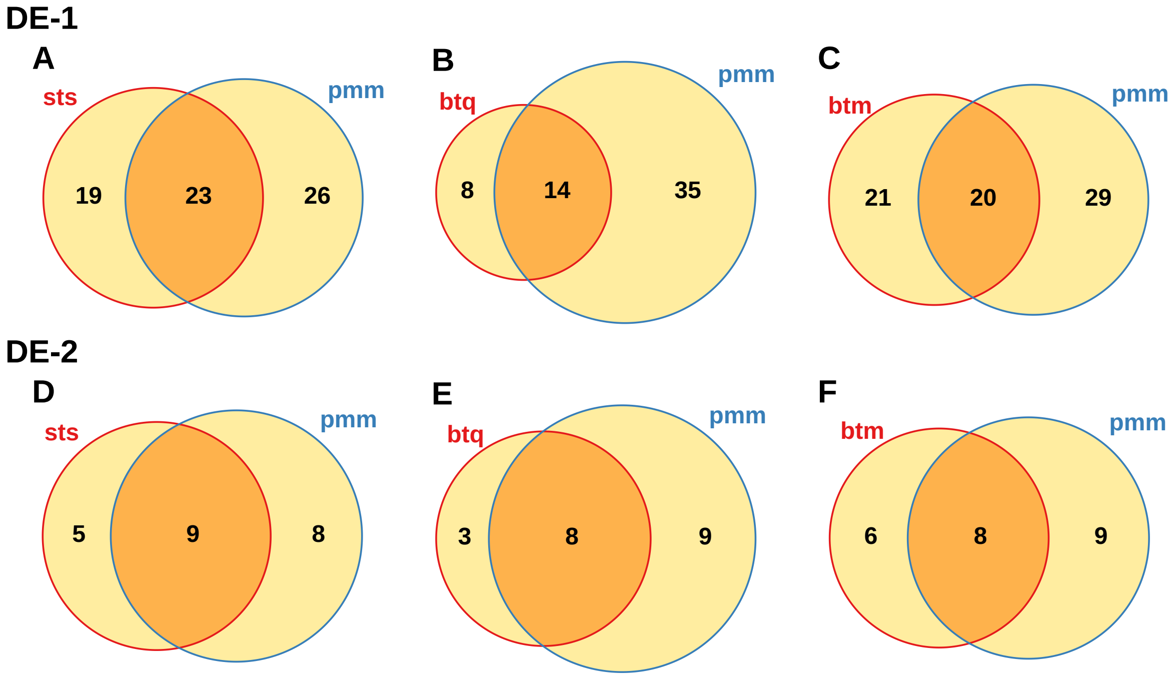


Panel 'DE-1' is showing in A to C the overlap between the three different alignment and quantification approaches and their significant microRNA candidates for the first differential analysis. The nomenclature is already described in Table 2 in the right column.
In panel 'DE-2' the same is done in D to E for the second differential analysis and again on microRNAs.
The two differential analyses differentiate by absolute count number and a more pronounced higher set number for the premature microRNAs (pmm).
Acronyms sts, btq, btm, pmm explainded in supplementary Table S1.
