## Supplementary material for "Evaluating the effectiveness of various small RNA alignment techniques in transcriptomic analysis by examining different sources of variability through a multi-alignment approach": Supplemetary MAF file: Figure S2.pdf

**Figure S2.**
**Overview on all significant candidates.**

Further examples for the ambiguity of significance
and robustness of candidates.

Mature microRNA / tRNA: sts, stm, btm, btq, bbm,
piRNA: pi,
premature microRNA: pmm,
1/2: differential set 1 (g1-g2) or 2 (g3-g4),
all Homo sapiens.

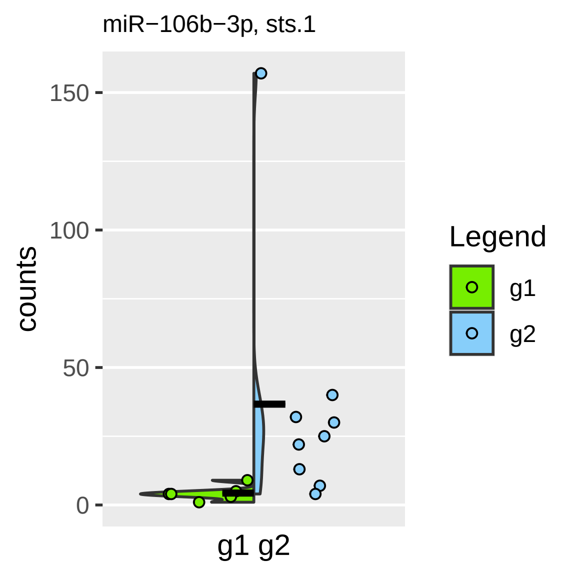

miR−106b−3p, sts.1
counts
150
100
50
0
g1 g2
Legend
g1
g2

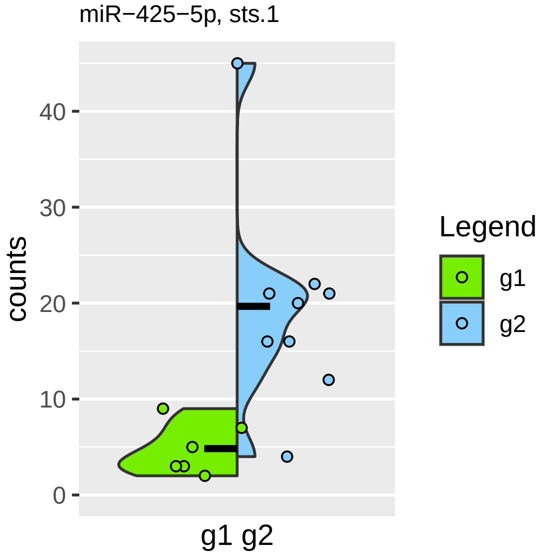

miR−425−5p, sts.1
counts
40
30
20
10
0
g1 g2
Legend
g1
g2

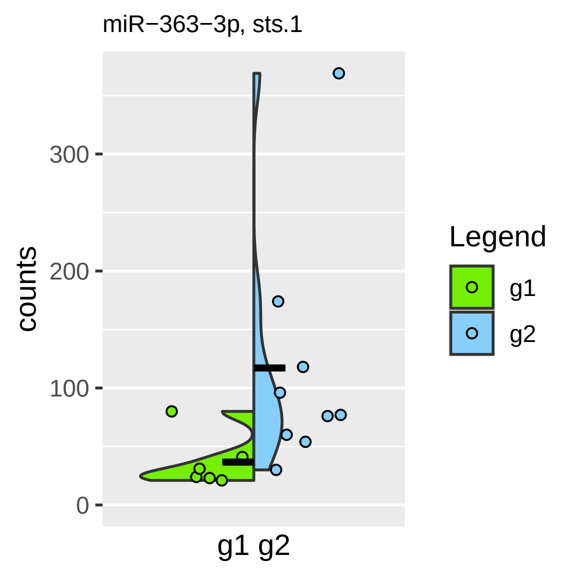

miR−363−3p, sts.1
counts
300
200
100
0
g1 g2
Legend
g1
g2

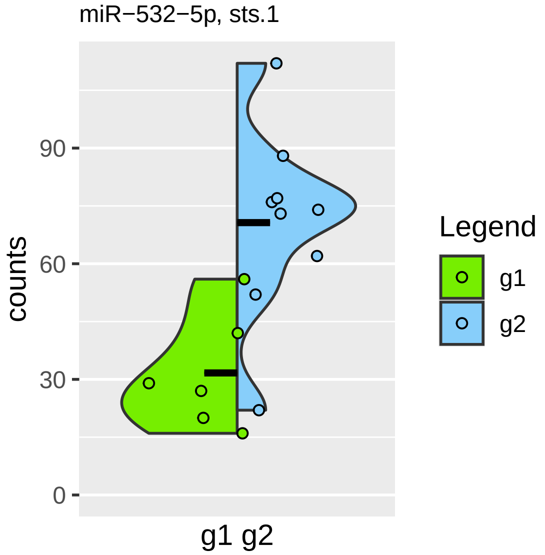
miR−532−5p, sts.1
counts
90
60
30
0
g1 g2
Legend
g1
g2

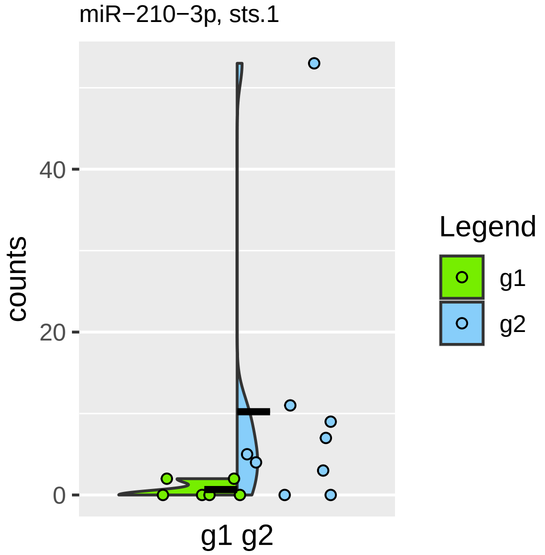
miR−210−3p, sts.1
counts
40
20
0
g1 g2
Legend
g1
g2

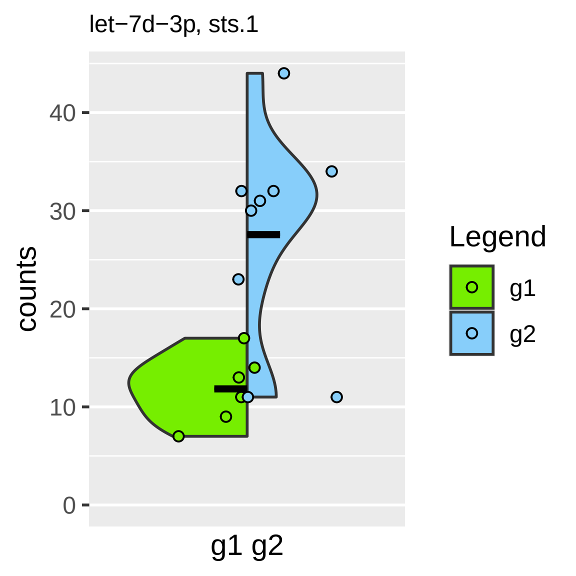
let−7d−3p, sts.1
counts
40
30
20
10
0
g1 g2
Legend
g1
g2

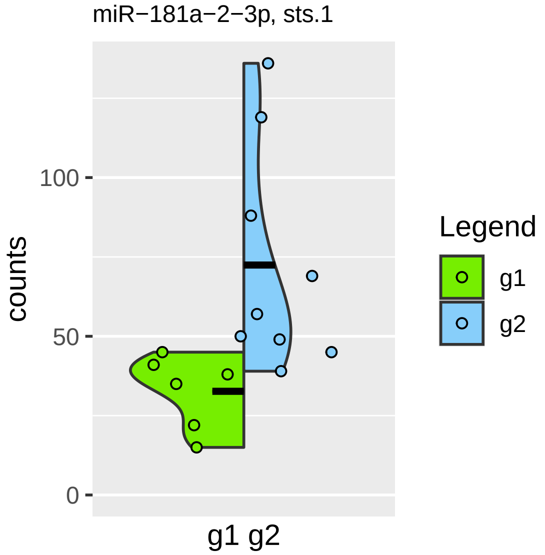
miR−181a−2−3p, sts.1
counts
100
50
0
g1 g2
Legend
g1
g2

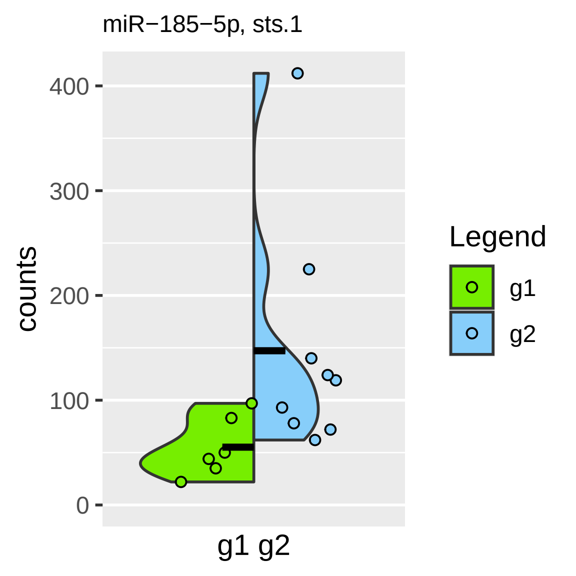
miR−185−5p, sts.1
400
300
200
100
0
counts
g1 g2
Legend
g1
g2

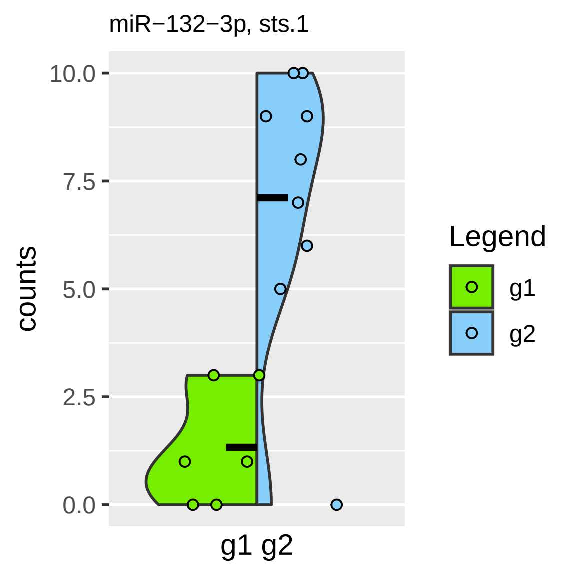
miR−132−3p, sts.1
10.0
7.5
5.0
2.5
0.0
counts
g1 g2
Legend
g1
g2

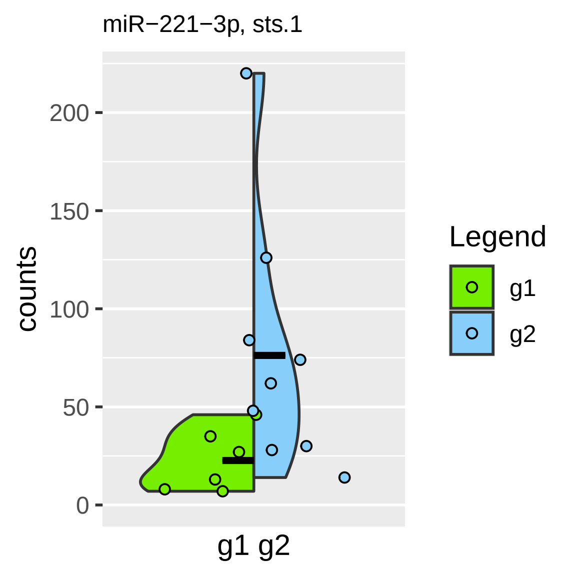
miR−221−3p, sts.1
counts
200
150
100
50
0
g1 g2
Legend
g1
g2

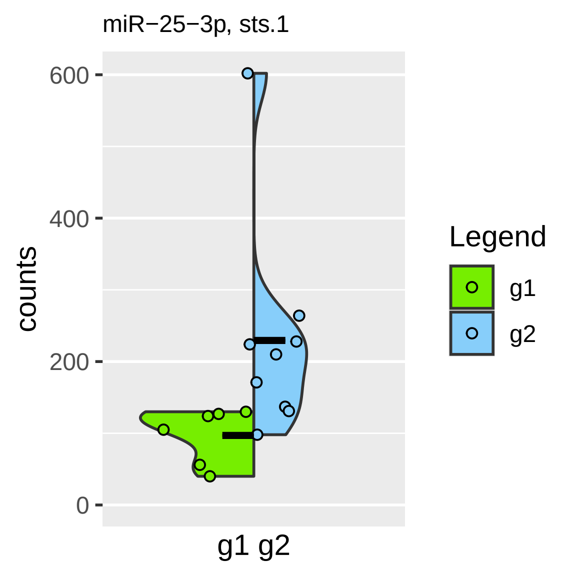

miR−25−3p, sts.1
600
400
200
0
counts
g1 g2
Legend
g1
g2

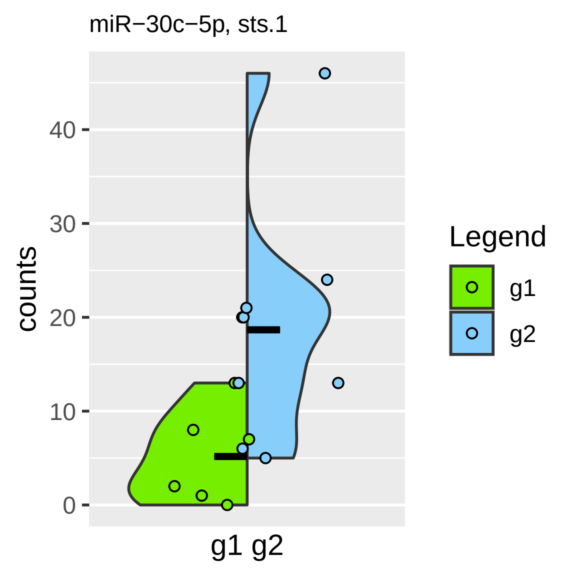

miR−30c−5p, sts.1
counts
0
10
20
30
40
g1 g2
Legend
g1
g2

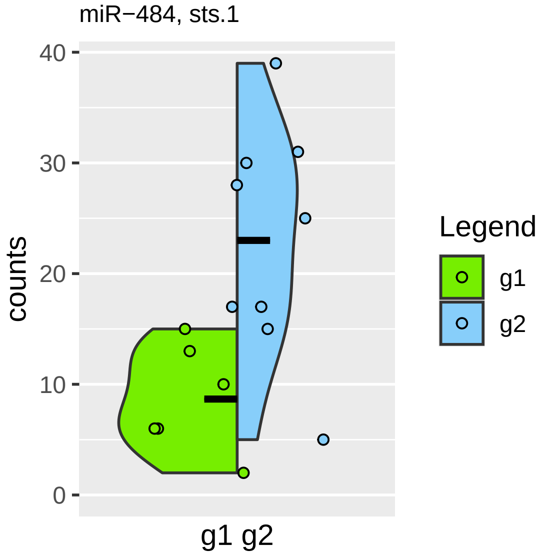

miR−484, sts.1
counts
40
30
20
10
0
g1 g2
Legend
g1
g2

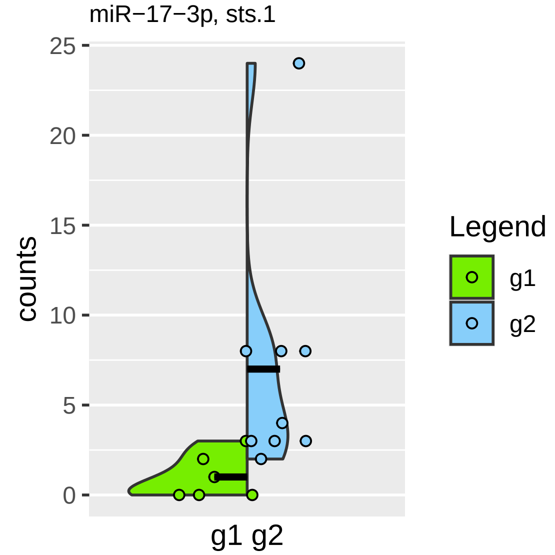
miR−17−3p, sts.1
counts
25
20
15
10
5
0
g1 g2
Legend
g1
g2

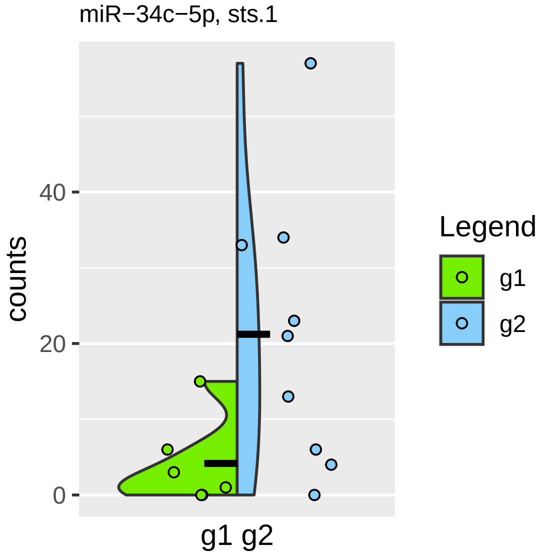
miR−34c−5p, sts.1
counts
0
20
40
g1 g2
Legend
g1
g2

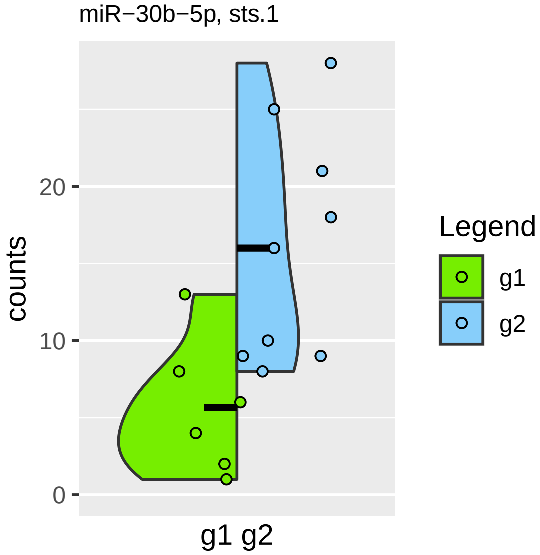

miR−30b−5p, sts.1
counts
20
10
0
g1 g2
Legend
g1
g2

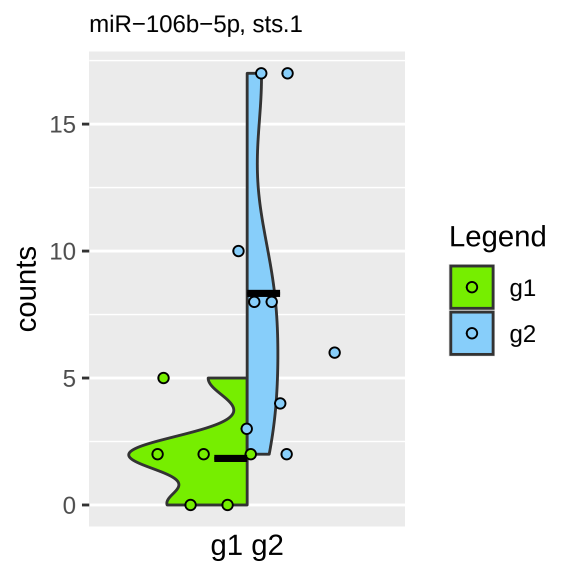

miR−106b−5p, sts.1
counts
15
10
5
0
g1 g2
Legend
g1
g2

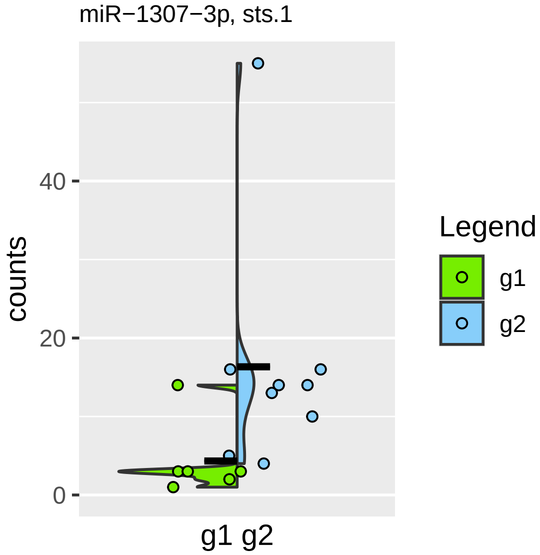

miR−1307−3p, sts.1
counts
0
20
40
g1 g2
Legend
g1
g2

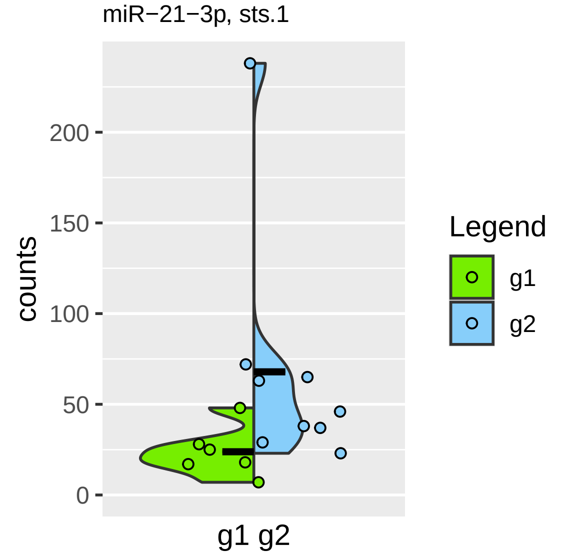
miR−21−3p, sts.1
counts
200
150
100
50
0
g1 g2
Legend
g1
g2

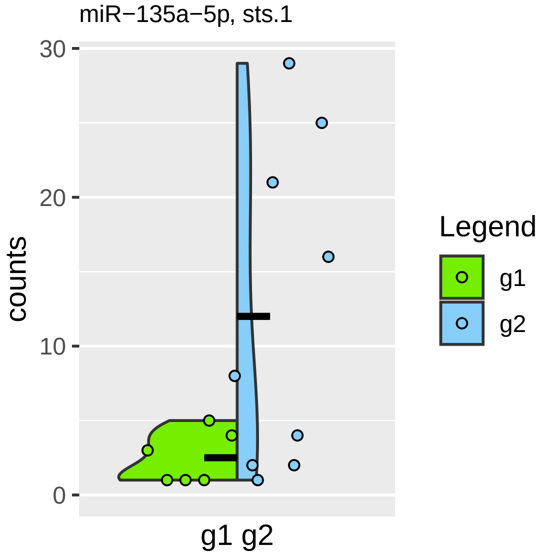

miR−135a−5p, sts.1
30
20
10
0
counts
g1 g2
Legend
g1
g2

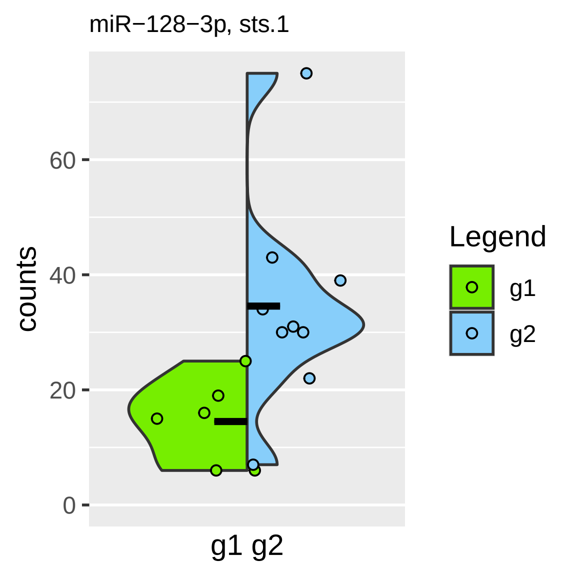

miR−128−3p, sts.1
counts
60
40
20
0
g1 g2
Legend
g1
g2

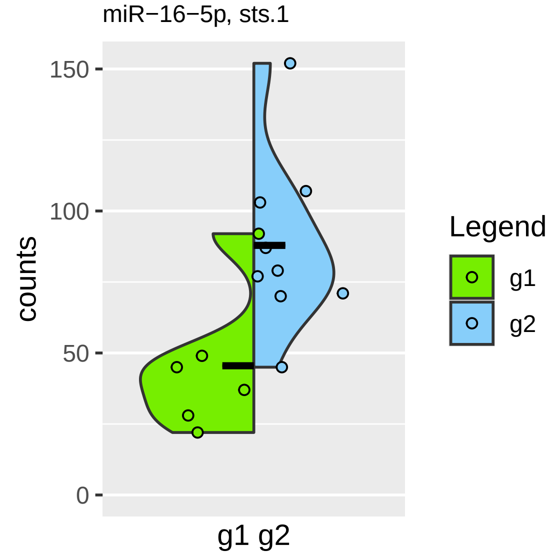
miR−16−5p, sts.1
150
100
50
0
counts
g1 g2
Legend
g1
g2

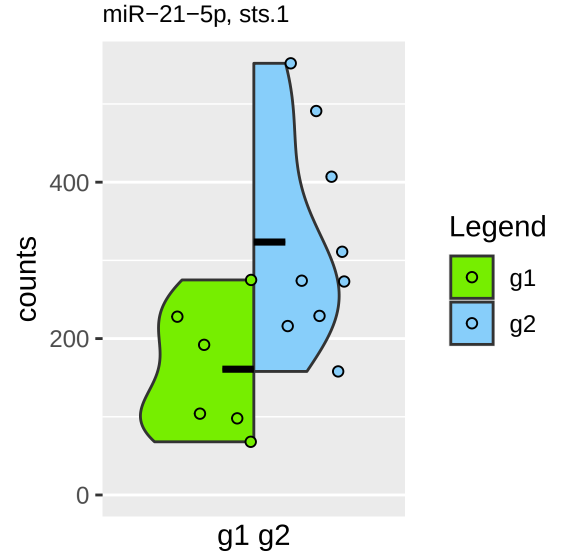

miR−21−5p, sts.1
counts
400
200
0
g1 g2
Legend
g1
g2

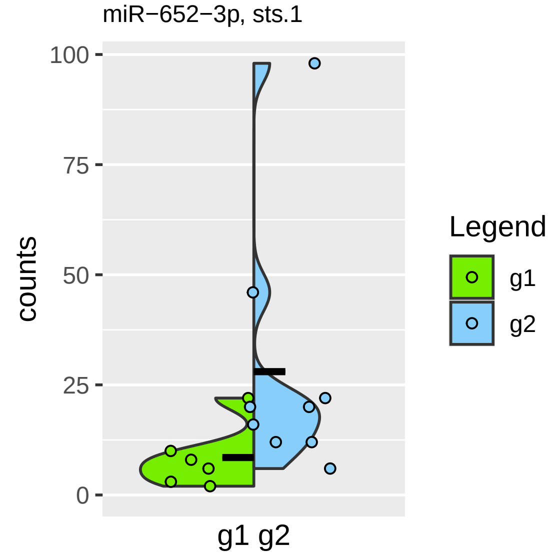

miR−652−3p, sts.1
100
75
50
25
0
counts
g1 g2
Legend
g1
g2

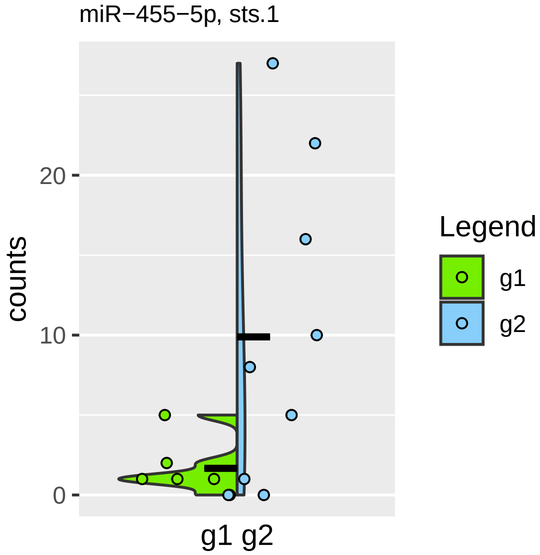
miR−455−5p, sts.1
counts
20
10
0
g1 g2
Legend
g1
g2

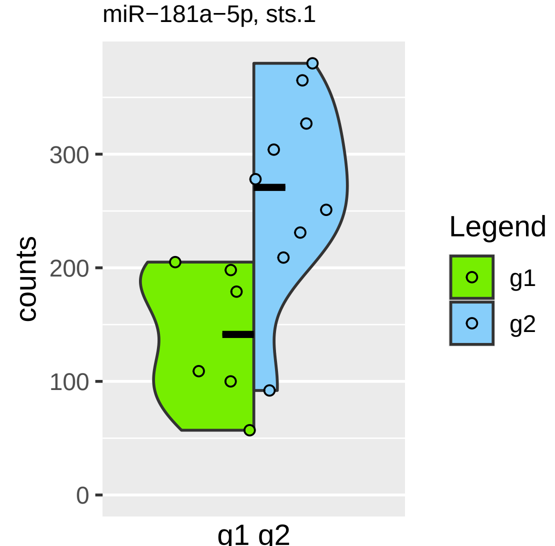
miR−181a−5p, sts.1
counts
300
200
100
0
g1 g2
Legend
g1
g2

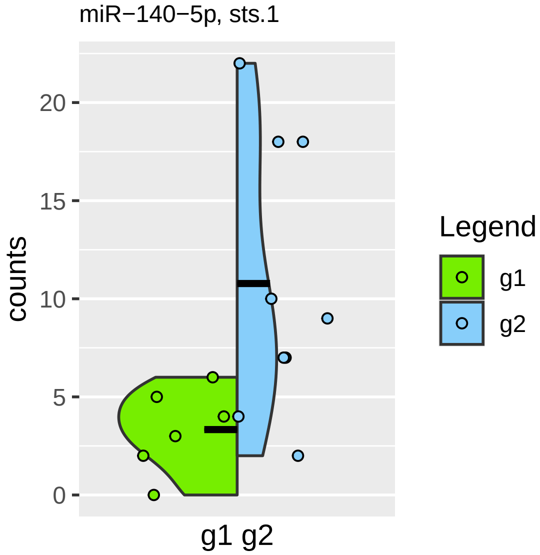

miR−140−5p, sts.1
counts
20
15
10
5
0
g1 g2
Legend
g1
g2

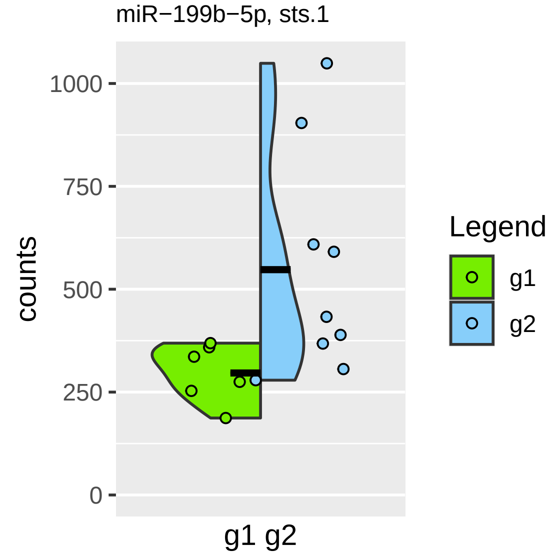

miR−199b−5p, sts.1
counts
1000
750
500
250
0
g1 g2
Legend
g1
g2

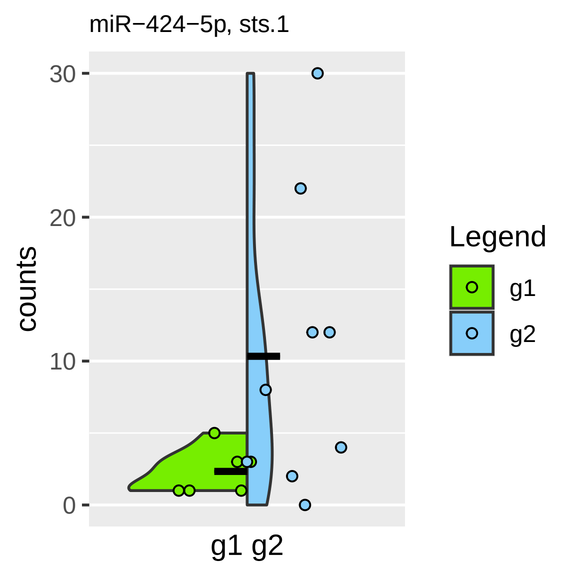
miR−424−5p, sts.1
counts
30
20
10
0
g1 g2
Legend
g1
g2

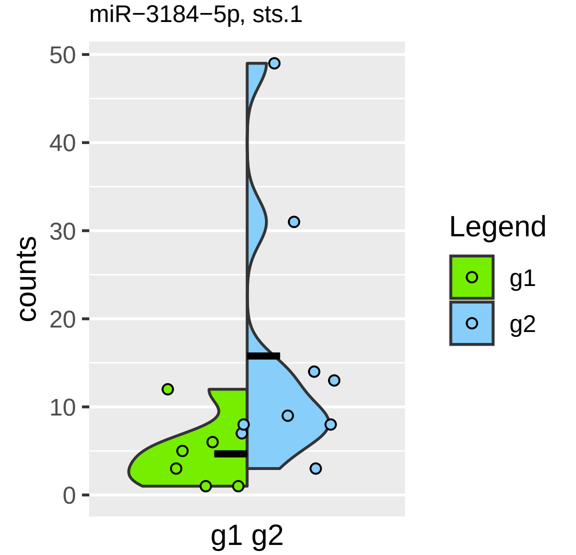
miR−3184−5p, sts.1
counts
50
40
30
20
10
0
g1 g2
Legend
g1
g2

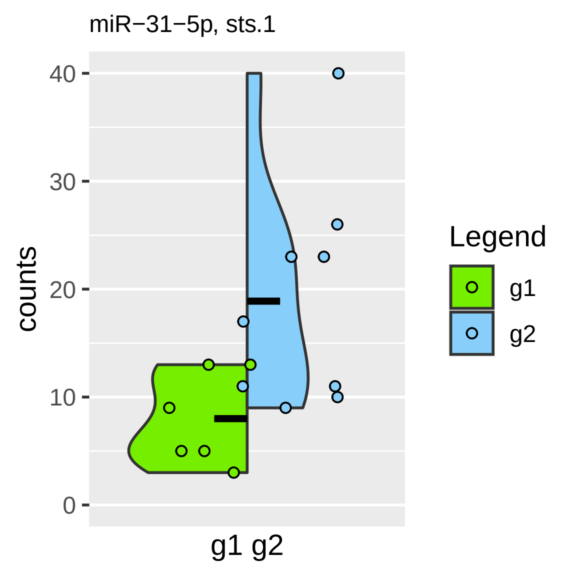

miR−31−5p, sts.1
counts
40
30
20
10
0
g1 g2
Legend
g1
g2

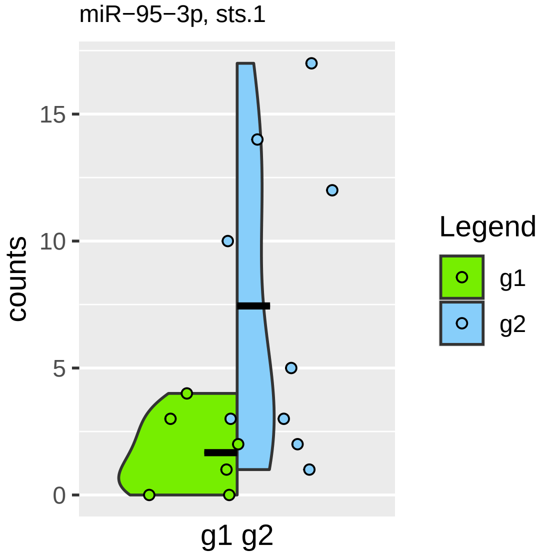

miR−95−3p, sts.1
counts
15
10
5
0
g1 g2
Legend
g1
g2

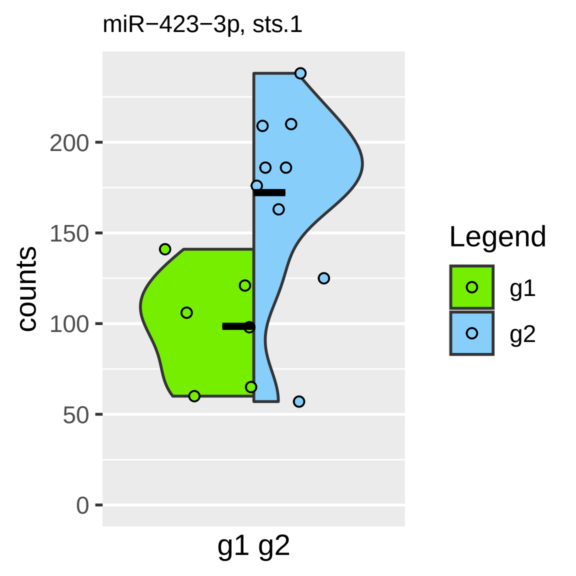

miR−423−3p, sts.1
counts
200
150
100
50
0
g1 g2
Legend
g1
g2

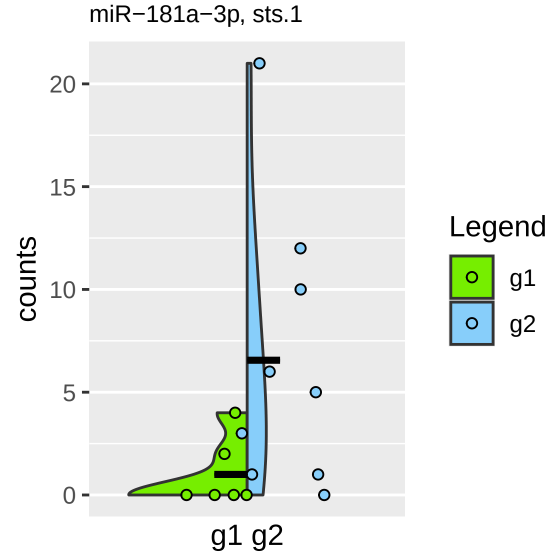
miR−181a−3p, sts.1
counts
20
15
10
5
0
g1 g2
Legend
g1
g2

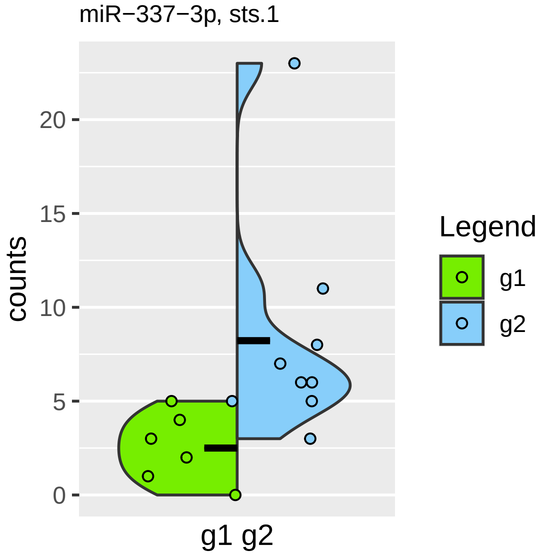

miR−337−3p, sts.1
counts
0
5
10
15
20
g1 g2
Legend
g1
g2

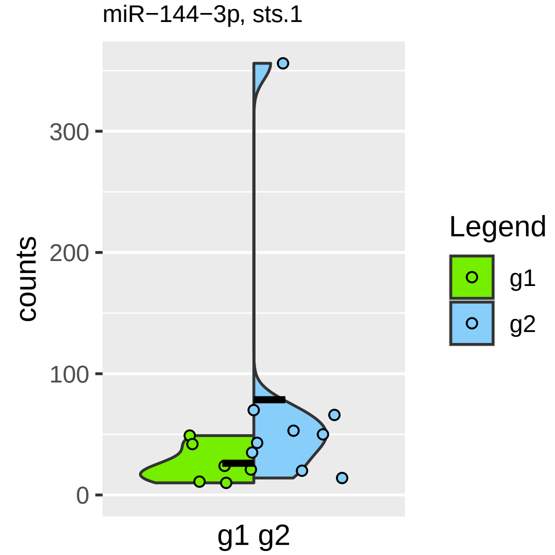
miR−144−3p, sts.1
counts
300
200
100
0
g1 g2
Legend
g1
g2

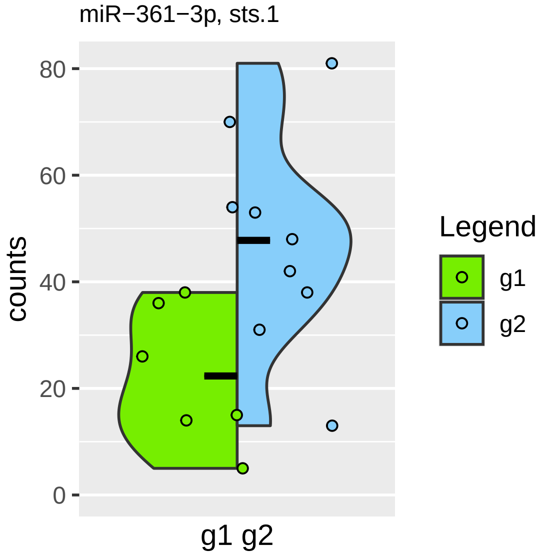

miR−361−3p, sts.1
80
60
40
20
0
counts
g1 g2
Legend
g1
g2

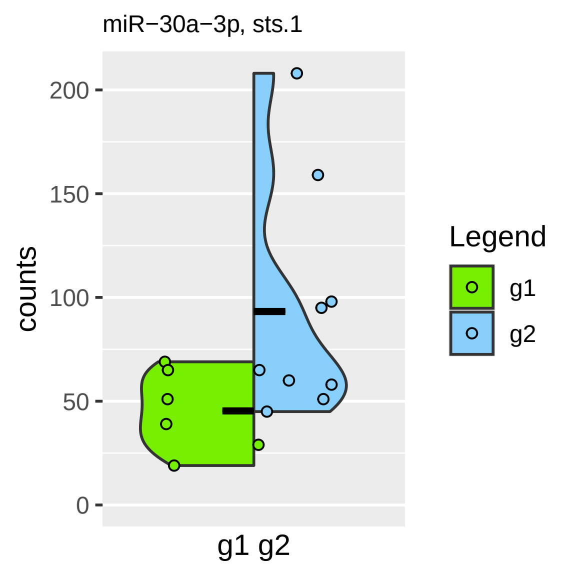

miR−30a−3p, sts.1
counts
200
150
100
50
0
g1 g2
Legend
g1
g2

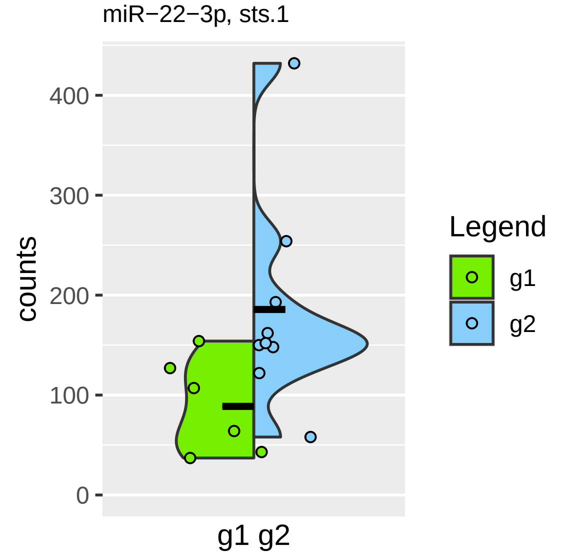

miR−22−3p, sts.1
counts
400
300
200
100
0
g1 g2
Legend
g1
g2

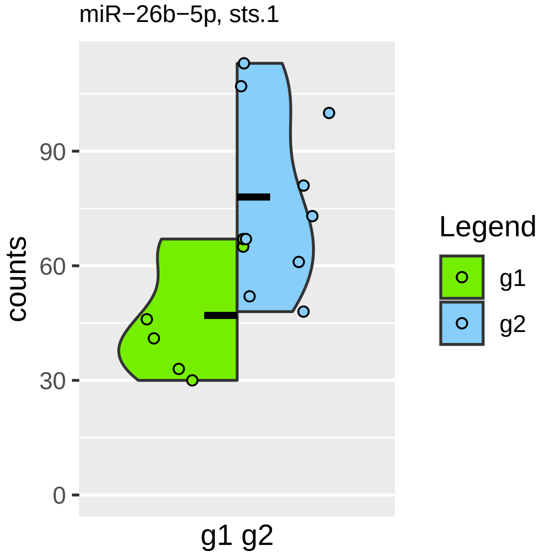

miR−26b−5p, sts.1
counts
90
60
30
0
g1 g2
Legend
g1
g2

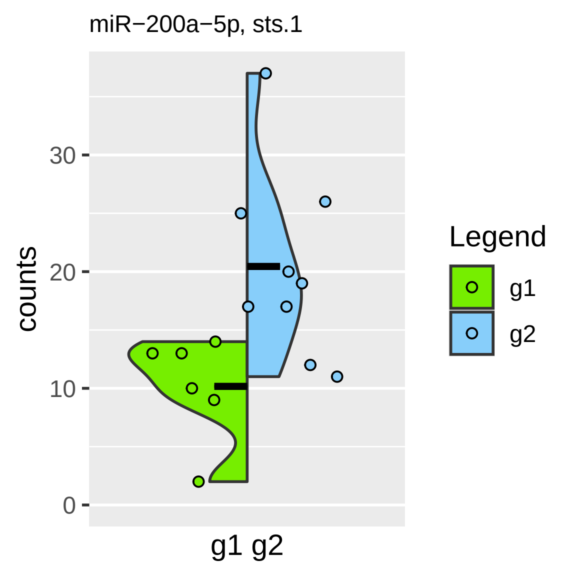
miR−200a−5p, sts.1
counts
30
20
10
0
g1 g2
Legend
g1
g2

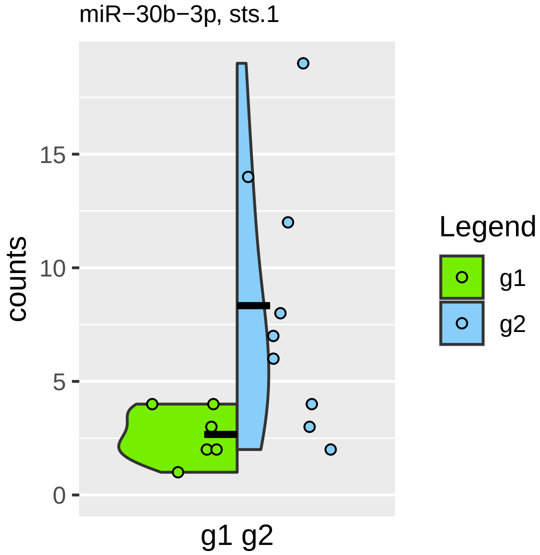

miR−30b−3p, sts.1
counts
15
10
5
0
g1 g2
Legend
g1
g2

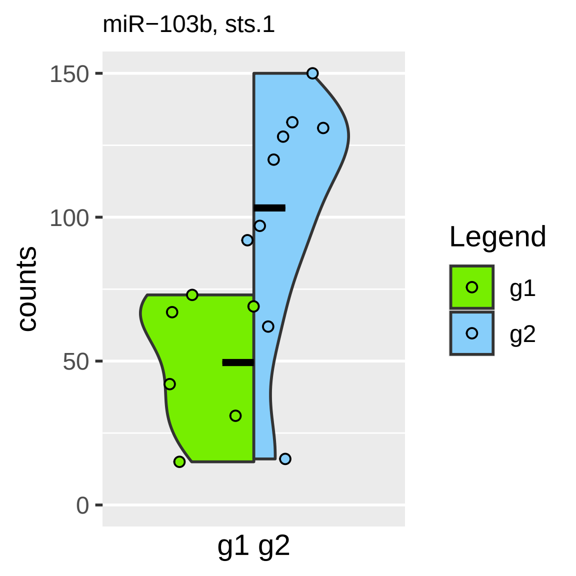

miR−103b, sts.1
150
100
50
0
counts
g1 g2
Legend
g1
g2

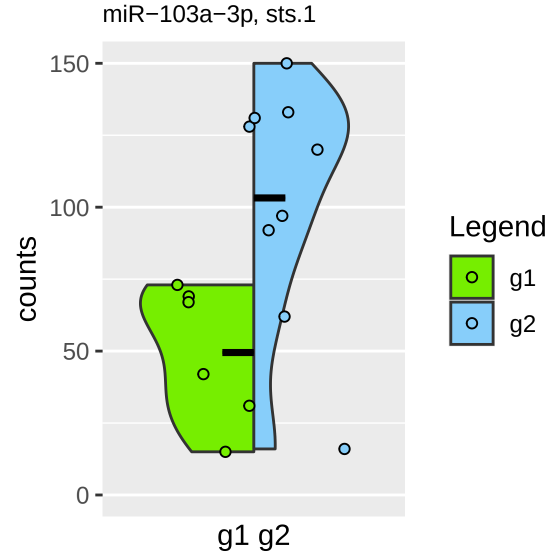

miR−103a−3p, sts.1
150
100
50
0
counts
g1 g2
Legend
g1
g2

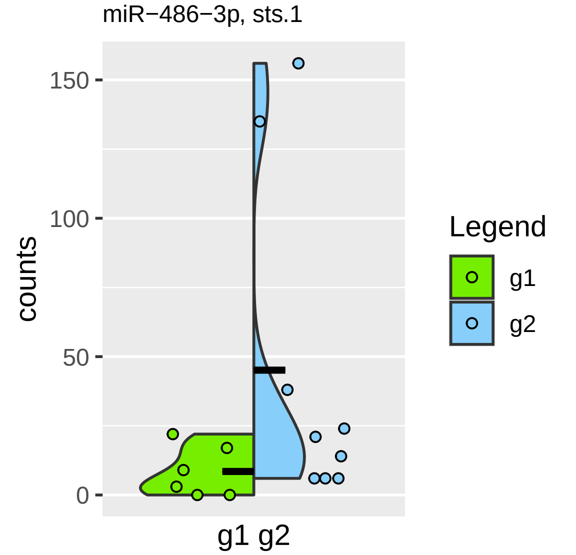

miR−486−3p, sts.1
150
100
50
0
counts
g1 g2
Legend
g1
g2

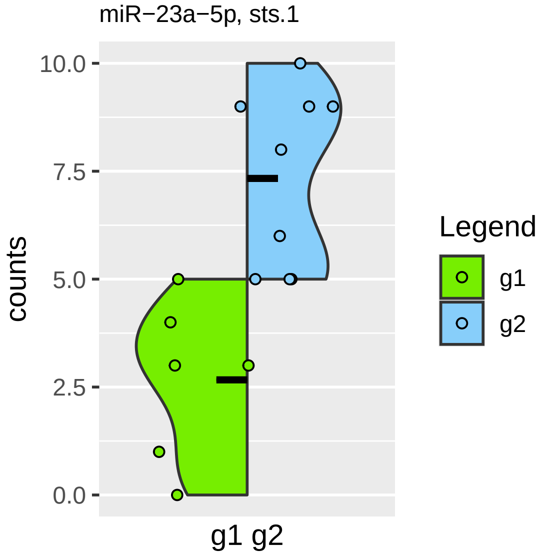

miR−23a−5p, sts.1
10.0
7.5
5.0
2.5
0.0
counts
g1 g2
Legend
g1
g2

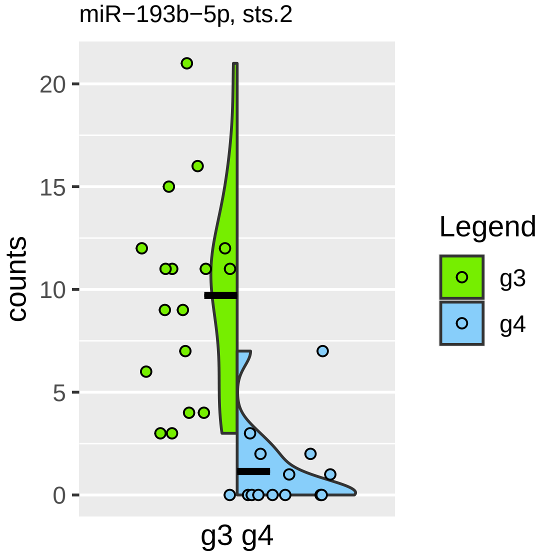

miR−193b−5p, sts.2
counts
20
15
10
5
0
g3 g4
Legend
g3
g4

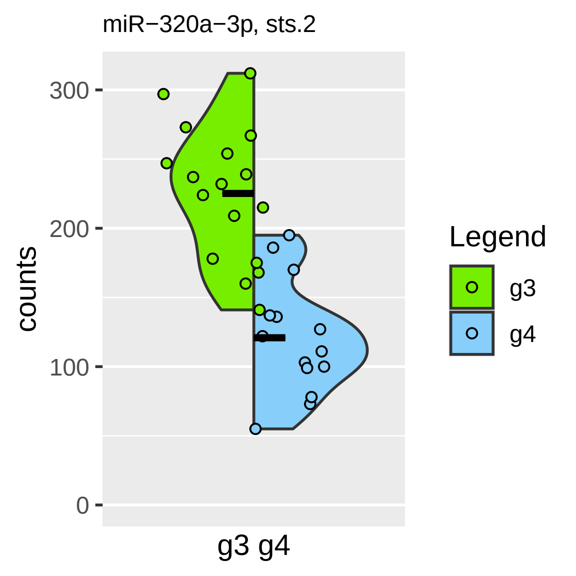

miR−320a−3p, sts.2
counts
300
200
100
0
g3 g4
Legend
g3
g4

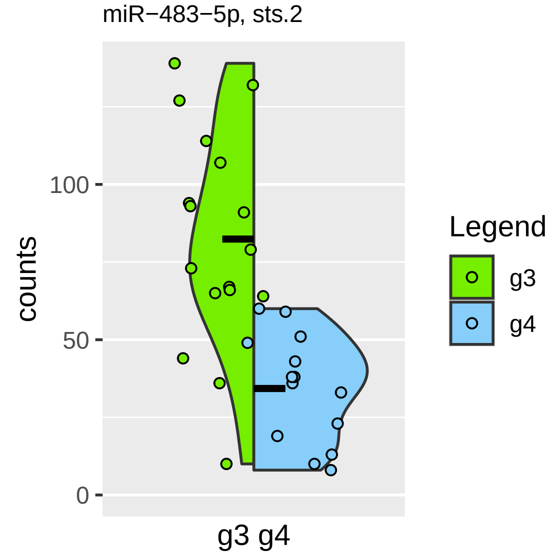

miR−483−5p, sts.2
counts
100
50
0
g3 g4
Legend
g3
g4

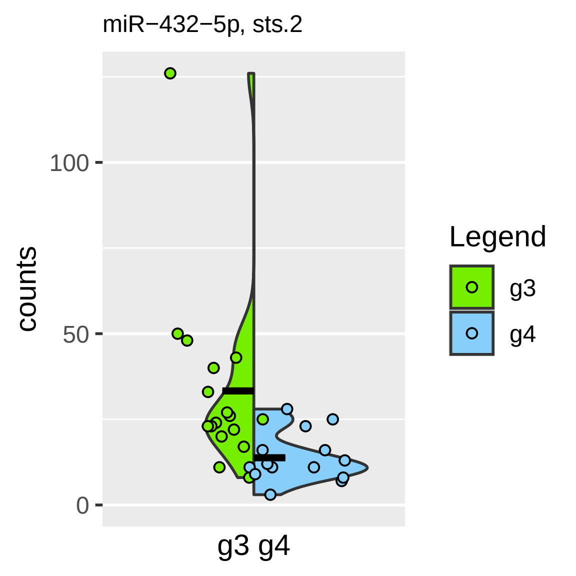

miR−432−5p, sts.2
counts
100
50
0
g3 g4
Legend
g3
g4

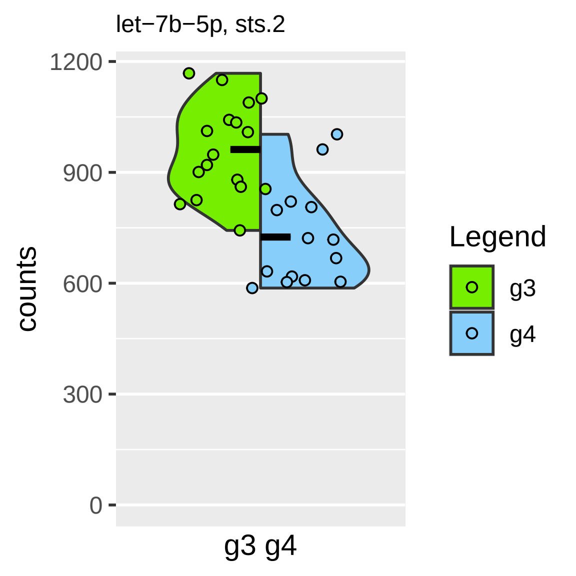
let−7b−5p, sts.2
1200
900
600
300
0
counts
g3 g4
Legend
g3
g4

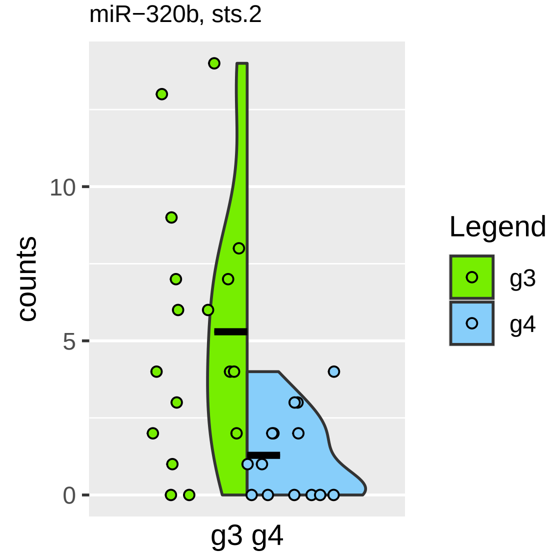

miR−320b, sts.2
counts
10
5
0
g3 g4
Legend
g3
g4

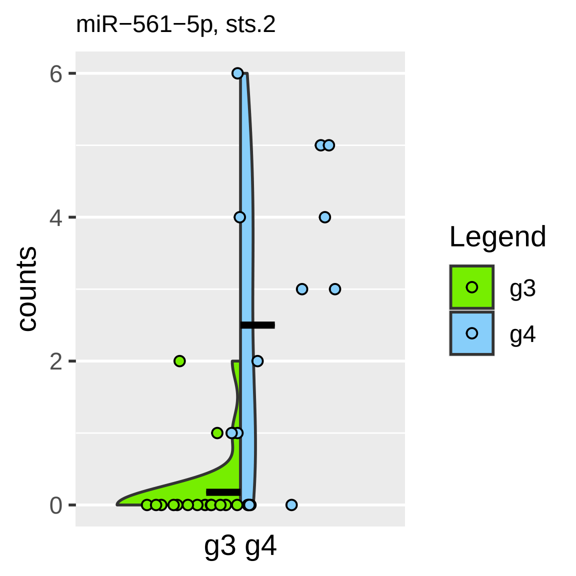

miR−561−5p, sts.2
counts
6
4
2
0
g3 g4
Legend
g3
g4

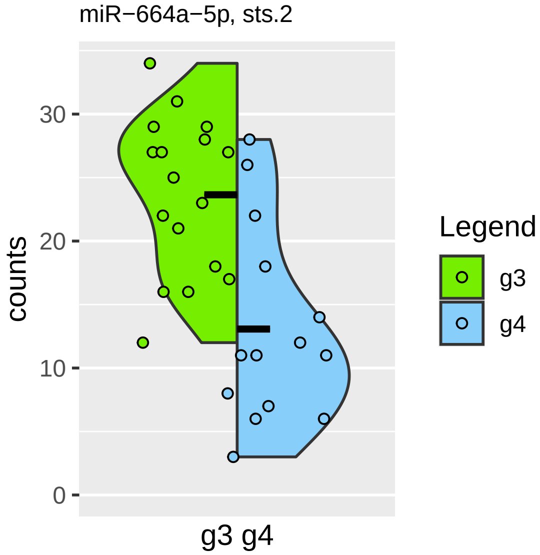

miR−664a−5p, sts.2
counts
30
20
10
0
g3 g4
Legend
g3
g4

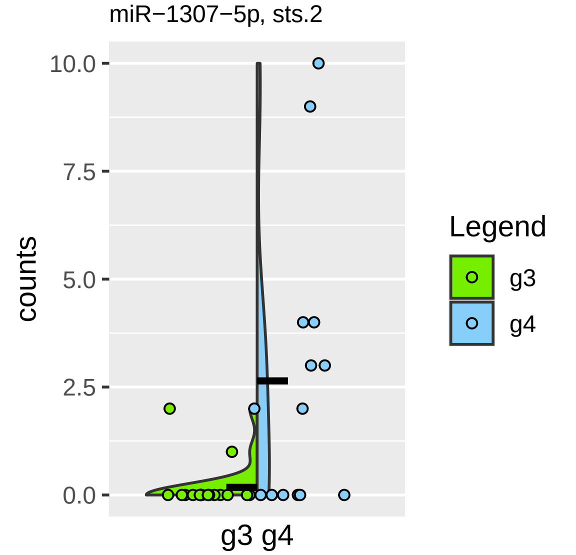

miR−1307−5p, sts.2
10.0
7.5
5.0
2.5
0.0
counts
g3 g4
Legend
g3
g4

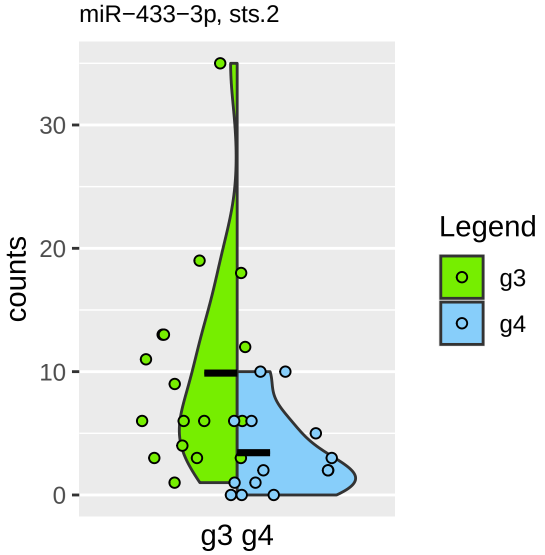

miR−433−3p, sts.2
counts
0
10
20
30
g3 g4
Legend
g3
g4

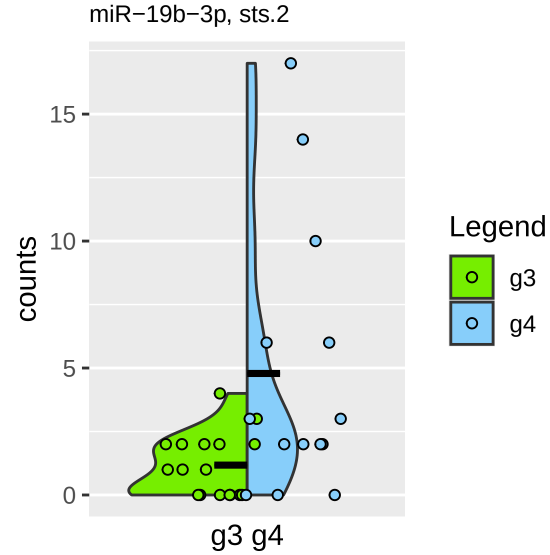

miR−19b−3p, sts.2
counts
15
10
5
0
g3 g4
Legend
g3
g4

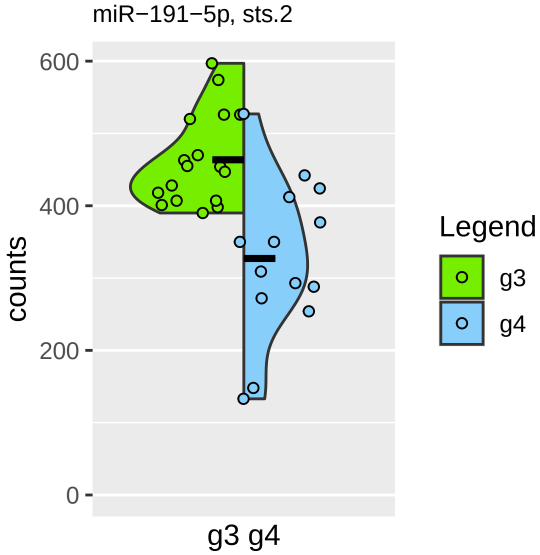

miR−191−5p, sts.2
600
400
200
0
counts
g3 g4
Legend
g3
g4

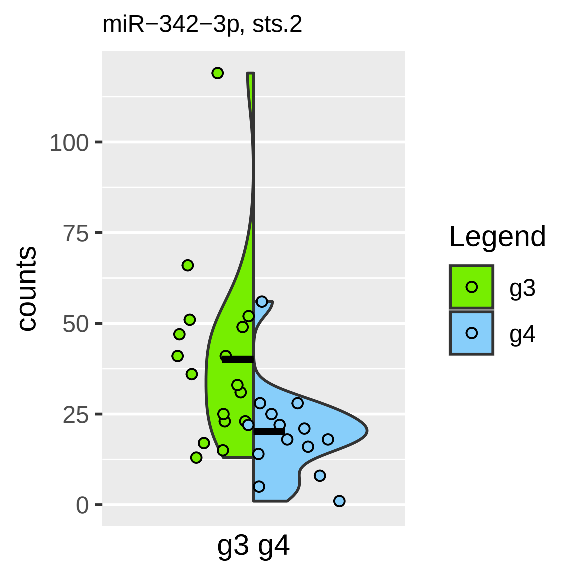

miR−342−3p, sts.2
counts
100
75
50
25
0
g3 g4
Legend
g3
g4

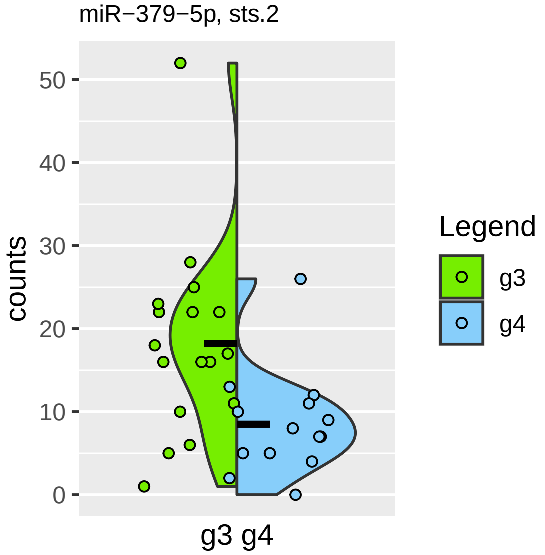
miR−379−5p, sts.2
counts
50
40
30
20
10
0
g3 g4
Legend
g3
g4

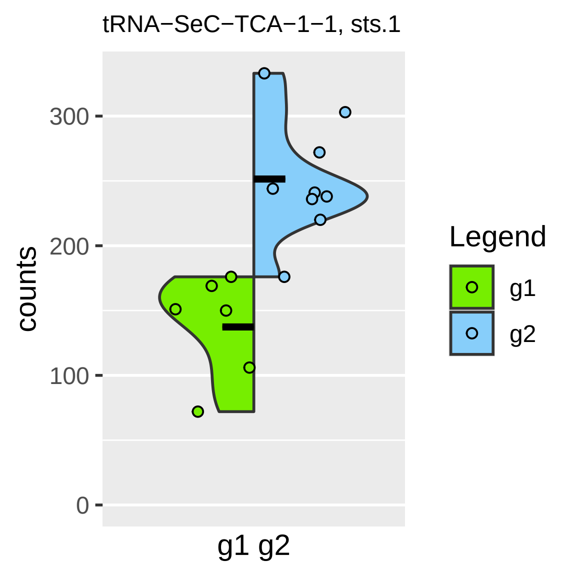
tRNA−SeC−TCA−1−1, sts.1
counts
300
200
100
0
g1 g2
Legend
g1
g2

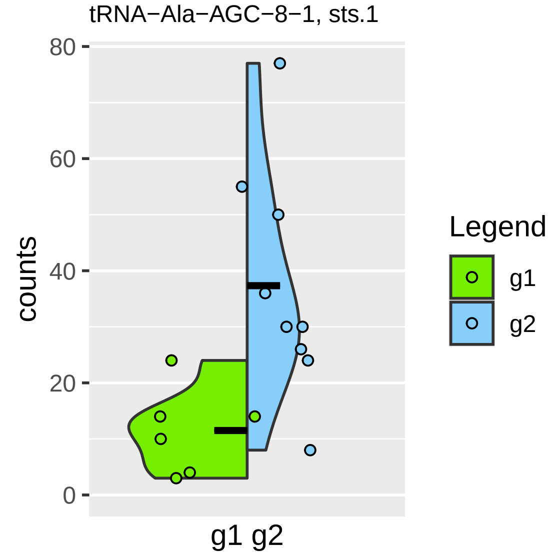
tRNA−Ala−AGC−8−1, sts.1
counts
0
20
40
60
80
g1 g2
Legend
g1
g2

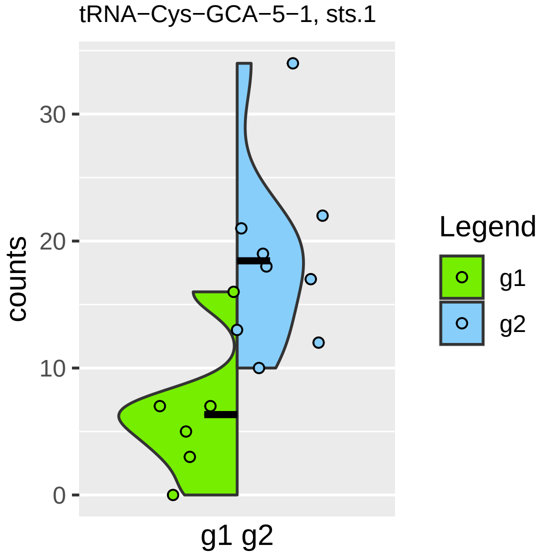

tRNA−Cys−GCA−5−1, sts.1
counts
0
10
20
30
g1 g2
Legend
g1
g2

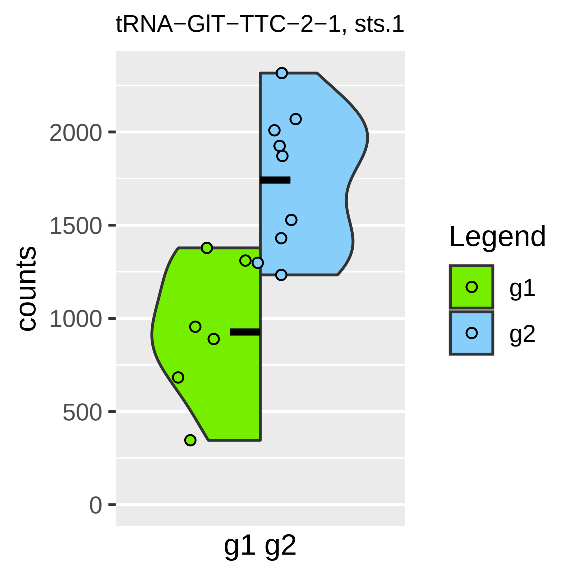
tRNA−GlT−TTC−2−1, sts.1
counts
2000
1500
1000
500
0
g1 g2
Legend
g1
g2

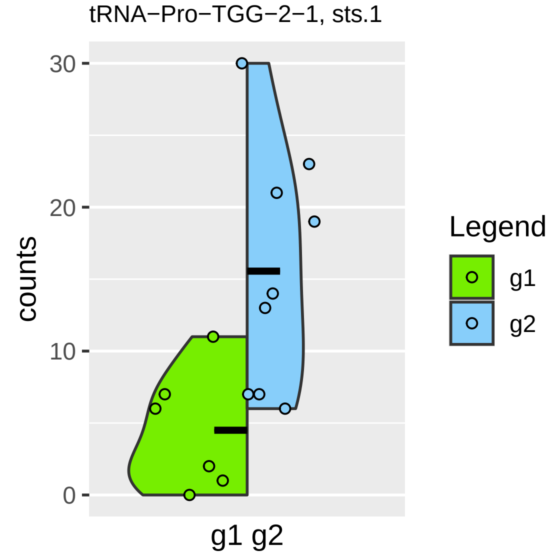
tRNA−Pro−TGG−2−1, sts.1
30
20
10
0
counts
g1 g2
Legend
g1
g2

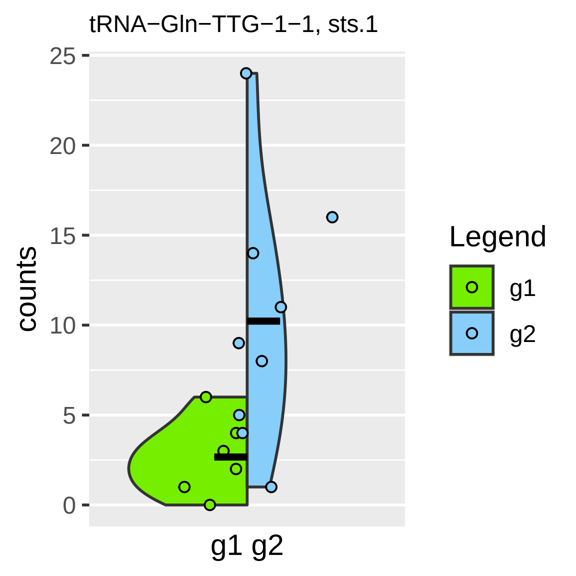
tRNA−Gln−TTG−1−1, sts.1
counts
25
20
15
10
5
0
g1 g2
Legend
g1
g2

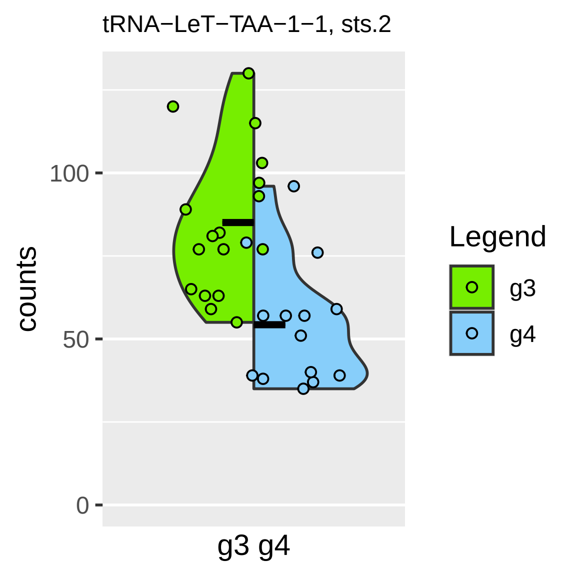
tRNA−LeT−TAA−1−1, sts.2
counts
100
50
0
g3 g4
Legend
g3
g4

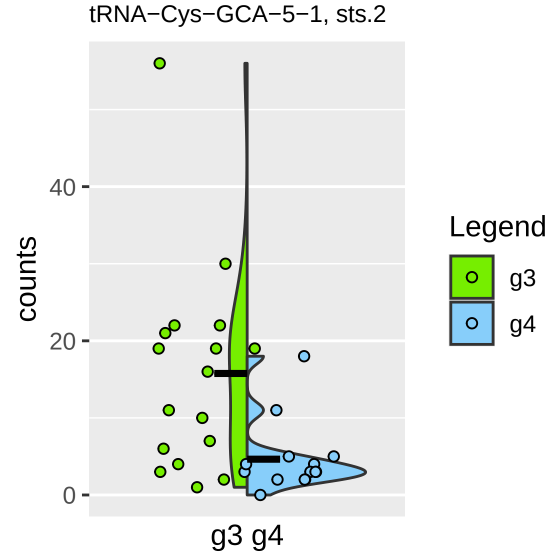

tRNA−Cys−GCA−5−1, sts.2
counts
40
20
0
g3 g4
Legend
g3
g4

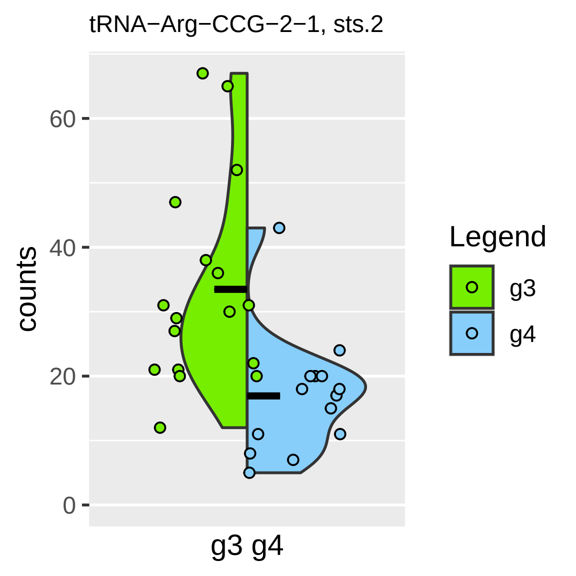

tRNA−Arg−CCG−2−1, sts.2
counts
60
40
20
0
g3 g4
Legend
g3
g4

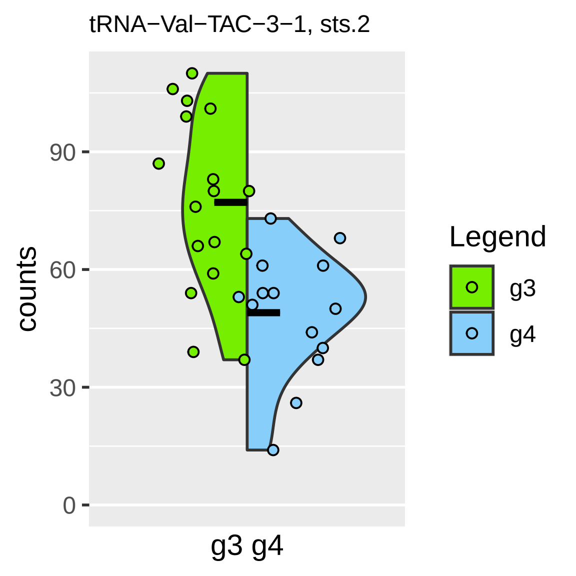

tRNA−Val−TAC−3−1, sts.2
counts
0
30
60
90
g3 g4
Legend
g3
g4

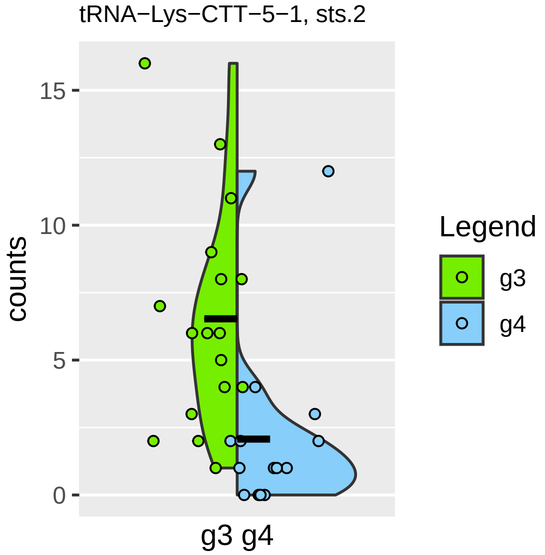
tRNA−Lys−CTT−5−1, sts.2
counts
15
10
5
0
g3 g4
Legend
g3
g4

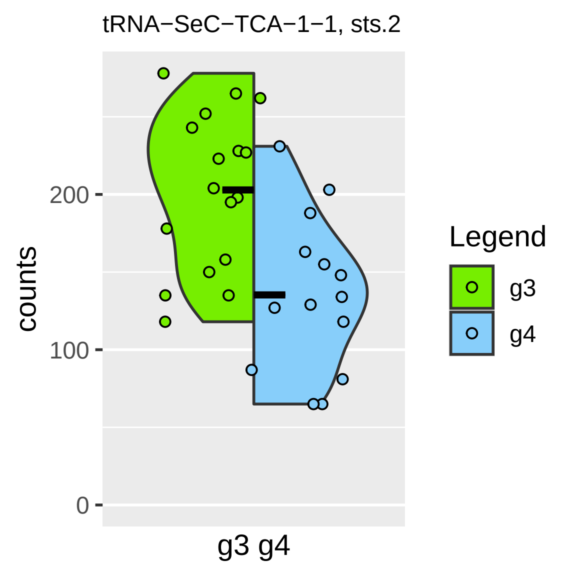

tRNA−SeC−TCA−1−1, sts.2
counts
0
100
200
g3 g4
Legend
g3
g4

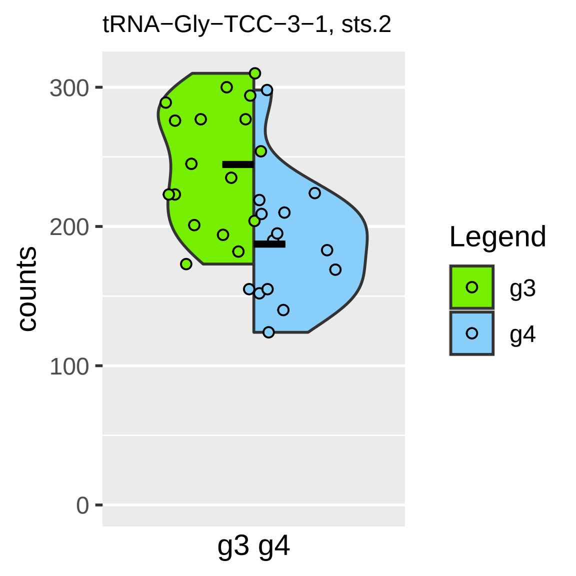
tRNA−Gly−TCC−3−1, sts.2
counts
300
200
100
0
g3 g4
Legend
g3
g4

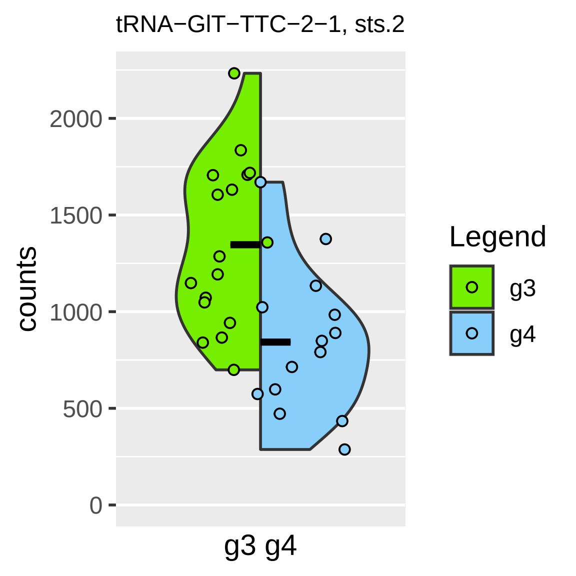

tRNA−GlT−TTC−2−1, sts.2
counts
2000
1500
1000
500
0
g3 g4
Legend
g3
g4

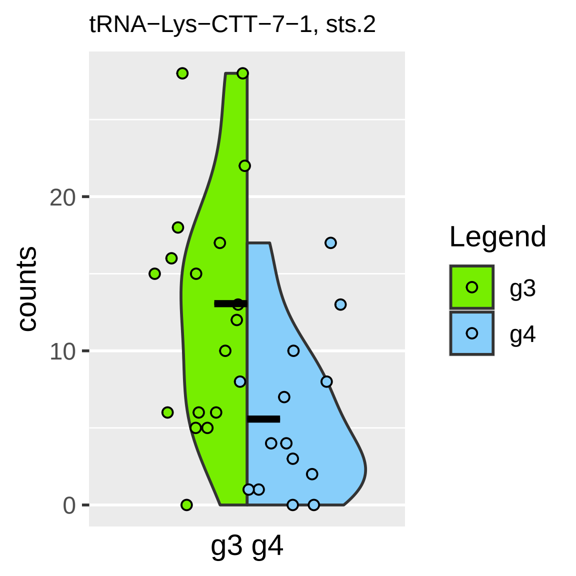

tRNA−Lys−CTT−7−1, sts.2
counts
20
10
0
g3 g4
Legend
g3
g4

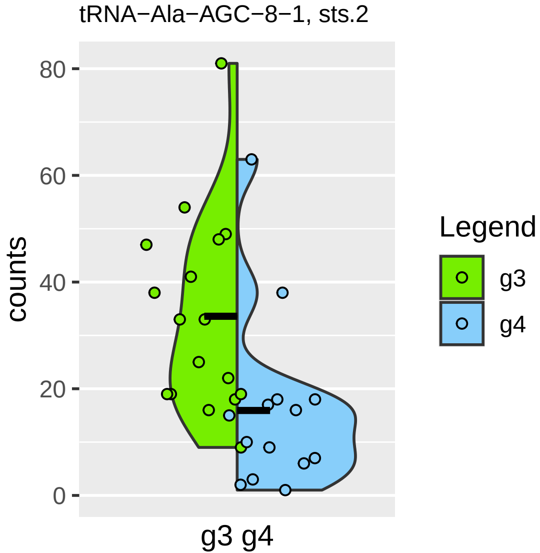

tRNA−Ala−AGC−8−1, sts.2
counts
80
60
40
20
0
g3 g4
Legend
g3
g4

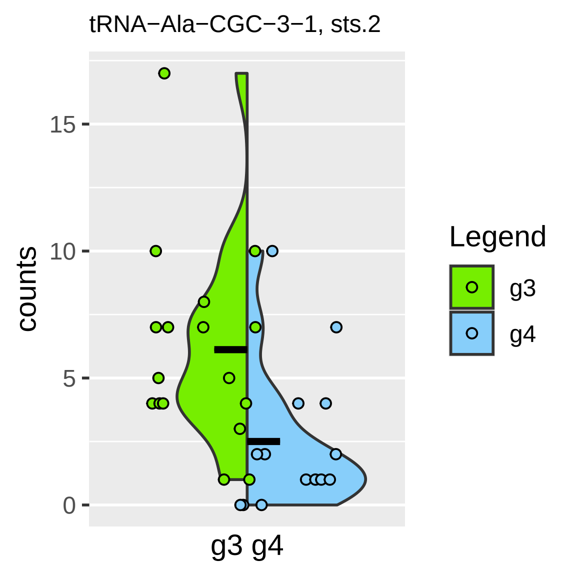

tRNA−Ala−CGC−3−1, sts.2
counts
15
10
5
0
g3 g4
Legend
g3
g4

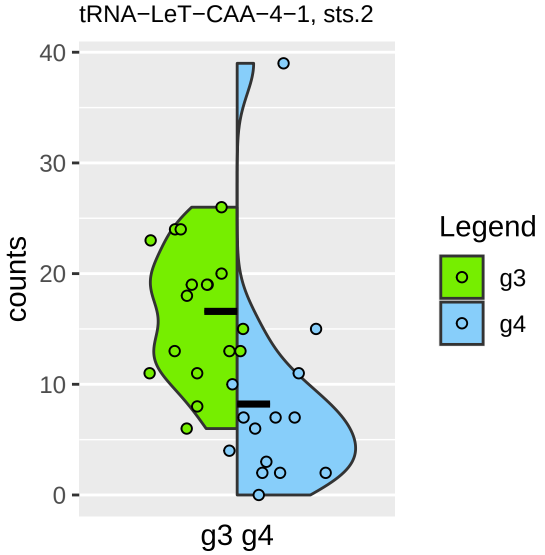
tRNA−LeT−CAA−4−1, sts.2
counts
40
30
20
10
0
g3 g4
Legend
g3
g4

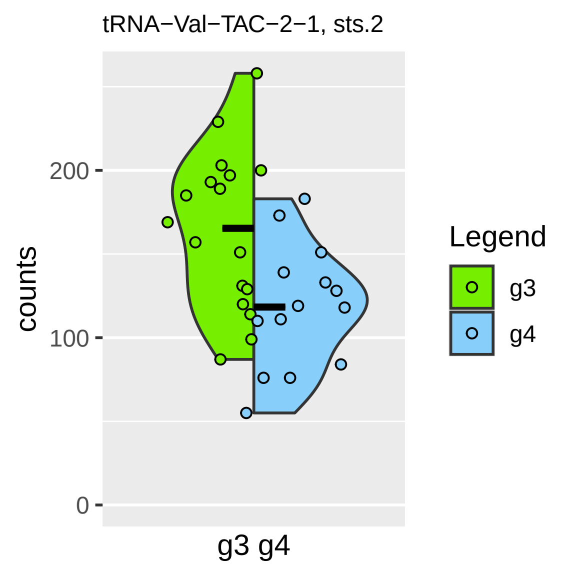
tRNA−Val−TAC−2−1, sts.2
counts
200
100
0
g3 g4
Legend
g3
g4

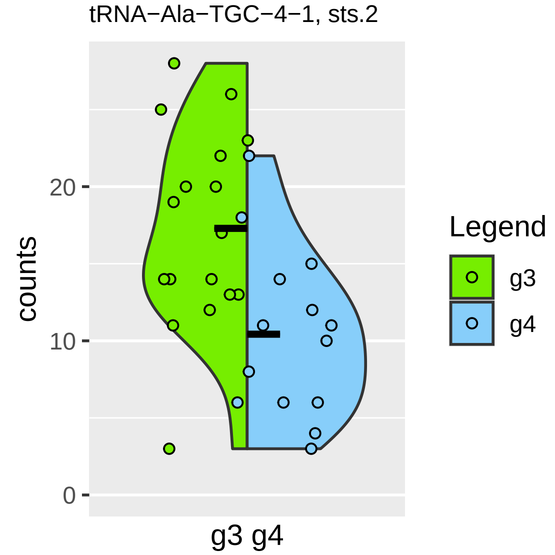
tRNA−Ala−TGC−4−1, sts.2
counts
0
10
20
g3 g4
Legend
g3
g4

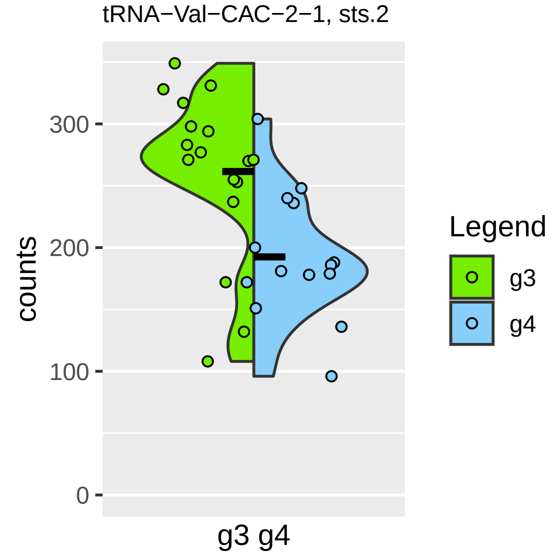

tRNA−Val−CAC−2−1, sts.2
counts
300
200
100
0
g3 g4
Legend
g3
g4

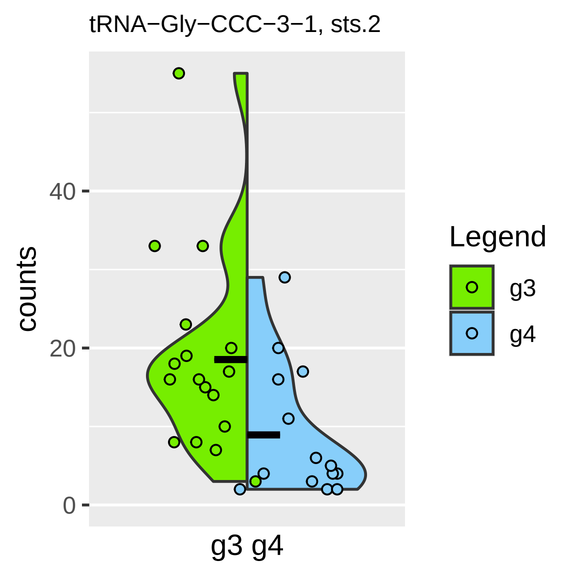
tRNA−Gly−CCC−3−1, sts.2
counts
40
20
0
g3 g4
Legend
g3
g4

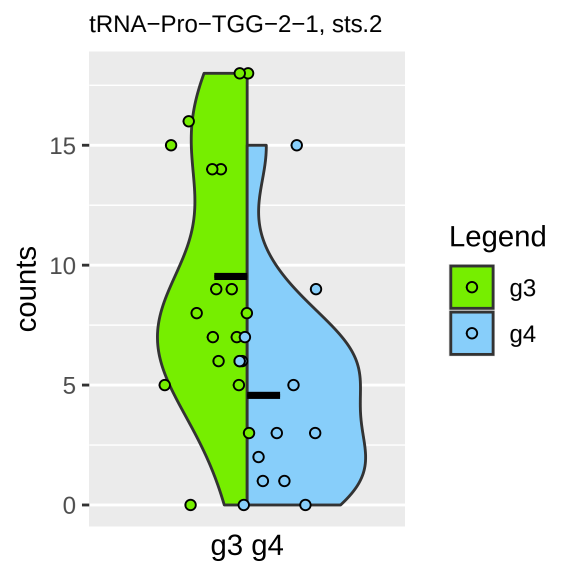

tRNA−Pro−TGG−2−1, sts.2
counts
15
10
5
0
g3 g4
Legend
g3
g4

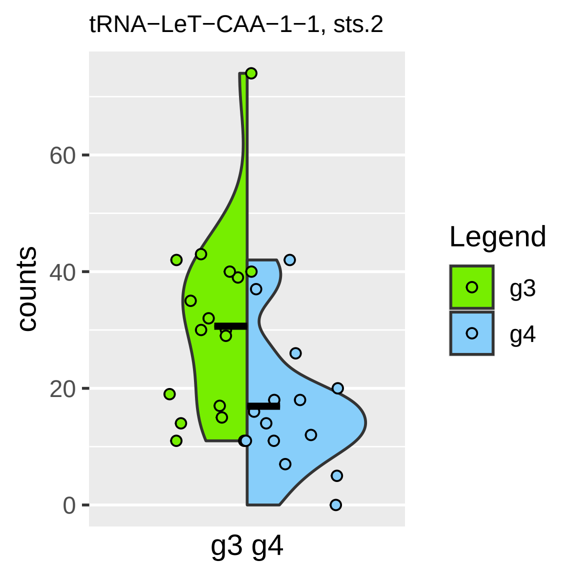
tRNA−LeT−CAA−1−1, sts.2
counts
0
20
40
60
g3 g4
Legend
g3
g4

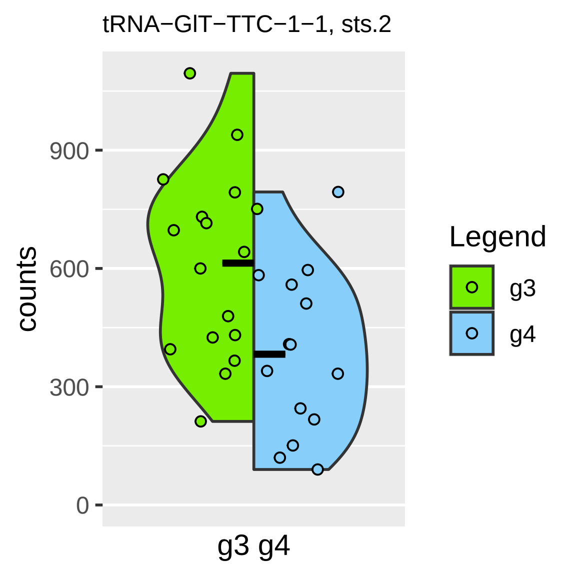

tRNA−GlT−TTC−1−1, sts.2
counts
900
600
300
0
g3 g4
Legend
g3
g4

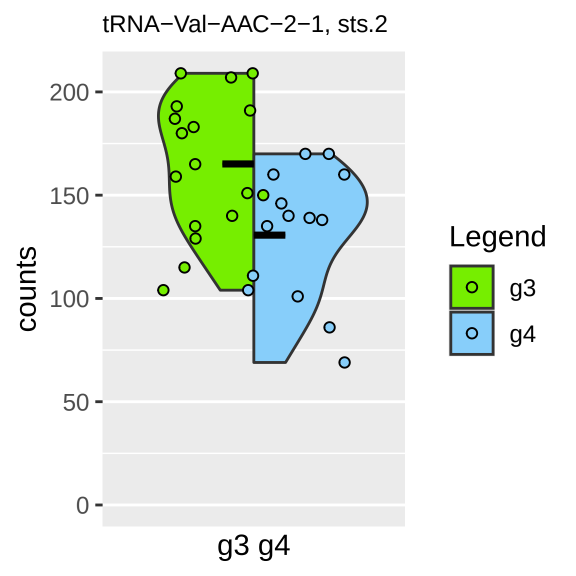

tRNA−Val−AAC−2−1, sts.2
counts
200
150
100
50
0
g3 g4
Legend
g3
g4

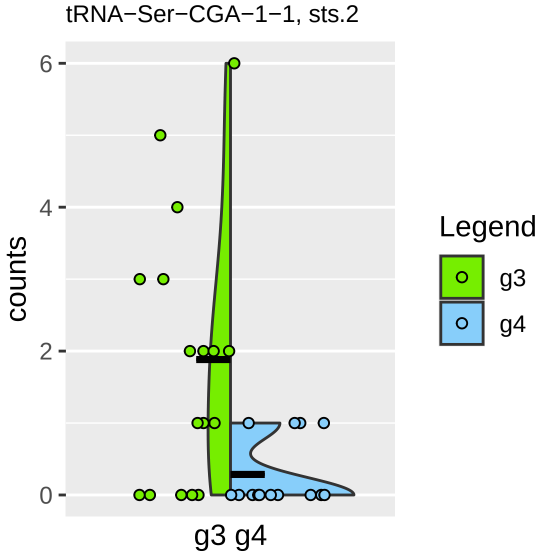
tRNA−Ser−CGA−1−1, sts.2
counts
6
4
2
0
g3 g4
Legend
g3
g4

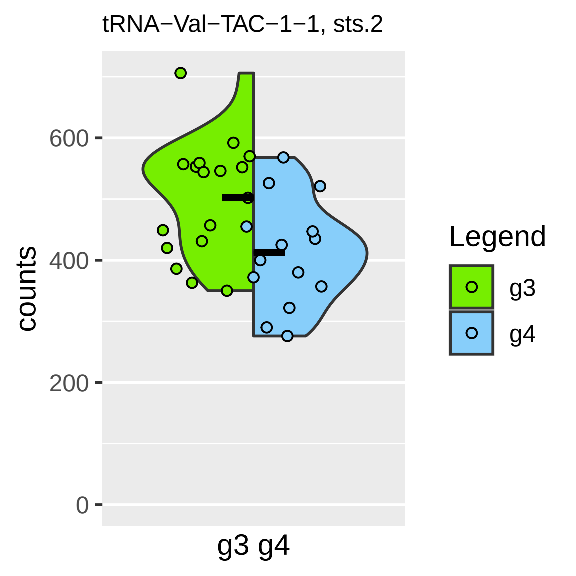

tRNA−Val−TAC−1−1, sts.2
counts
600
400
200
0
g3 g4
Legend
g3
g4

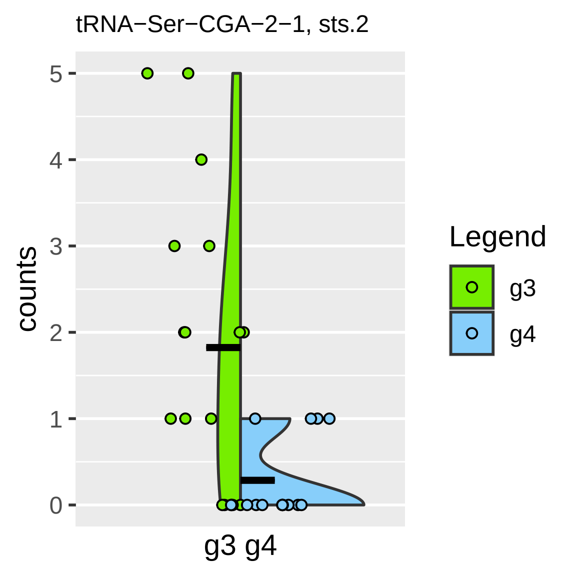
tRNA−Ser−CGA−2−1, sts.2
counts
5
4
3
2
1
0
g3 g4
Legend
g3
g4

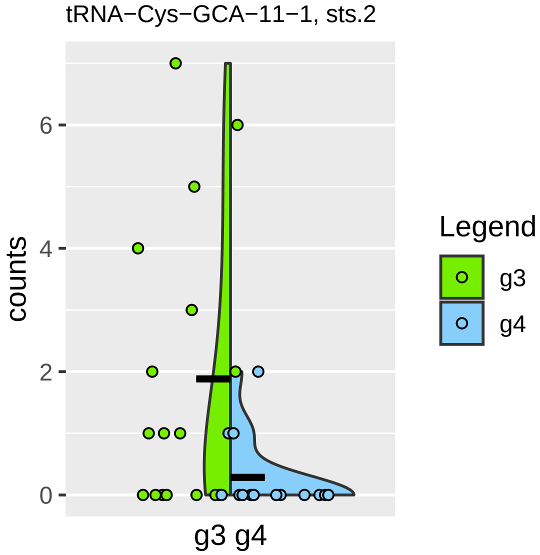

tRNA−Cys−GCA−11−1, sts.2
counts
0
2
4
6
g3 g4
Legend
g3
g4

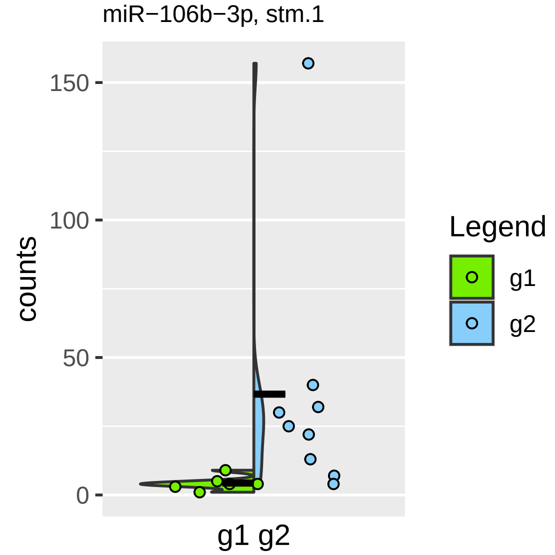

miR−106b−3p, stm.1
counts
150
100
50
0
g1 g2
Legend
g1
g2

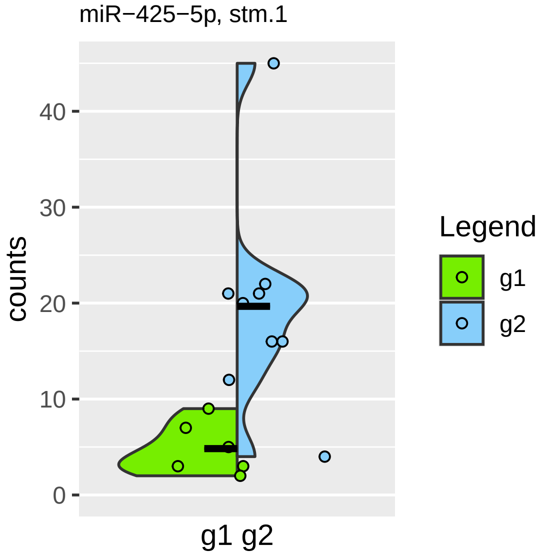
miR−425−5p, stm.1
counts
0
10
20
30
40
g1 g2
Legend
g1
g2

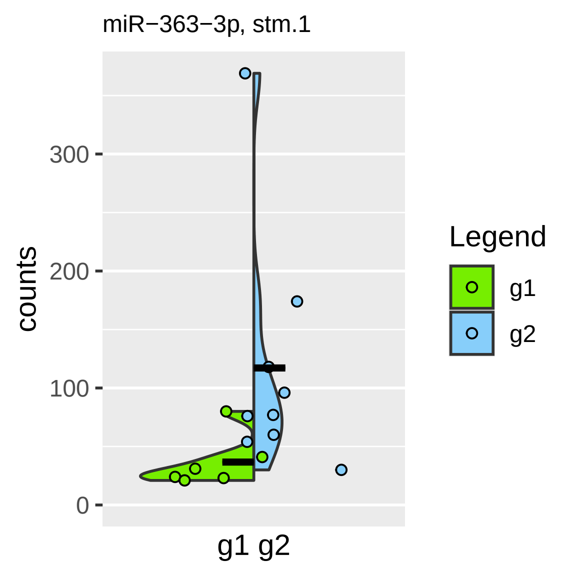

miR−363−3p, stm.1
counts
300
200
100
0
g1 g2
Legend
g1
g2

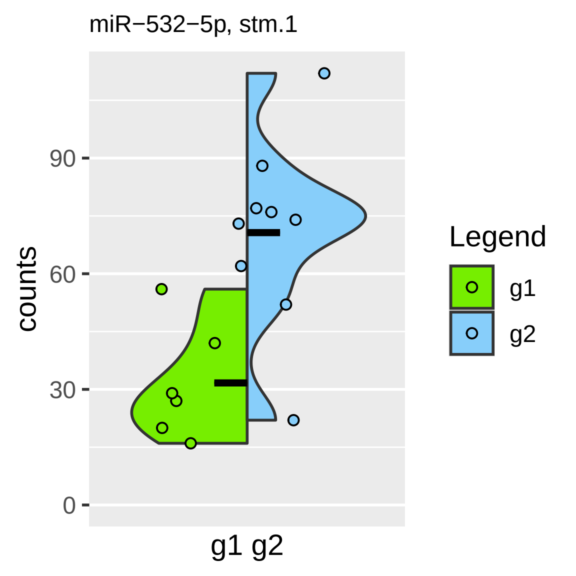

miR−532−5p, stm.1
counts
90
60
30
0
g1 g2
Legend
g1
g2

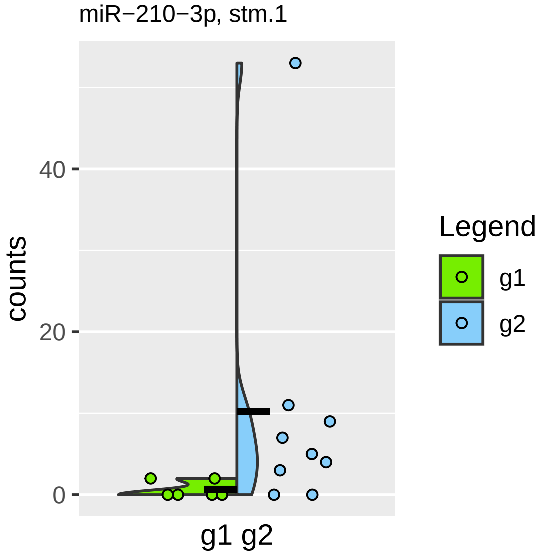

miR−210−3p, stm.1
counts
40
20
0
g1 g2
Legend
g1
g2

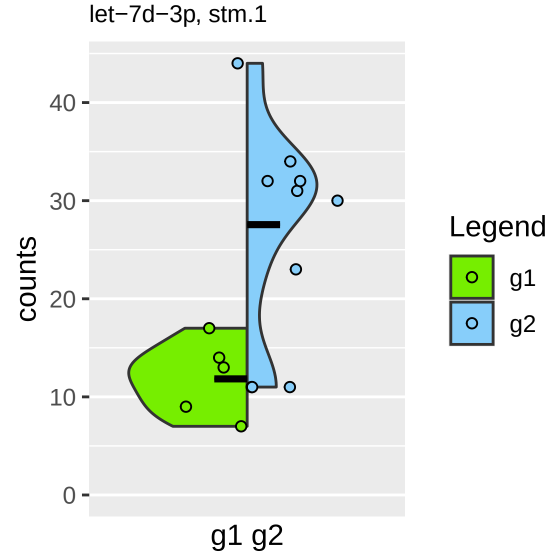
let−7d−3p, stm.1
counts
40
30
20
10
0
g1 g2
Legend
g1
g2

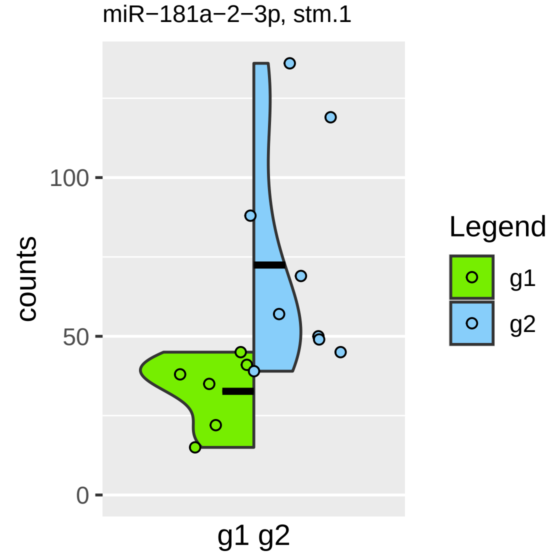

miR−181a−2−3p, stm.1
counts
100
50
0
g1 g2
Legend
g1
g2

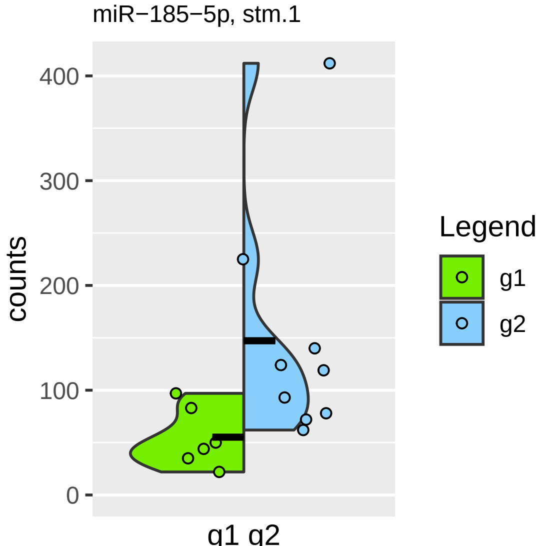

miR−185−5p, stm.1
counts
400
300
200
100
0
g1 g2
Legend
g1
g2

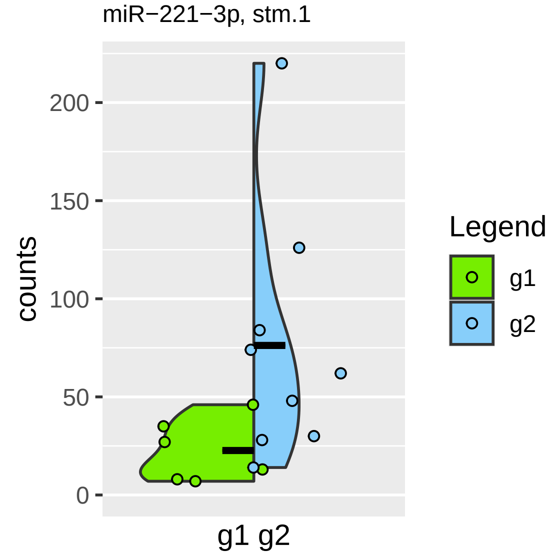
miR−221−3p, stm.1
counts
200
150
100
50
0
g1 g2
Legend
g1
g2

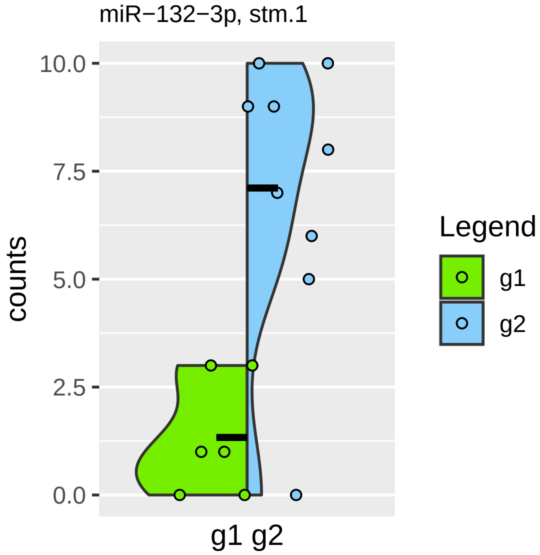
miR−132−3p, stm.1
10.0
7.5
5.0
2.5
0.0
counts
g1 g2
Legend
g1
g2

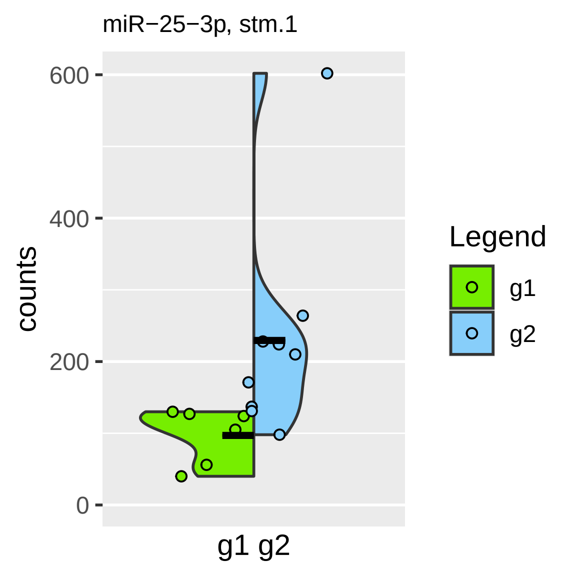

miR−25−3p, stm.1
600
400
200
0
counts
g1 g2
Legend
g1
g2

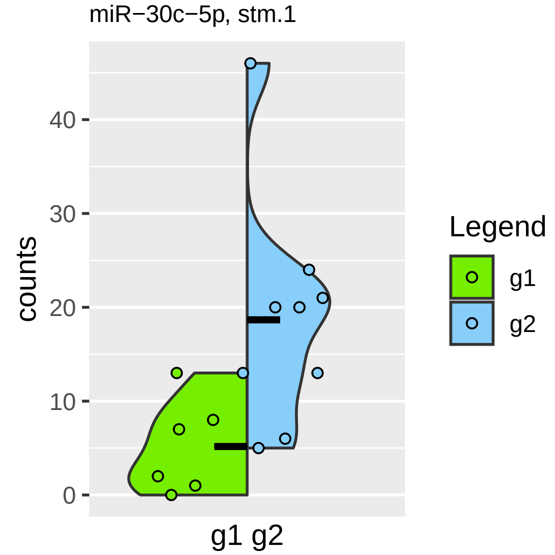

miR−30c−5p, stm.1
counts
0
10
20
30
40
g1 g2
Legend
g1
g2

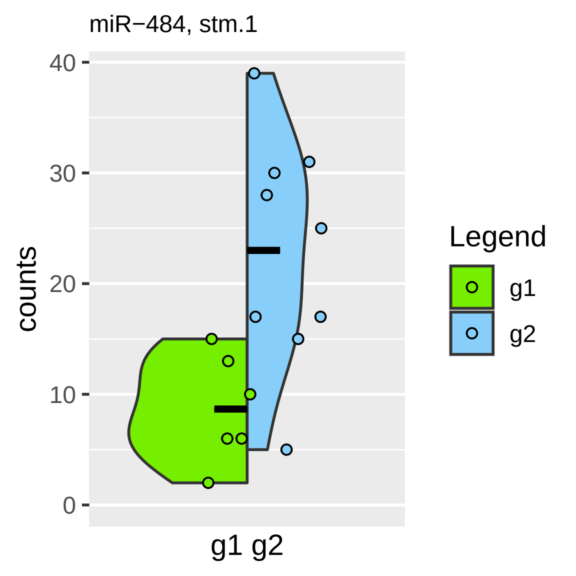
miR−484, stm.1
40
30
20
10
0
counts
g1 g2
Legend
g1
g2

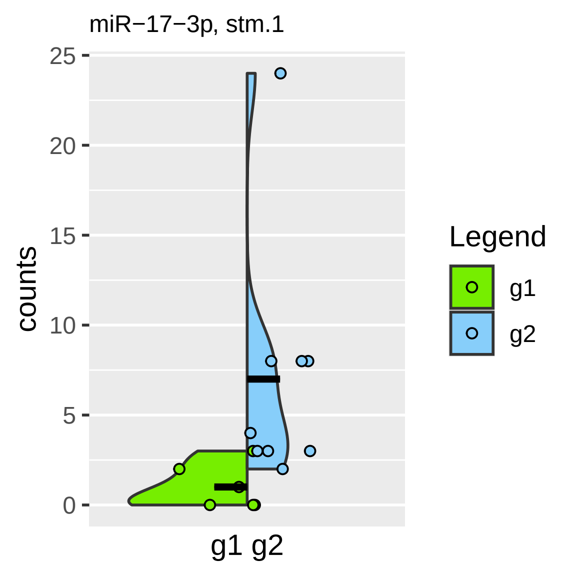

miR−17−3p, stm.1
counts
25
20
15
10
5
0
g1 g2
Legend
g1
g2

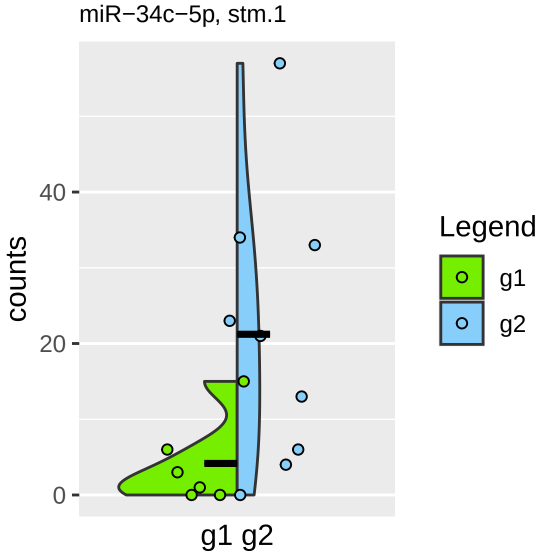

miR−34c−5p, stm.1
counts
40
20
0
g1 g2
Legend
g1
g2

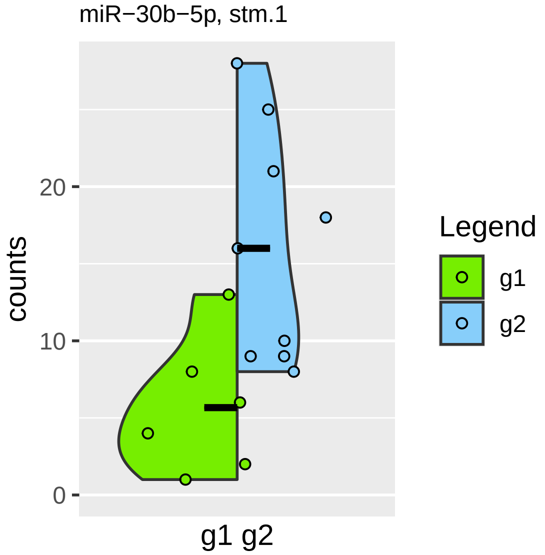

miR−30b−5p, stm.1
counts
20
10
0
g1 g2
Legend
g1
g2

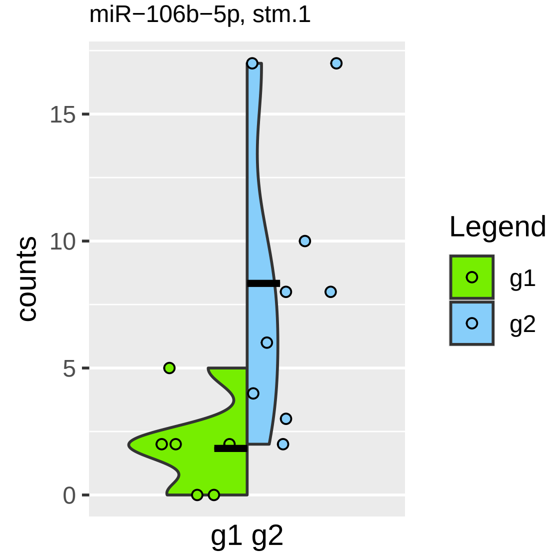
miR−106b−5p, stm.1
counts
15
10
5
0
g1 g2
Legend
g1
g2

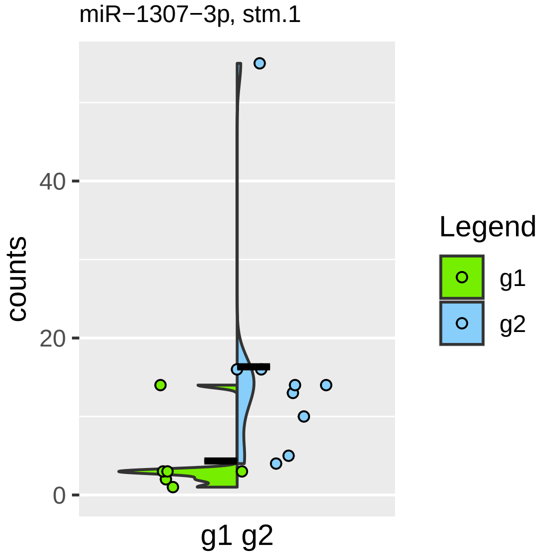

miR−1307−3p, stm.1
counts
40
20
0
g1 g2
Legend
g1
g2

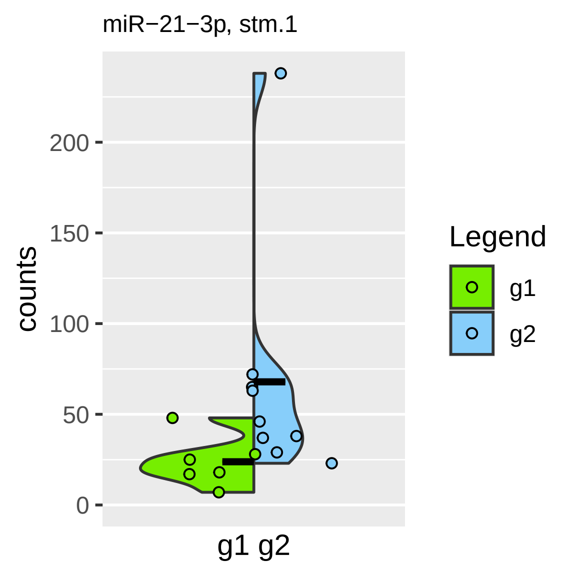

miR−21−3p, stm.1
counts
200
150
100
50
0
g1 g2
Legend
g1
g2

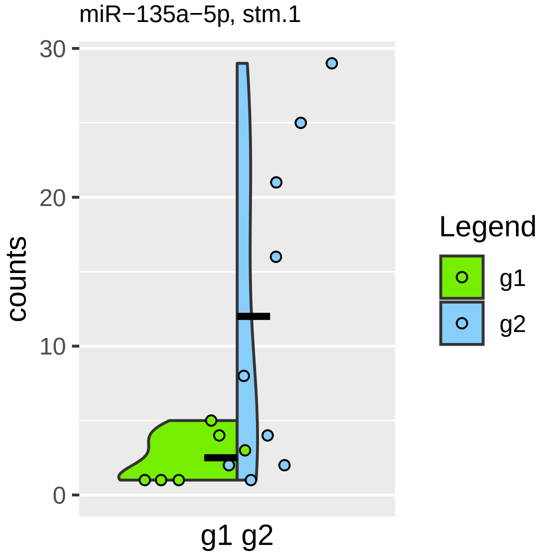

miR−135a−5p, stm.1
counts
30
20
10
0
g1 g2
Legend
g1
g2

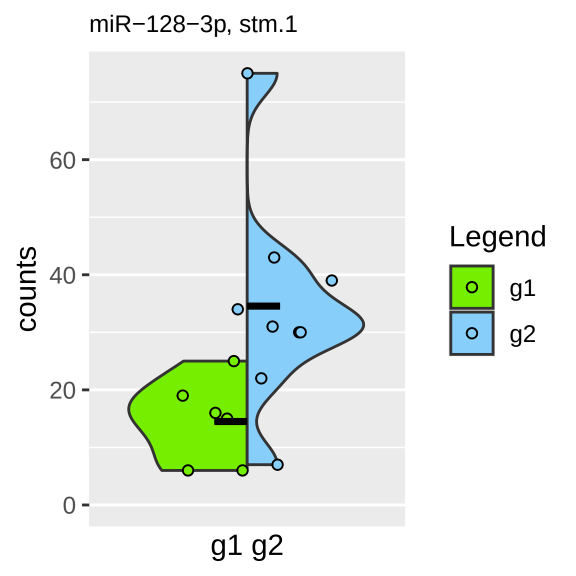
miR−128−3p, stm.1
counts
0
20
40
60
g1 g2
Legend
g1
g2

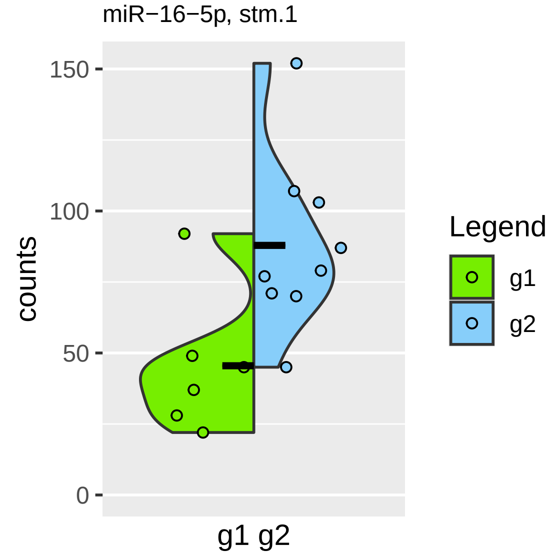

miR−16−5p, stm.1
150
100
50
0
counts
g1 g2
Legend
g1
g2

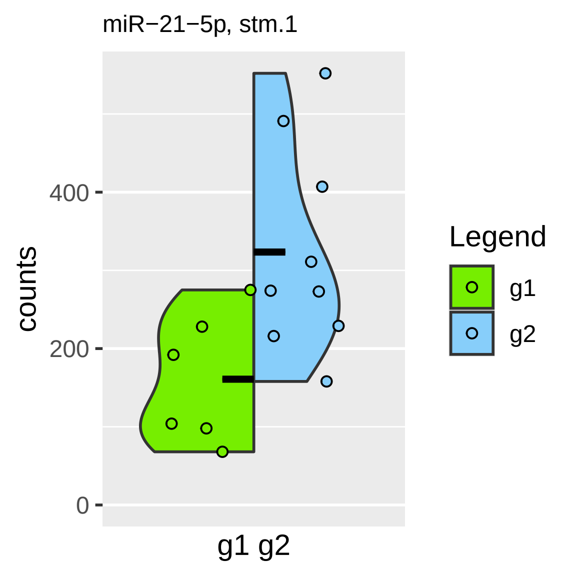
miR−21−5p, stm.1
counts
400
200
0
g1 g2
Legend
g1
g2

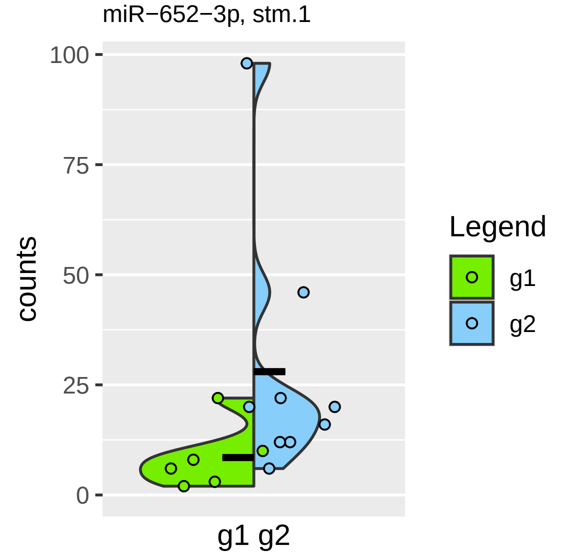
miR−652−3p, stm.1
100
75
50
25
0
counts
g1 g2
Legend
g1
g2

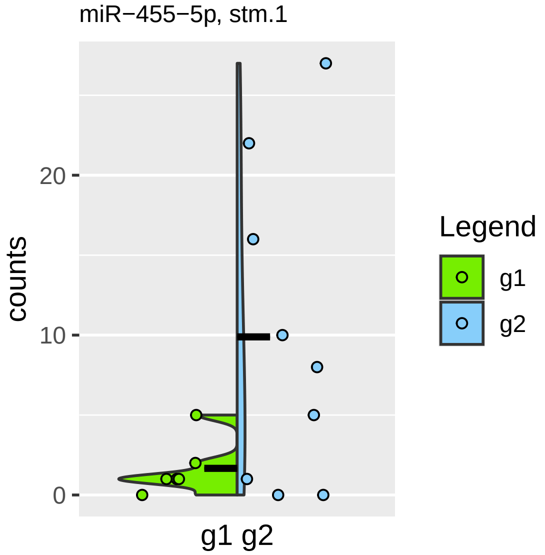

miR−455−5p, stm.1
counts
0
10
20
g1 g2
Legend
g1
g2

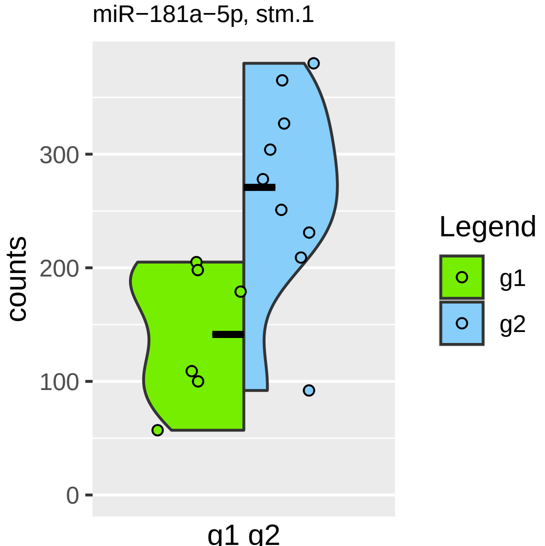

miR−181a−5p, stm.1
counts
300
200
100
0
g1 g2
Legend
g1
g2

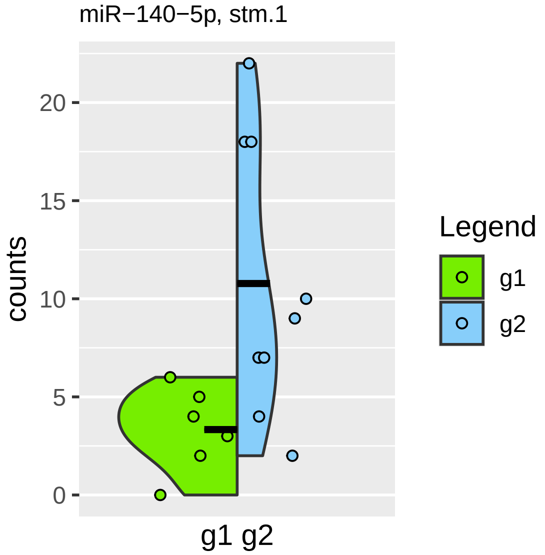

miR−140−5p, stm.1
counts
20
15
10
5
0
g1 g2
Legend
g1
g2

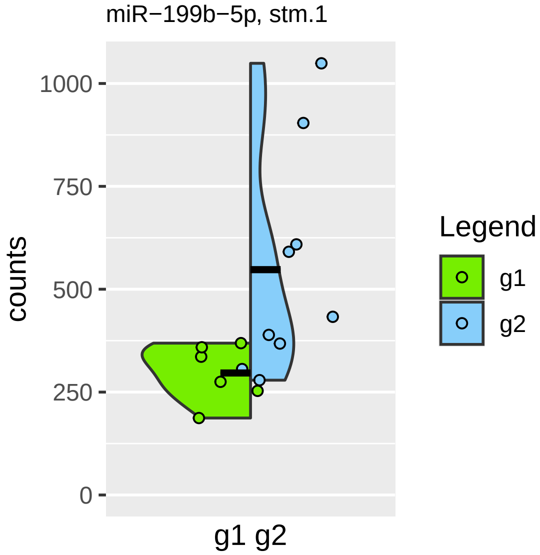

miR−199b−5p, stm.1
counts
1000
750
500
250
0
g1 g2
Legend
g1
g2

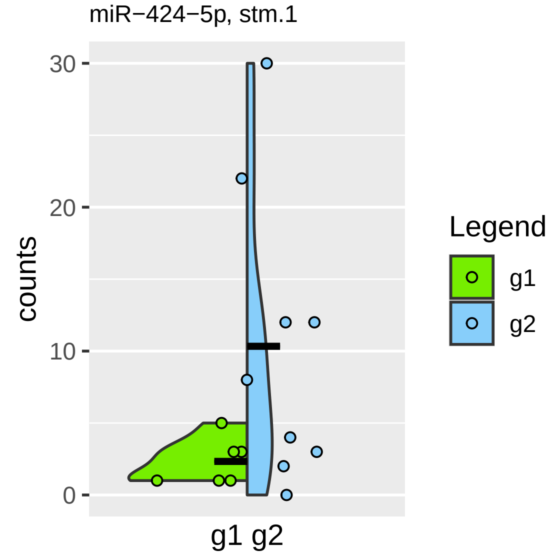

miR−424−5p, stm.1
30
20
10
0
counts
g1 g2
Legend
g1
g2

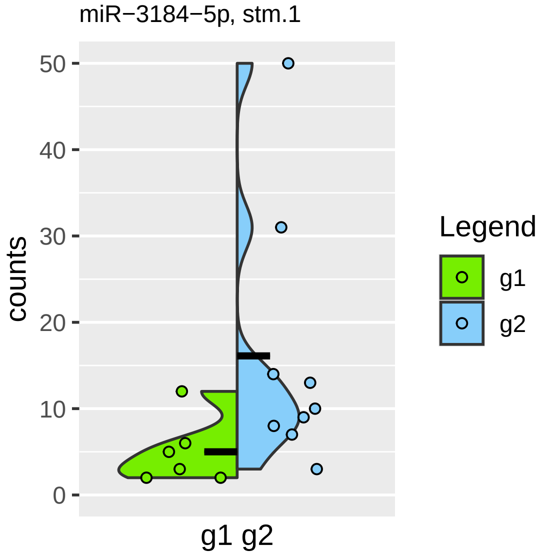
miR−3184−5p, stm.1
counts
50
40
30
20
10
0
g1 g2
Legend
g1
g2

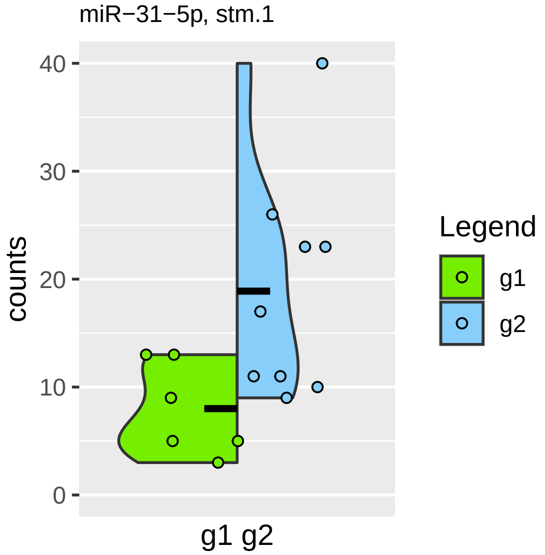

miR−31−5p, stm.1
40
30
20
10
0
counts
g1 g2
Legend
g1
g2

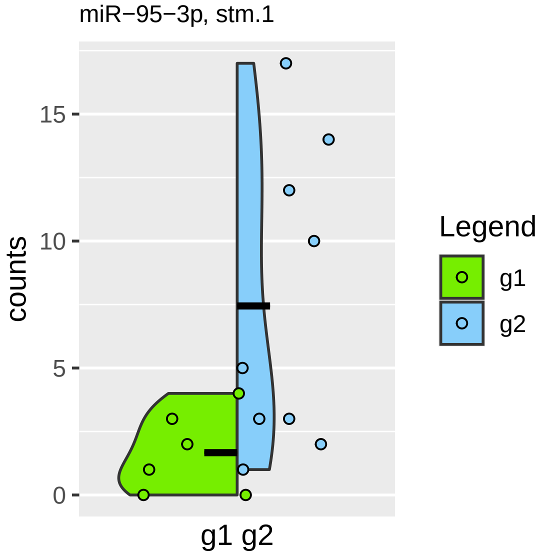

miR−95−3p, stm.1
counts
0
5
10
15
g1 g2
Legend
g1
g2

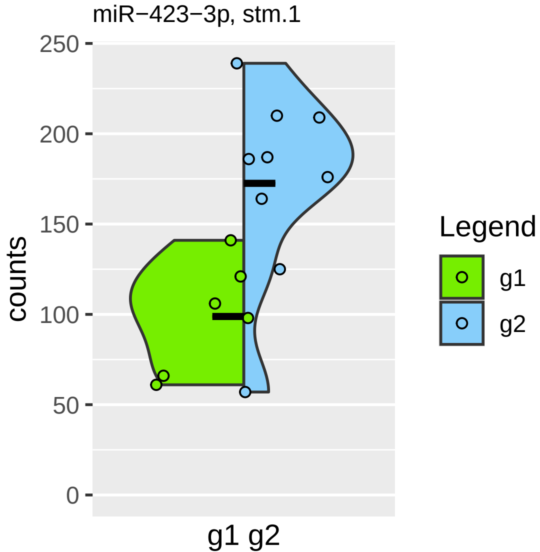
miR−423−3p, stm.1
counts
250
200
150
100
50
0
g1 g2
Legend
g1
g2

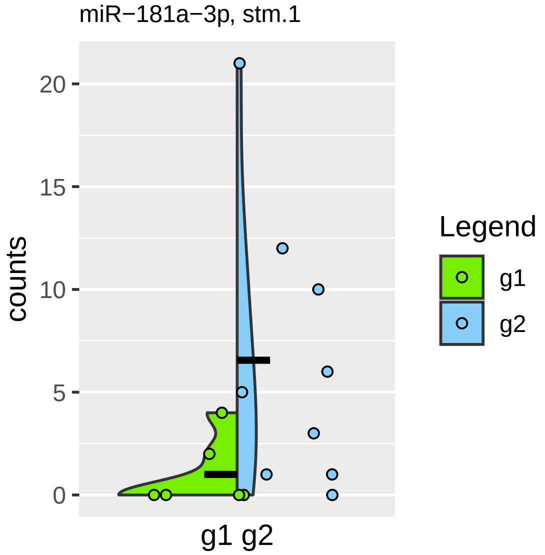

miR−181a−3p, stm.1
counts
20
15
10
5
0
g1 g2
Legend
g1
g2

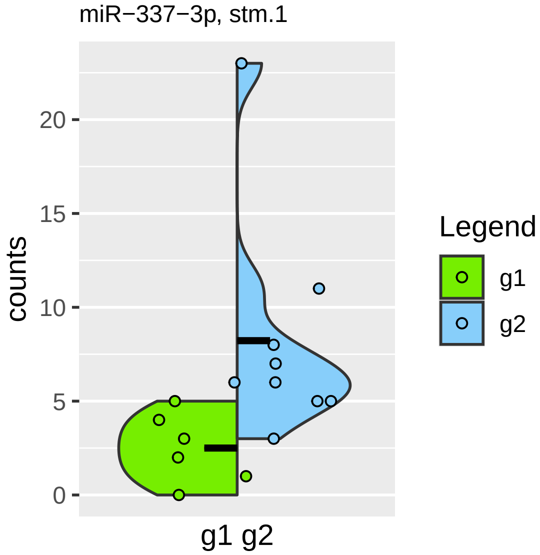
miR−337−3p, stm.1
counts
0
5
10
15
20
g1 g2
Legend
g1
g2

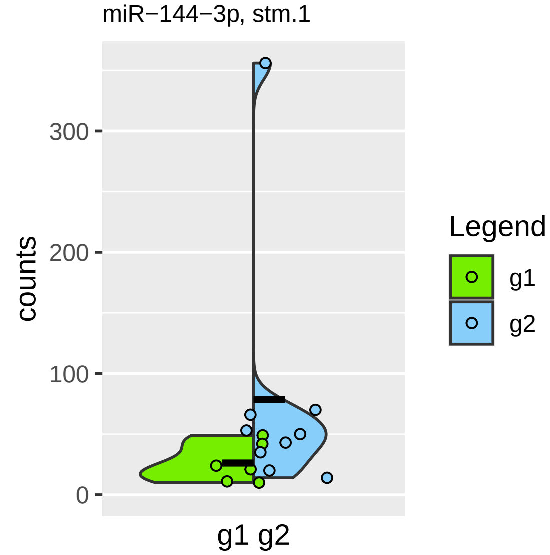

miR−144−3p, stm.1
counts
300
200
100
0
g1 g2
Legend
g1
g2

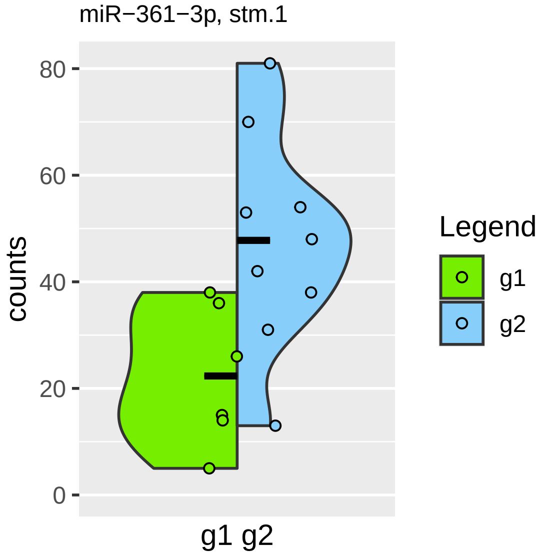

miR−361−3p, stm.1
counts
80
60
40
20
0
g1 g2
Legend
g1
g2

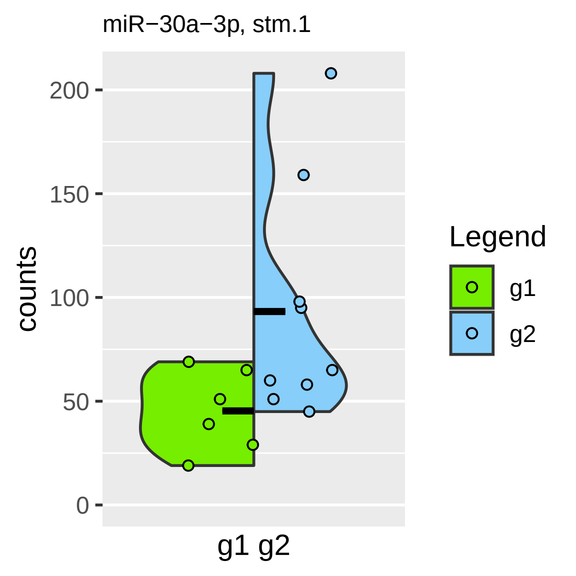

miR−30a−3p, stm.1
200
150
100
50
0
counts
g1 g2
Legend
g1
g2

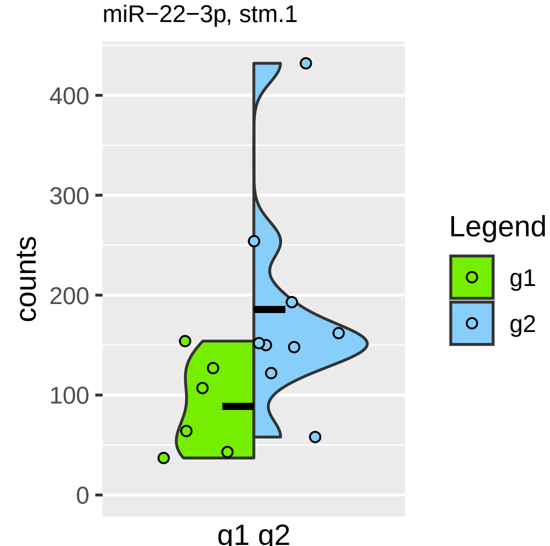

miR−22−3p, stm.1
400
300
200
100
0
counts
g1 g2
Legend
g1
g2

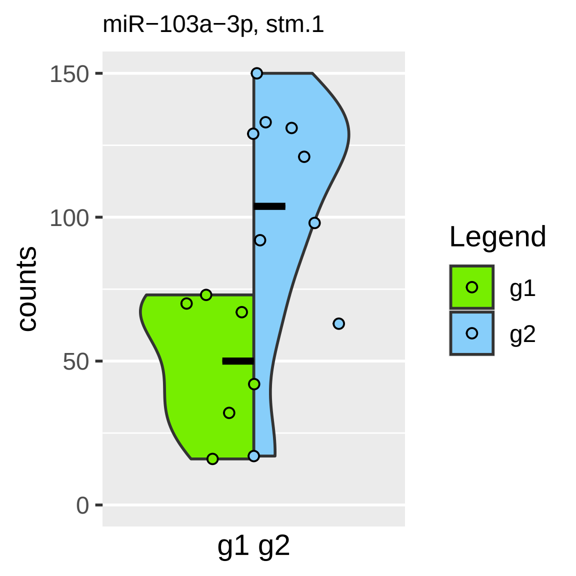

miR−103a−3p, stm.1
150
100
50
0
counts
g1 g2
Legend
g1
g2

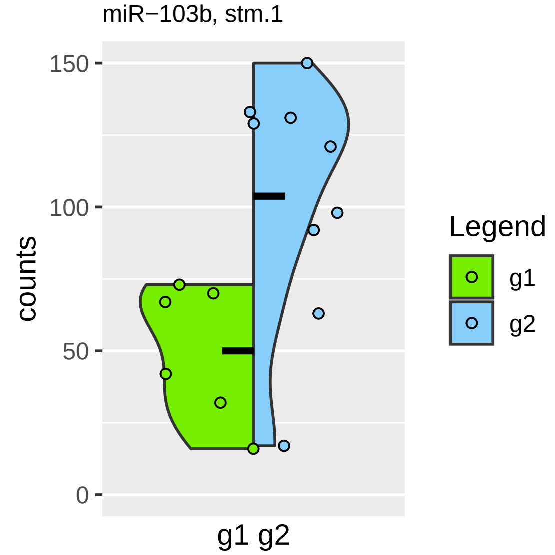
miR−103b, stm.1
150
100
50
0
counts
g1 g2
Legend
g1
g2

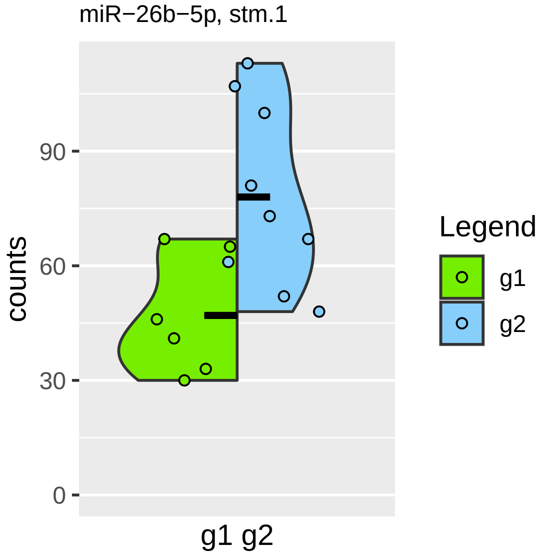

miR−26b−5p, stm.1
counts
90
60
30
0
g1 g2
Legend
g1
g2

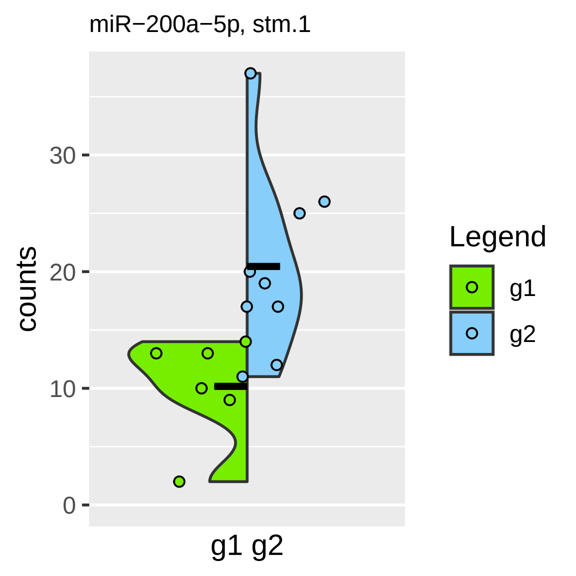
miR−200a−5p, stm.1
counts
30
20
10
0
g1 g2
Legend
g1
g2

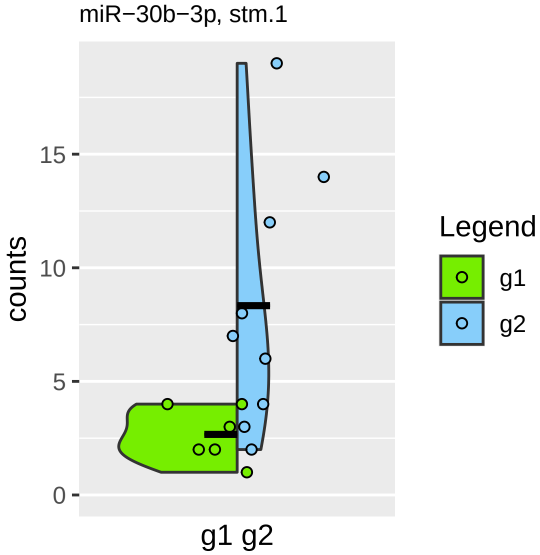
miR−30b−3p, stm.1
counts
0
5
10
15
g1 g2
Legend
g1
g2

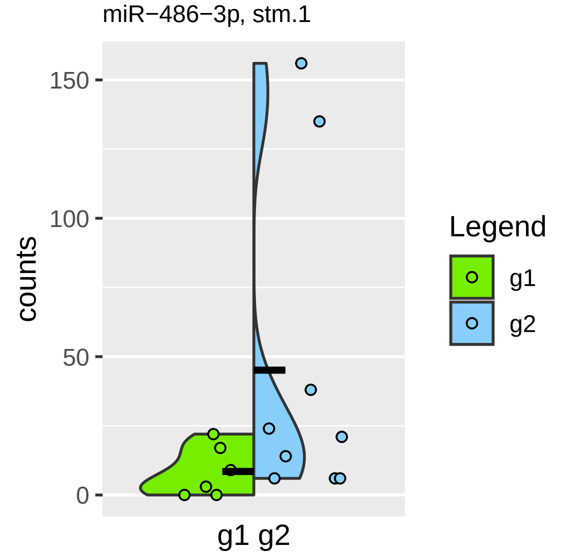

miR−486−3p, stm.1
150
100
50
0
counts
g1 g2
Legend
g1
g2

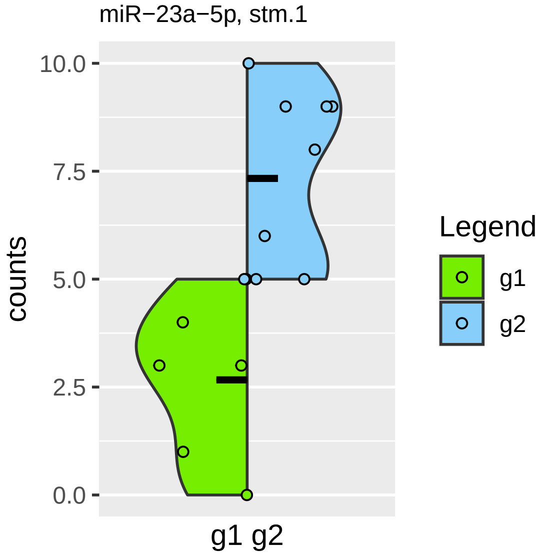

miR−23a−5p, stm.1
10.0
7.5
5.0
2.5
0.0
counts
g1 g2
Legend
g1
g2

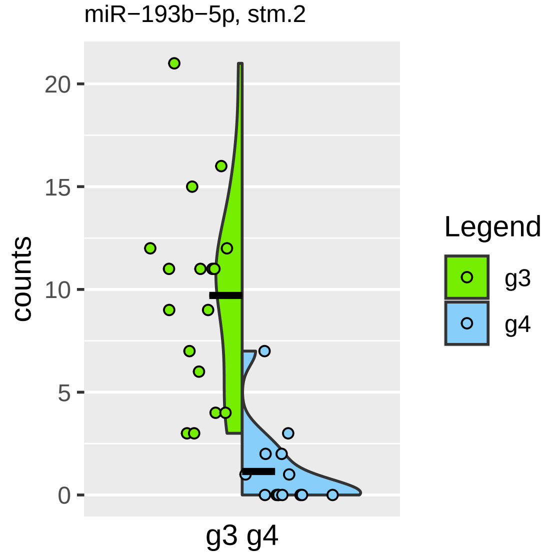

miR−193b−5p, stm.2
counts
20
15
10
5
0
g3 g4
Legend
g3
g4

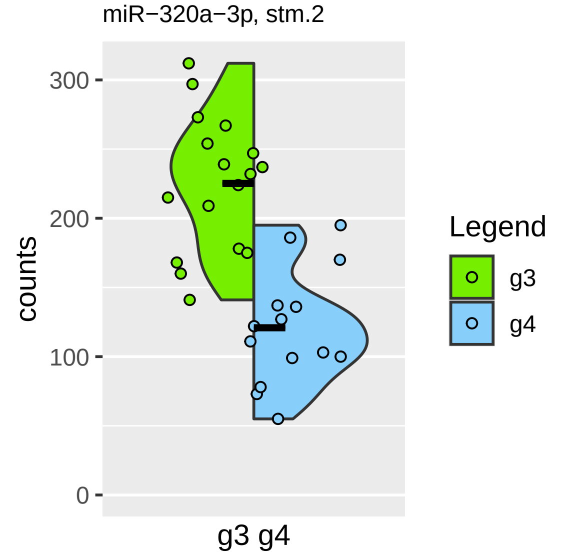
miR−320a−3p, stm.2
counts
300
200
100
0
g3 g4
Legend
g3
g4

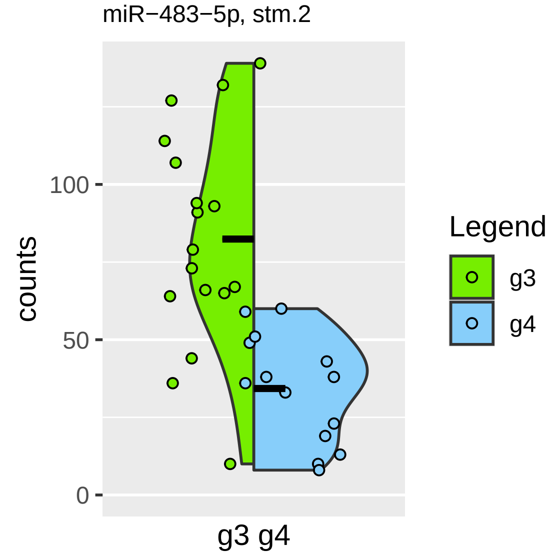

miR−483−5p, stm.2
counts
100
50
0
g3 g4
Legend
g3
g4

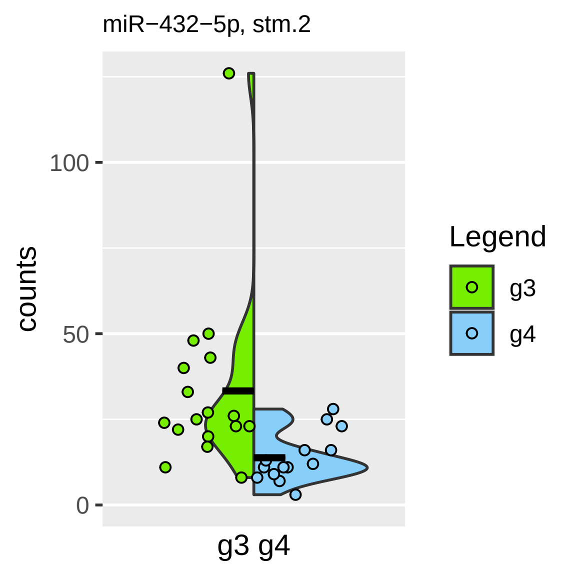

miR−432−5p, stm.2
counts
100
50
0
g3 g4
Legend
g3
g4

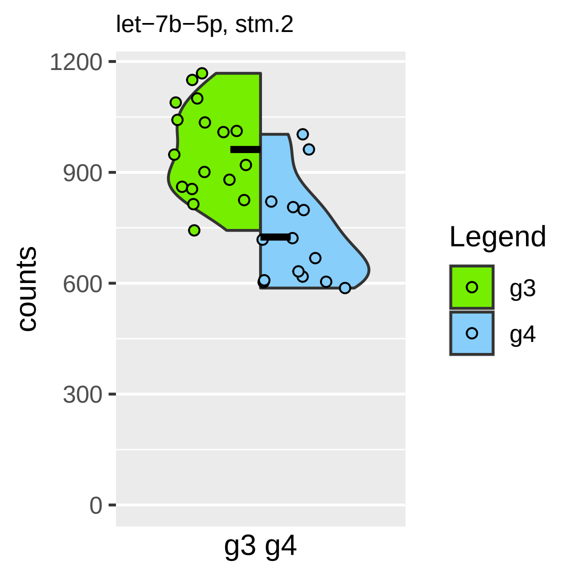

let−7b−5p, stm.2
1200
900
600
300
0
counts
g3 g4
Legend
g3
g4

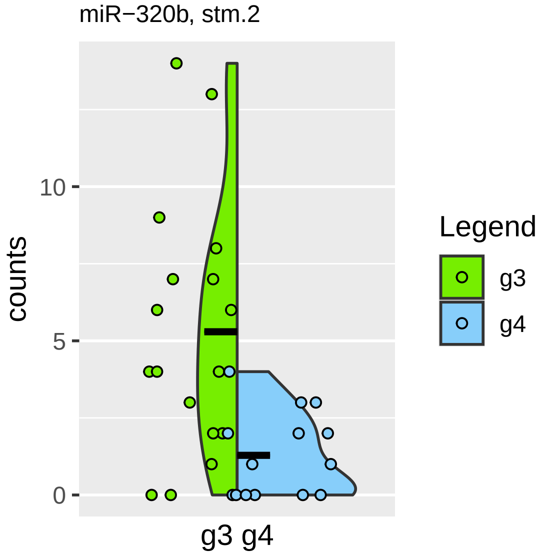
miR−320b, stm.2
counts
10
5
0
g3 g4
Legend
g3
g4

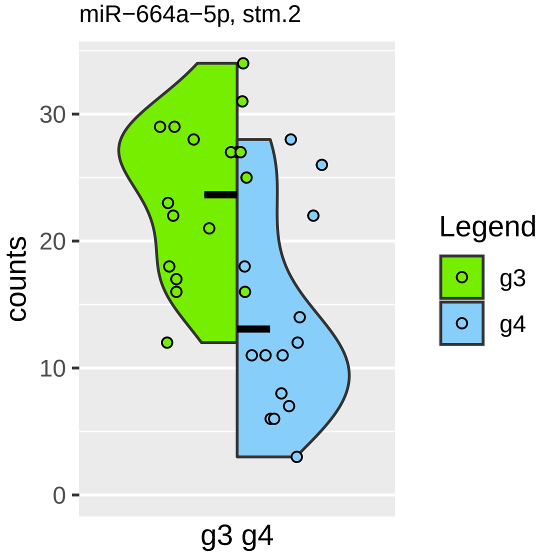

miR−664a−5p, stm.2
counts
30
20
10
0
g3 g4
Legend
g3
g4

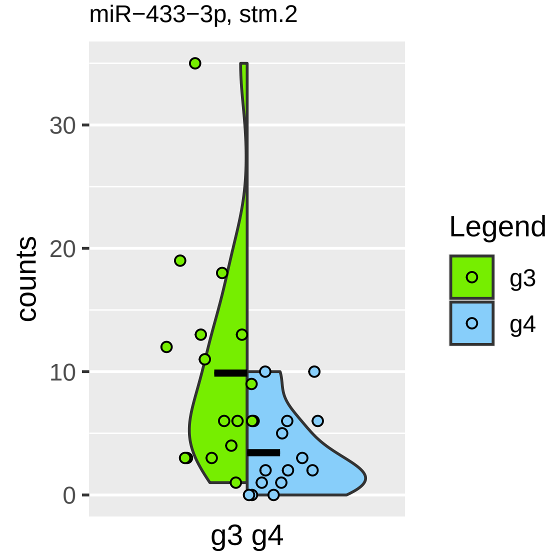

miR−433−3p, stm.2
counts
30
20
10
0
g3 g4
Legend
g3
g4

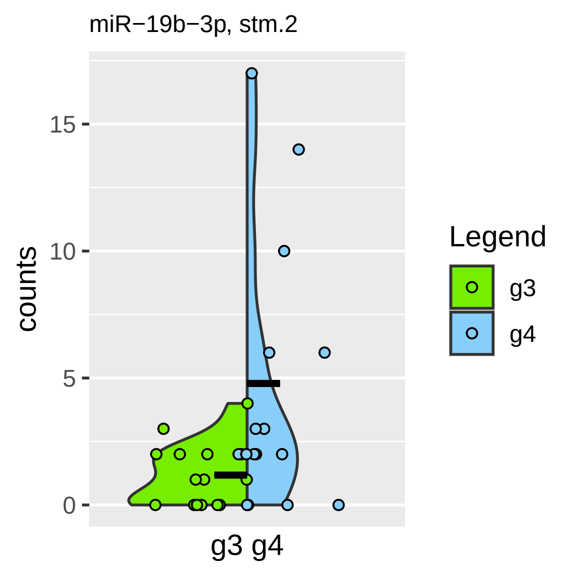
miR−19b−3p, stm.2
counts
15
10
5
0
g3 g4
Legend
g3
g4

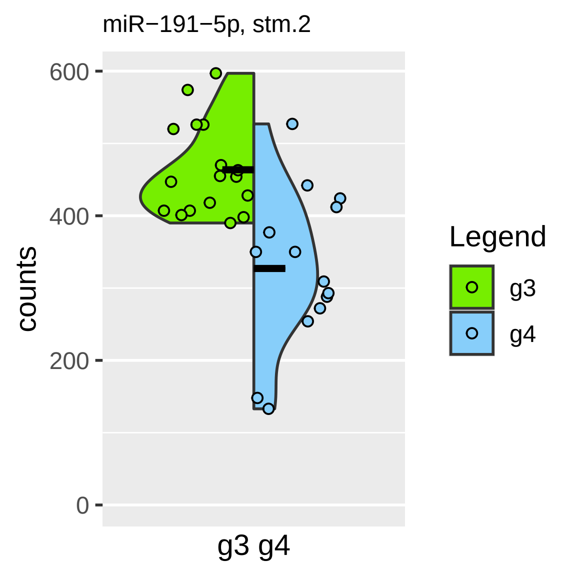

miR−191−5p, stm.2
600
400
200
0
counts
g3 g4
Legend
g3
g4

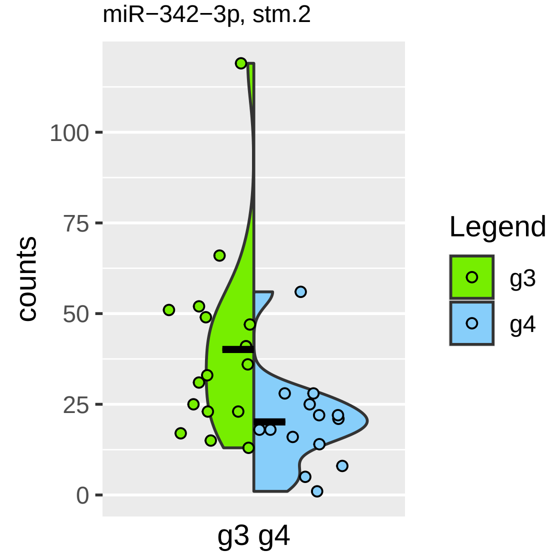

miR−342−3p, stm.2
counts
100
75
50
25
0
g3 g4
Legend
g3
g4

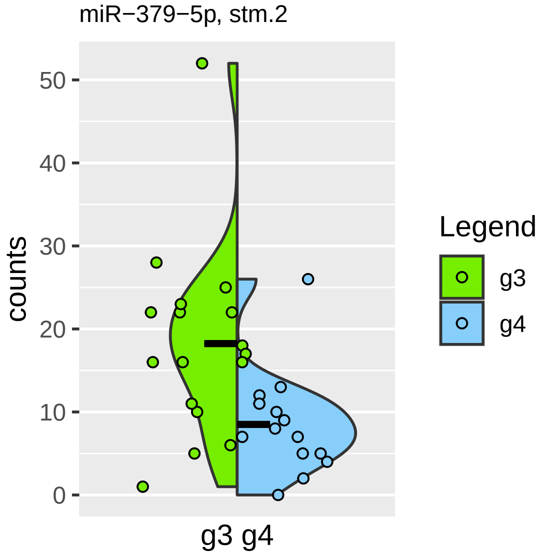

miR−379−5p, stm.2
counts
50
40
30
20
10
0
g3 g4
Legend
g3
g4

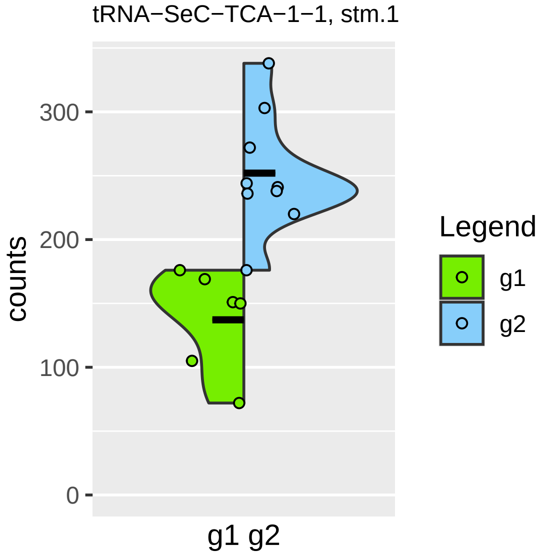
tRNA−SeC−TCA−1−1, stm.1
counts
300
200
100
0
g1 g2
Legend
g1
g2

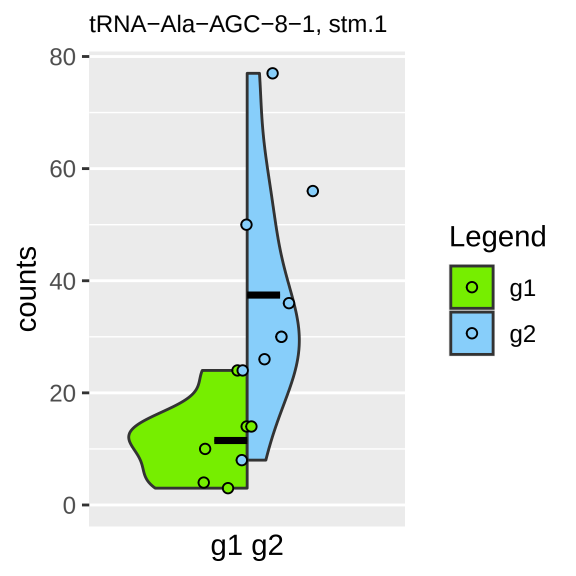

tRNA−Ala−AGC−8−1, stm.1
counts
80
60
40
20
0
g1 g2
Legend
g1
g2

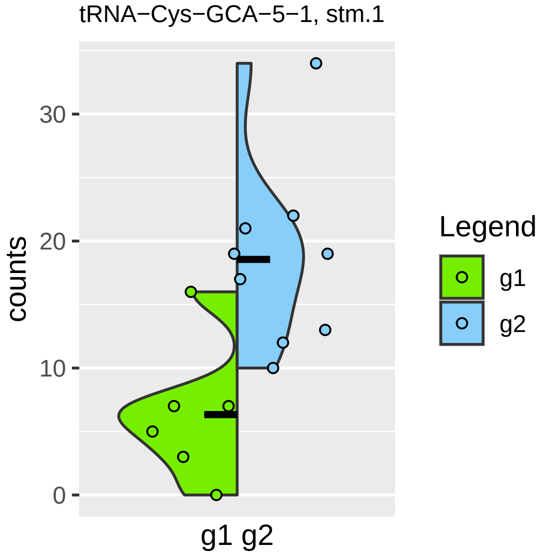
tRNA−Cys−GCA−5−1, stm.1
counts
30
20
10
0
g1 g2
Legend
g1
g2

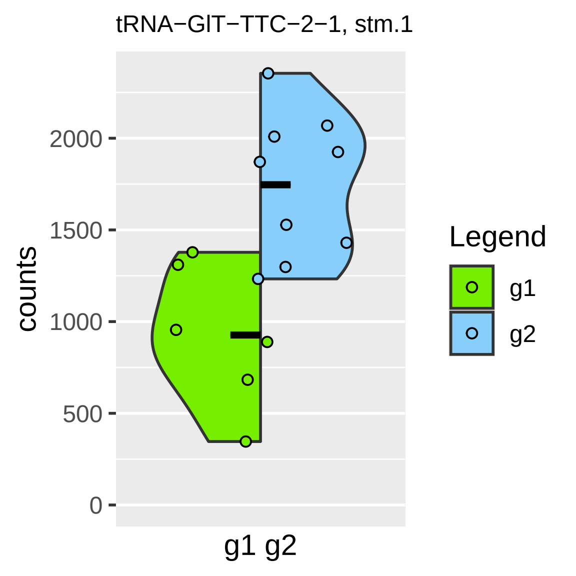
tRNA−GlT−TTC−2−1, stm.1
counts
2000
1500
1000
500
0
g1 g2
Legend
g1
g2

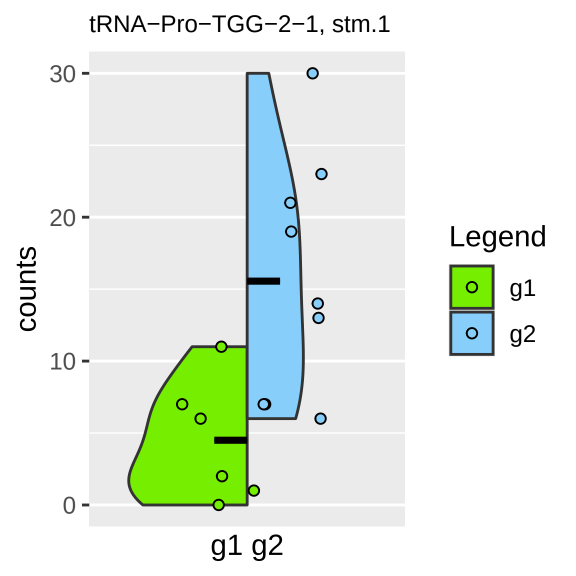
tRNA−Pro−TGG−2−1, stm.1
counts
30
20
10
0
g1 g2
Legend
g1
g2

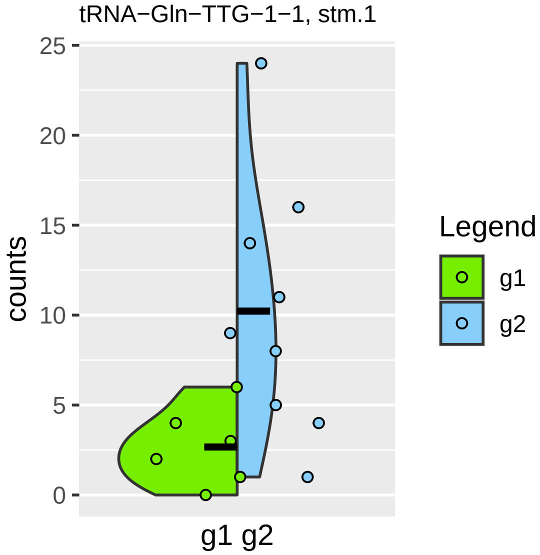

tRNA−Gln−TTG−1−1, stm.1
counts
25
20
15
10
5
0
g1 g2
Legend
g1
g2

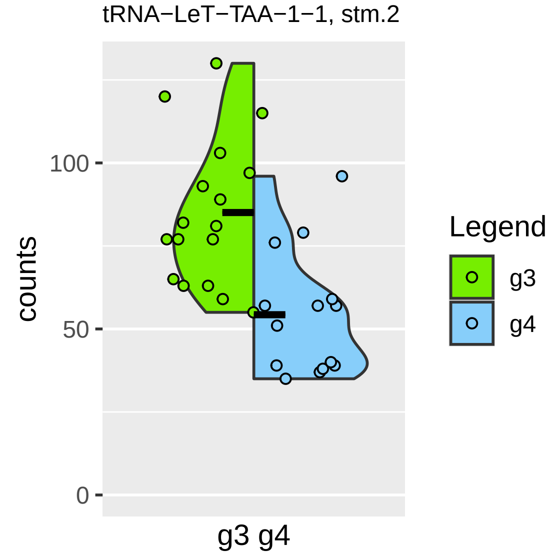
tRNA−LeT−TAA−1−1, stm.2
counts
100
50
0
g3 g4
Legend
g3
g4

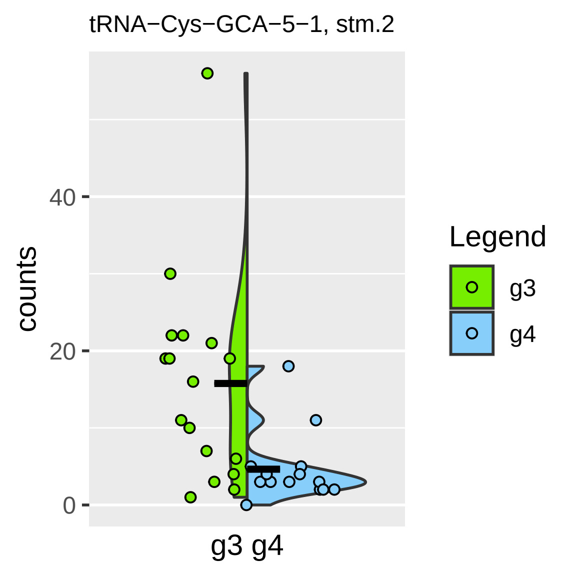
tRNA−Cys−GCA−5−1, stm.2
counts
40
20
0
g3 g4
Legend
g3
g4

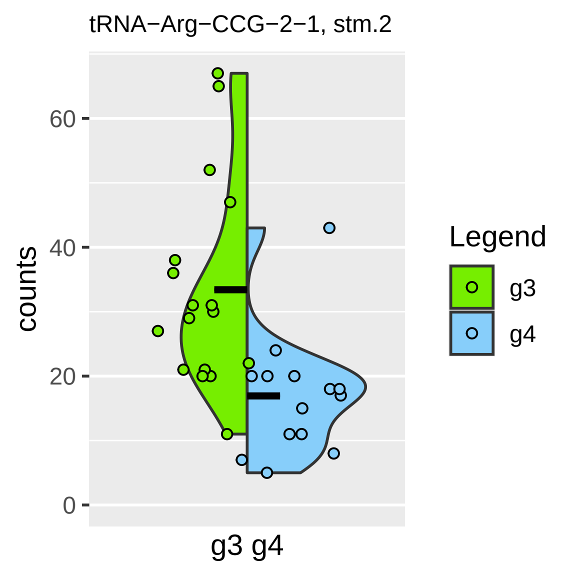
tRNA−Arg−CCG−2−1, stm.2
counts
60
40
20
0
g3 g4
Legend
g3
g4

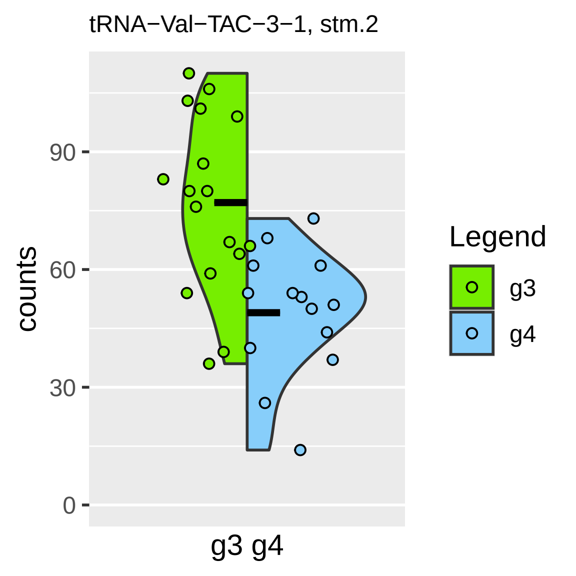
tRNA−Val−TAC−3−1, stm.2
counts
90
60
30
0
g3 g4
Legend
g3
g4

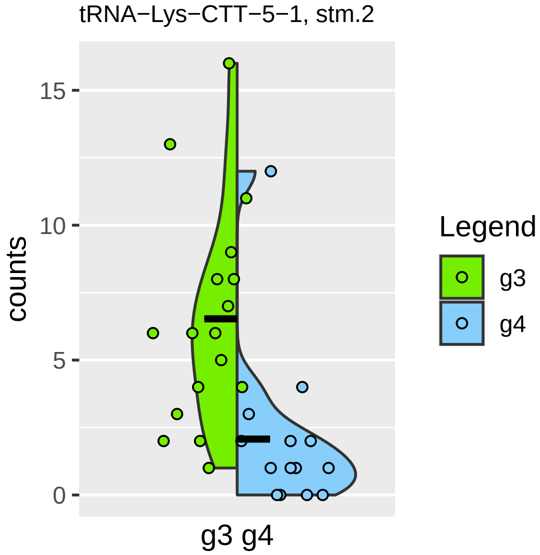
tRNA−Lys−CTT−5−1, stm.2
counts
15
10
5
0
g3 g4
Legend
g3
g4

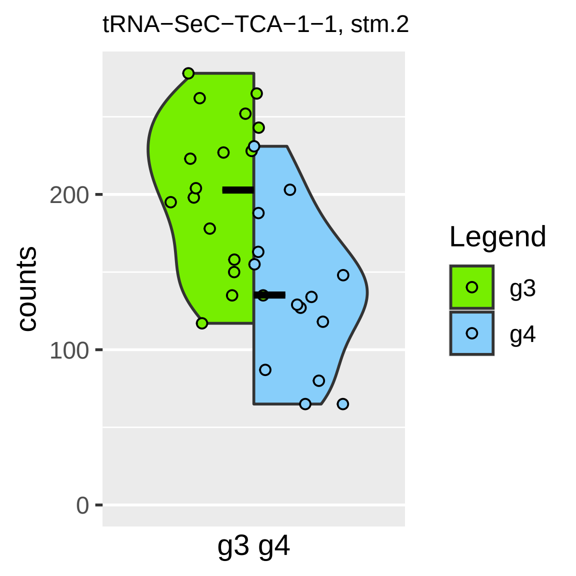
tRNA−SeC−TCA−1−1, stm.2
counts
200
100
0
g3 g4
Legend
g3
g4

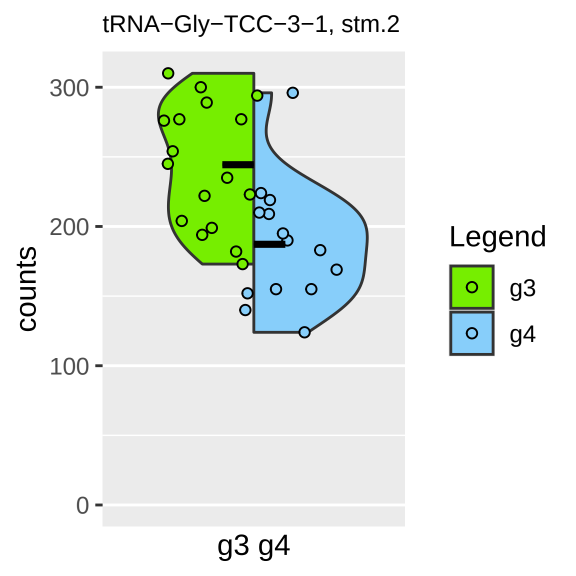

tRNA−Gly−TCC−3−1, stm.2
counts
300
200
100
0
g3 g4
Legend
g3
g4

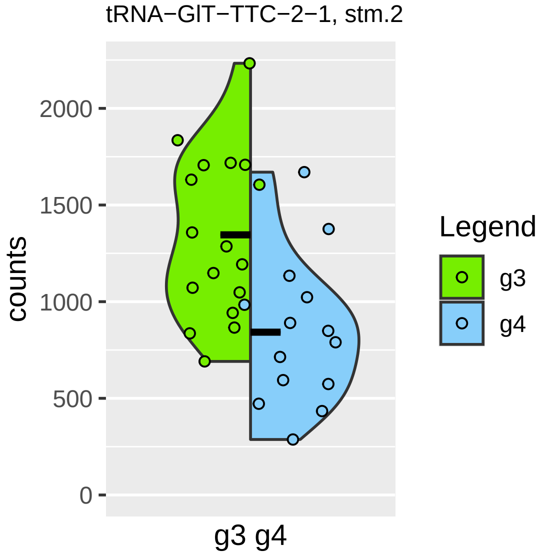

tRNA−GlT−TTC−2−1, stm.2
counts
2000
1500
1000
500
0
g3 g4
Legend
g3
g4

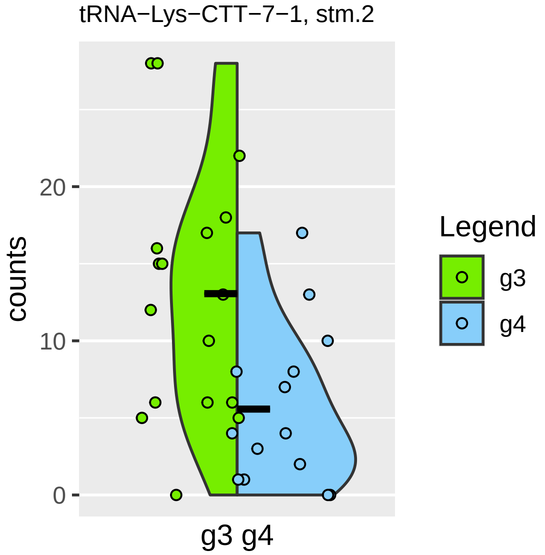
tRNA−Lys−CTT−7−1, stm.2
counts
20
10
0
g3 g4
Legend
g3
g4

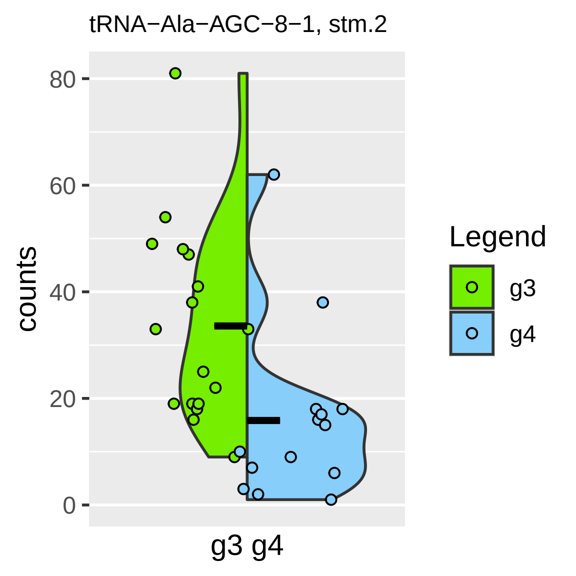

tRNA−Ala−AGC−8−1, stm.2
counts
0
20
40
60
80
g3 g4
Legend
g3
g4

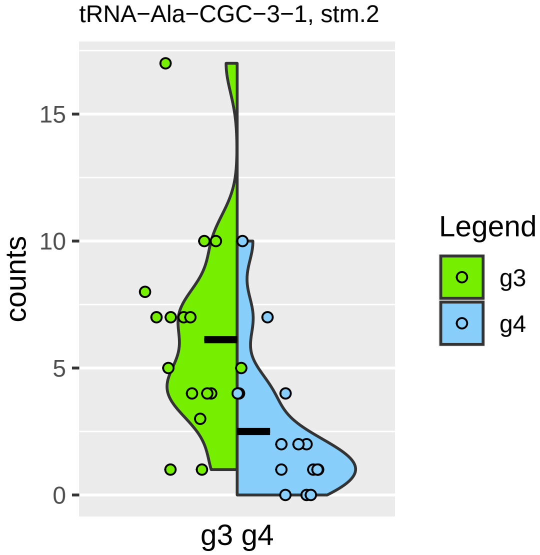

tRNA−Ala−CGC−3−1, stm.2
counts
15
10
5
0
g3 g4
Legend
g3
g4

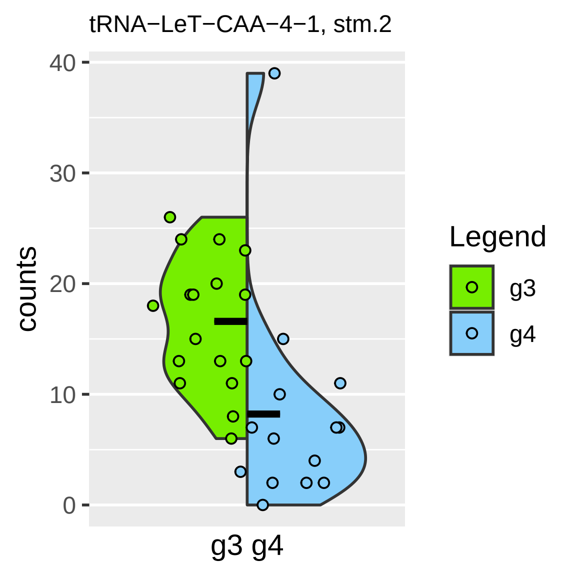
tRNA−LeT−CAA−4−1, stm.2
counts
40
30
20
10
0
g3 g4
Legend
g3
g4

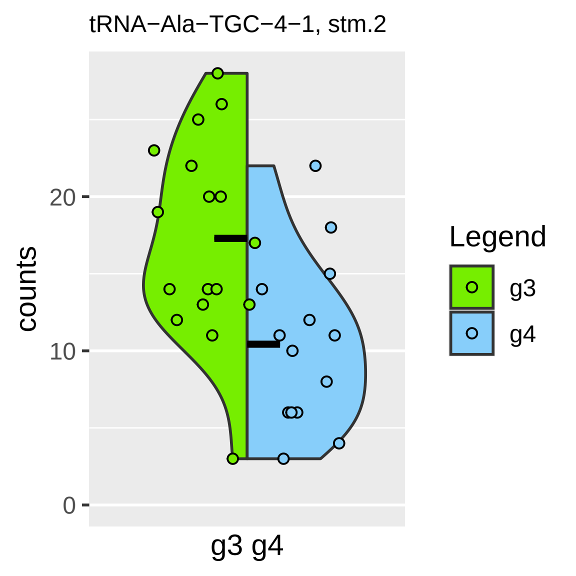
tRNA−Ala−TGC−4−1, stm.2
counts
20
10
0
g3 g4
Legend
g3
g4

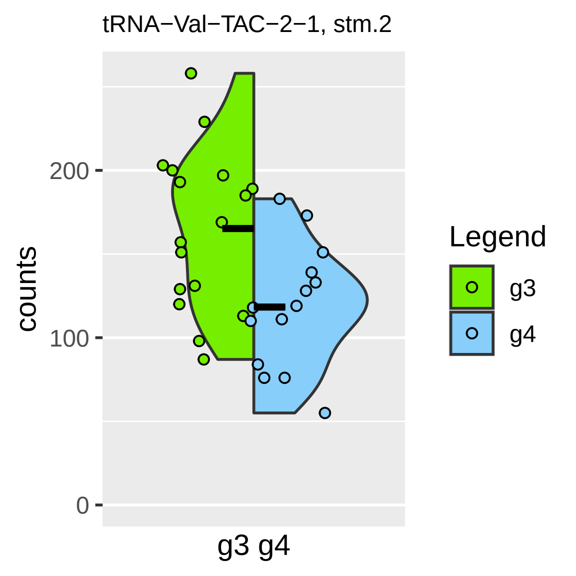

tRNA−Val−TAC−2−1, stm.2
counts
200
100
0
g3 g4
Legend
g3
g4

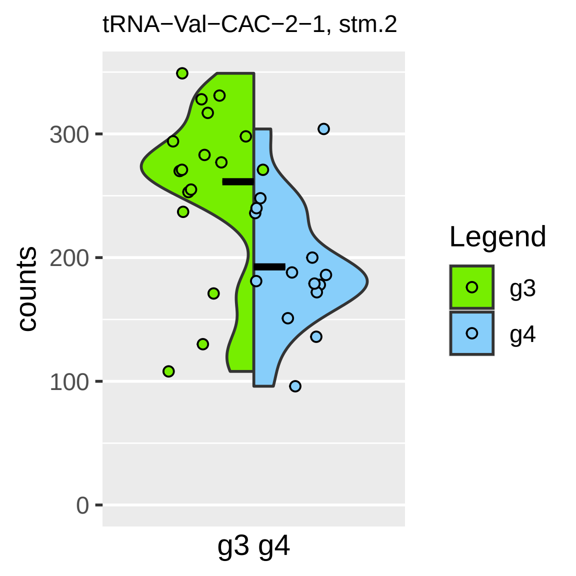

tRNA−Val−CAC−2−1, stm.2
300
200
100
0
counts
g3 g4
Legend
g3
g4

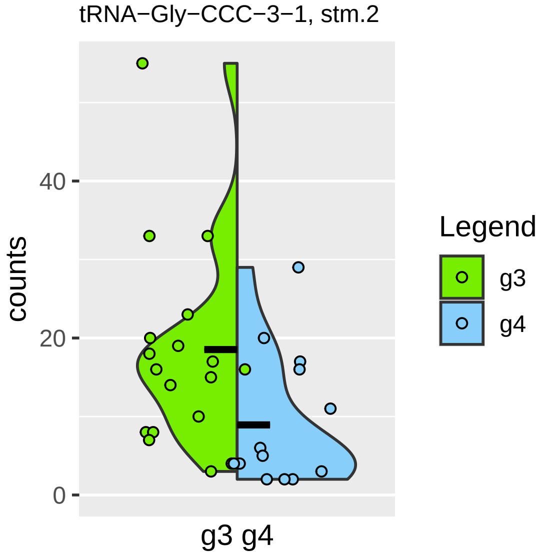

tRNA−Gly−CCC−3−1, stm.2
counts
0
20
40
g3 g4
Legend
g3
g4

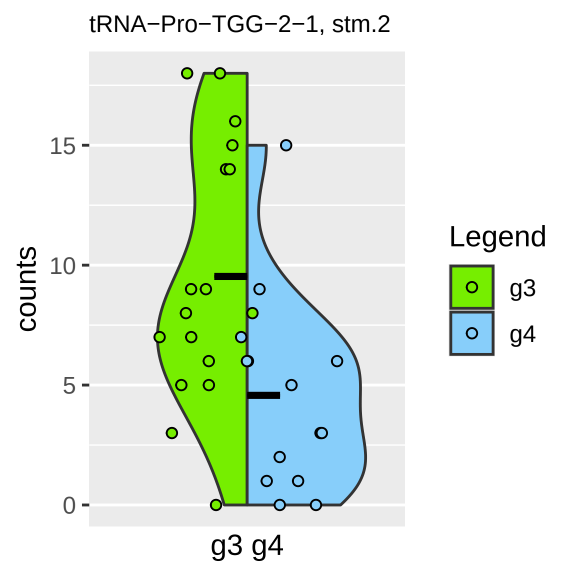

tRNA−Pro−TGG−2−1, stm.2
counts
0
5
10
15
g3 g4
Legend
g3
g4

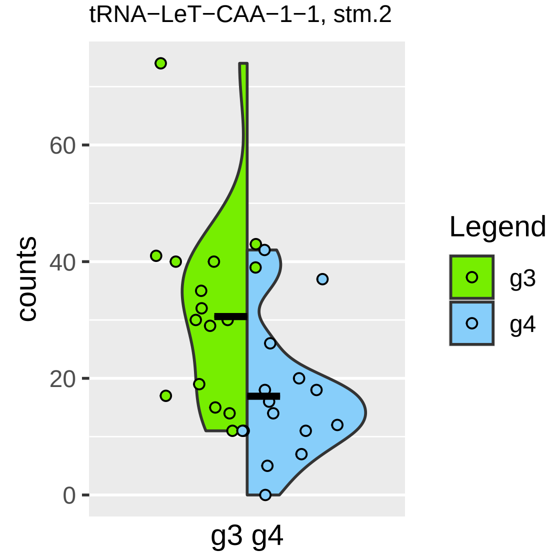
tRNA−LeT−CAA−1−1, stm.2
counts
60
40
20
0
g3 g4
Legend
g3
g4

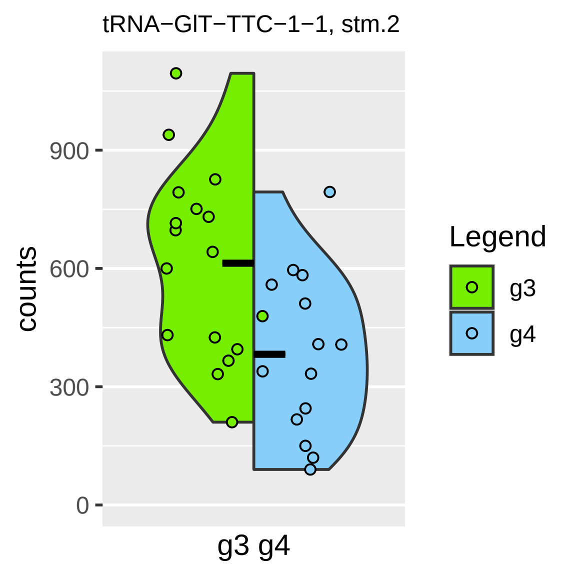

tRNA−GlT−TTC−1−1, stm.2
counts
900
600
300
0
g3 g4
Legend
g3
g4

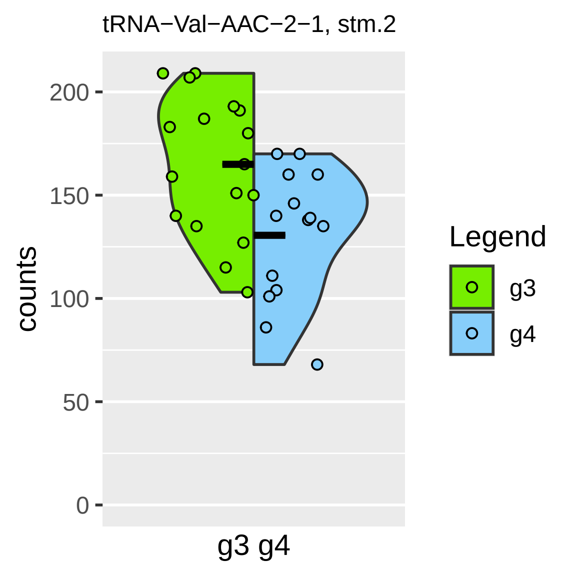

tRNA−Val−AAC−2−1, stm.2
counts
200
150
100
50
0
g3 g4
Legend
g3
g4

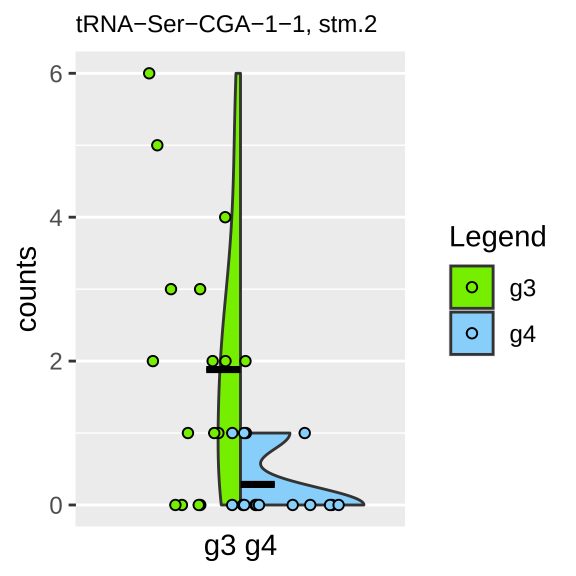

tRNA−Ser−CGA−1−1, stm.2
counts
6
4
2
0
g3 g4
Legend
g3
g4

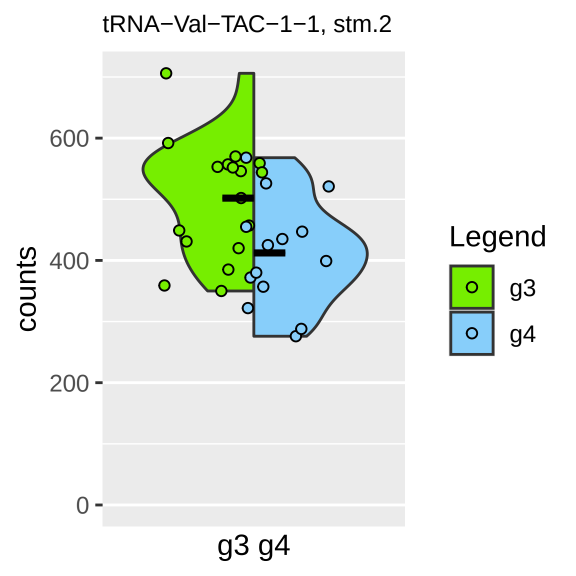

tRNA−Val−TAC−1−1, stm.2
counts
600
400
200
0
g3 g4
Legend
g3
g4

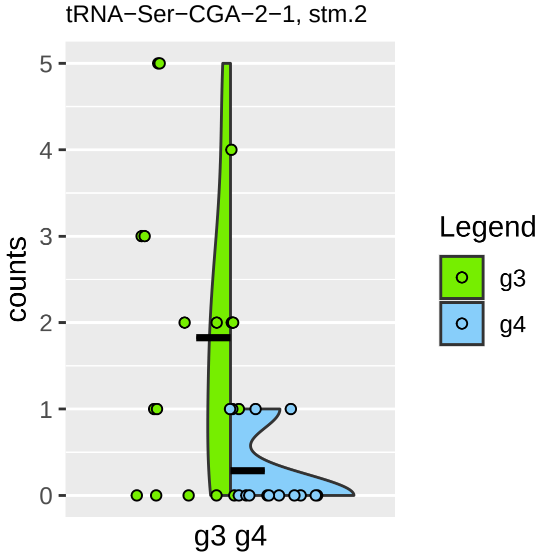
tRNA−Ser−CGA−2−1, stm.2
counts
5
4
3
2
1
0
g3 g4
Legend
g3
g4

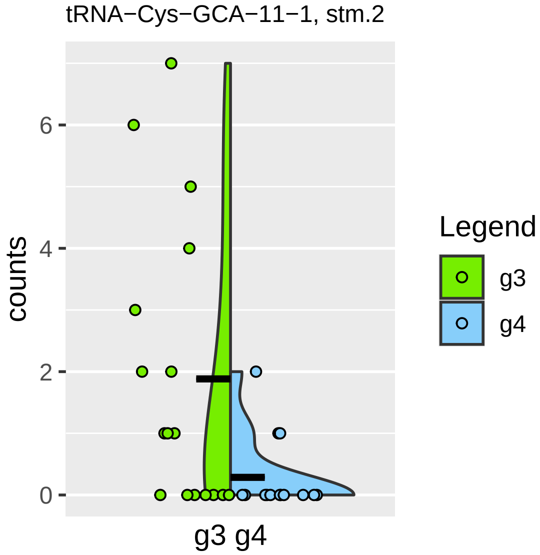

tRNA−Cys−GCA−11−1, stm.2
counts
0
2
4
6
g3 g4
Legend
g3
g4

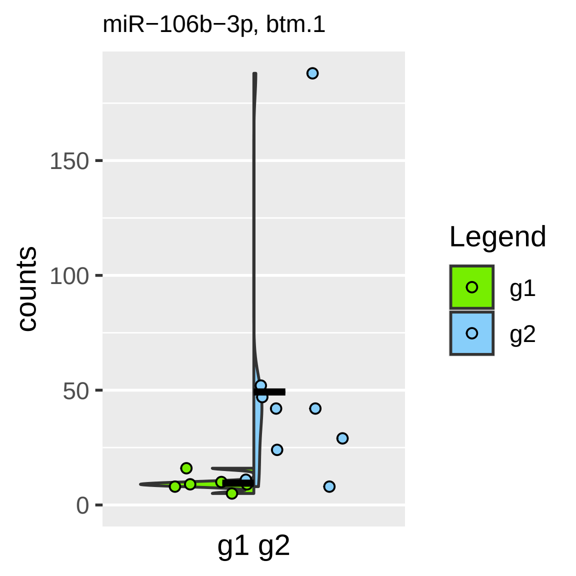
miR−106b−3p, btm.1
counts
150
100
50
0
g1 g2
Legend
g1
g2

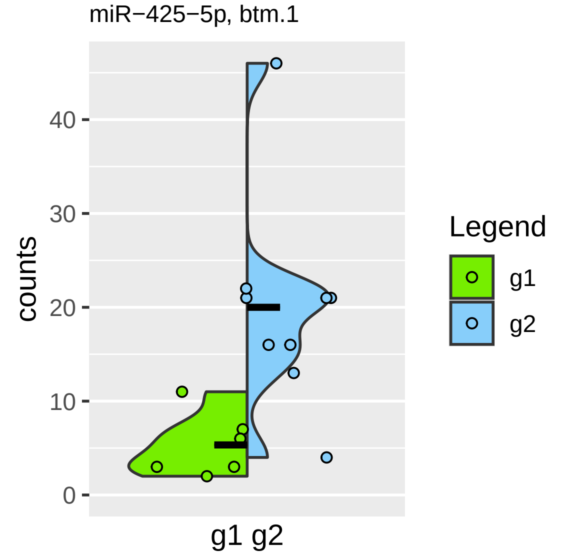

miR−425−5p, btm.1
counts
0
10
20
30
40
g1 g2
Legend
g1
g2

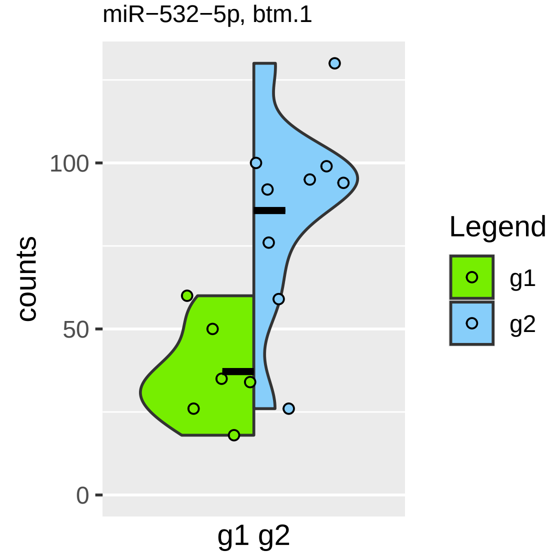
miR−532−5p, btm.1
counts
100
50
0
g1 g2
Legend
g1
g2

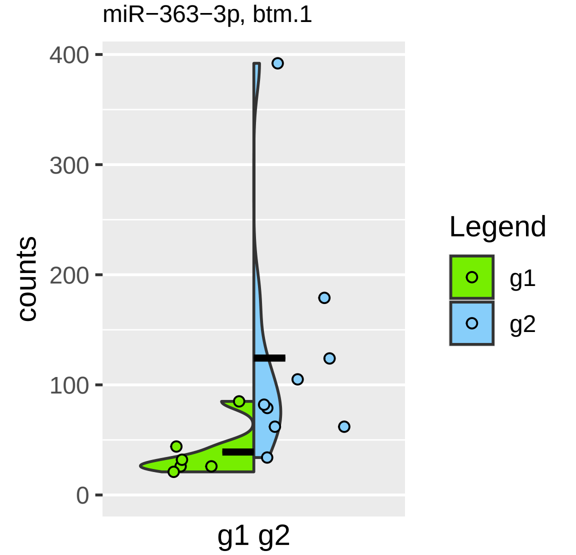

miR−363−3p, btm.1
400
300
200
100
0
counts
g1 g2
Legend
g1
g2

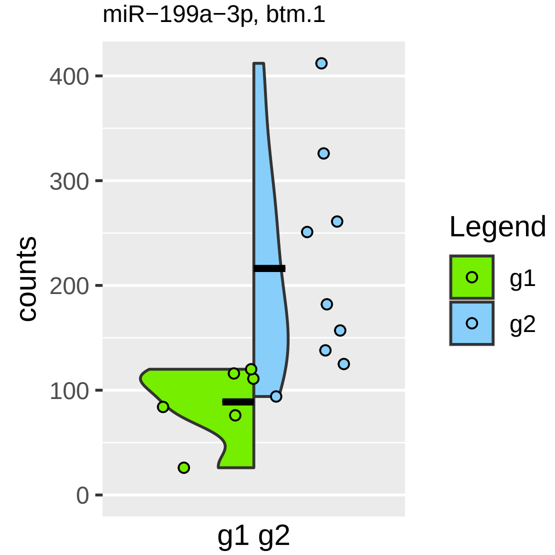

miR−199a−3p, btm.1
counts
400
300
200
100
0
g1 g2
Legend
g1
g2

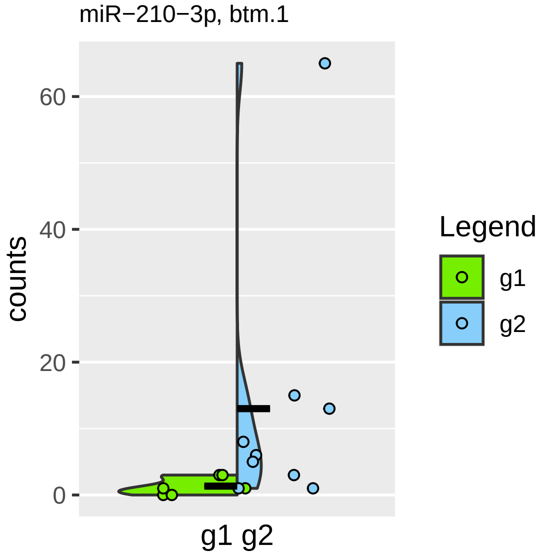
miR−210−3p, btm.1
counts
60
40
20
0
g1 g2
Legend
g1
g2

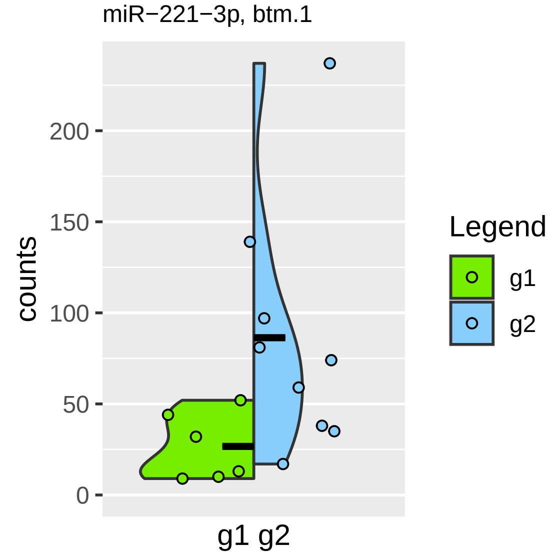

miR−221−3p, btm.1
counts
0
50
100
150
200
g1 g2
Legend
g1
g2

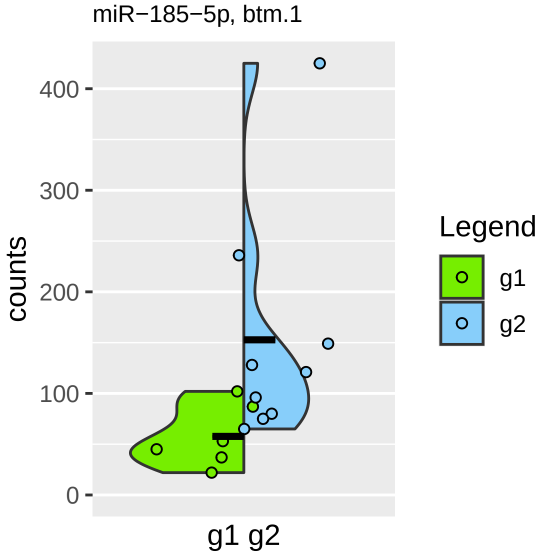

miR−185−5p, btm.1
counts
400
300
200
100
0
g1 g2
Legend
g1
g2

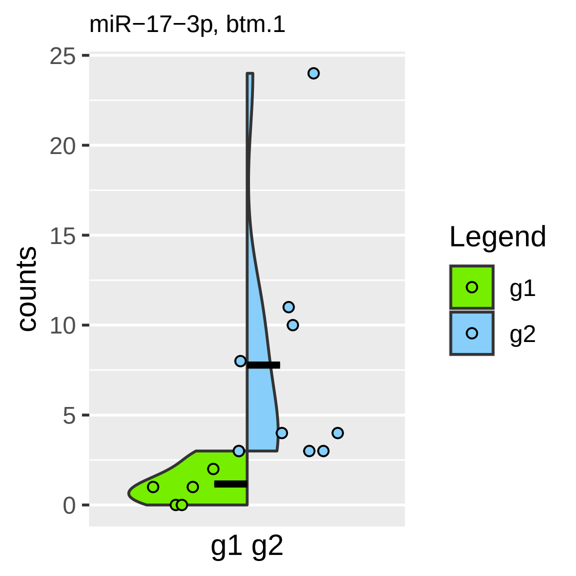
miR−17−3p, btm.1
counts
25
20
15
10
5
0
g1 g2
Legend
g1
g2

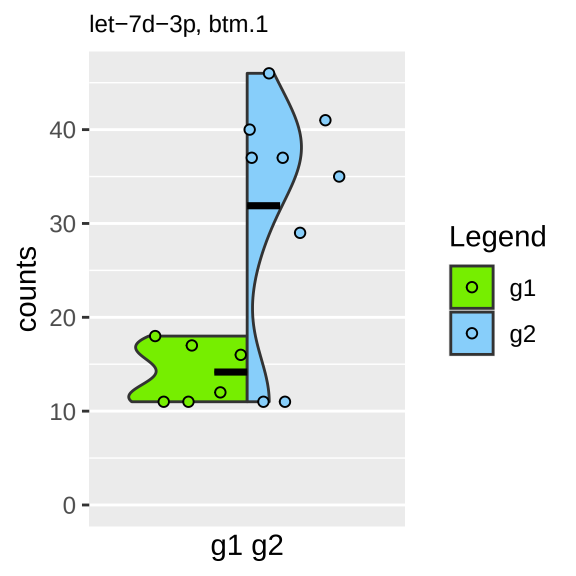

let−7d−3p, btm.1
counts
0
10
20
30
40
g1 g2
Legend
g1
g2

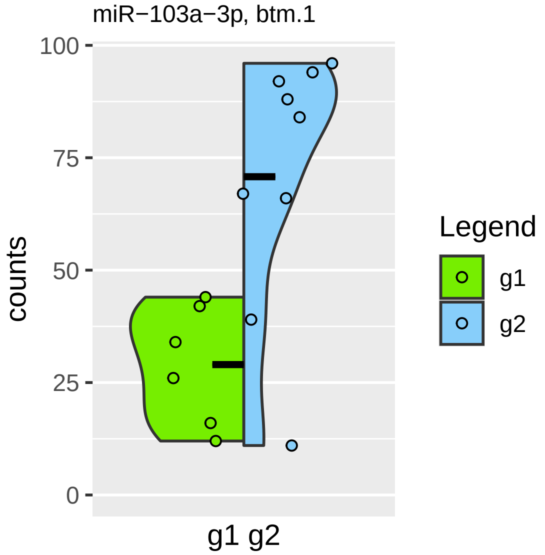
miR−103a−3p, btm.1
100
75
50
25
0
counts
g1 g2
Legend
g1
g2

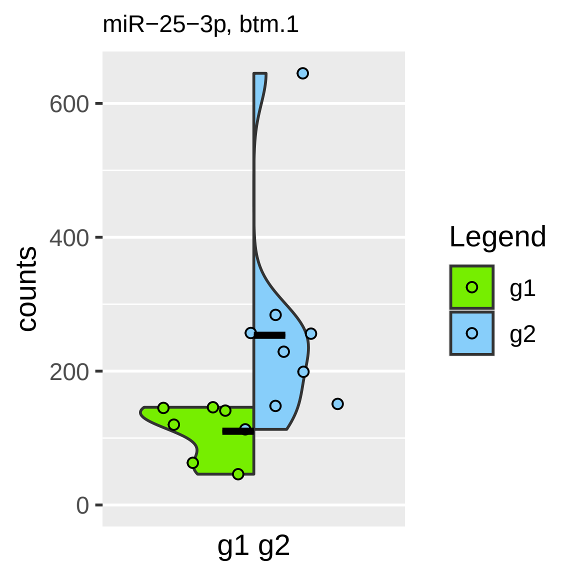

miR−25−3p, btm.1
counts
600
400
200
0
g1 g2
Legend
g1
g2

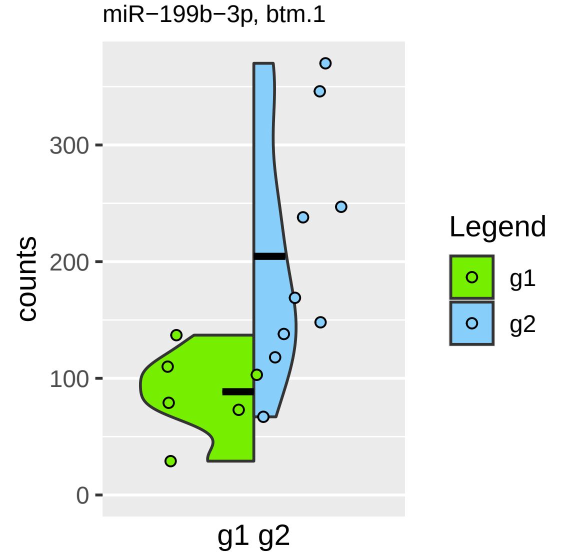

miR−199b−3p, btm.1
counts
300
200
100
0
g1 g2
Legend
g1
g2

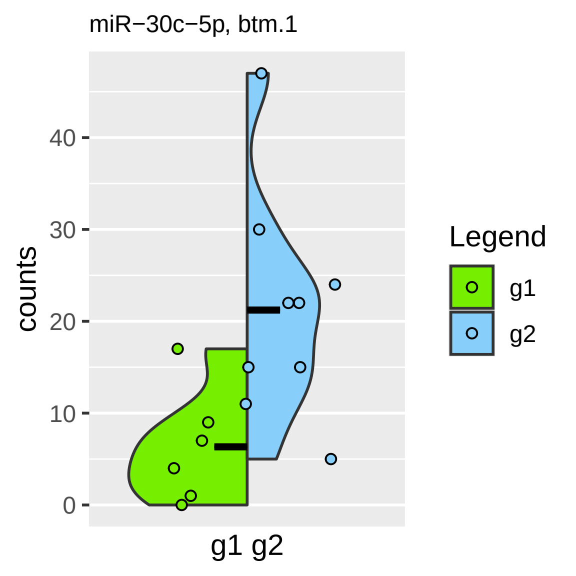

miR−30c−5p, btm.1
counts
0
10
20
30
40
g1 g2
Legend
g1
g2

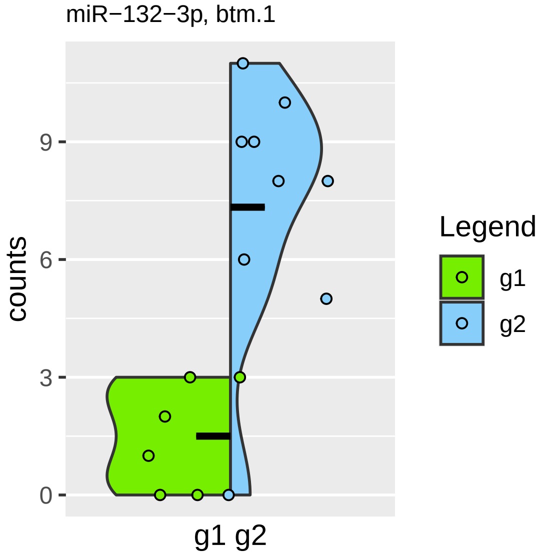
miR−132−3p, btm.1
counts
0
3
6
9
g1 g2
Legend
g1
g2

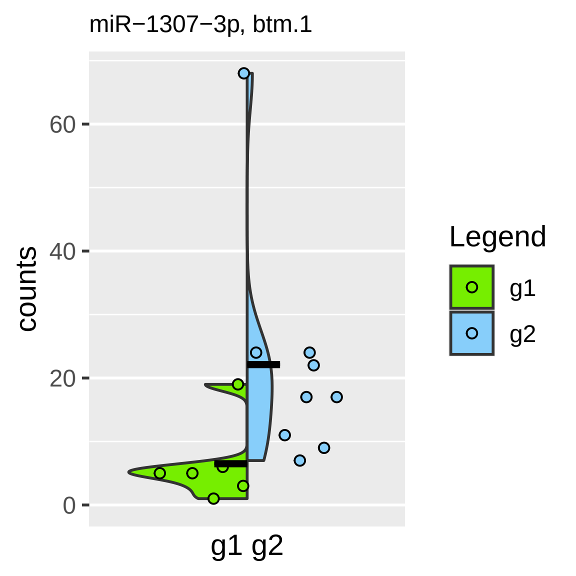

miR−1307−3p, btm.1
counts
0
20
40
60
g1 g2
Legend
g1
g2

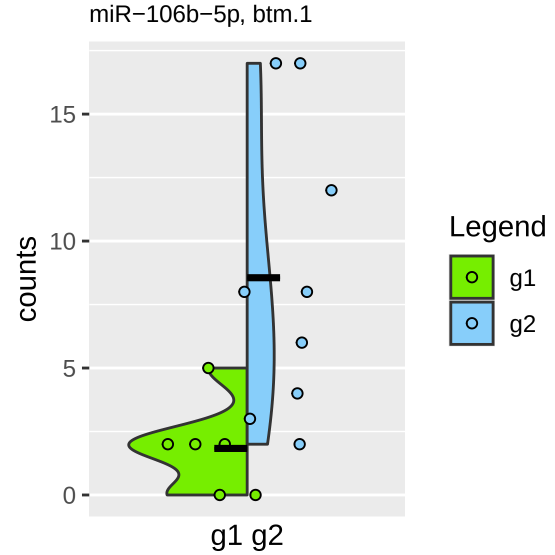

miR−106b−5p, btm.1
counts
0
5
10
15
g1 g2
Legend
g1
g2

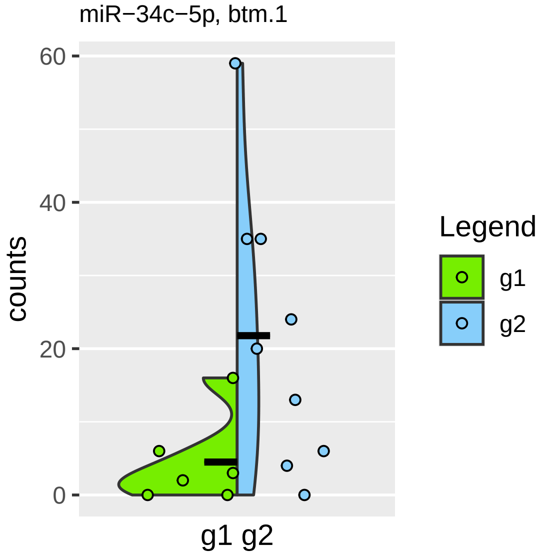

miR−34c−5p, btm.1
60
40
20
0
counts
g1 g2
Legend
g1
g2

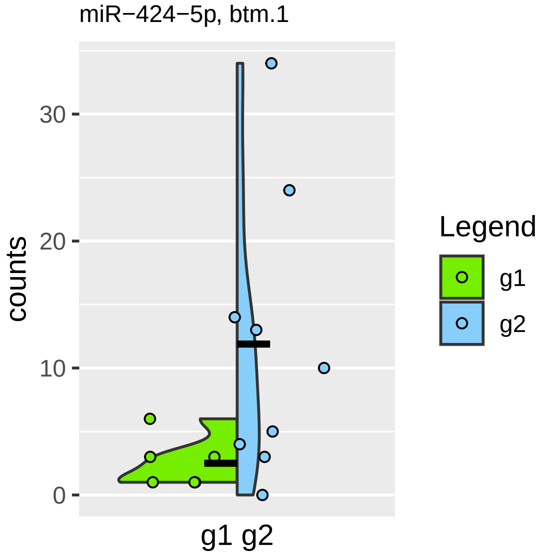
miR−424−5p, btm.1
counts
30
20
10
0
g1 g2
Legend
g1
g2

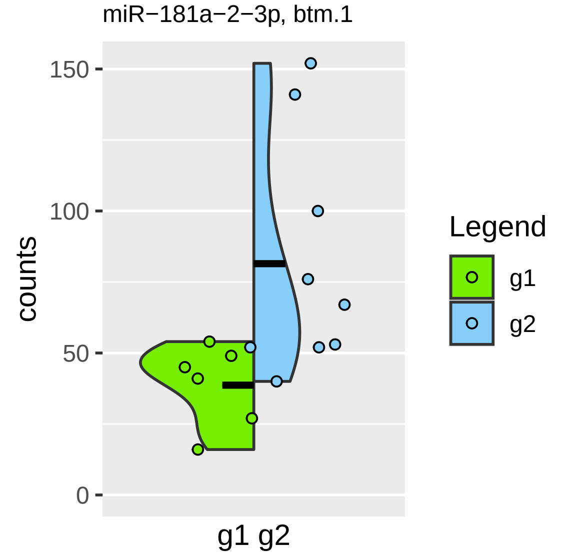

miR−181a−2−3p, btm.1
150
100
50
0
counts
g1 g2
Legend
g1
g2

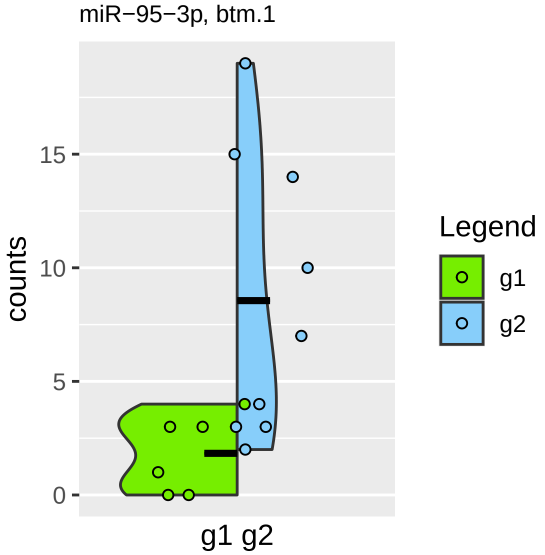

miR−95−3p, btm.1
counts
0
5
10
15
g1 g2
Legend
g1
g2

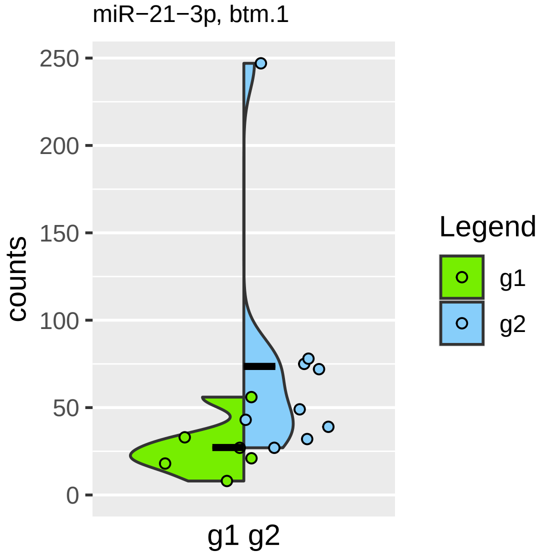

miR−21−3p, btm.1
counts
250
200
150
100
50
0
g1 g2
Legend
g1
g2

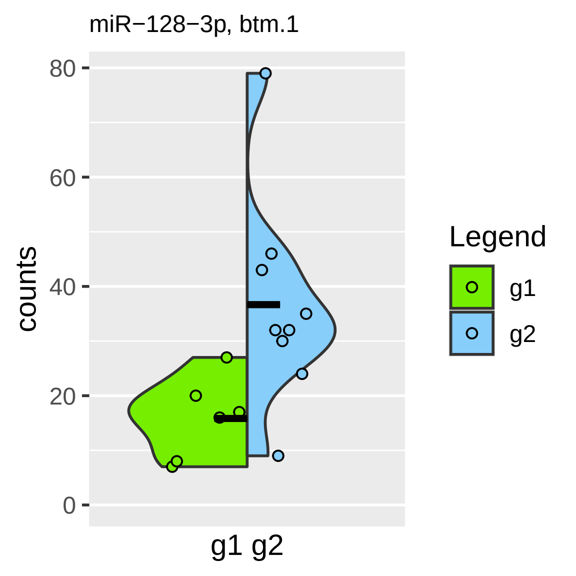

miR−128−3p, btm.1
counts
0
20
40
60
80
g1 g2
Legend
g1
g2

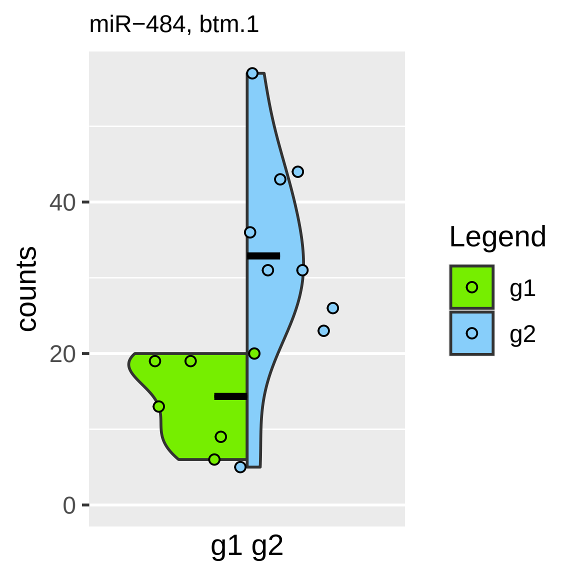
miR−484, btm.1
counts
0
20
40
g1 g2
Legend
g1
g2

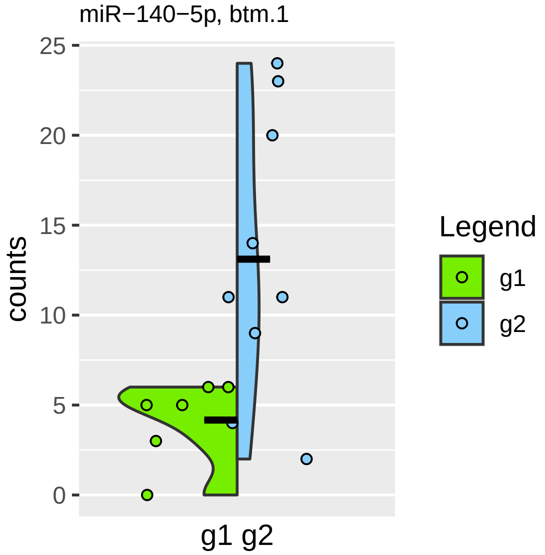

miR−140−5p, btm.1
counts
25
20
15
10
5
0
g1 g2
Legend
g1
g2

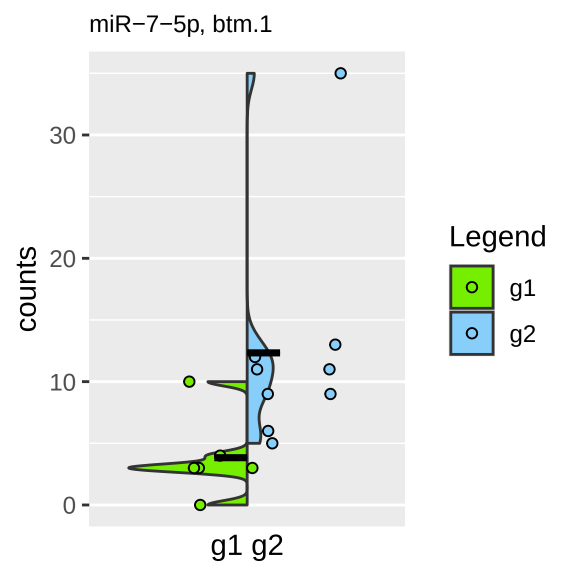
miR−7−5p, btm.1
counts
30
20
10
0
g1 g2
Legend
g1
g2

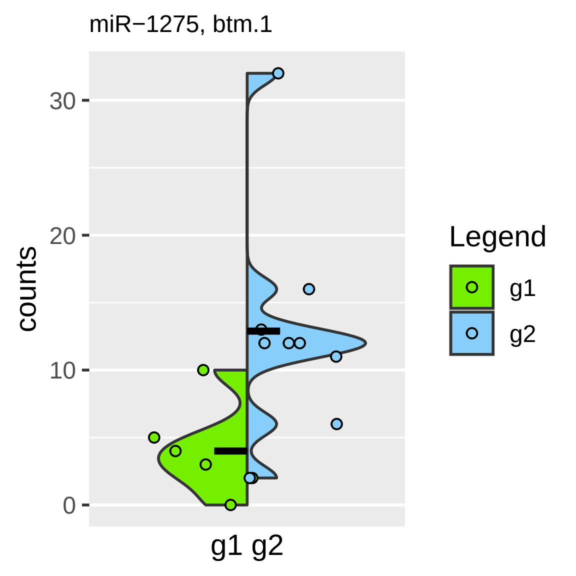

miR−1275, btm.1
counts
30
20
10
0
g1 g2
Legend
g1
g2

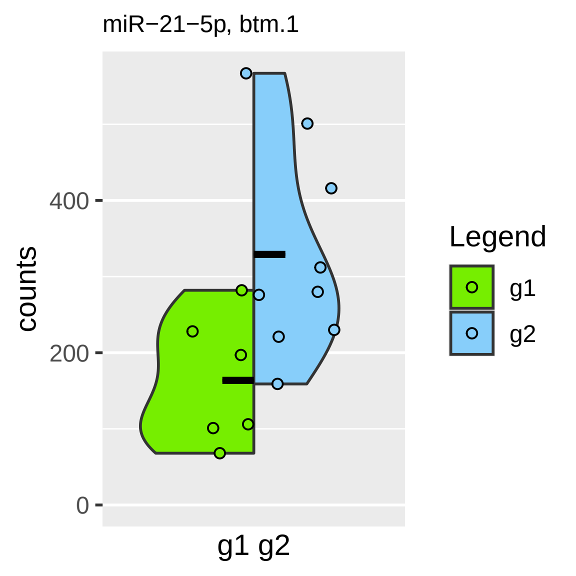

miR−21−5p, btm.1
counts
400
200
0
g1 g2
Legend
g1
g2

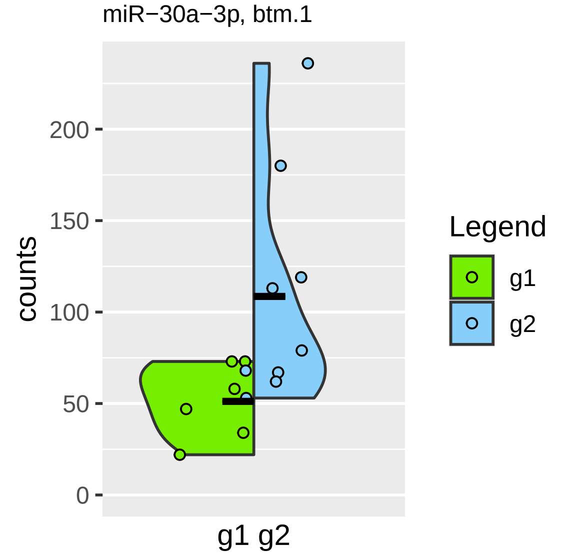

miR−30a−3p, btm.1
counts
200
150
100
50
0
g1 g2
Legend
g1
g2

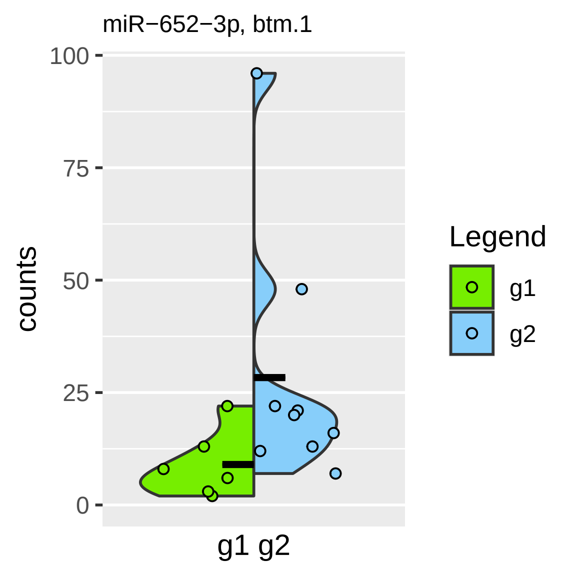
miR−652−3p, btm.1
100
75
50
25
0
counts
g1 g2
Legend
g1
g2

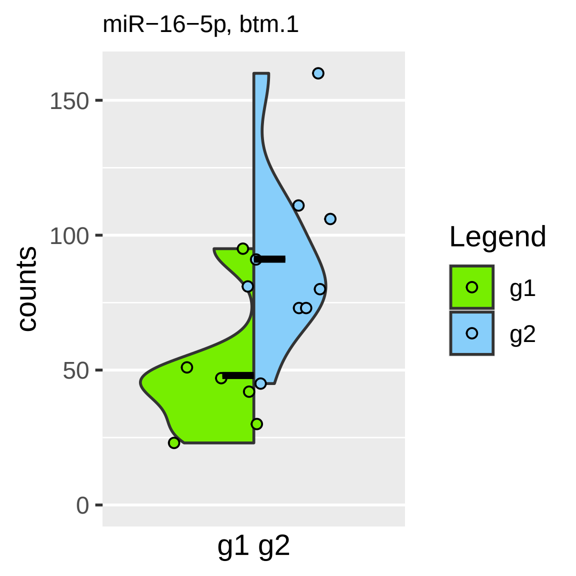
miR−16−5p, btm.1
counts
150
100
50
0
g1 g2
Legend
g1
g2

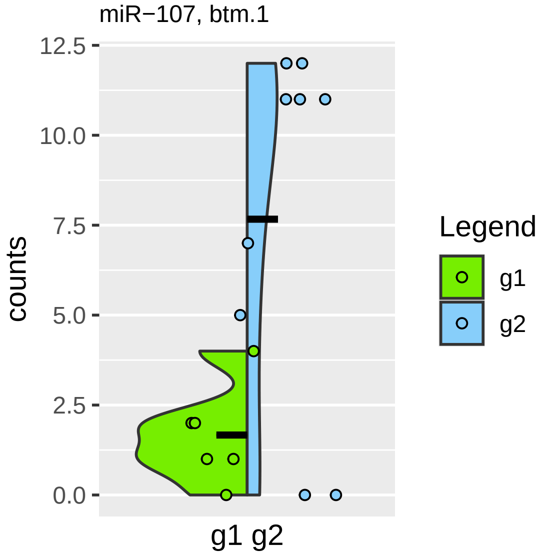
miR−107, btm.1
counts
12.5
10.0
7.5
5.0
2.5
0.0
g1 g2
Legend
g1
g2

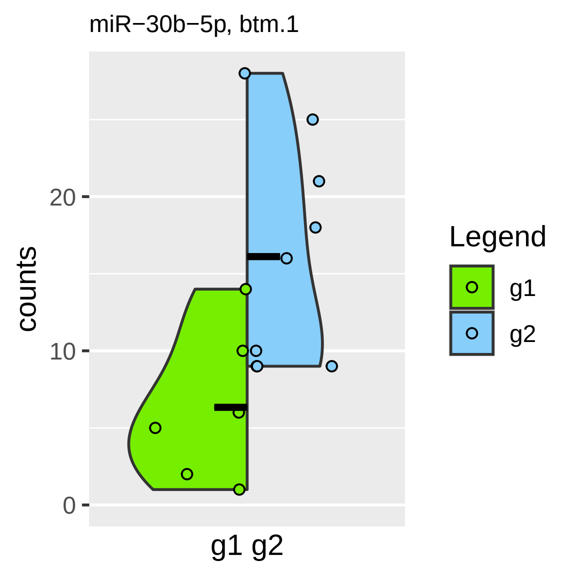
miR−30b−5p, btm.1
counts
20
10
0
g1 g2
Legend
g1
g2

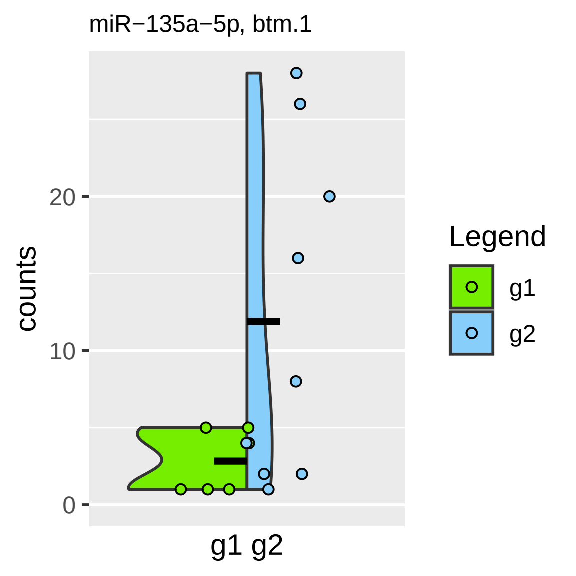
miR−135a−5p, btm.1
counts
20
10
0
g1 g2
Legend
g1
g2

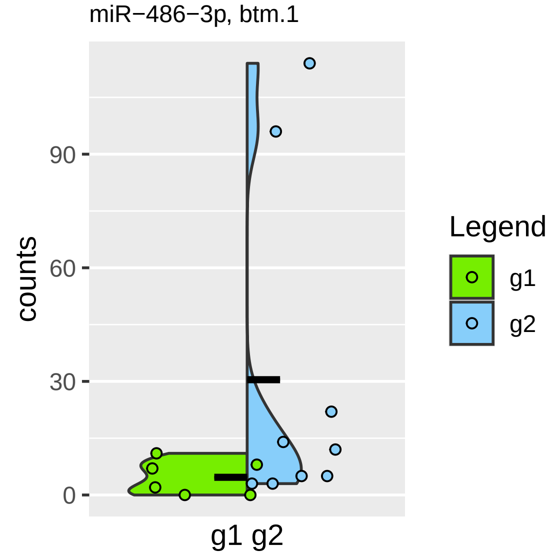
miR−486−3p, btm.1
counts
0
30
60
90
g1 g2
Legend
g1
g2

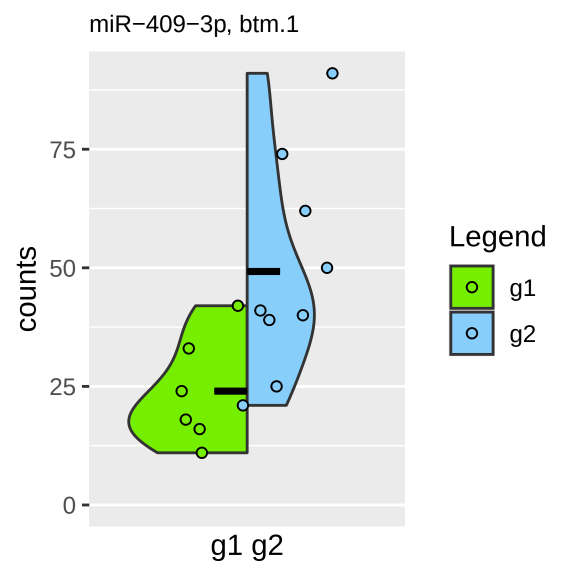

miR−409−3p, btm.1
counts
75
50
25
0
g1 g2
Legend
g1
g2

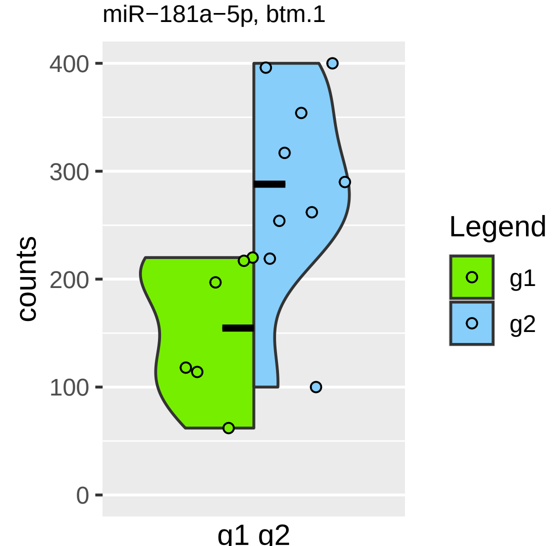

miR−181a−5p, btm.1
counts
400
300
200
100
0
g1 g2
Legend
g1
g2

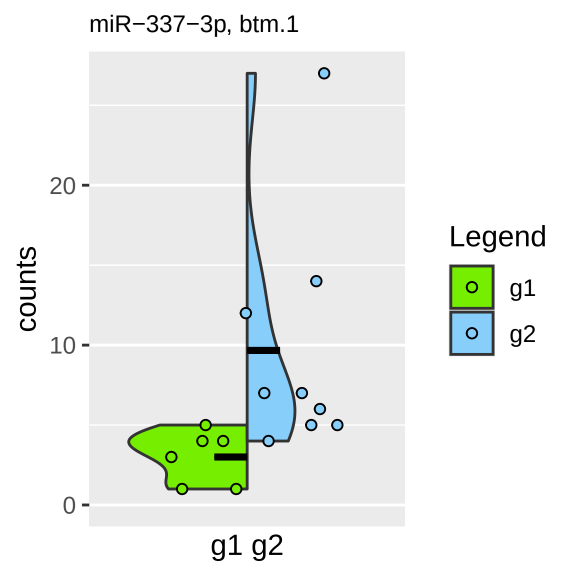

miR−337−3p, btm.1
counts
0
10
20
g1 g2
Legend
g1
g2

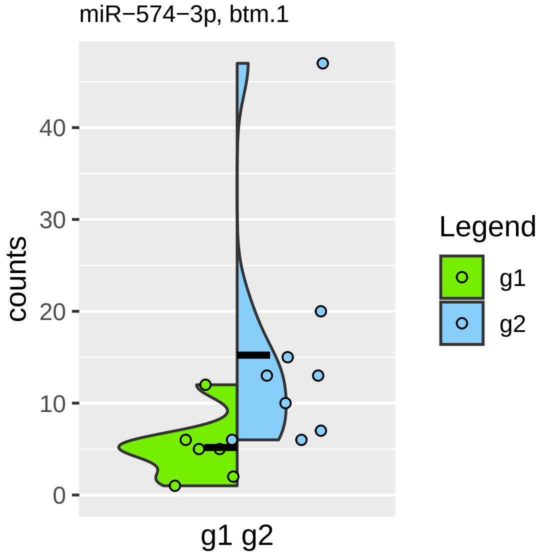

miR−574−3p, btm.1
counts
0
10
20
30
40
g1 g2
Legend
g1
g2

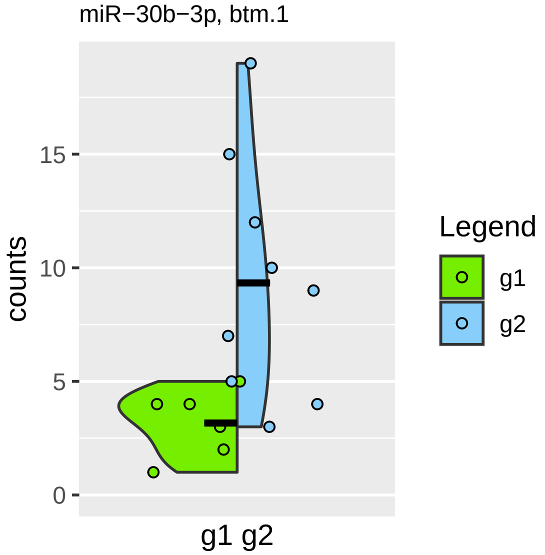

miR−30b−3p, btm.1
counts
15
10
5
0
g1 g2
Legend
g1
g2

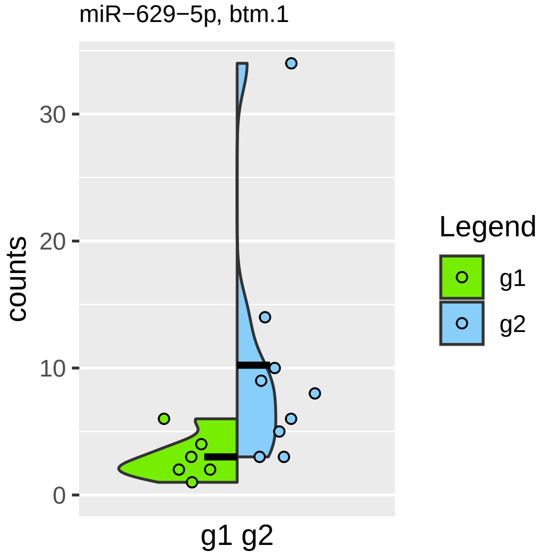

miR−629−5p, btm.1
counts
30
20
10
0
g1 g2
Legend
g1
g2

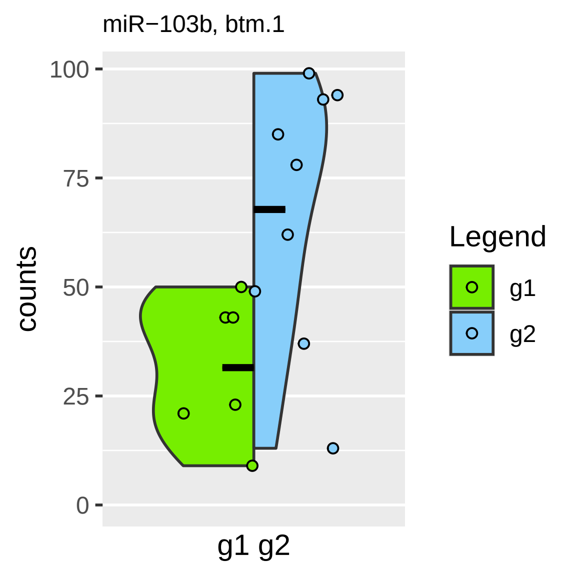
miR−103b, btm.1
100
75
50
25
0
counts
g1 g2
Legend
g1
g2

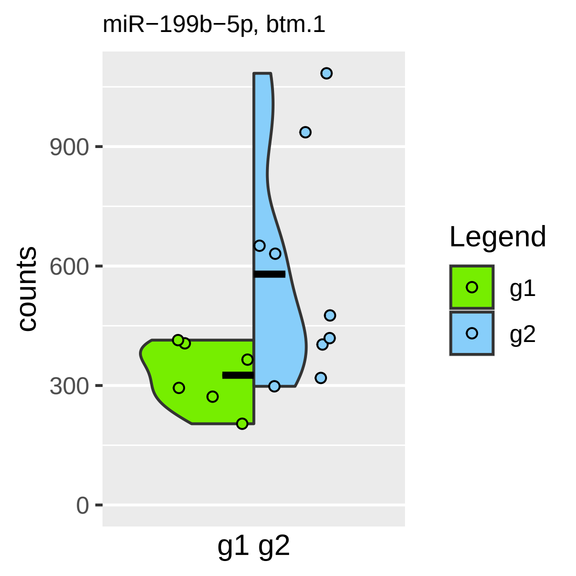

miR−199b−5p, btm.1
counts
900
600
300
0
g1 g2
Legend
g1
g2

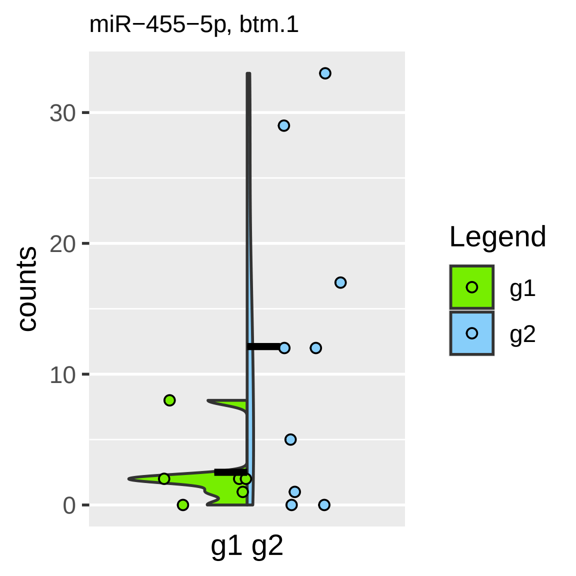
miR−455−5p, btm.1
counts
0
10
20
30
g1 g2
Legend
g1
g2

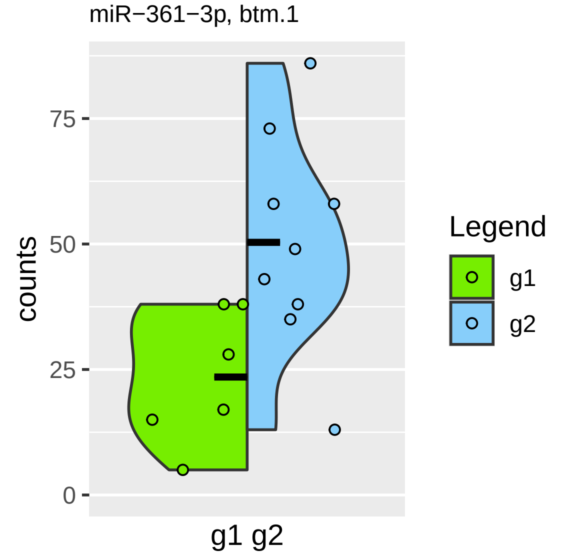

miR−361−3p, btm.1
counts
75
50
25
0
g1 g2
Legend
g1
g2

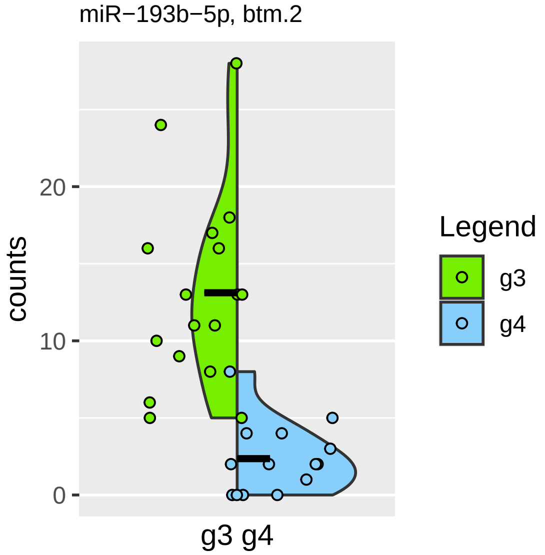

miR−193b−5p, btm.2
counts
0
10
20
g3 g4
Legend
g3
g4

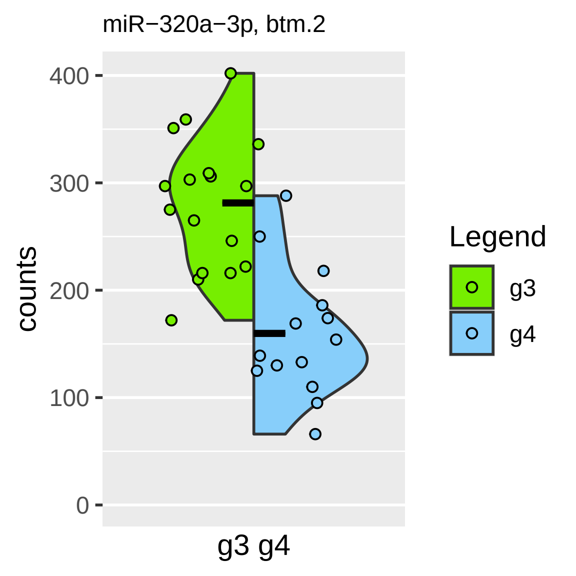

miR−320a−3p, btm.2
400
300
200
100
0
counts
g3 g4
Legend
g3
g4

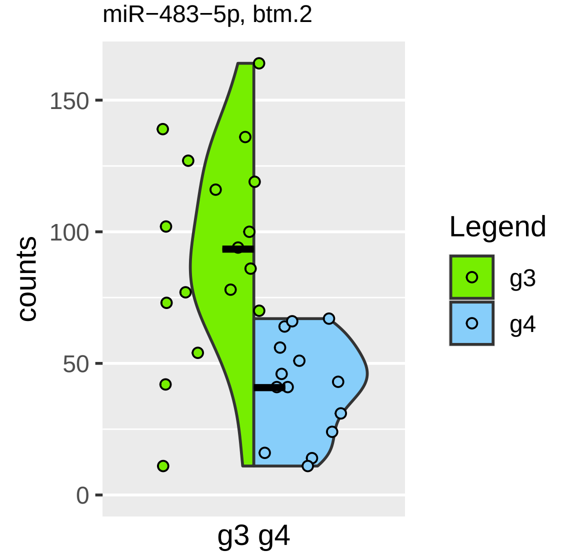

miR−483−5p, btm.2
counts
150
100
50
0
g3 g4
Legend
g3
g4

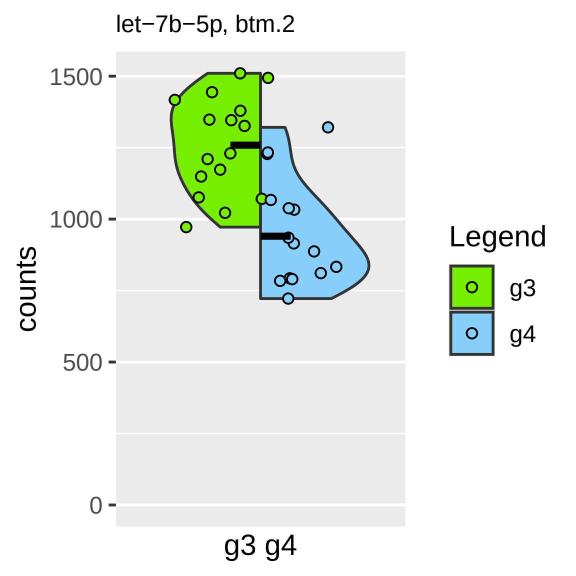

let−7b−5p, btm.2
1500
1000
500
0
counts
g3 g4
Legend
g3
g4

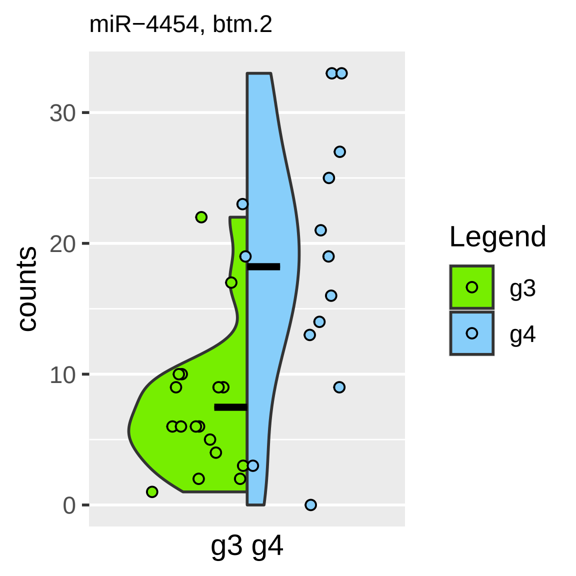

miR−4454, btm.2
counts
30
20
10
0
g3 g4
Legend
g3
g4

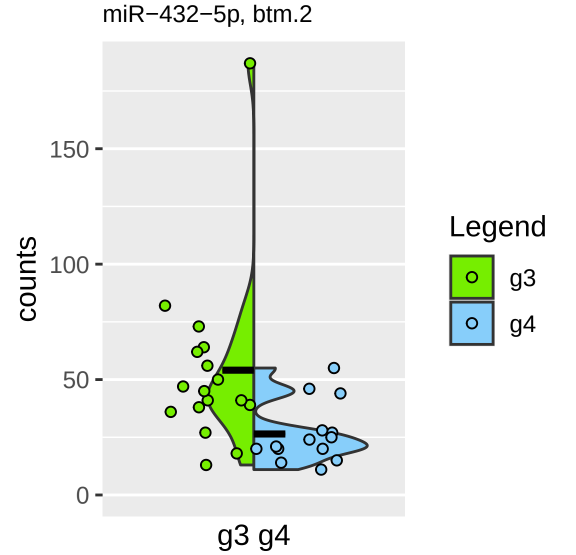

miR−432−5p, btm.2
counts
150
100
50
0
g3 g4
Legend
g3
g4

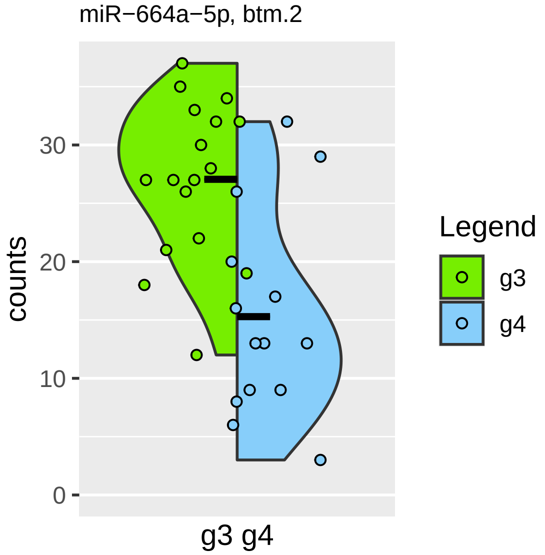
miR−664a−5p, btm.2
counts
0
10
20
30
g3 g4
Legend
g3
g4

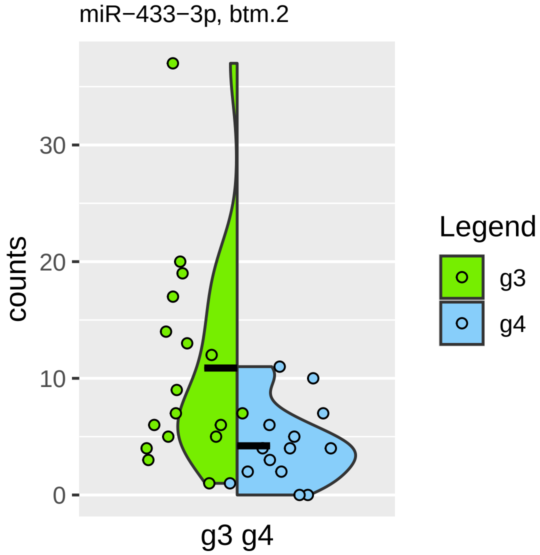

miR−433−3p, btm.2
counts
30
20
10
0
g3 g4
Legend
g3
g4

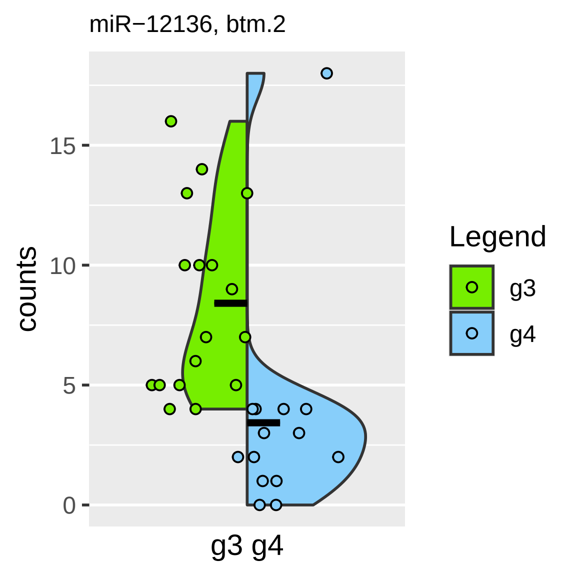

miR−12136, btm.2
counts
15
10
5
0
g3 g4
Legend
g3
g4

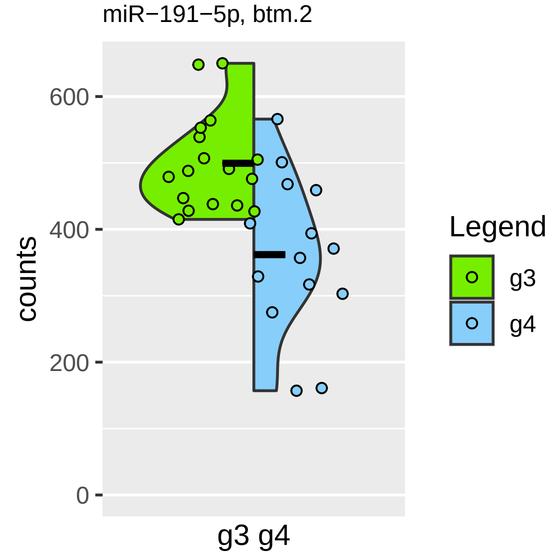
miR−191−5p, btm.2
600
400
200
0
counts
g3 g4
Legend
g3
g4

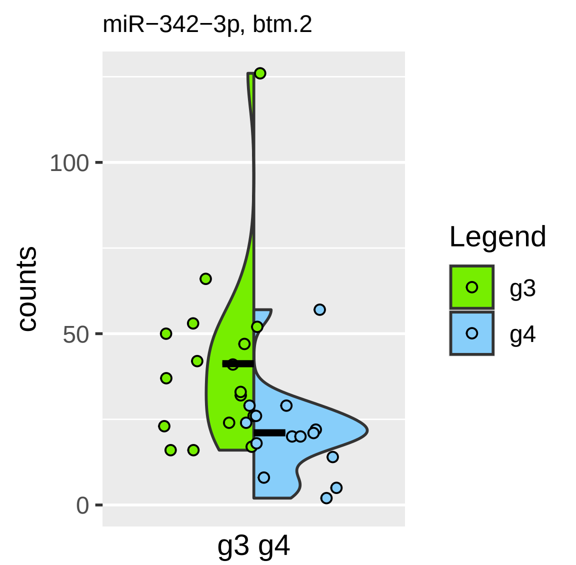
miR−342−3p, btm.2
counts
100
50
0
g3 g4
Legend
g3
g4

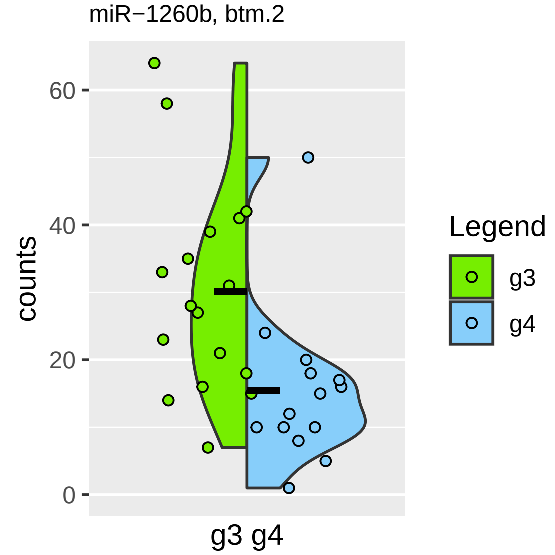

miR−1260b, btm.2
60
40
20
0
counts
g3 g4
Legend
g3
g4

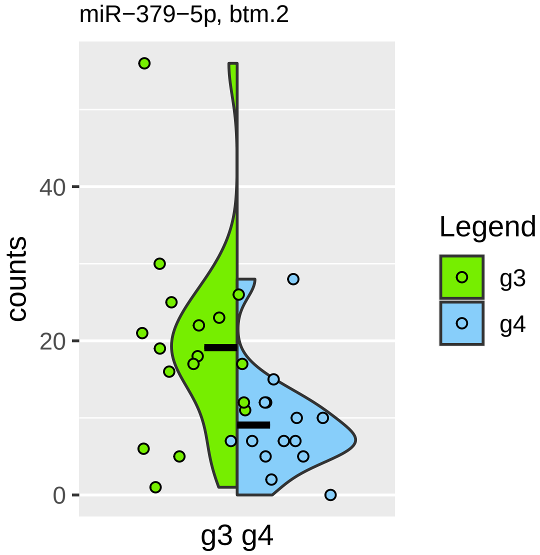

miR−379−5p, btm.2
counts
40
20
0
g3 g4
Legend
g3
g4

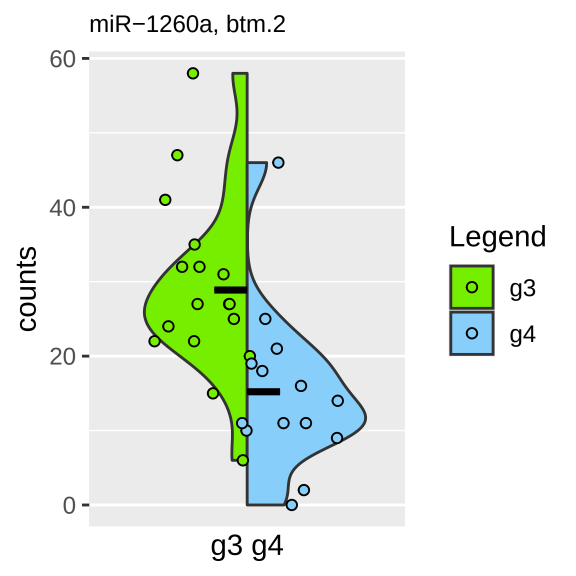

miR−1260a, btm.2
60
40
20
0
counts
g3 g4
Legend
g3
g4

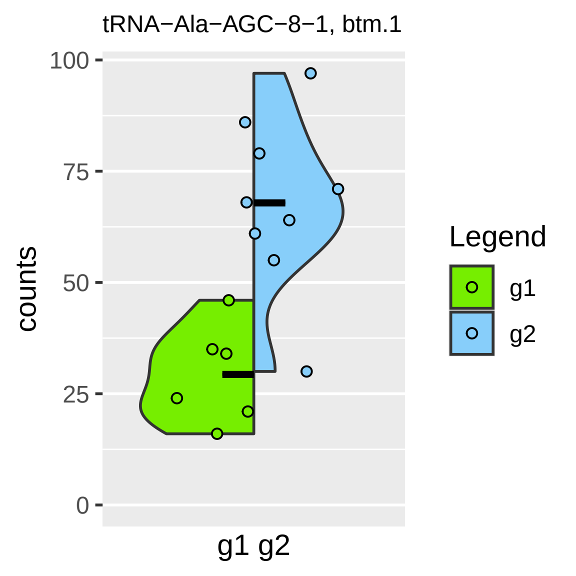
tRNA−Ala−AGC−8−1, btm.1
counts
100
75
50
25
0
g1 g2
Legend
g1
g2

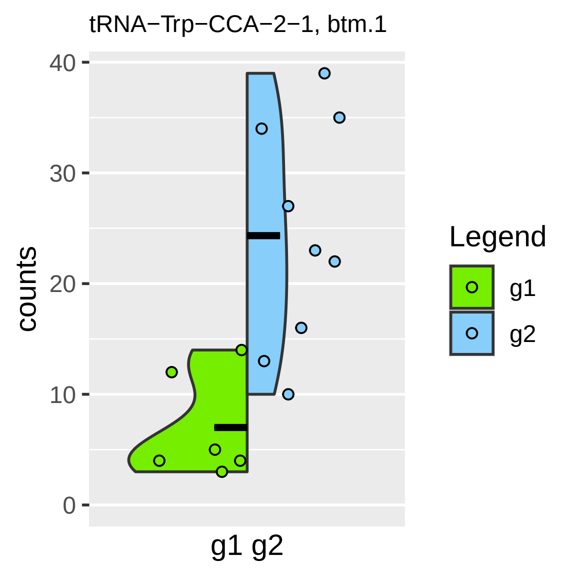
tRNA−Trp−CCA−2−1, btm.1
counts
40
30
20
10
0
g1 g2
Legend
g1
g2

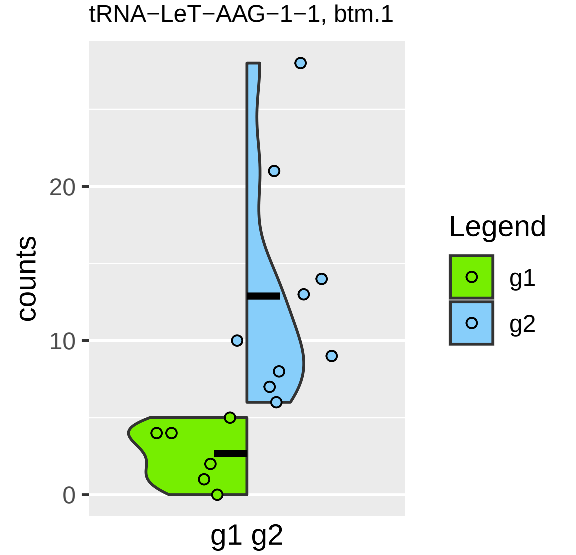

tRNA−LeT−AAG−1−1, btm.1
counts
0
10
20
g1 g2
Legend
g1
g2

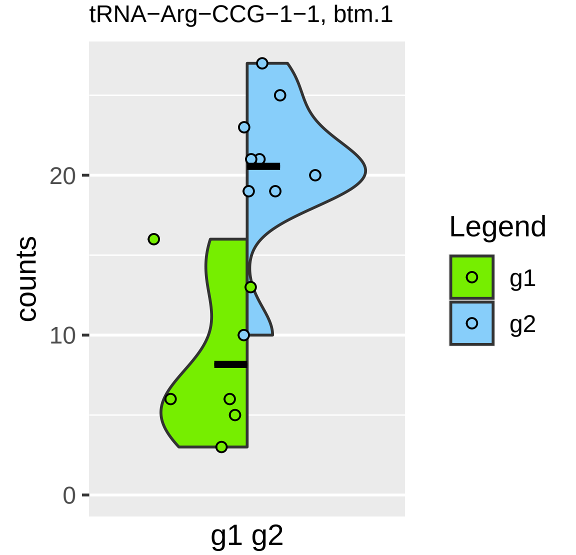

tRNA−Arg−CCG−1−1, btm.1
counts
0
10
20
g1 g2
Legend
g1
g2

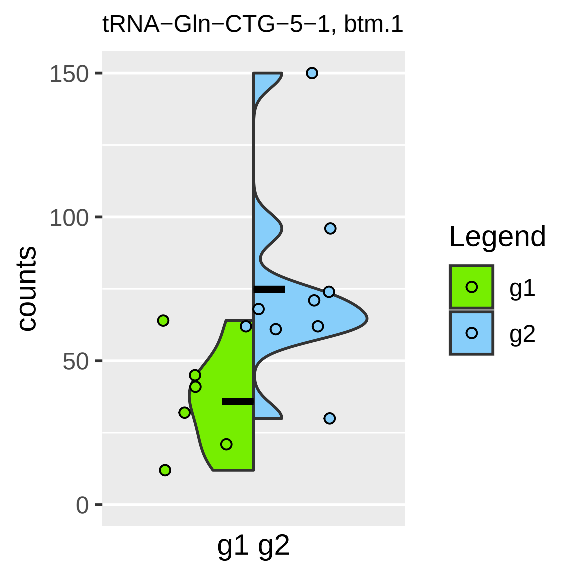
tRNA−Gln−CTG−5−1, btm.1
counts
150
100
50
0
g1 g2
Legend
g1
g2

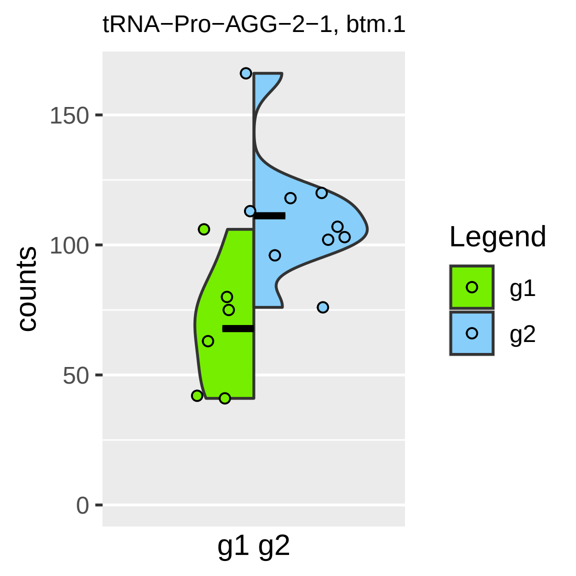

tRNA−Pro−AGG−2−1, btm.1
counts
150
100
50
0
g1 g2
Legend
g1
g2

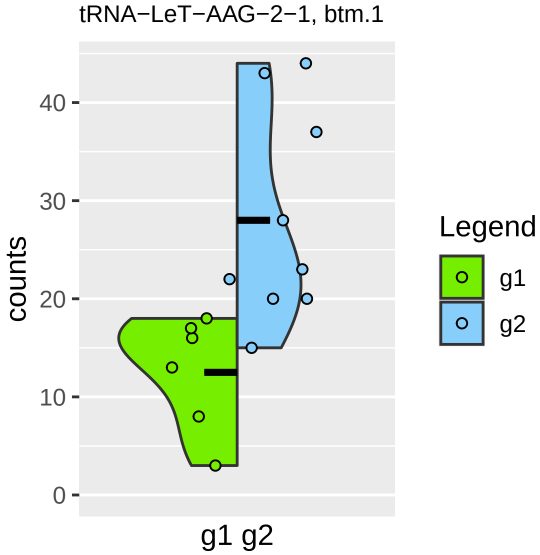

tRNA−LeT−AAG−2−1, btm.1
counts
0
10
20
30
40
g1 g2
Legend
g1
g2

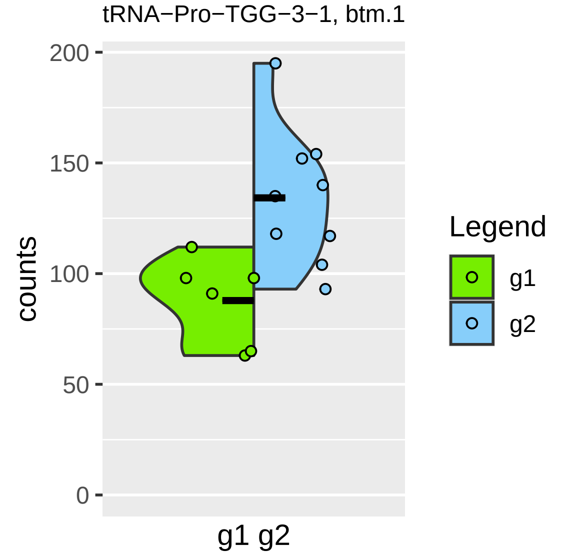
tRNA−Pro−TGG−3−1, btm.1
200
150
100
50
0
counts
g1 g2
Legend
g1
g2

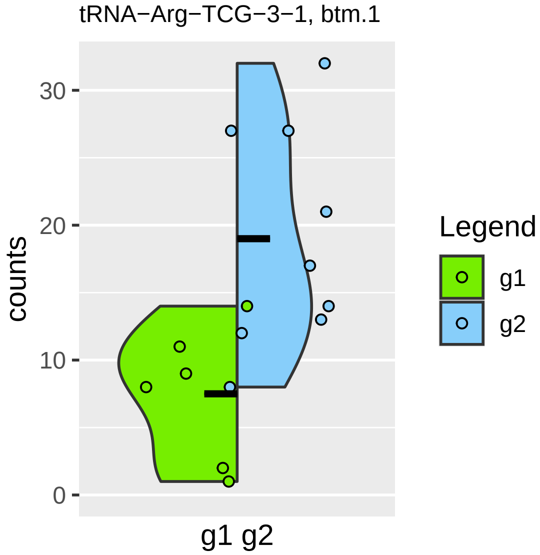

tRNA−Arg−TCG−3−1, btm.1
counts
0
10
20
30
g1 g2
Legend
g1
g2

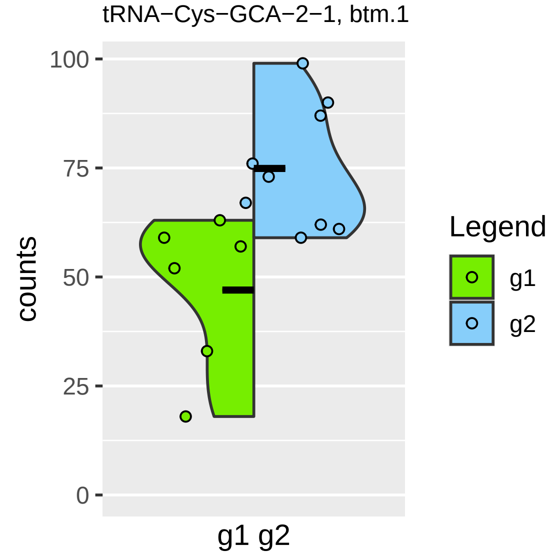

tRNA−Cys−GCA−2−1, btm.1
100
75
50
25
0
counts
g1 g2
Legend
g1
g2

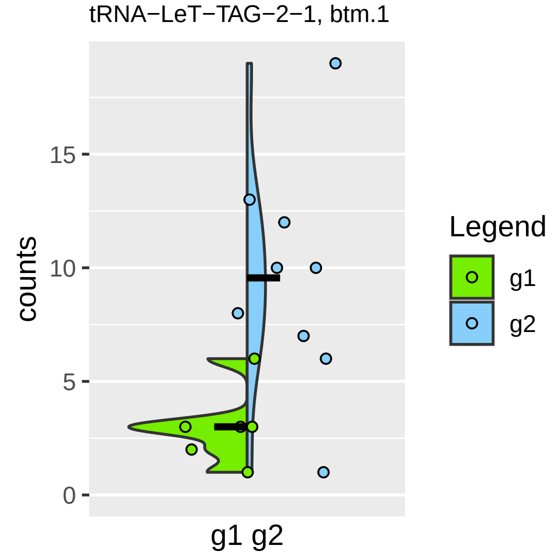

tRNA−LeT−TAG−2−1, btm.1
counts
0
5
10
15
g1 g2
Legend
g1
g2

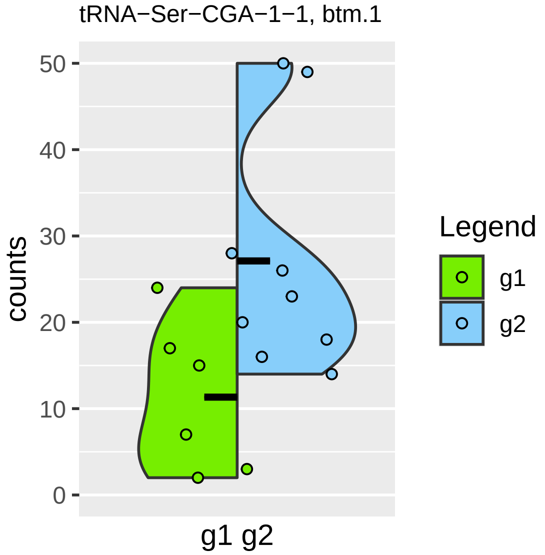

tRNA−Ser−CGA−1−1, btm.1
counts
50
40
30
20
10
0
g1 g2
Legend
g1
g2

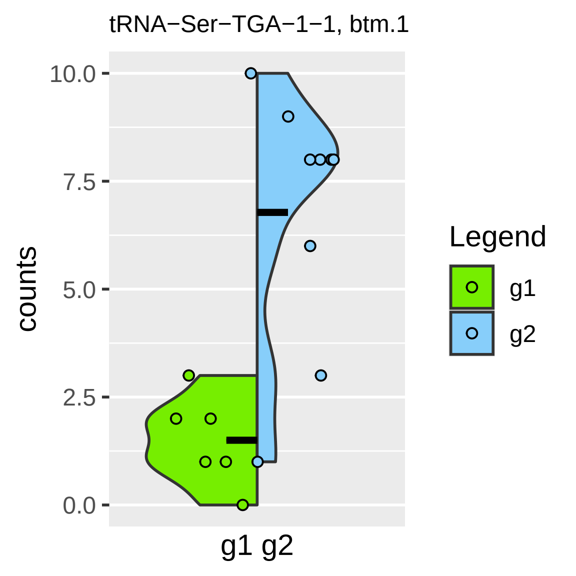
tRNA−Ser−TGA−1−1, btm.1
10.0
7.5
5.0
2.5
0.0
counts
g1 g2
Legend
g1
g2

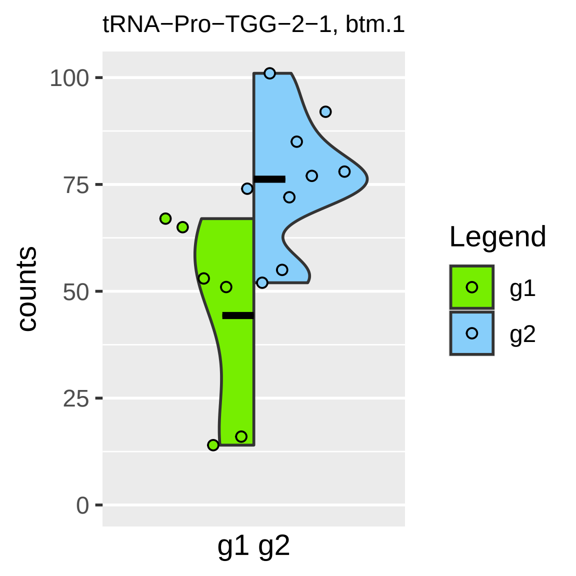

tRNA−Pro−TGG−2−1, btm.1
counts
100
75
50
25
0
g1 g2
Legend
g1
g2

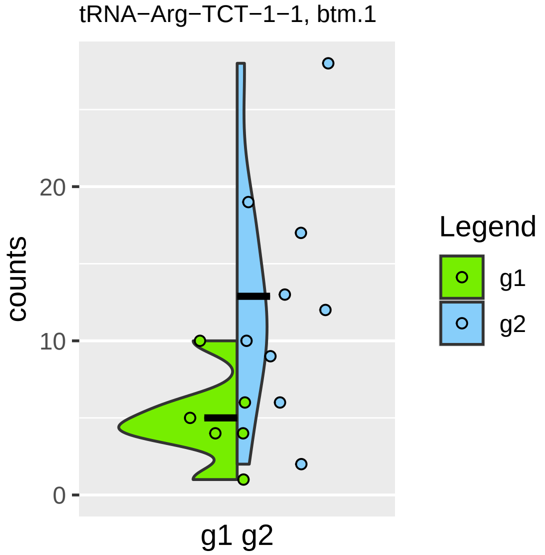

tRNA−Arg−TCT−1−1, btm.1
counts
20
10
0
g1 g2
Legend
g1
g2

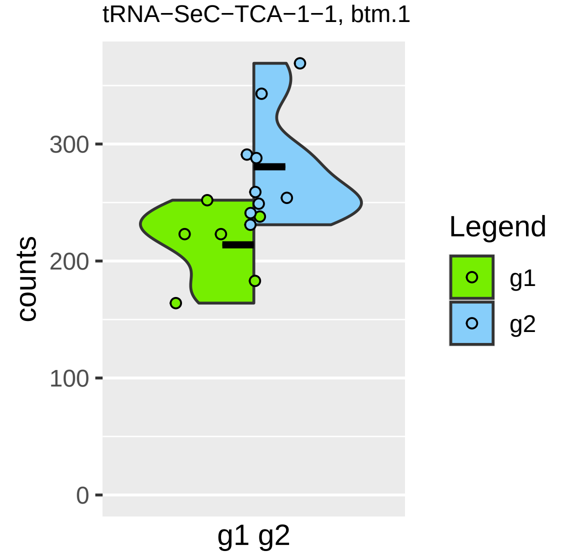
tRNA−SeC−TCA−1−1, btm.1
counts
300
200
100
0
g1 g2
Legend
g1
g2

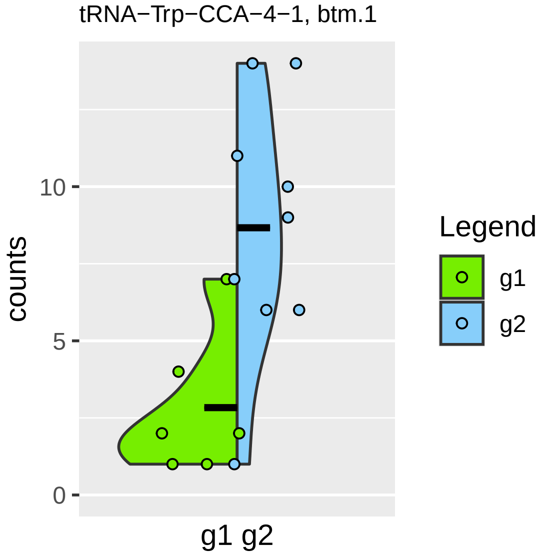

tRNA−Trp−CCA−4−1, btm.1
counts
0
5
10
g1 g2
Legend
g1
g2

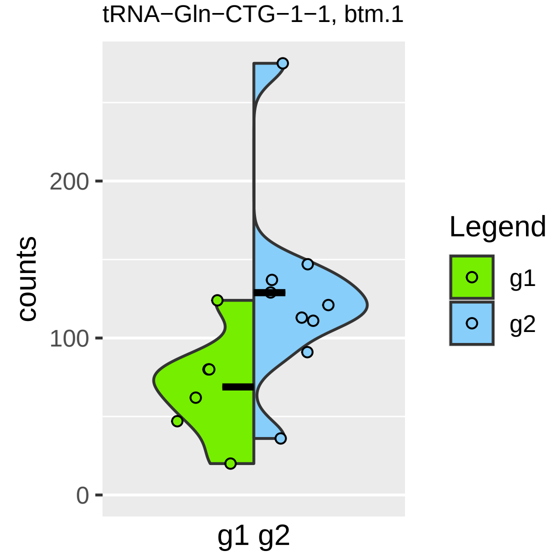

tRNA−Gln−CTG−1−1, btm.1
counts
200
100
0
g1 g2
Legend
g1
g2

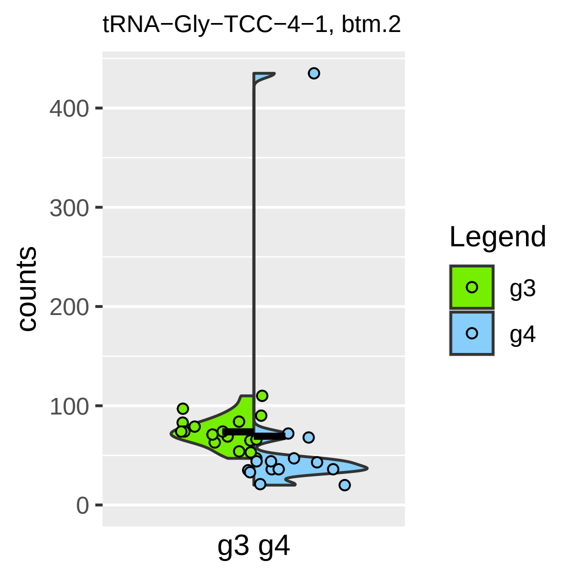

tRNA−Gly−TCC−4−1, btm.2
counts
0
100
200
300
400
g3 g4
Legend
g3
g4

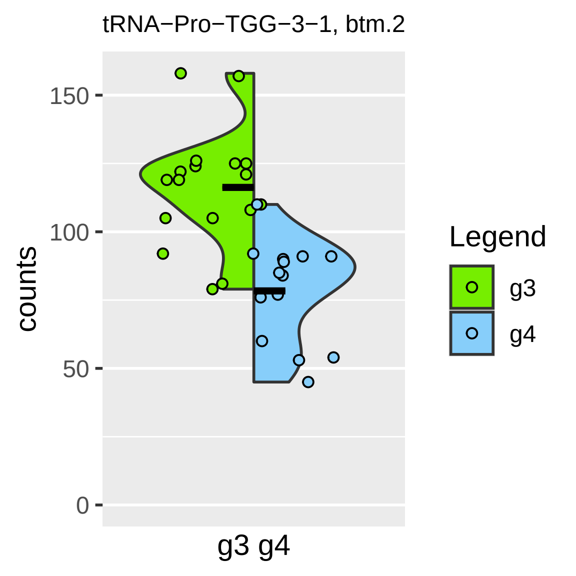
tRNA−Pro−TGG−3−1, btm.2
counts
150
100
50
0
g3 g4
Legend
g3
g4

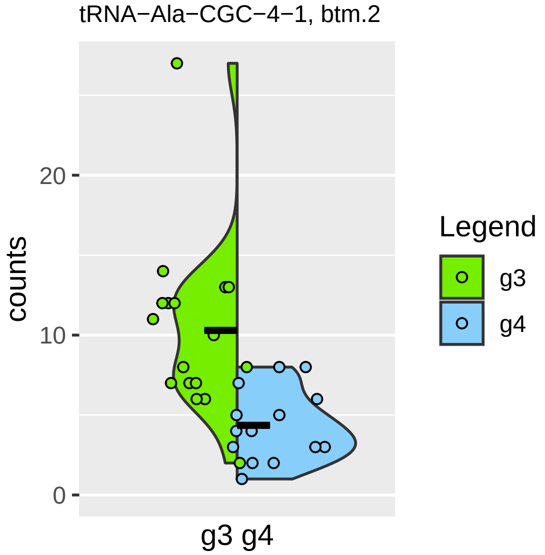

tRNA−Ala−CGC−4−1, btm.2
counts
20
10
0
g3 g4
Legend
g3
g4

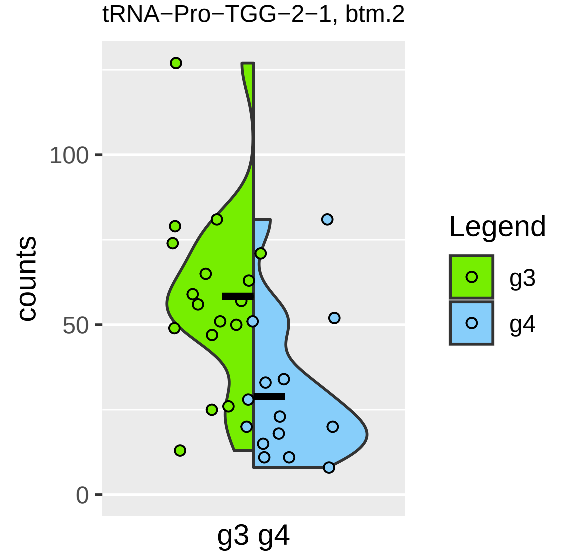
tRNA−Pro−TGG−2−1, btm.2
counts
100
50
0
g3 g4
Legend
g3
g4

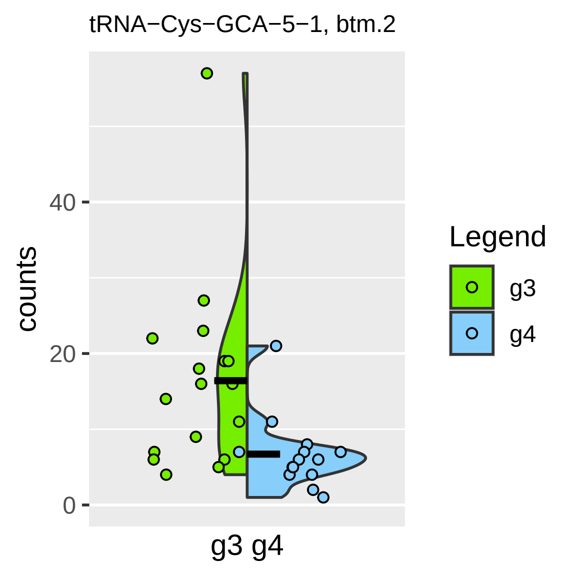

tRNA−Cys−GCA−5−1, btm.2
counts
0
20
40
g3 g4
Legend
g3
g4

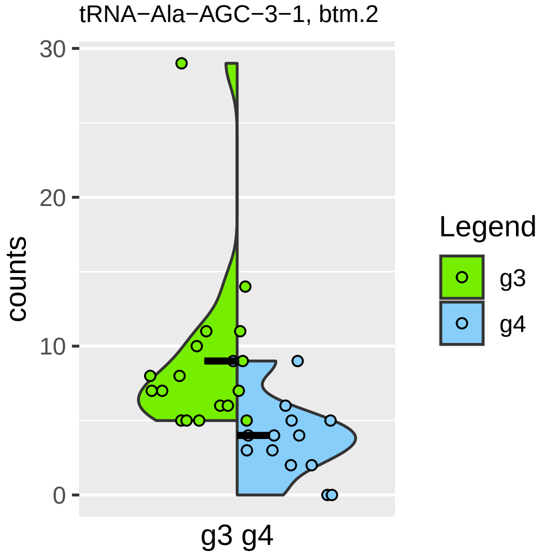

tRNA−Ala−AGC−3−1, btm.2
30
20
10
0
counts
g3 g4
Legend
g3
g4

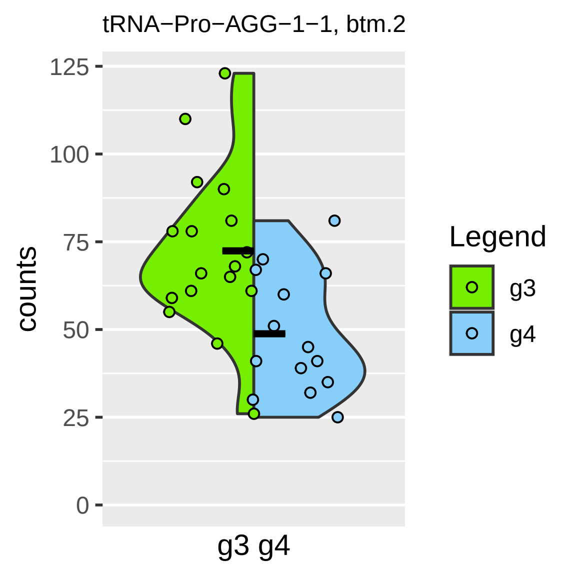

tRNA−Pro−AGG−1−1, btm.2
counts
125
100
75
50
25
0
g3 g4
Legend
g3
g4

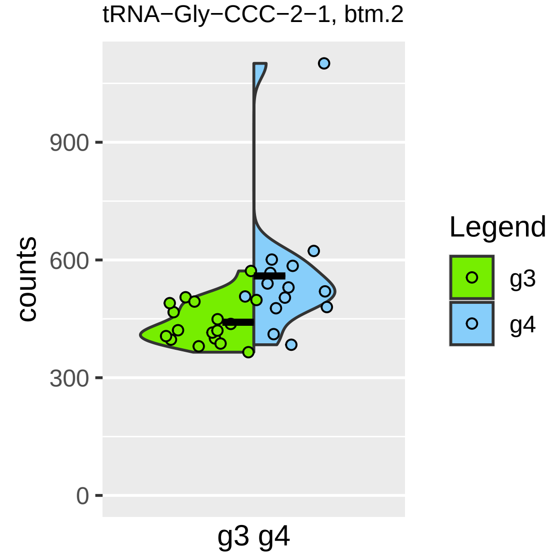
tRNA−Gly−CCC−2−1, btm.2
900
600
300
0
counts
g3 g4
Legend
g3
g4

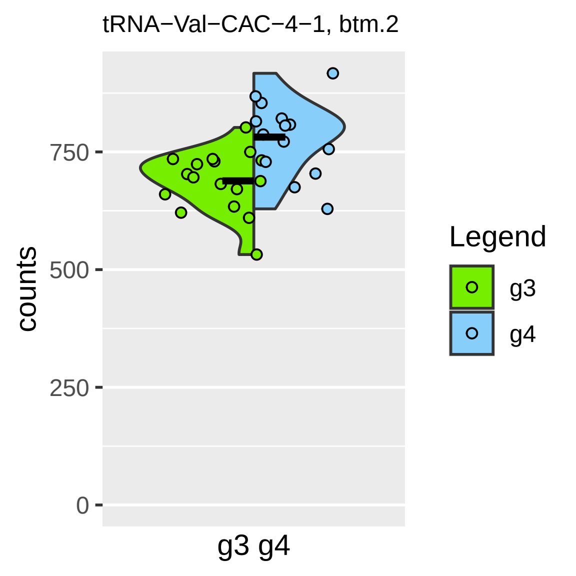

tRNA−Val−CAC−4−1, btm.2
counts
750
500
250
0
g3 g4
Legend
g3
g4

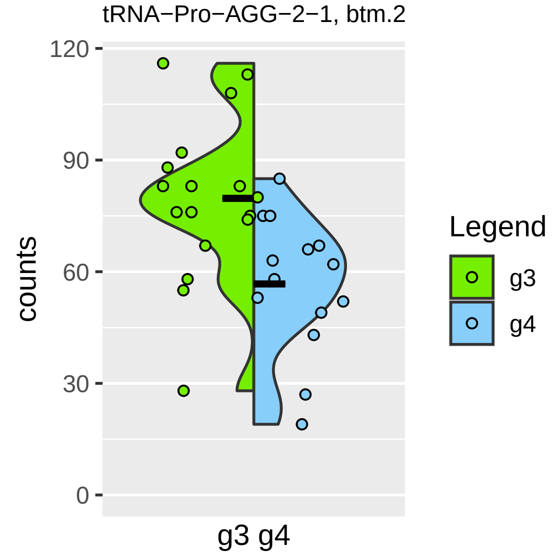
tRNA−Pro−AGG−2−1, btm.2
120
90
60
30
0
counts
g3 g4
Legend
g3
g4

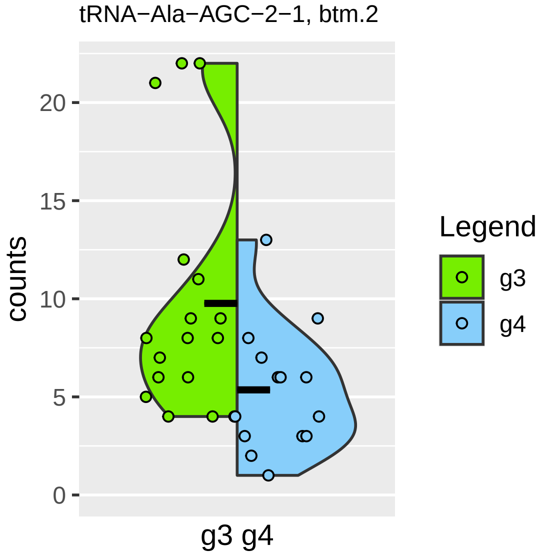

tRNA−Ala−AGC−2−1, btm.2
counts
0
5
10
15
20
g3 g4
Legend
g3
g4

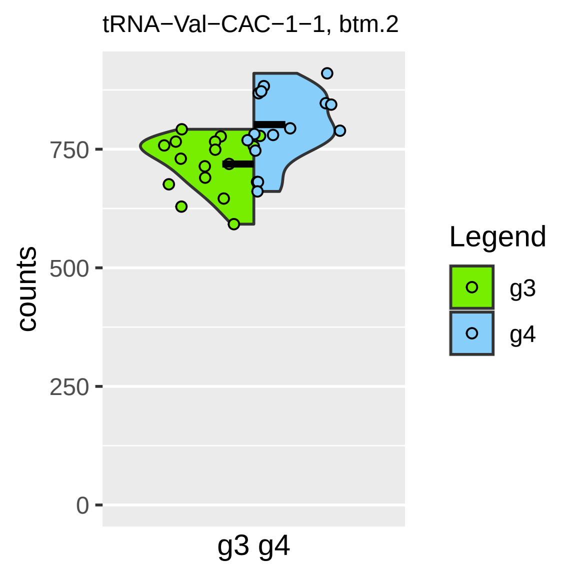
tRNA−Val−CAC−1−1, btm.2
counts
750
500
250
0
g3 g4
Legend
g3
g4

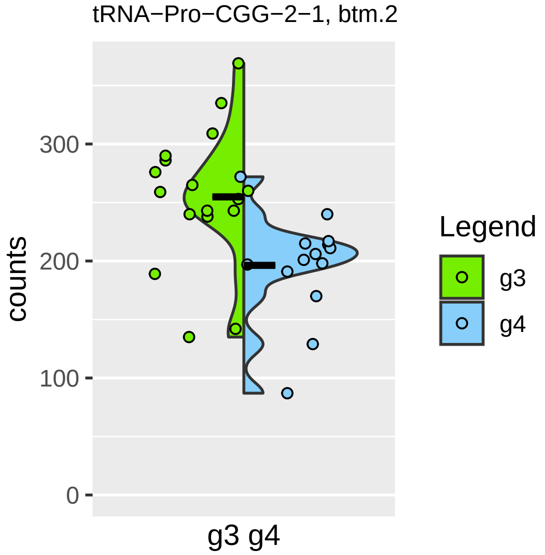
tRNA−Pro−CGG−2−1, btm.2
counts
300
200
100
0
g3 g4
Legend
g3
g4

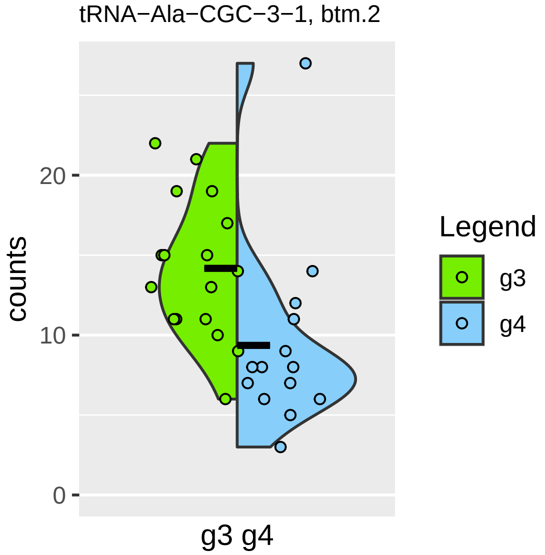
tRNA−Ala−CGC−3−1, btm.2
counts
20
10
0
g3 g4
Legend
g3
g4

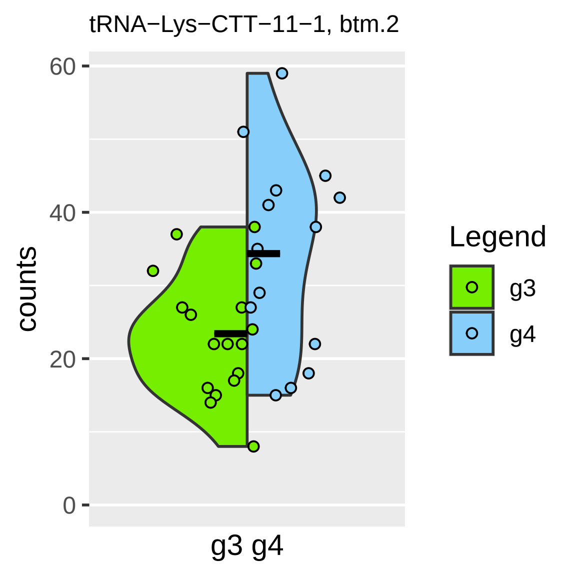

tRNA−Lys−CTT−11−1, btm.2
counts
60
40
20
0
g3 g4
Legend
g3
g4

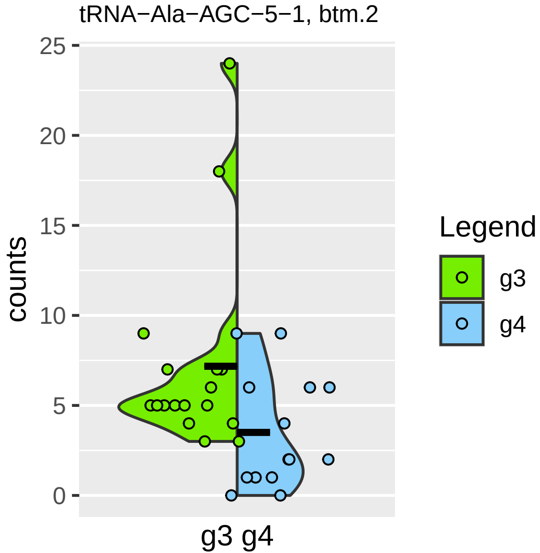

tRNA−Ala−AGC−5−1, btm.2
counts
25
20
15
10
5
0
g3 g4
Legend
g3
g4

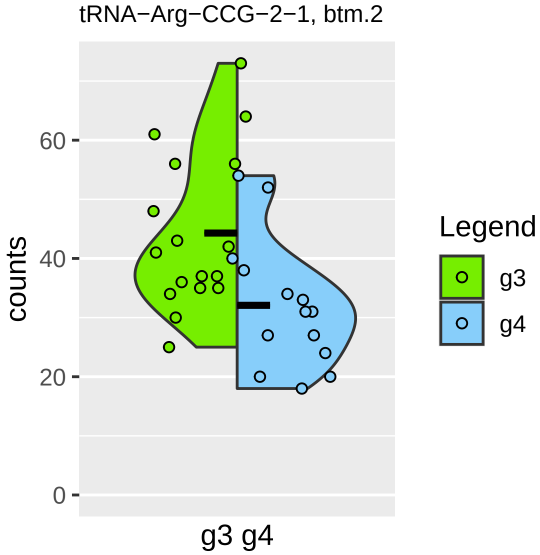

tRNA−Arg−CCG−2−1, btm.2
counts
0
20
40
60
g3 g4
Legend
g3
g4

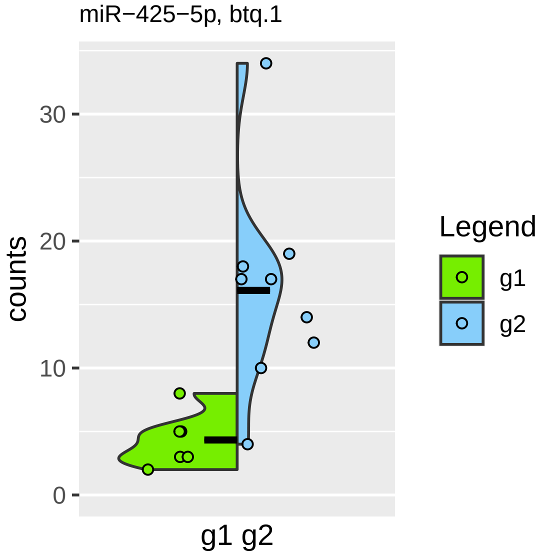

miR−425−5p, btq.1
counts
30
20
10
0
g1 g2
Legend
g1
g2

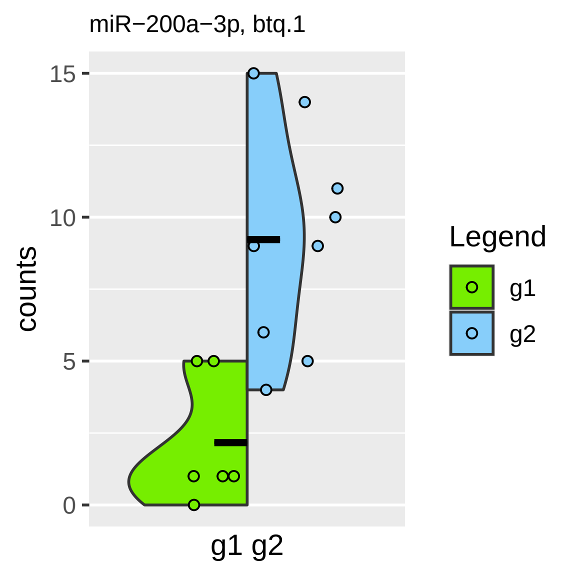

miR−200a−3p, btq.1
counts
15
10
5
0
g1 g2
Legend
g1
g2

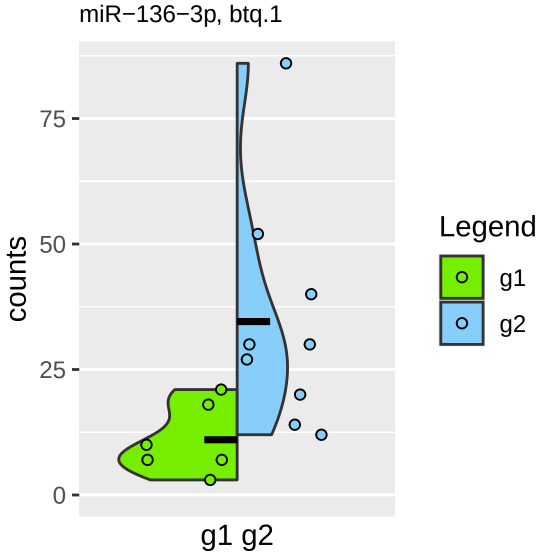
miR−136−3p, btq.1
counts
75
50
25
0
g1 g2
Legend
g1
g2

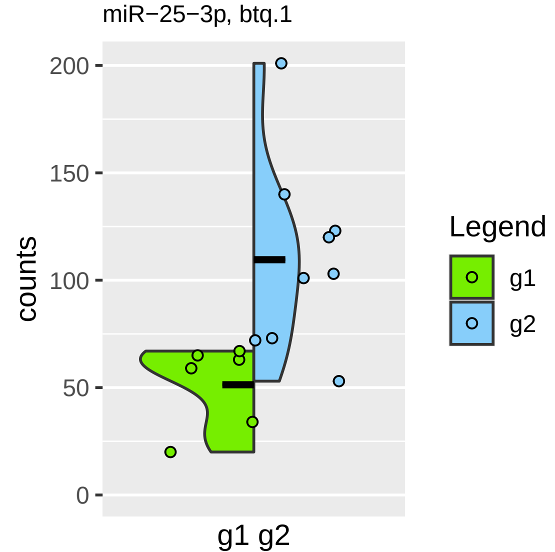

miR−25−3p, btq.1
counts
200
150
100
50
0
g1 g2
Legend
g1
g2

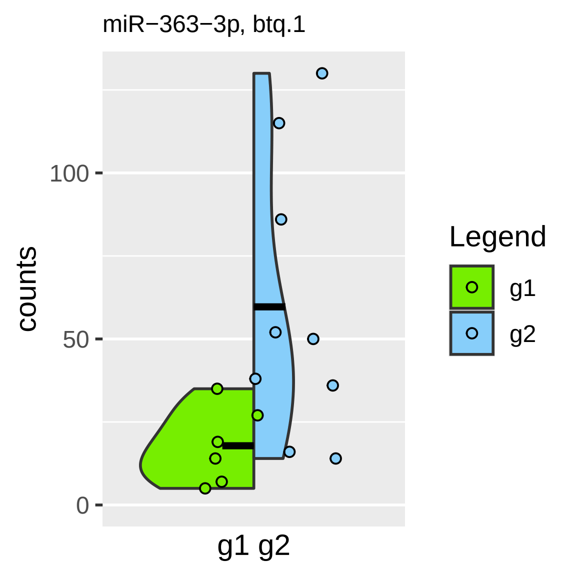

miR−363−3p, btq.1
counts
100
50
0
g1 g2
Legend
g1
g2

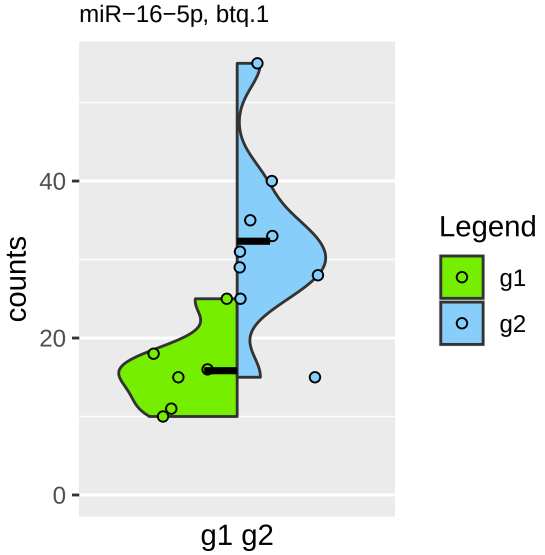

miR−16−5p, btq.1
counts
40
20
0
g1 g2
Legend
g1
g2

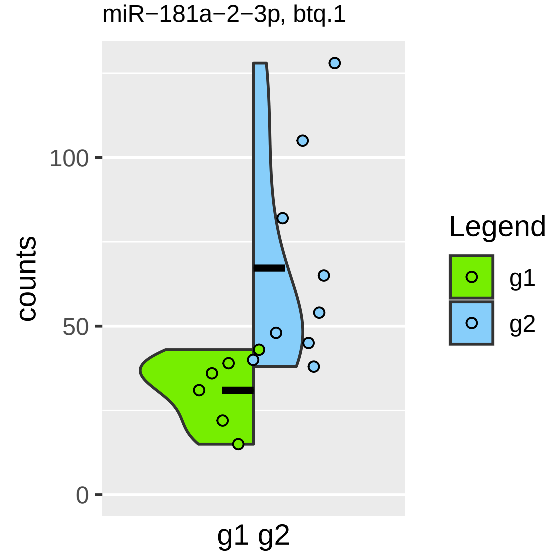

miR−181a−2−3p, btq.1
counts
100
50
0
g1 g2
Legend
g1
g2

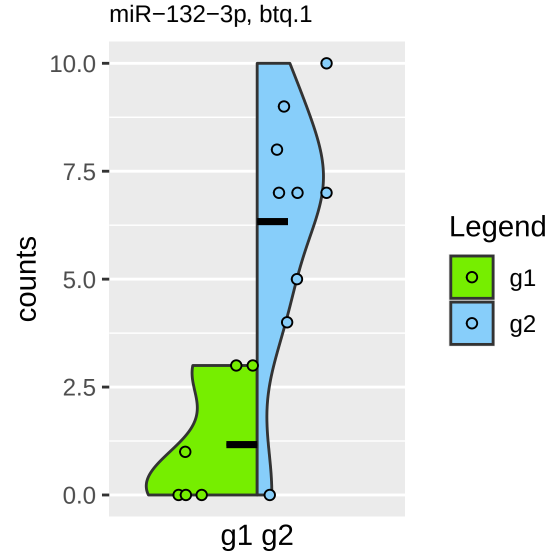
miR−132−3p, btq.1
10.0
7.5
5.0
2.5
0.0
counts
g1 g2
Legend
g1
g2

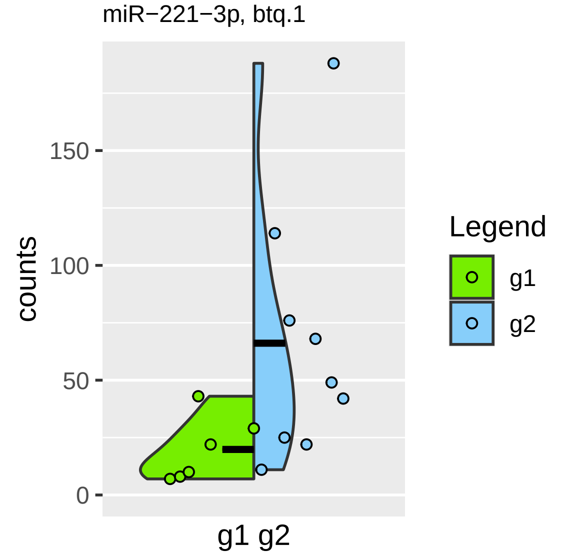

miR−221−3p, btq.1
counts
150
100
50
0
g1 g2
Legend
g1
g2

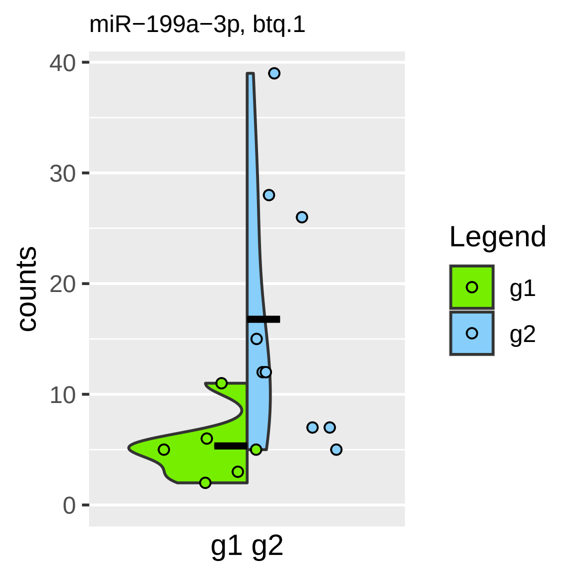

miR−199a−3p, btq.1
40
30
20
10
0
counts
g1 g2
Legend
g1
g2

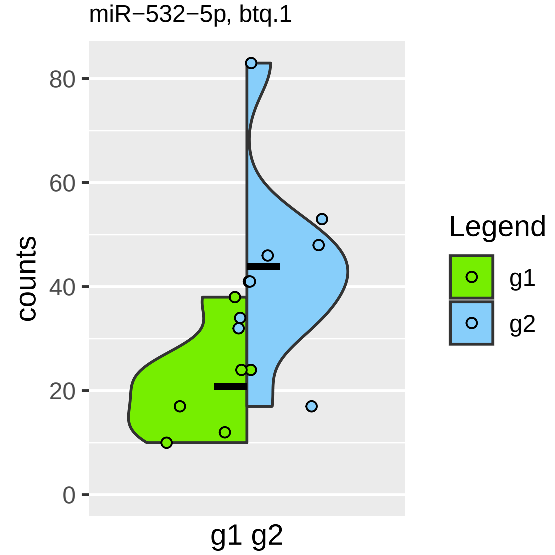

miR−532−5p, btq.1
80
60
40
20
0
counts
g1 g2
Legend
g1
g2

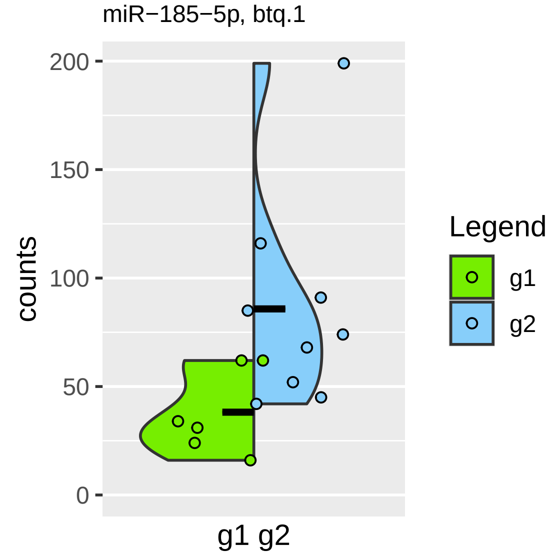

miR−185−5p, btq.1
200
150
100
50
0
counts
g1 g2
Legend
g1
g2

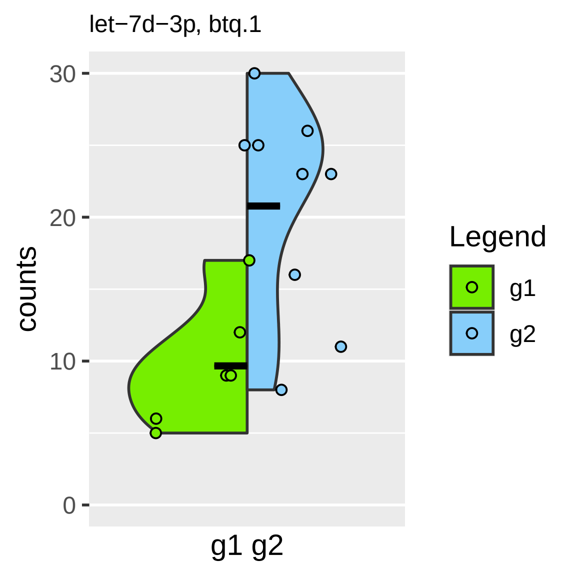

let−7d−3p, btq.1
30
20
10
0
counts
g1 g2
Legend
g1
g2

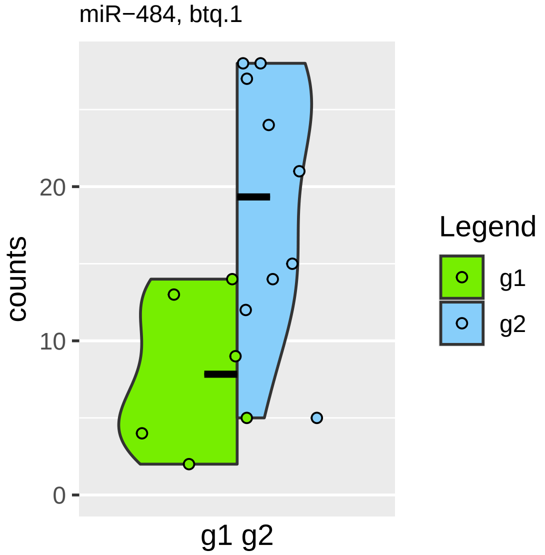

miR−484, btq.1
counts
20
10
0
g1 g2
Legend
g1
g2

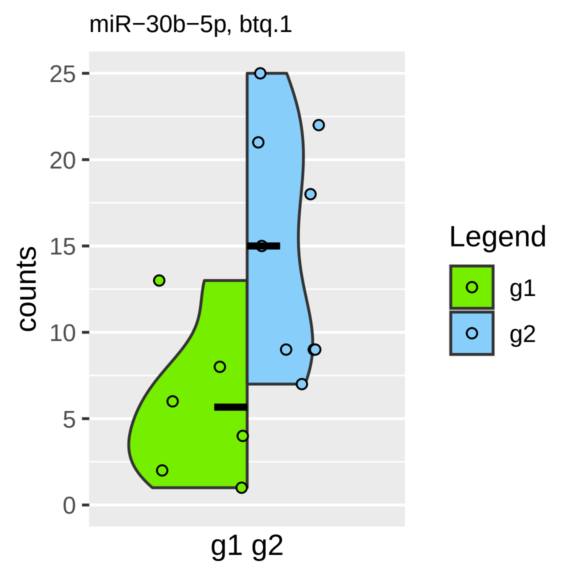
miR−30b−5p, btq.1
counts
25
20
15
10
5
0
g1 g2
Legend
g1
g2

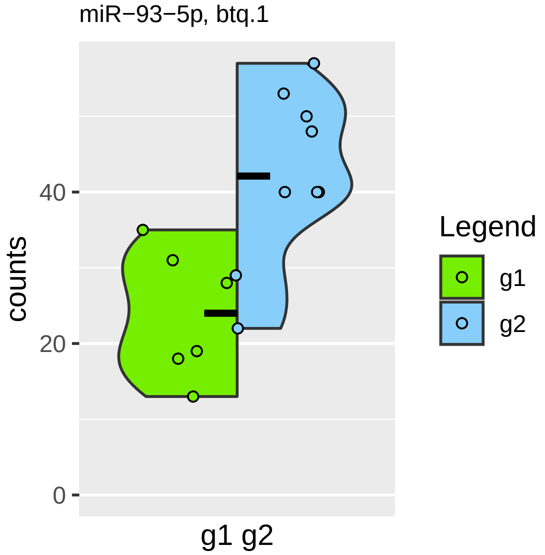

miR−93−5p, btq.1
counts
0
20
40
g1 g2
Legend
g1
g2

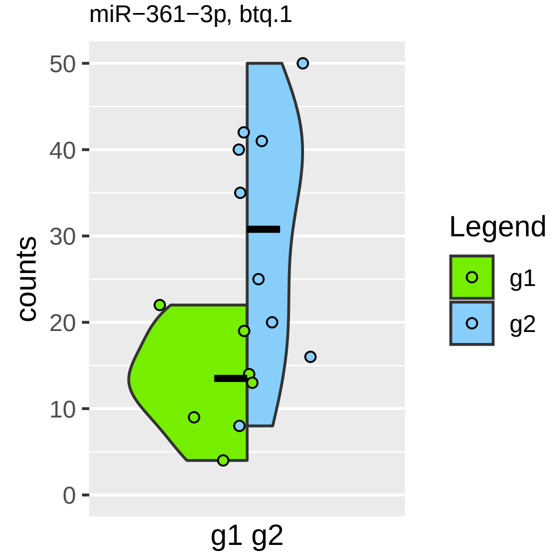

miR−361−3p, btq.1
counts
50
40
30
20
10
0
g1 g2
Legend
g1
g2

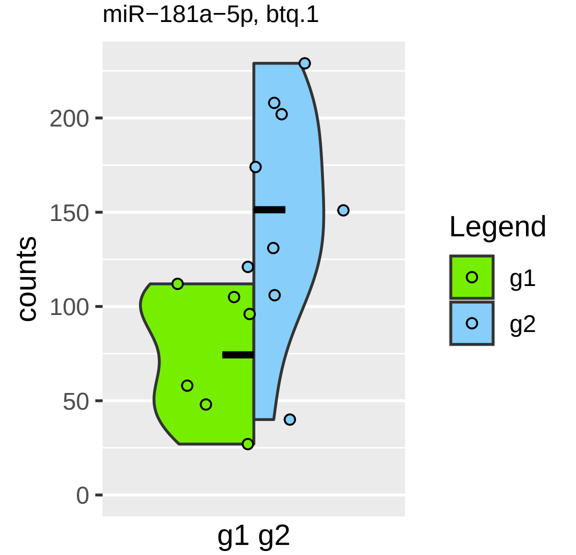
miR−181a−5p, btq.1
counts
200
150
100
50
0
g1 g2
Legend
g1
g2

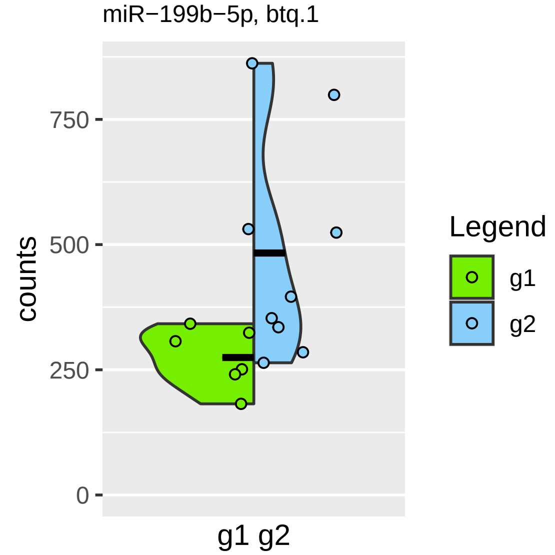

miR−199b−5p, btq.1
counts
750
500
250
0
g1 g2
Legend
g1
g2

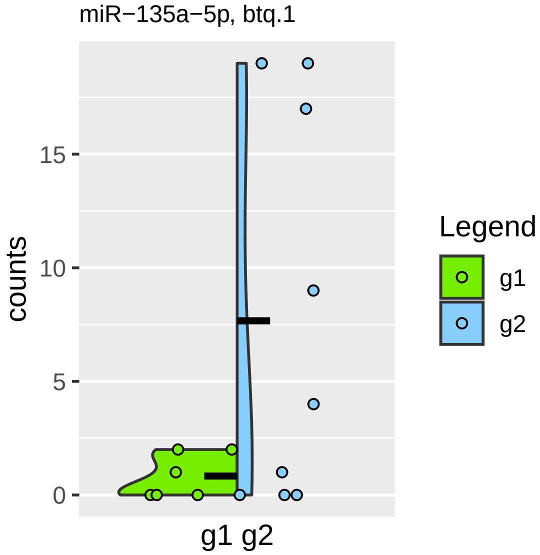

miR−135a−5p, btq.1
counts
0
5
10
15
g1 g2
Legend
g1
g2

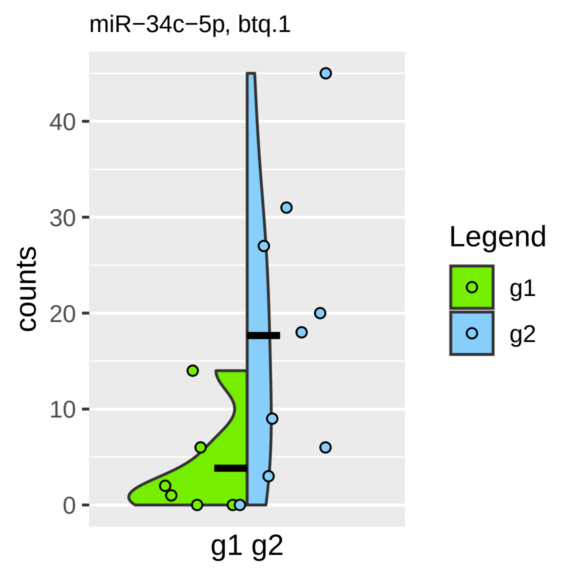
miR−34c−5p, btq.1
counts
40
30
20
10
0
g1 g2
Legend
g1
g2

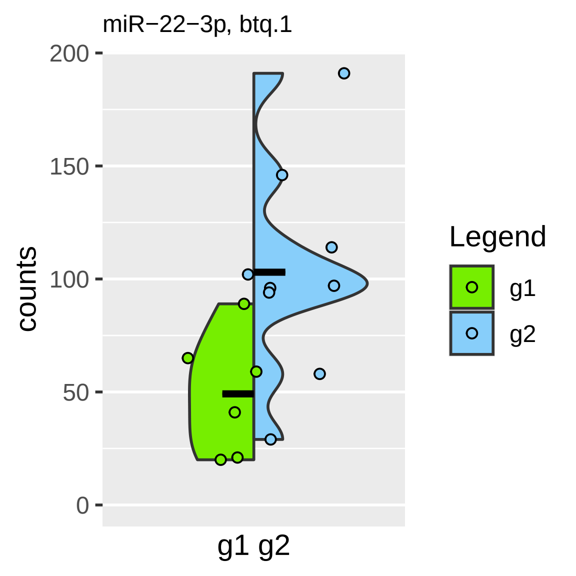

miR−22−3p, btq.1
200
150
100
50
0
counts
g1 g2
Legend
g1
g2

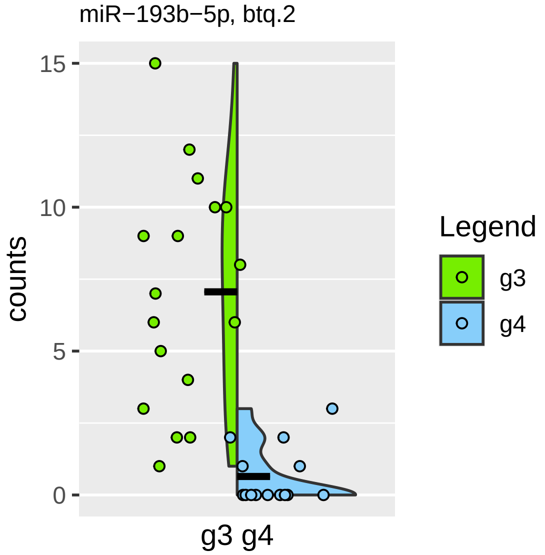
miR−193b−5p, btq.2
15
10
5
0
counts
g3 g4
Legend
g3
g4

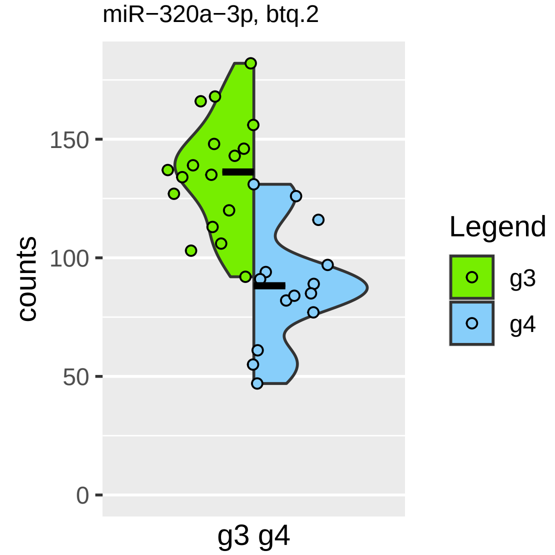

miR−320a−3p, btq.2
counts
150
100
50
0
g3 g4
Legend
g3
g4

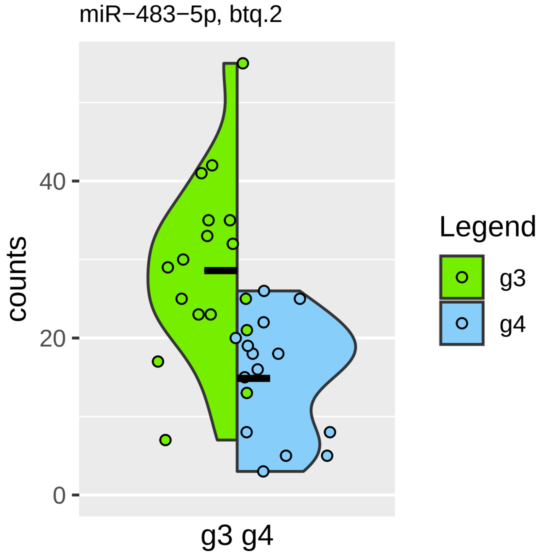

miR−483−5p, btq.2
counts
40
20
0
g3 g4
Legend
g3
g4

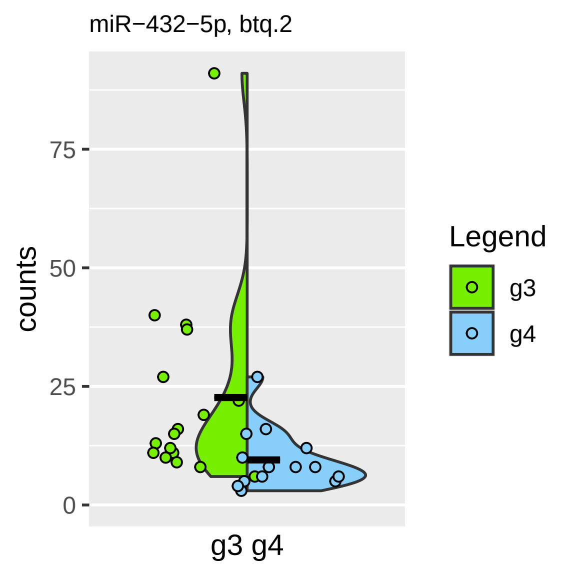
miR−432−5p, btq.2
counts
75
50
25
0
g3 g4
Legend
g3
g4

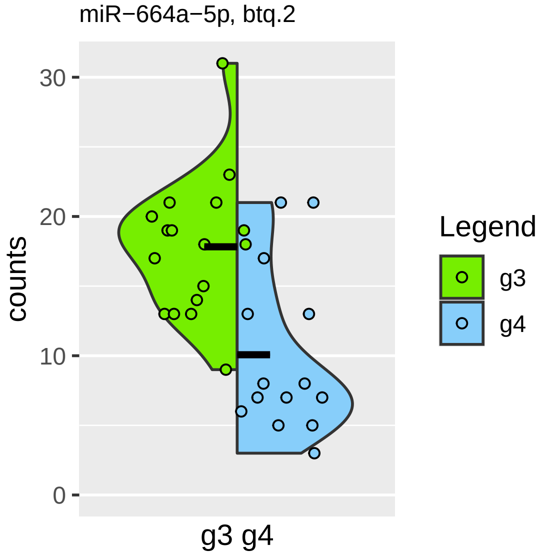

miR−664a−5p, btq.2
30
20
10
0
counts
g3 g4
Legend
g3
g4

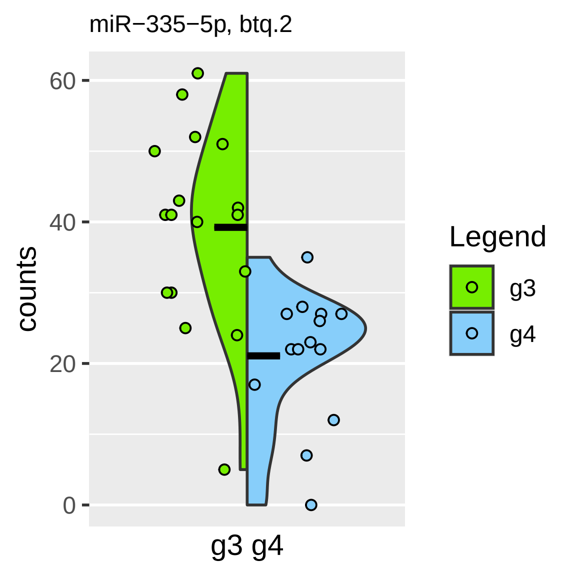

miR−335−5p, btq.2
60
40
20
0
counts
g3 g4
Legend
g3
g4

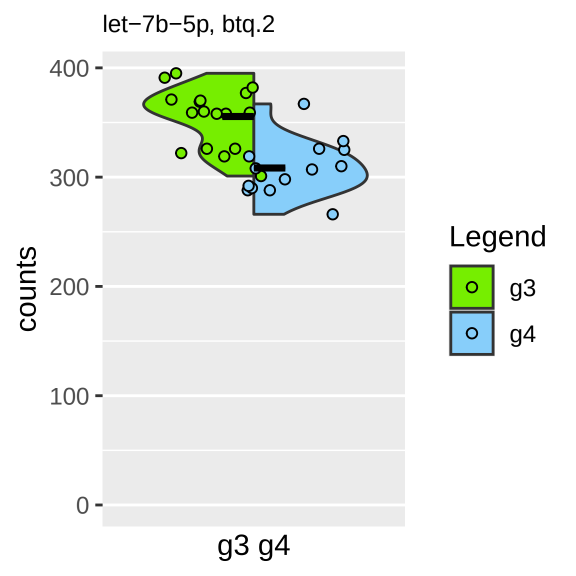

let−7b−5p, btq.2
400
300
200
100
0
counts
g3 g4
Legend
g3
g4

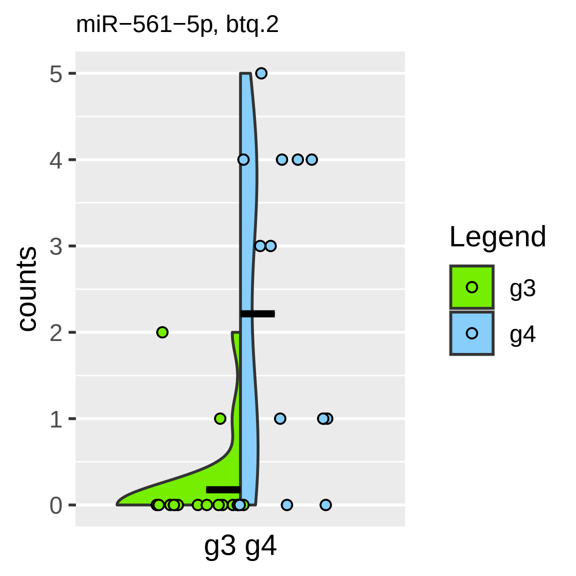
miR−561−5p, btq.2
counts
5
4
3
2
1
0
g3 g4
Legend
g3
g4

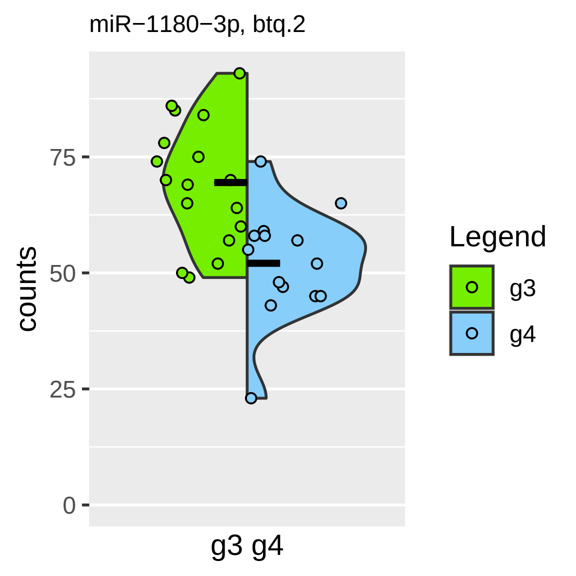

miR−1180−3p, btq.2
counts
75
50
25
0
g3 g4
Legend
g3
g4

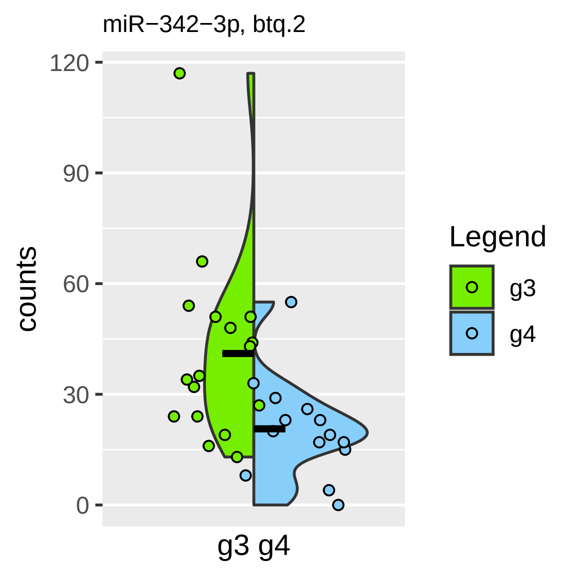

miR−342−3p, btq.2
counts
120
90
60
30
0
g3 g4
Legend
g3
g4

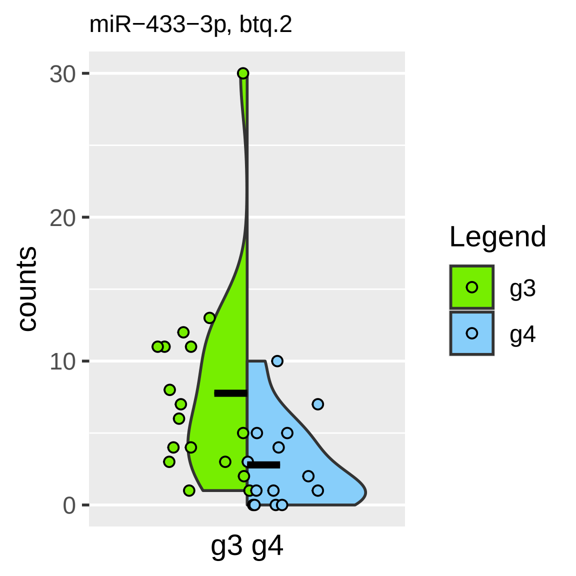

miR−433−3p, btq.2
counts
30
20
10
0
g3 g4
Legend
g3
g4

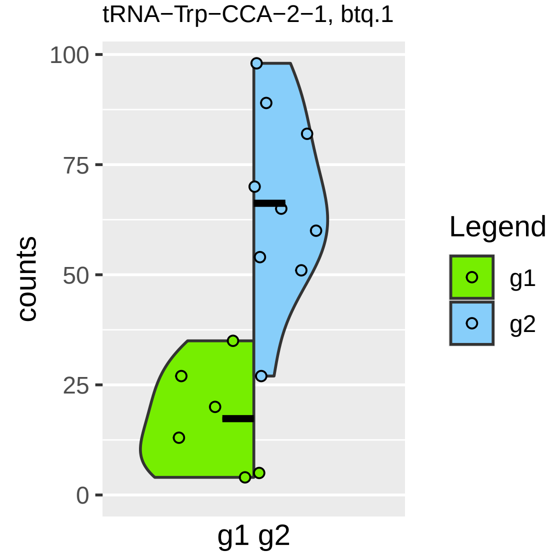
tRNA−Trp−CCA−2−1, btq.1
100
75
50
25
0
counts
g1 g2
Legend
g1
g2

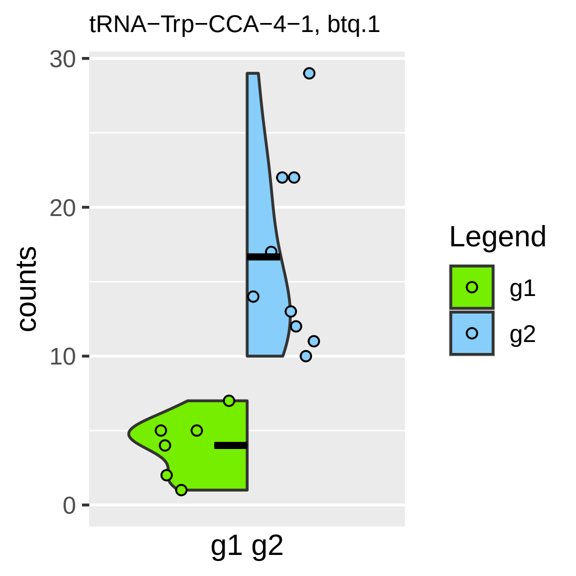

tRNA−Trp−CCA−4−1, btq.1
30
20
10
0
counts
g1 g2
Legend
g1
g2

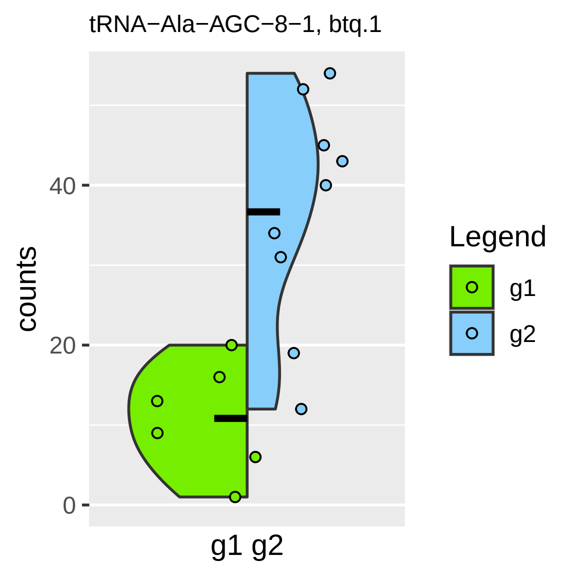

tRNA−Ala−AGC−8−1, btq.1
counts
0
20
40
g1 g2
Legend
g1
g2

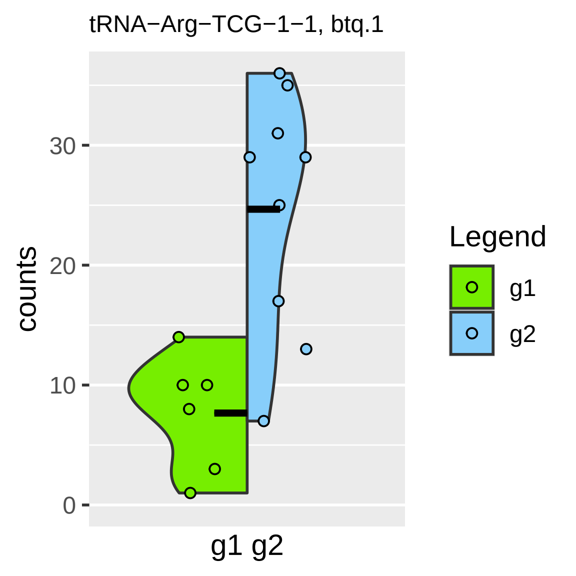

tRNA−Arg−TCG−1−1, btq.1
counts
0
10
20
30
g1 g2
Legend
g1
g2

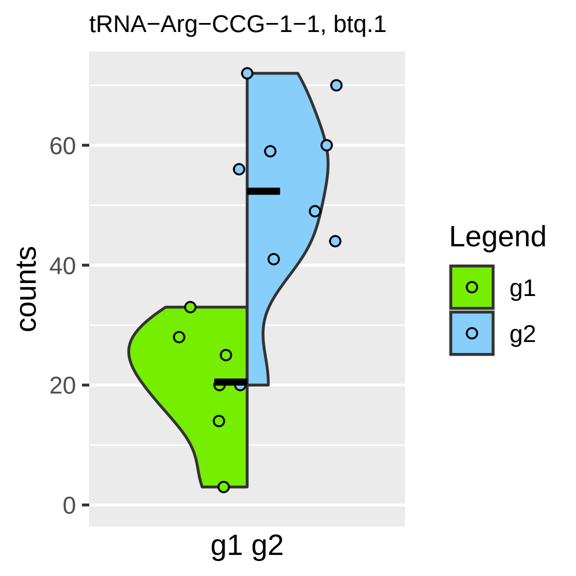

tRNA−Arg−CCG−1−1, btq.1
counts
60
40
20
0
g1 g2
Legend
g1
g2

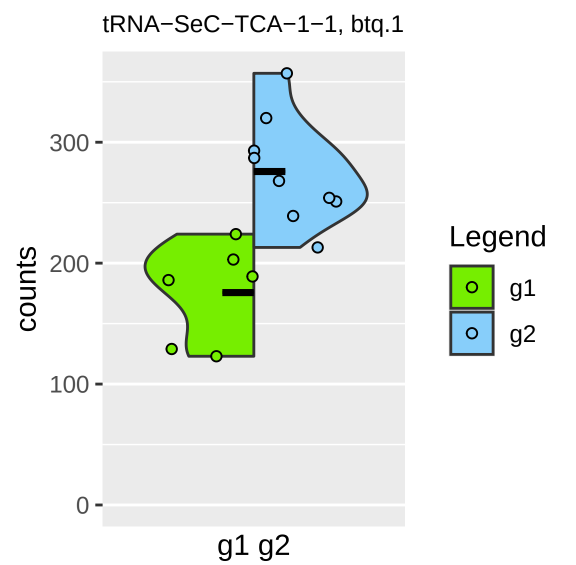

tRNA−SeC−TCA−1−1, btq.1
counts
300
200
100
0
g1 g2
Legend
g1
g2

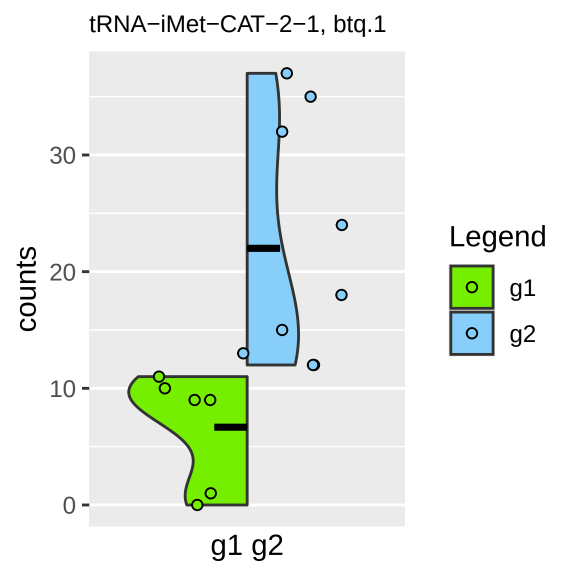
tRNA−iMet−CAT−2−1, btq.1
30
20
10
0
counts
g1 g2
Legend
g1
g2

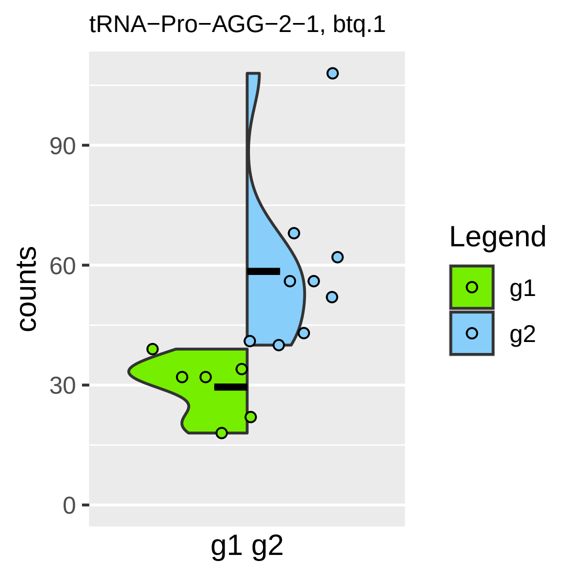
tRNA−Pro−AGG−2−1, btq.1
counts
90
60
30
0
g1 g2
Legend
g1
g2

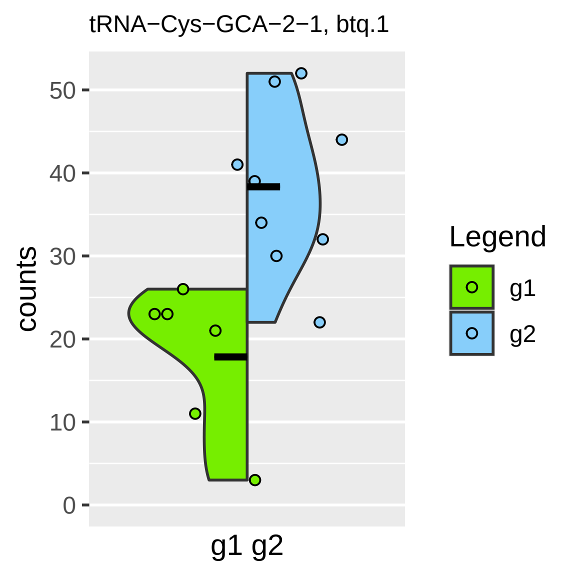
tRNA−Cys−GCA−2−1, btq.1
counts
0
10
20
30
40
50
g1 g2
Legend
g1
g2

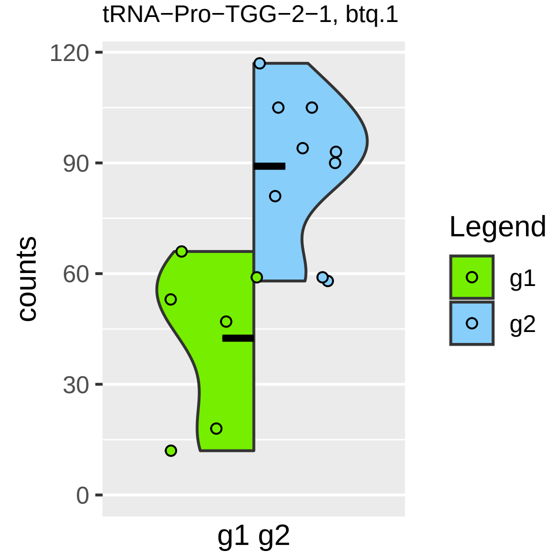

tRNA−Pro−TGG−2−1, btq.1
counts
120
90
60
30
0
g1 g2
Legend
g1
g2

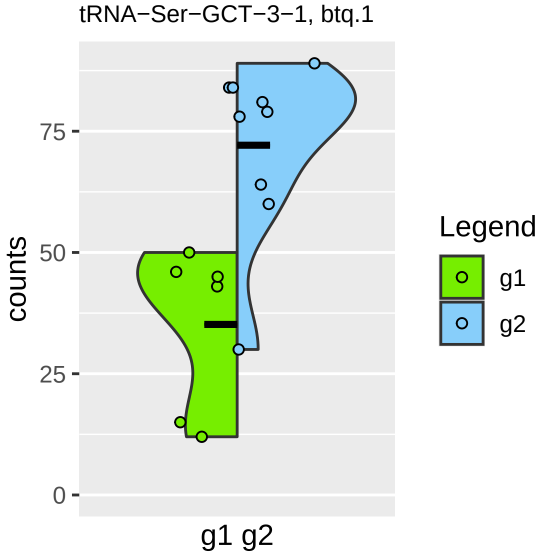

tRNA−Ser−GCT−3−1, btq.1
counts
75
50
25
0
g1 g2
Legend
g1
g2

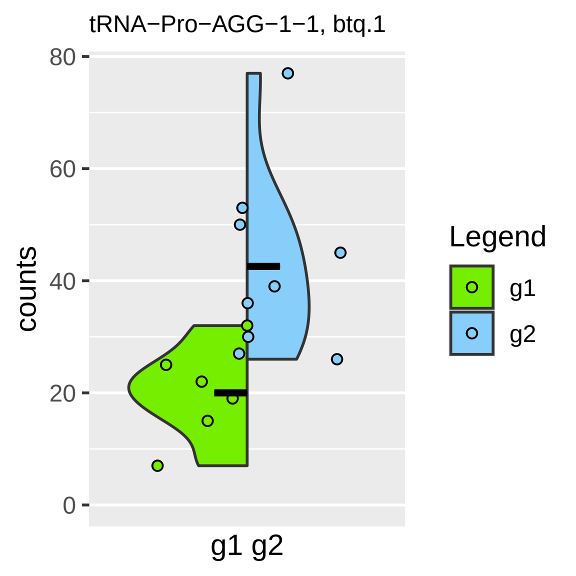

tRNA−Pro−AGG−1−1, btq.1
counts
80
60
40
20
0
g1 g2
Legend
g1
g2

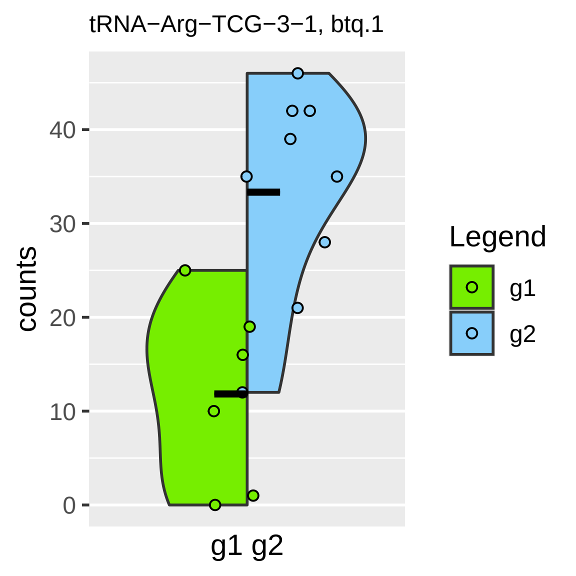
tRNA−Arg−TCG−3−1, btq.1
counts
40
30
20
10
0
g1 g2
Legend
g1
g2

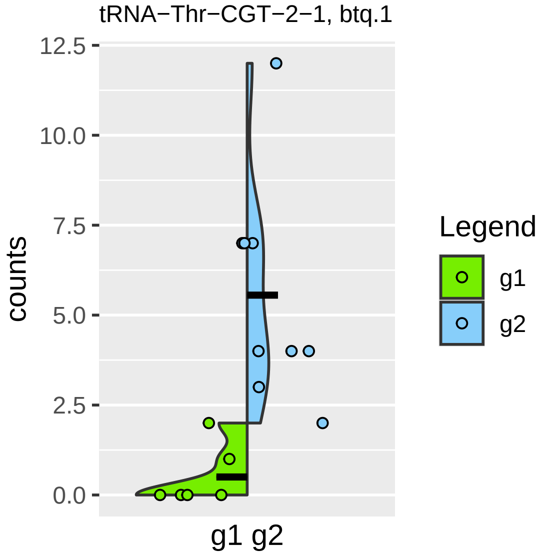
tRNA−Thr−CGT−2−1, btq.1
12.5
10.0
7.5
5.0
2.5
0.0
counts
g1 g2
Legend
g1
g2

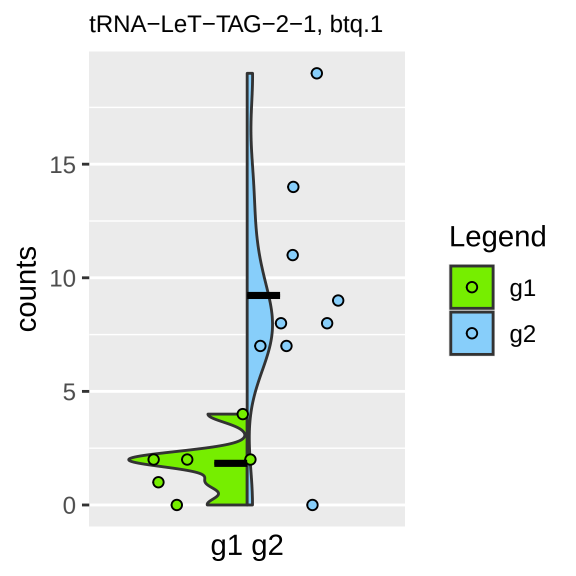
tRNA−LeT−TAG−2−1, btq.1
counts
15
10
5
0
g1 g2
Legend
g1
g2

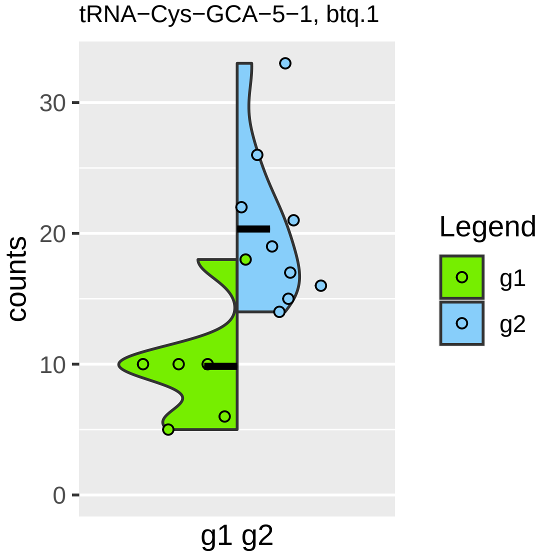
tRNA−Cys−GCA−5−1, btq.1
counts
30
20
10
0
g1 g2
Legend
g1
g2

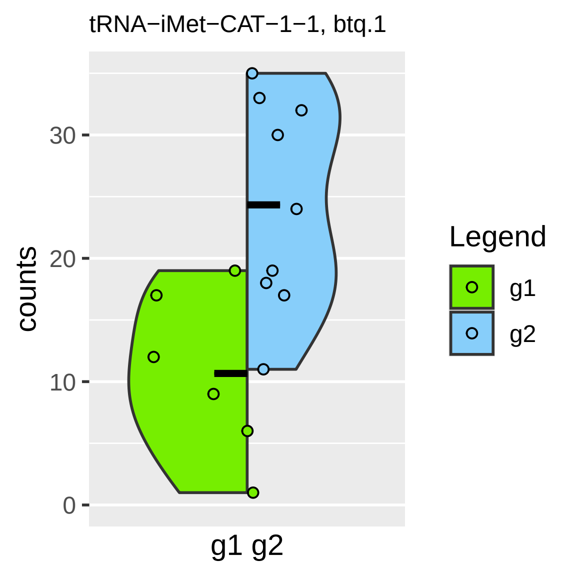

tRNA−iMet−CAT−1−1, btq.1
counts
0
10
20
30
g1 g2
Legend
g1
g2

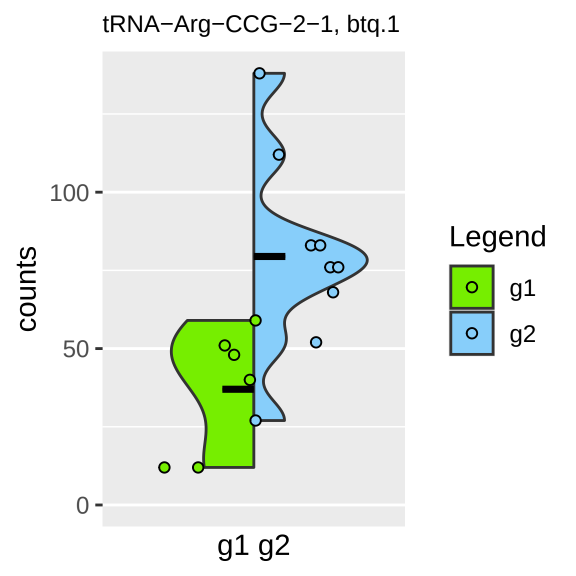
tRNA−Arg−CCG−2−1, btq.1
counts
100
50
0
g1 g2
Legend
g1
g2

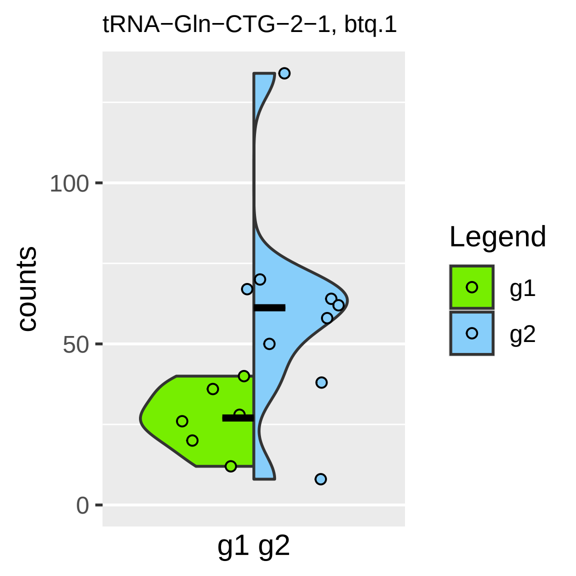
tRNA−Gln−CTG−2−1, btq.1
counts
100
50
0
g1 g2
Legend
g1
g2

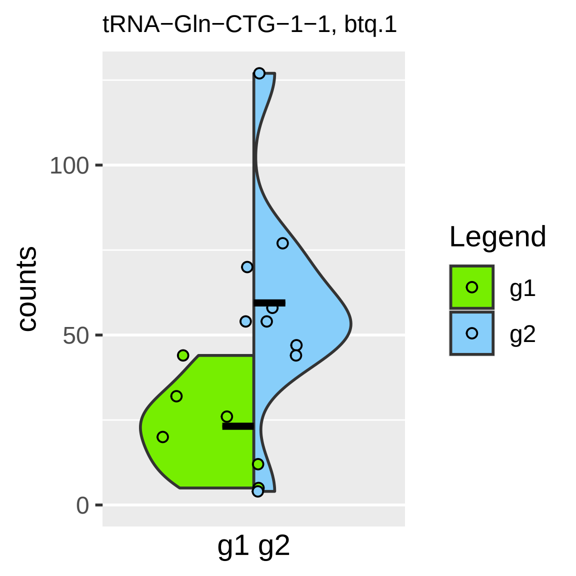

tRNA−Gln−CTG−1−1, btq.1
counts
100
50
0
g1 g2
Legend
g1
g2

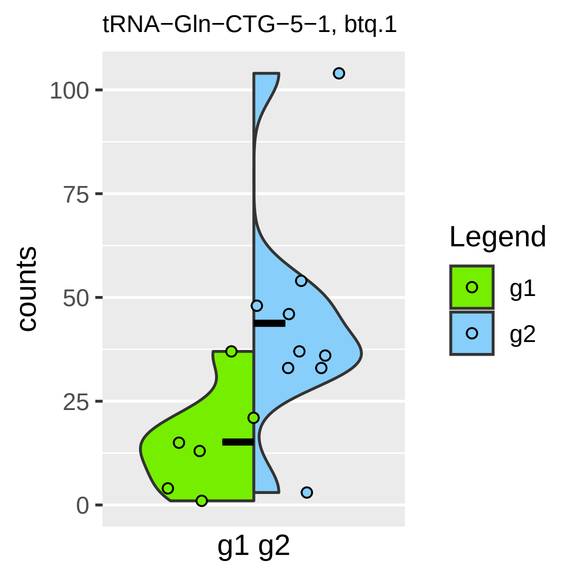
tRNA−Gln−CTG−5−1, btq.1
counts
100
75
50
25
0
g1 g2
Legend
g1
g2

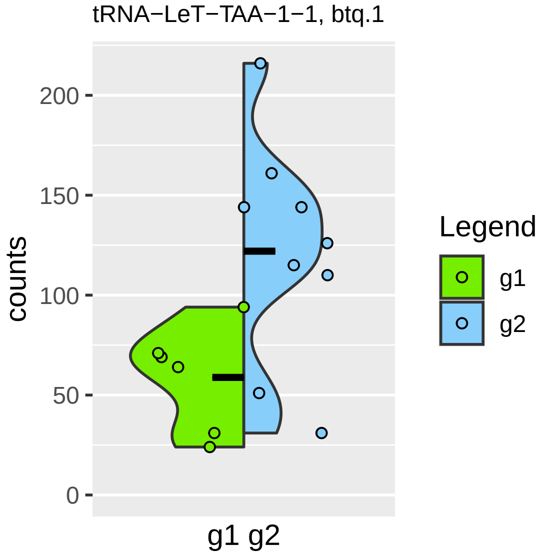

tRNA−LeT−TAA−1−1, btq.1
counts
200
150
100
50
0
g1 g2
Legend
g1
g2

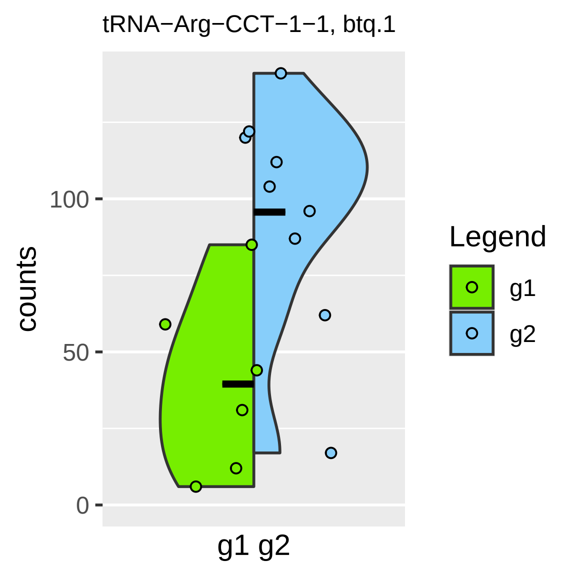
tRNA−Arg−CCT−1−1, btq.1
counts
100
50
0
g1 g2
Legend
g1
g2

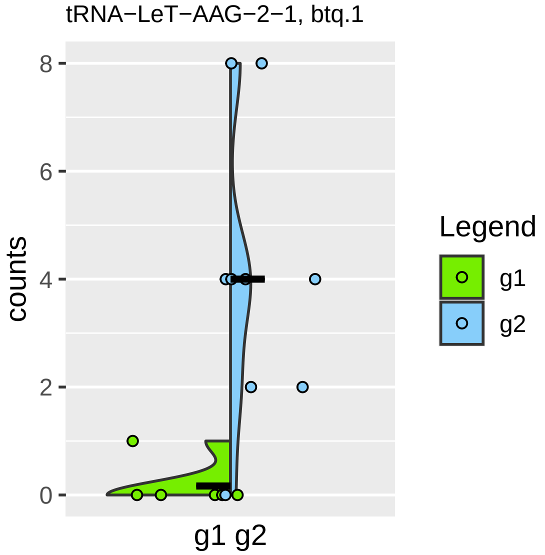

tRNA−LeT−AAG−2−1, btq.1
counts
8
6
4
2
0
g1 g2
Legend
g1
g2

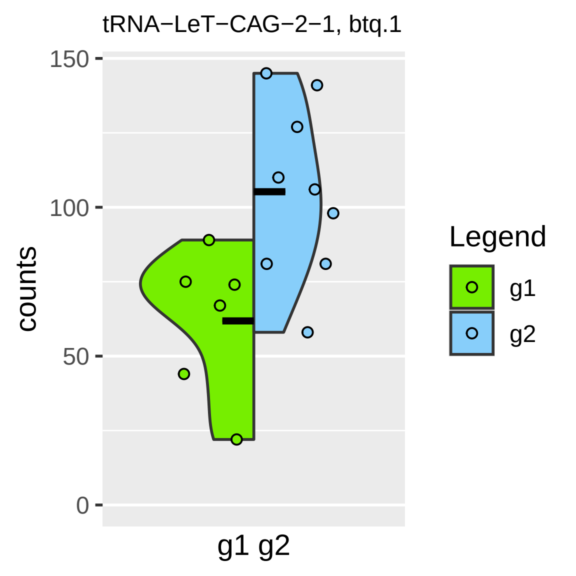
tRNA−LeT−CAG−2−1, btq.1
150
100
50
0
counts
g1 g2
Legend
g1
g2

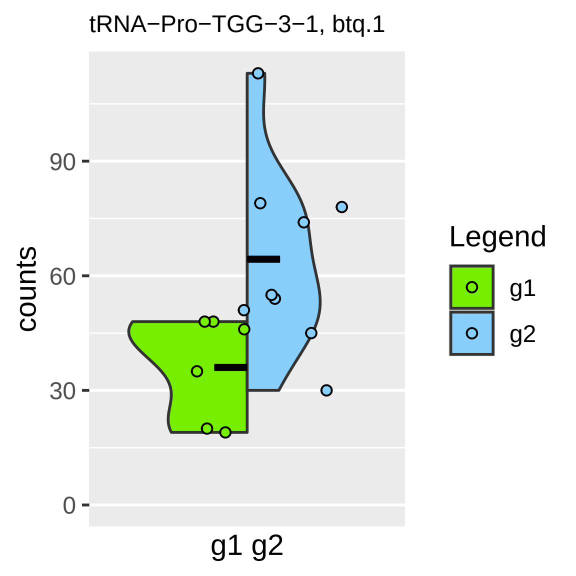

tRNA−Pro−TGG−3−1, btq.1
counts
90
60
30
0
g1 g2
Legend
g1
g2

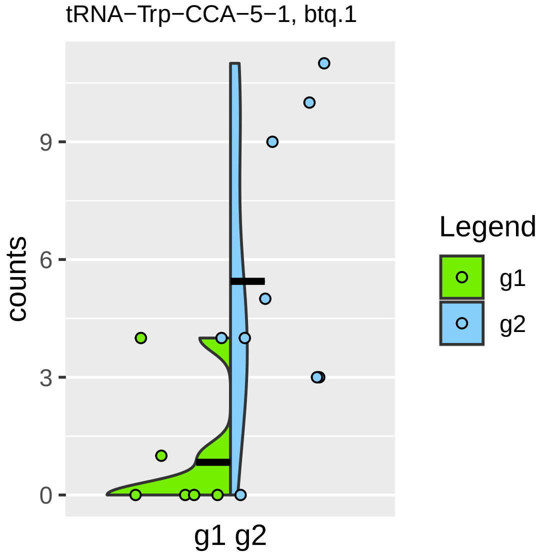

tRNA−Trp−CCA−5−1, btq.1
counts
0
3
6
9
g1 g2
Legend
g1
g2

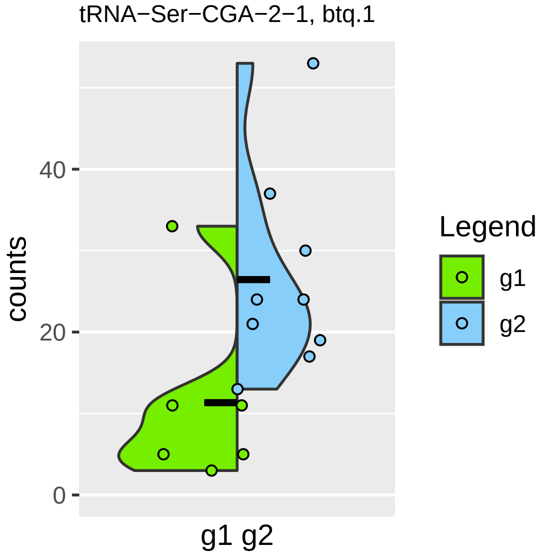

tRNA−Ser−CGA−2−1, btq.1
counts
40
20
0
g1 g2
Legend
g1
g2

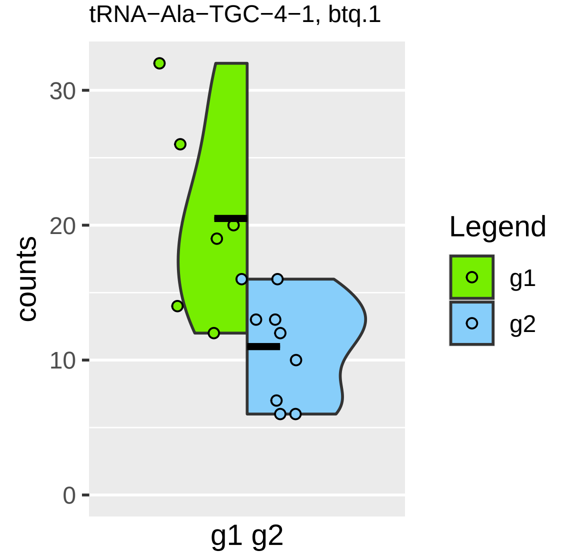
tRNA−Ala−TGC−4−1, btq.1
counts
30
20
10
0
g1 g2
Legend
g1
g2

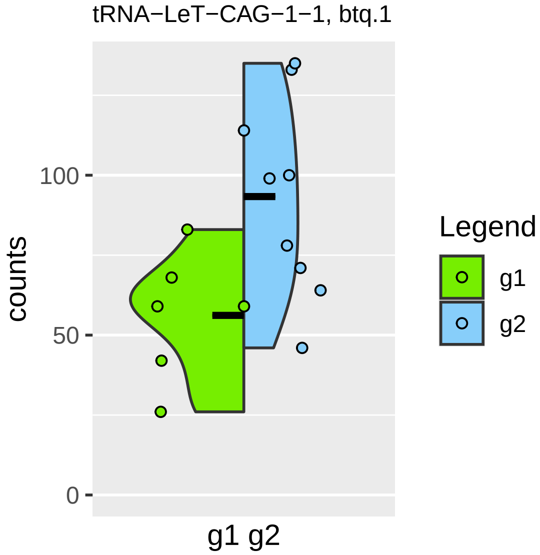

tRNA−LeT−CAG−1−1, btq.1
counts
100
50
0
g1 g2
Legend
g1
g2

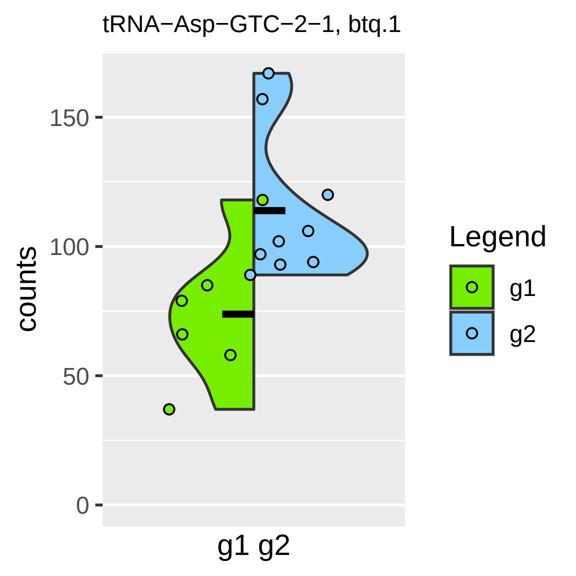
tRNA−Asp−GTC−2−1, btq.1
counts
150
100
50
0
g1 g2
Legend
g1
g2

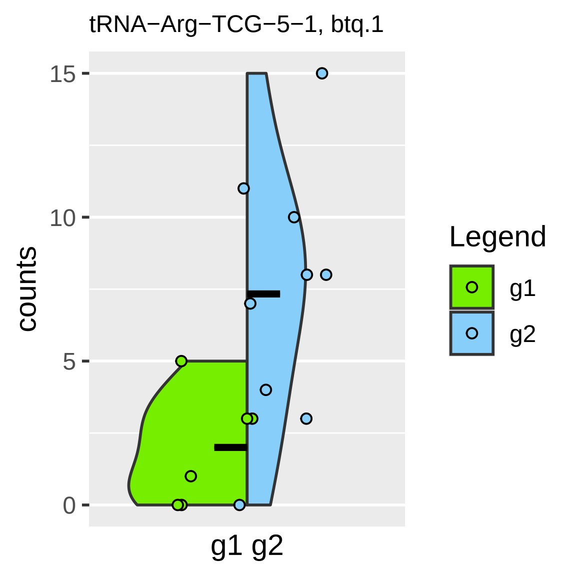

tRNA−Arg−TCG−5−1, btq.1
15
10
5
0
counts
g1 g2
Legend
g1
g2

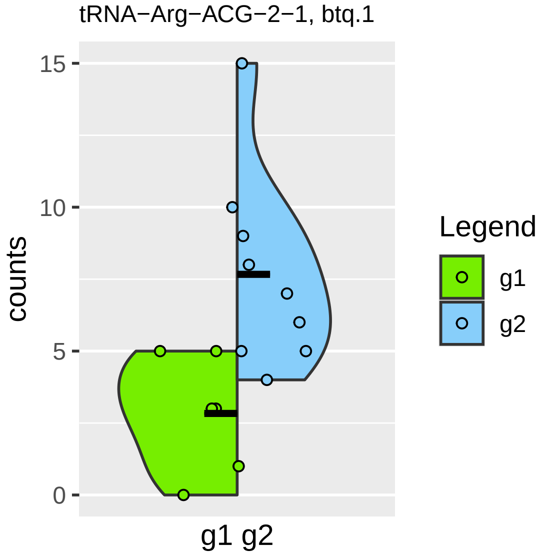

tRNA−Arg−ACG−2−1, btq.1
15
10
5
0
counts
g1 g2
Legend
g1
g2

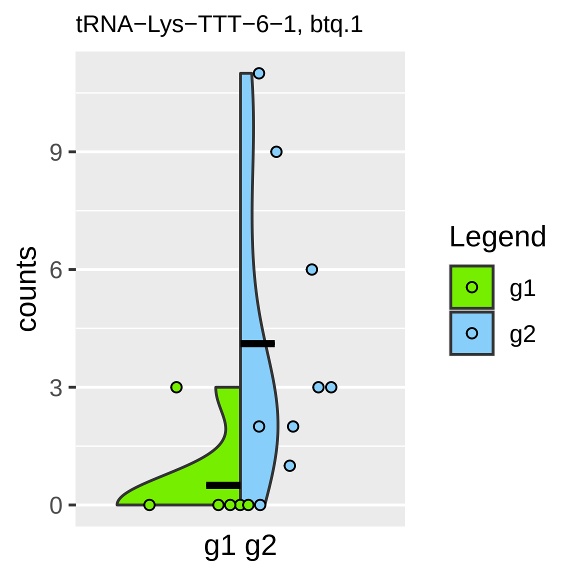

tRNA−Lys−TTT−6−1, btq.1
counts
0
3
6
9
g1 g2
Legend
g1
g2

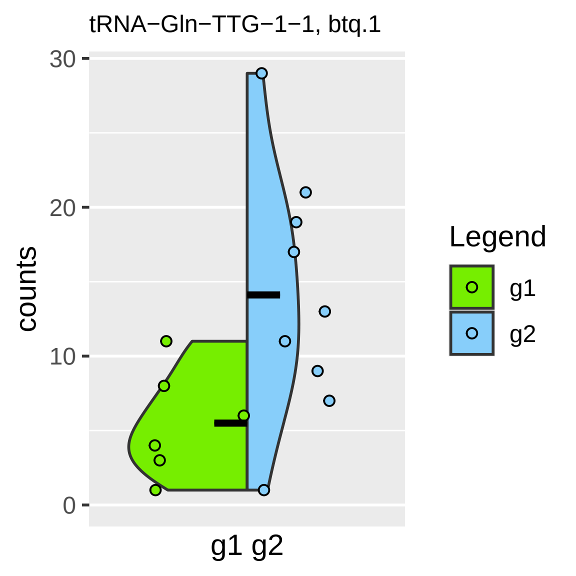

tRNA−Gln−TTG−1−1, btq.1
counts
30
20
10
0
g1 g2
Legend
g1
g2

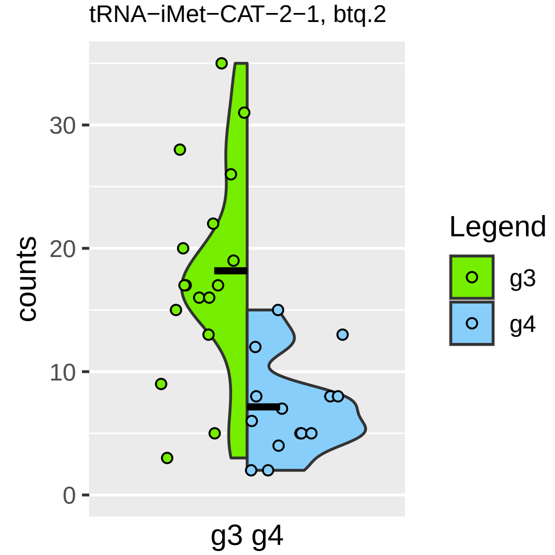
tRNA−iMet−CAT−2−1, btq.2
counts
30
20
10
0
g3 g4
Legend
g3
g4

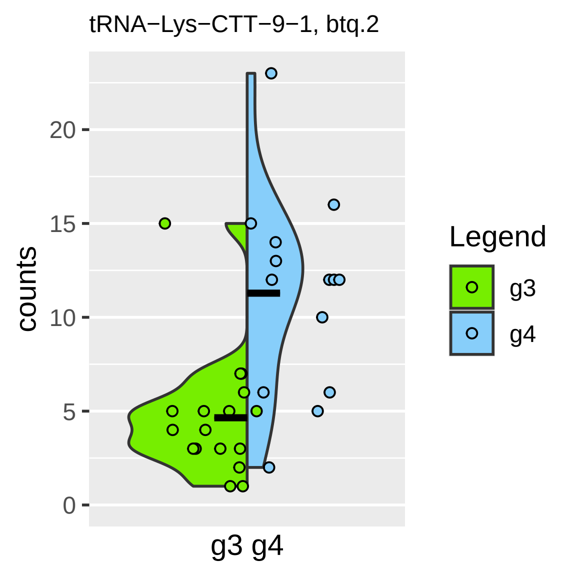
tRNA−Lys−CTT−9−1, btq.2
counts
0
5
10
15
20
g3 g4
Legend
g3
g4

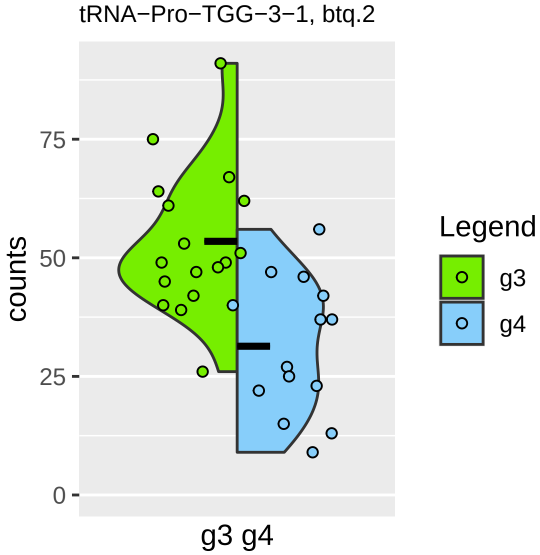
tRNA−Pro−TGG−3−1, btq.2
counts
75
50
25
0
g3 g4
Legend
g3
g4

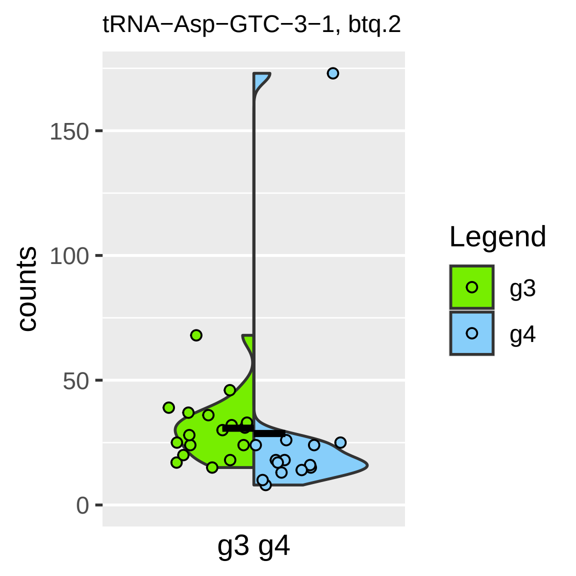

tRNA−Asp−GTC−3−1, btq.2
counts
150
100
50
0
g3 g4
Legend
g3
g4

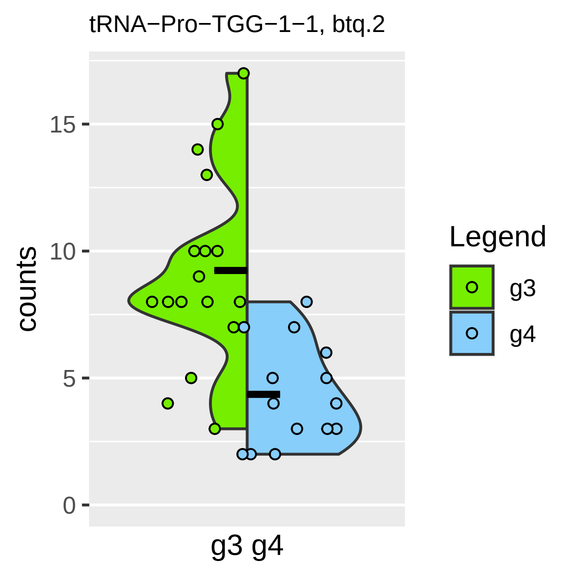
tRNA−Pro−TGG−1−1, btq.2
counts
15
10
5
0
g3 g4
Legend
g3
g4

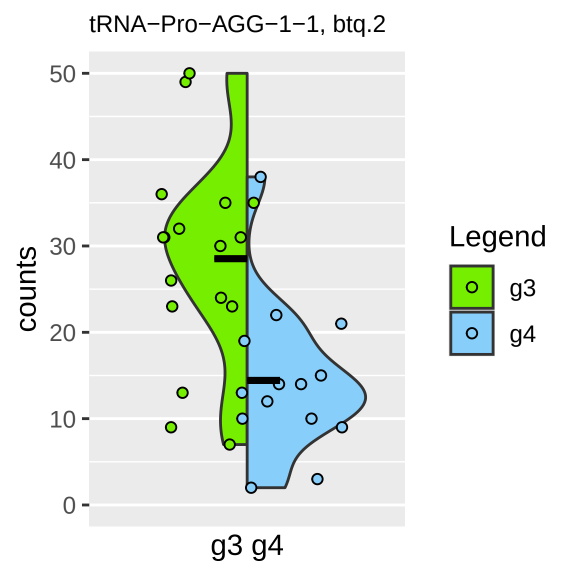

tRNA−Pro−AGG−1−1, btq.2
counts
50
40
30
20
10
0
g3 g4
Legend
g3
g4

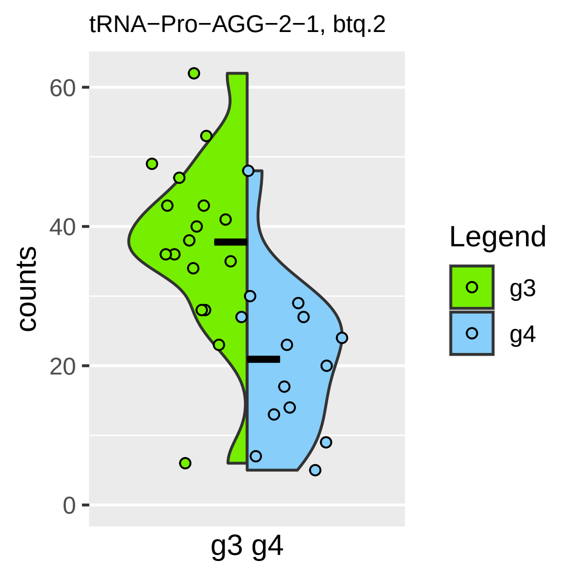

tRNA−Pro−AGG−2−1, btq.2
counts
60
40
20
0
g3 g4
Legend
g3
g4

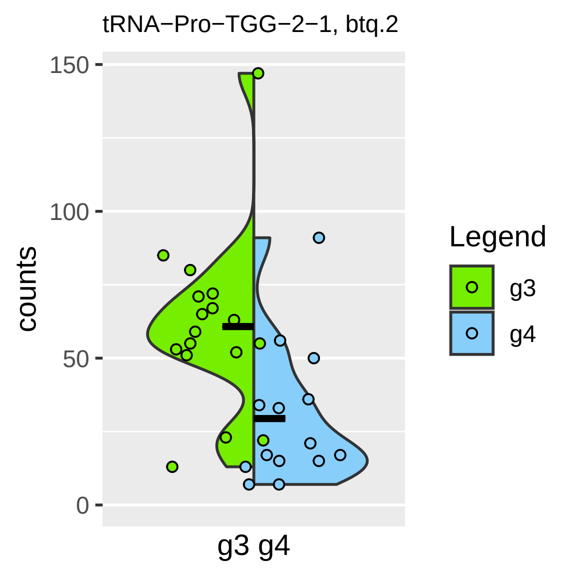
tRNA−Pro−TGG−2−1, btq.2
150
100
50
0
counts
g3 g4
Legend
g3
g4

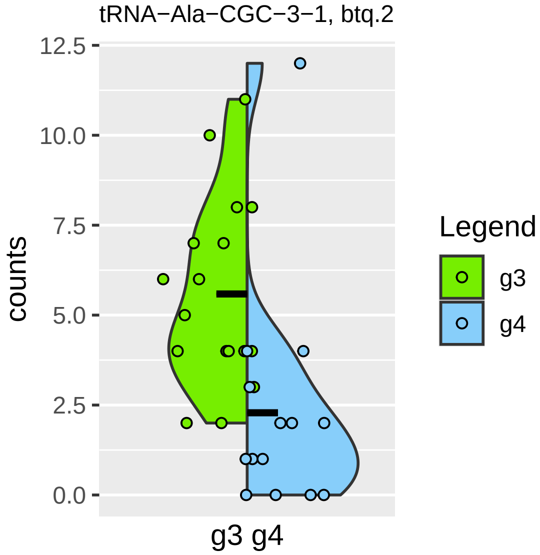

tRNA−Ala−CGC−3−1, btq.2
counts
12.5
10.0
7.5
5.0
2.5
0.0
g3 g4
Legend
g3
g4

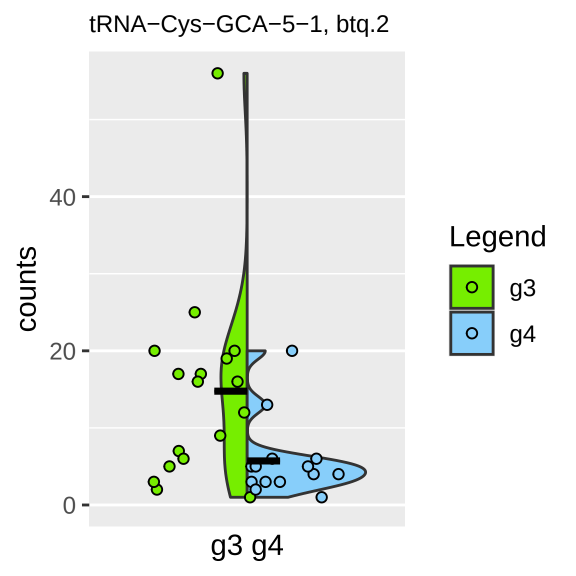

tRNA−Cys−GCA−5−1, btq.2
counts
0
20
40
g3 g4
Legend
g3
g4

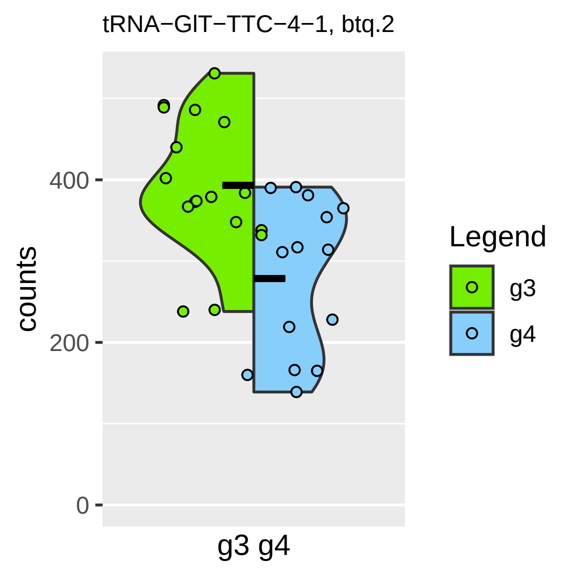
tRNA−GlT−TTC−4−1, btq.2
counts
0
200
400
g3 g4
Legend
g3
g4

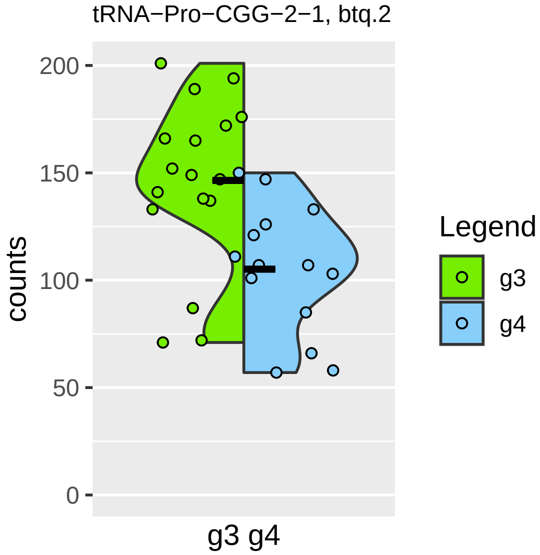

tRNA−Pro−CGG−2−1, btq.2
counts
200
150
100
50
0
g3 g4
Legend
g3
g4

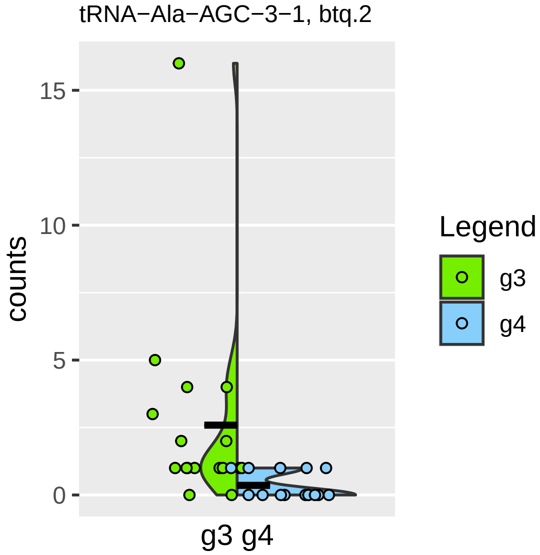

tRNA−Ala−AGC−3−1, btq.2
counts
15
10
5
0
g3 g4
Legend
g3
g4

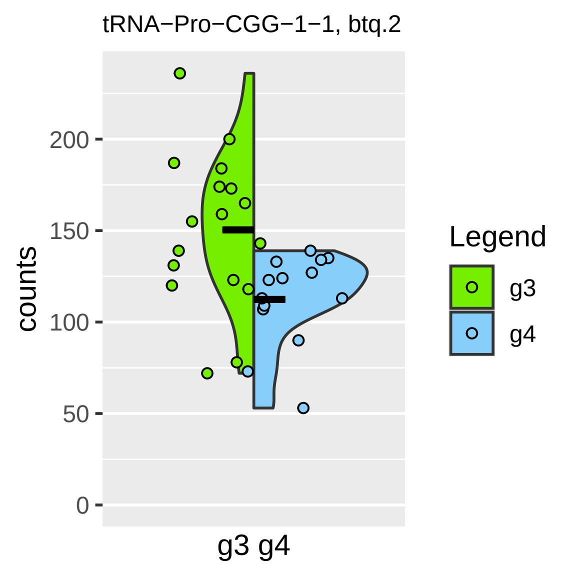
tRNA−Pro−CGG−1−1, btq.2
counts
200
150
100
50
0
g3 g4
Legend
g3
g4

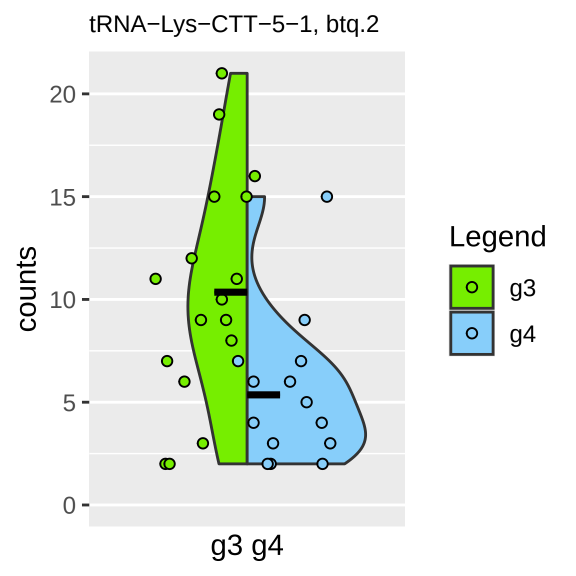
tRNA−Lys−CTT−5−1, btq.2
counts
0
5
10
15
20
g3 g4
Legend
g3
g4

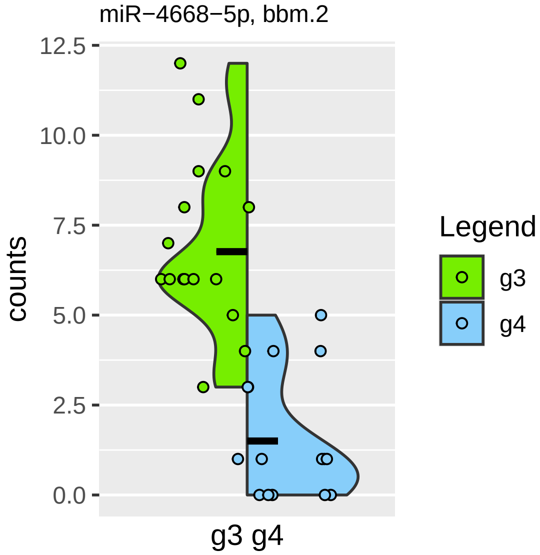

miR−4668−5p, bbm.2
counts
12.5
10.0
7.5
5.0
2.5
0.0
g3 g4
Legend
g3
g4

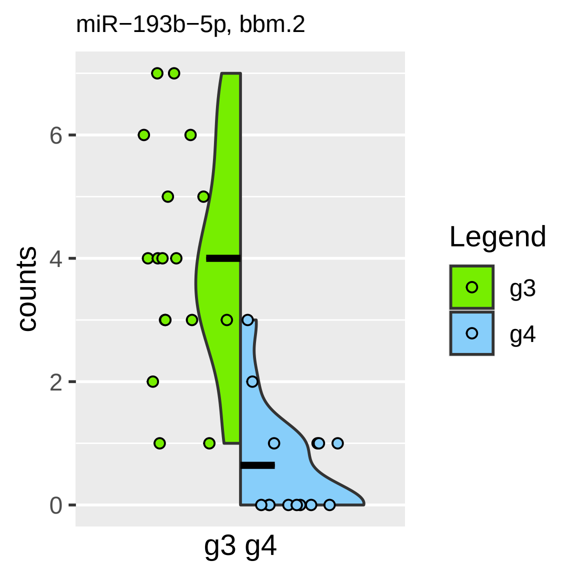

miR−193b−5p, bbm.2
counts
0
2
4
6
g3 g4
Legend
g3
g4

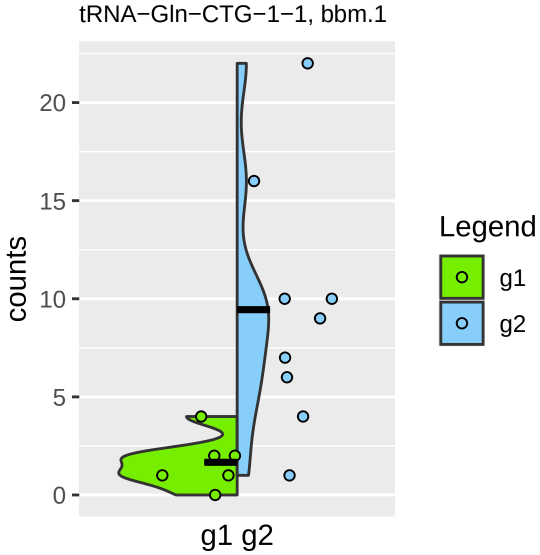

tRNA−Gln−CTG−1−1, bbm.1
counts
0
5
10
15
20
g1 g2
Legend
g1
g2

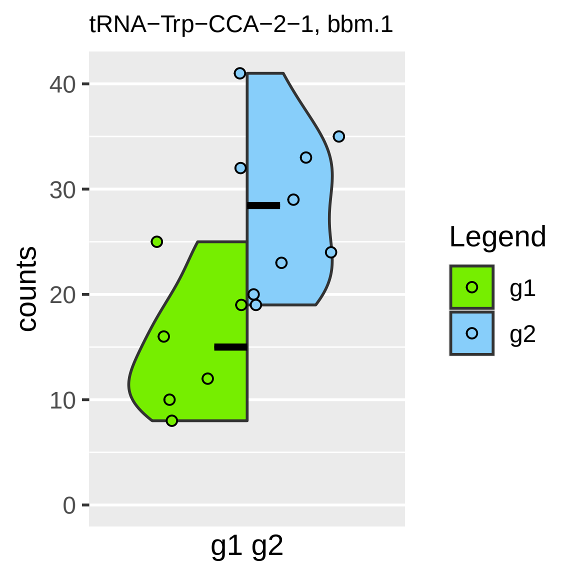

tRNA−Trp−CCA−2−1, bbm.1
counts
0
10
20
30
40
g1 g2
Legend
g1
g2

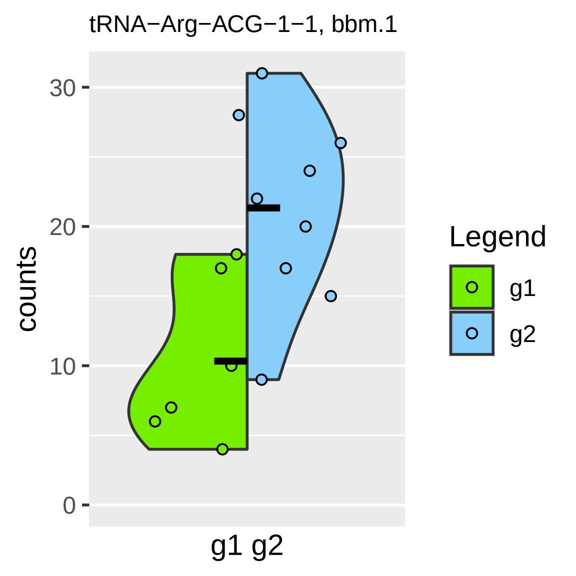
tRNA−Arg−ACG−1−1, bbm.1
counts
30
20
10
0
g1 g2
Legend
g1
g2

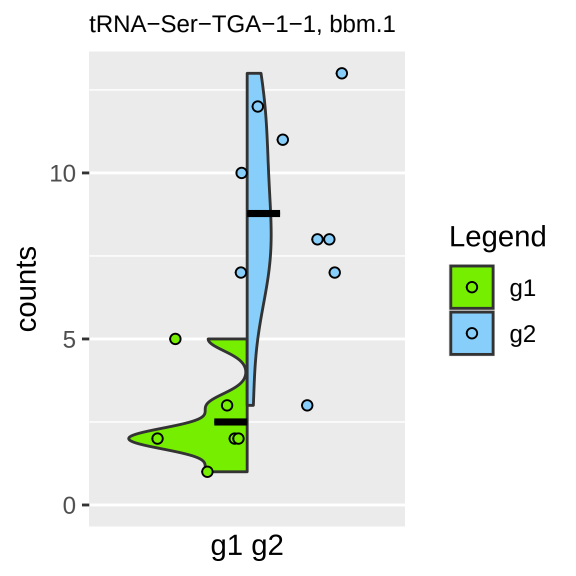

tRNA−Ser−TGA−1−1, bbm.1
counts
0
5
10
g1 g2
Legend
g1
g2

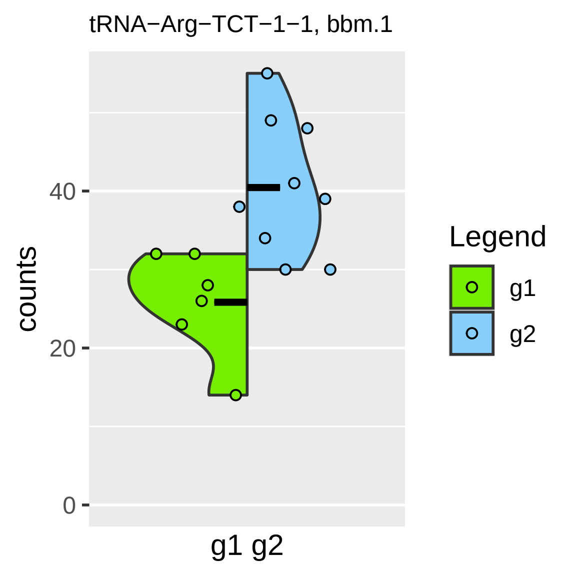

tRNA−Arg−TCT−1−1, bbm.1
counts
0
20
40
g1 g2
Legend
g1
g2

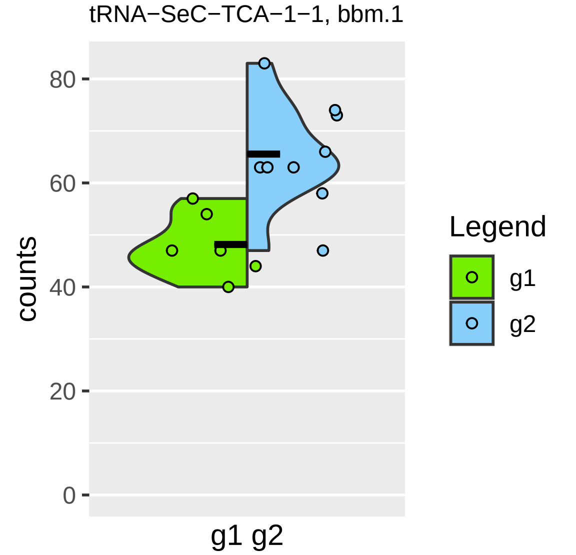

tRNA−SeC−TCA−1−1, bbm.1
counts
0
20
40
60
80
g1 g2
Legend
g1
g2

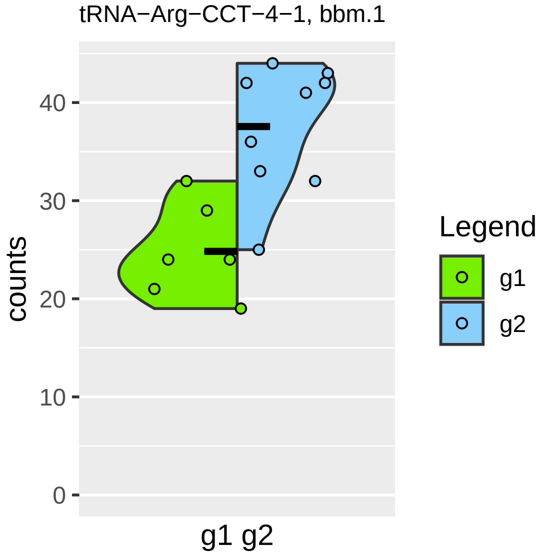
tRNA−Arg−CCT−4−1, bbm.1
counts
0
10
20
30
40
g1 g2
Legend
g1
g2

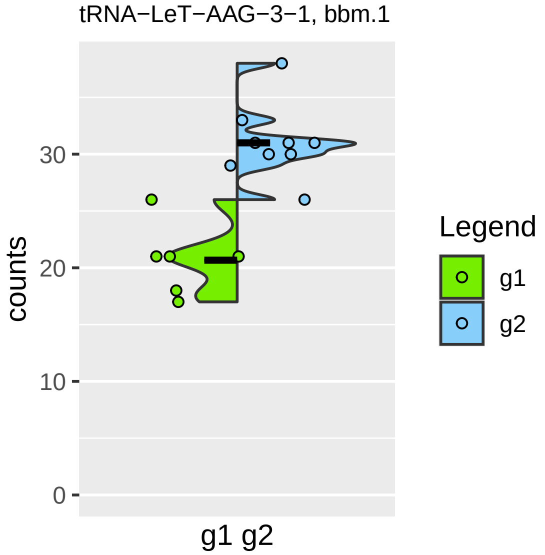
tRNA−LeT−AAG−3−1, bbm.1
counts
30
20
10
0
g1 g2
Legend
g1
g2

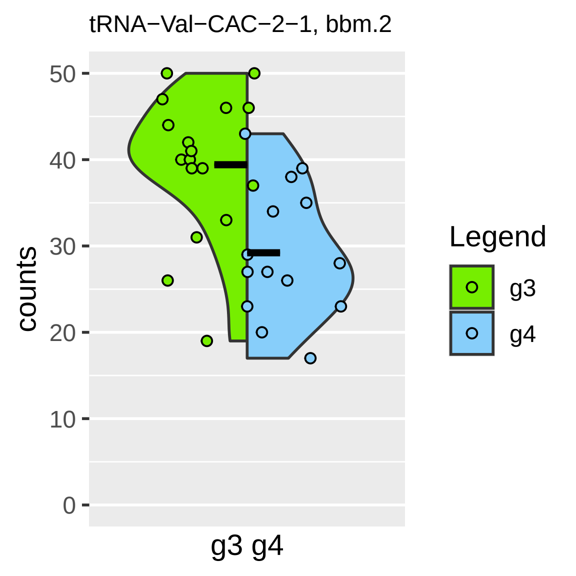

tRNA−Val−CAC−2−1, bbm.2
counts
50
40
30
20
10
0
g3 g4
Legend
g3
g4

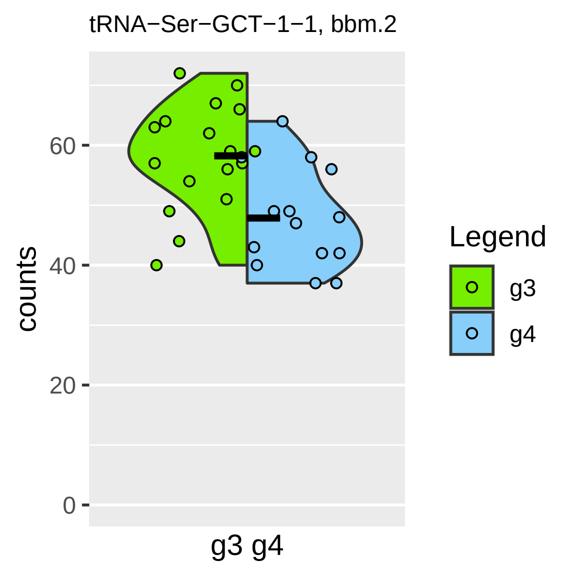

tRNA−Ser−GCT−1−1, bbm.2
counts
60
40
20
0
g3 g4
Legend
g3
g4

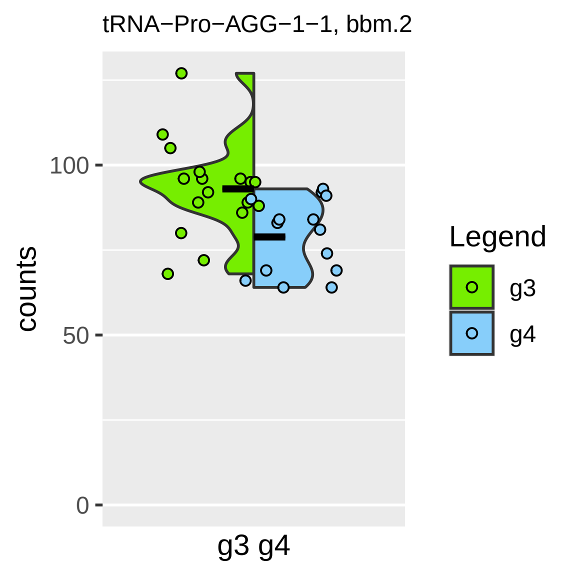

tRNA−Pro−AGG−1−1, bbm.2
counts
100
50
0
g3 g4
Legend
g3
g4

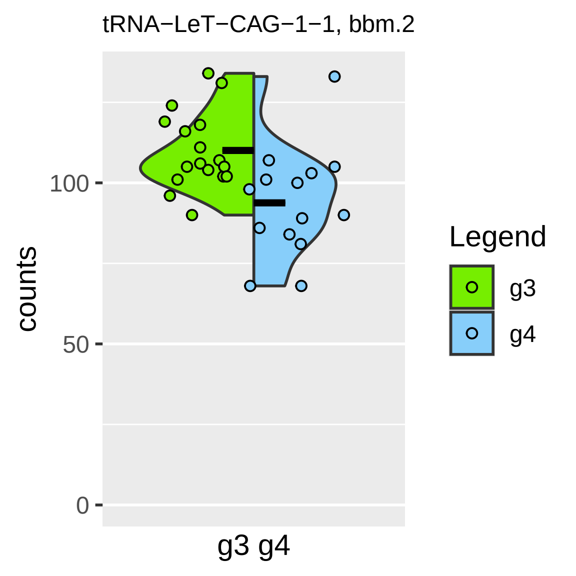
tRNA−LeT−CAG−1−1, bbm.2
counts
100
50
0
g3 g4
Legend
g3
g4

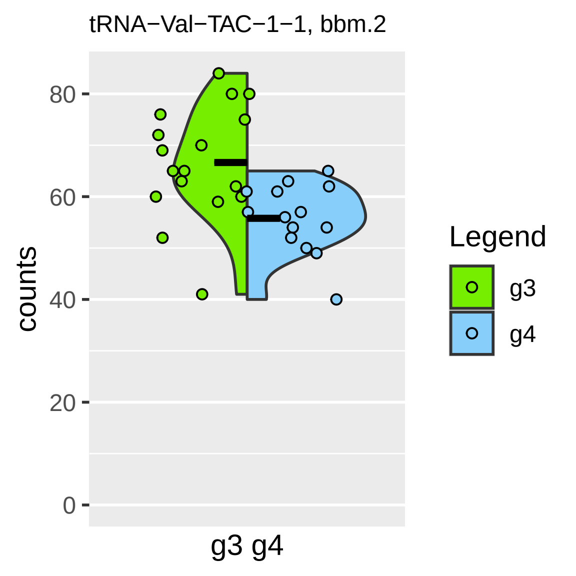

tRNA−Val−TAC−1−1, bbm.2
counts
80
60
40
20
0
g3 g4
Legend
g3
g4

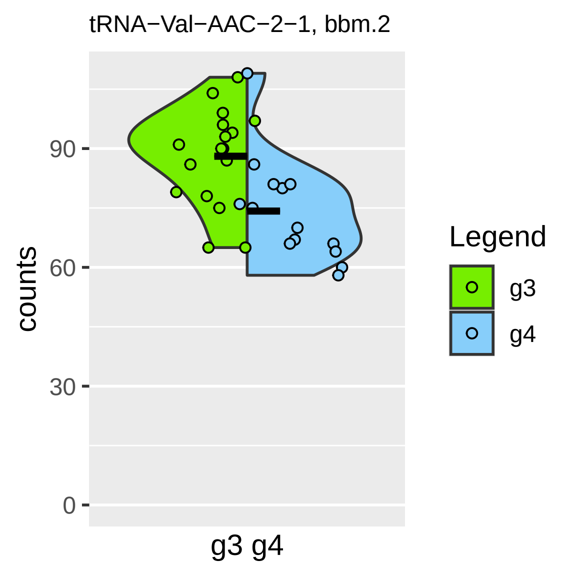
tRNA−Val−AAC−2−1, bbm.2
counts
0
30
60
90
g3 g4
Legend
g3
g4

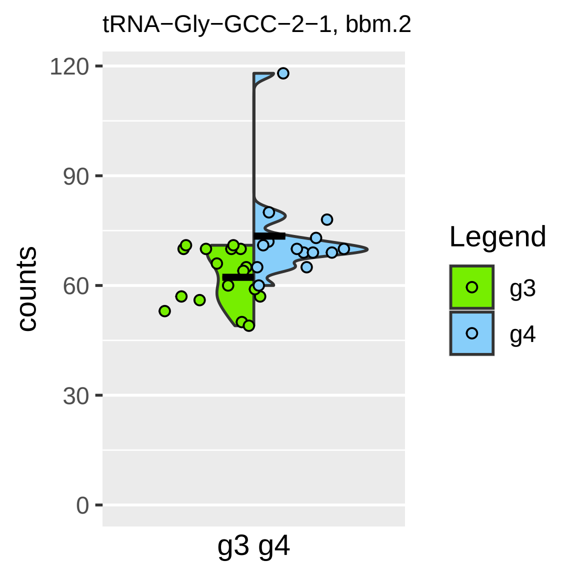

tRNA−Gly−GCC−2−1, bbm.2
120
90
60
30
0
counts
g3 g4
Legend
g3
g4

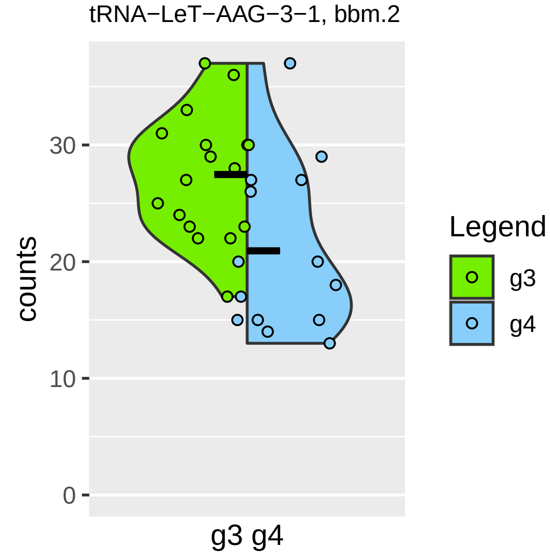

tRNA−LeT−AAG−3−1, bbm.2
counts
30
20
10
0
g3 g4
Legend
g3
g4

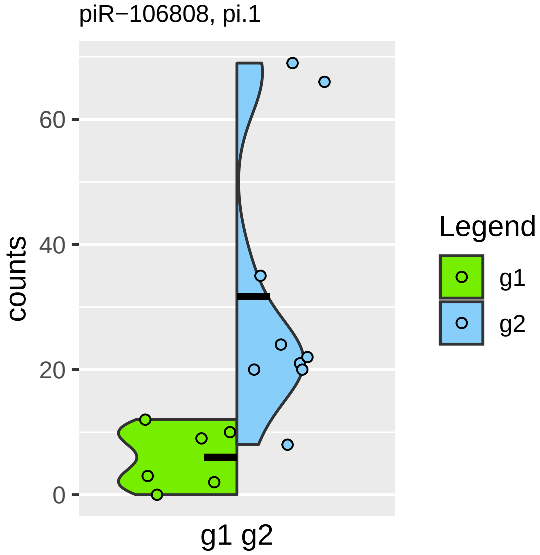

piR−106808, pi.1
counts
60
40
20
0
g1 g2
Legend
g1
g2

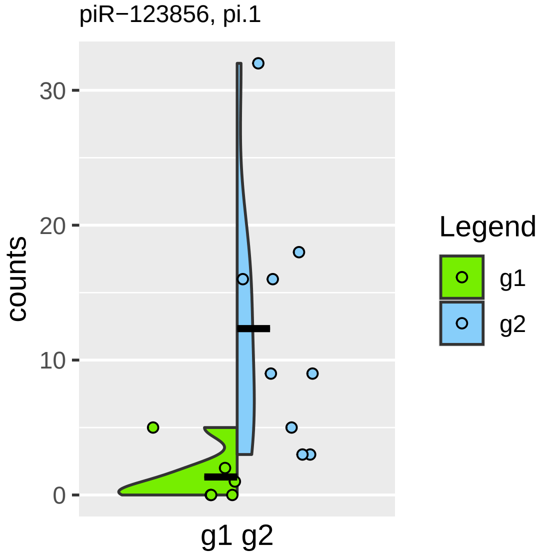

piR−123856, pi.1
counts
30
20
10
0
g1 g2
Legend
g1
g2

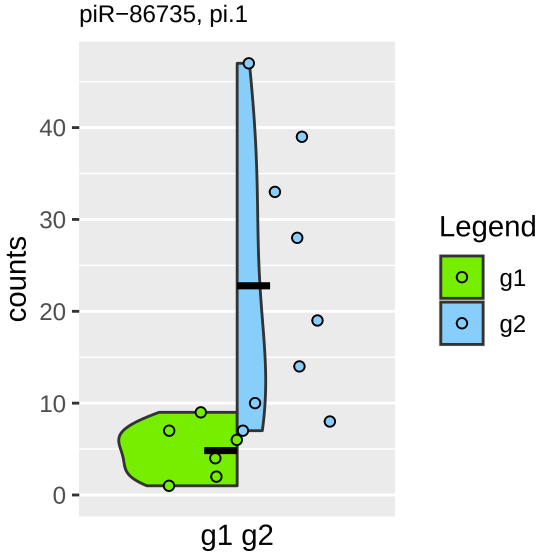

piR−86735, pi.1
counts
40
30
20
10
0
g1 g2
Legend
g1
g2

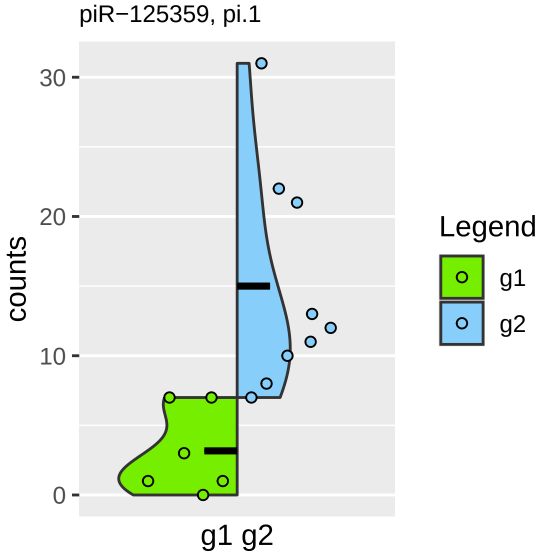
piR−125359, pi.1
counts
30
20
10
0
g1 g2
Legend
g1
g2

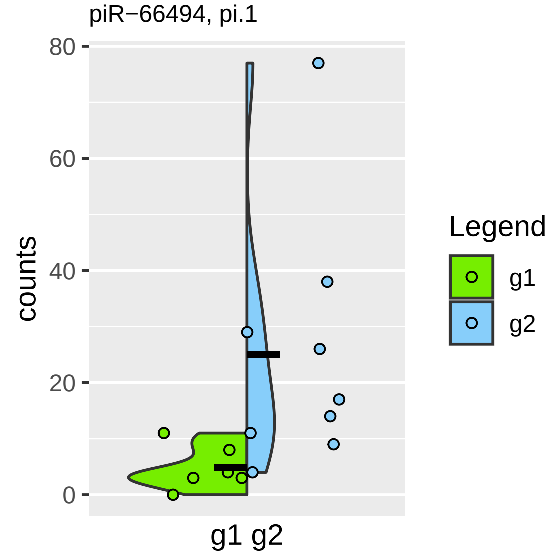

piR−66494, pi.1
80
60
40
20
0
counts
g1 g2
Legend
g1
g2

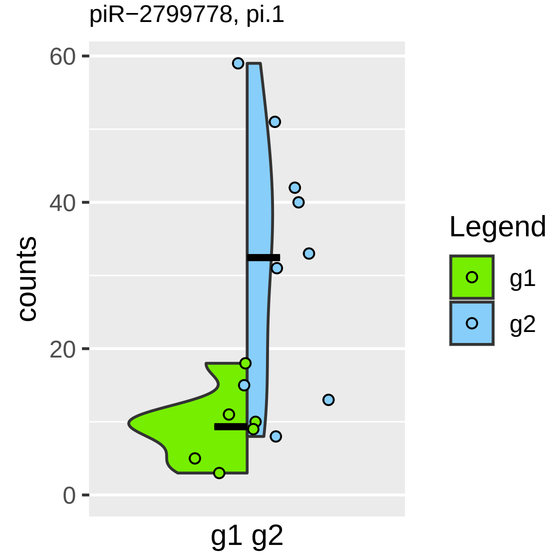
piR−2799778, pi.1
counts
60
40
20
0
g1 g2
Legend
g1
g2

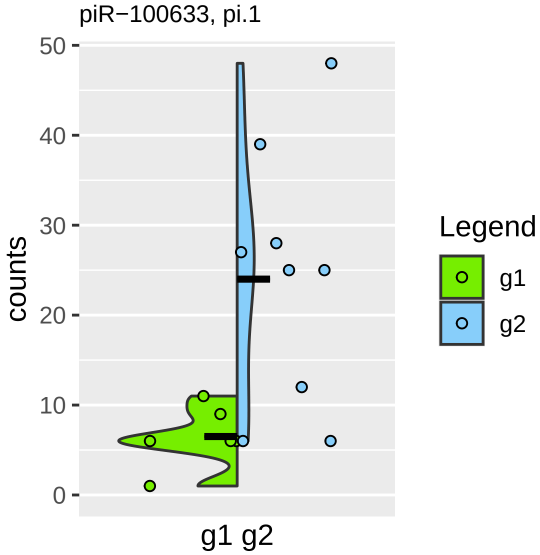

piR−100633, pi.1
counts
50
40
30
20
10
0
g1 g2
Legend
g1
g2

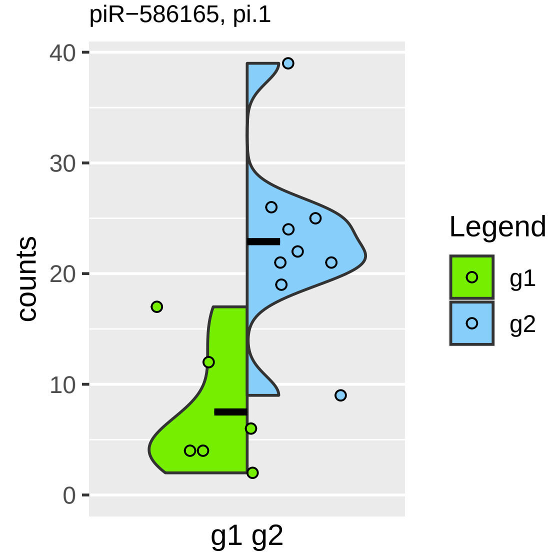
piR−586165, pi.1
counts
0
10
20
30
40
g1 g2
Legend
g1
g2

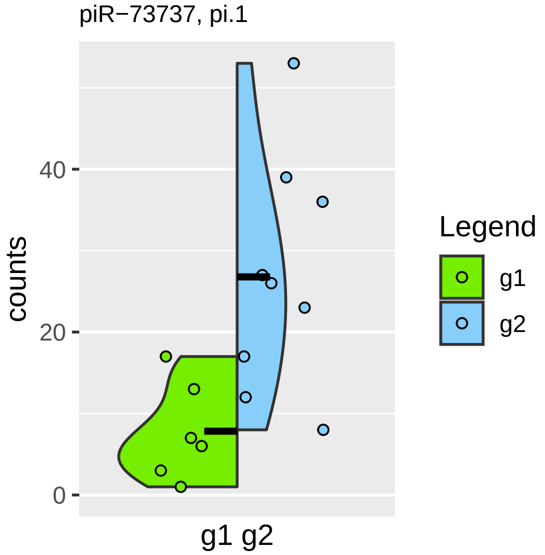
piR−73737, pi.1
counts
0
20
40
g1 g2
Legend
g1
g2

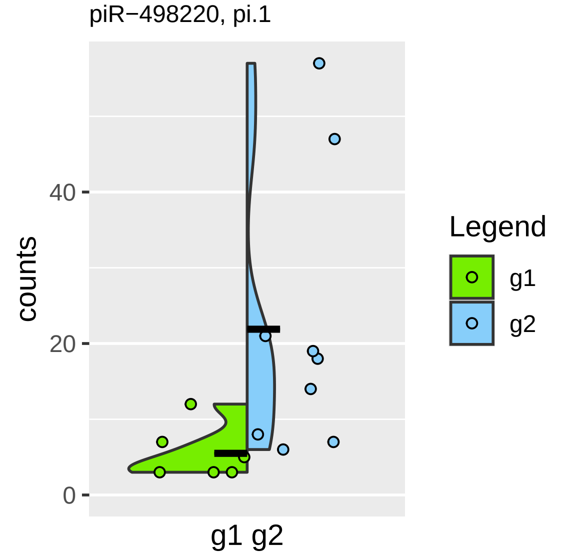

piR−498220, pi.1
counts
0
20
40
g1 g2
Legend
g1
g2

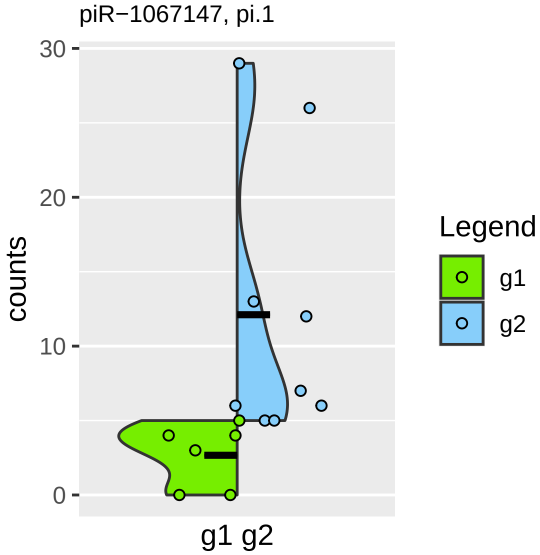

piR−1067147, pi.1
30
20
10
0
counts
g1 g2
Legend
g1
g2

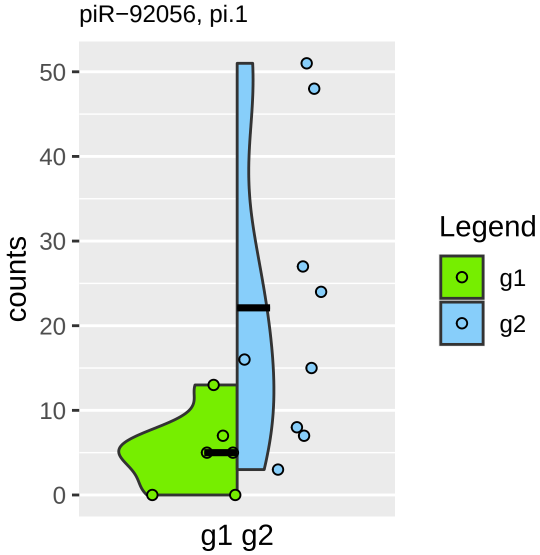
piR−92056, pi.1
counts
50
40
30
20
10
0
g1 g2
Legend
g1
g2

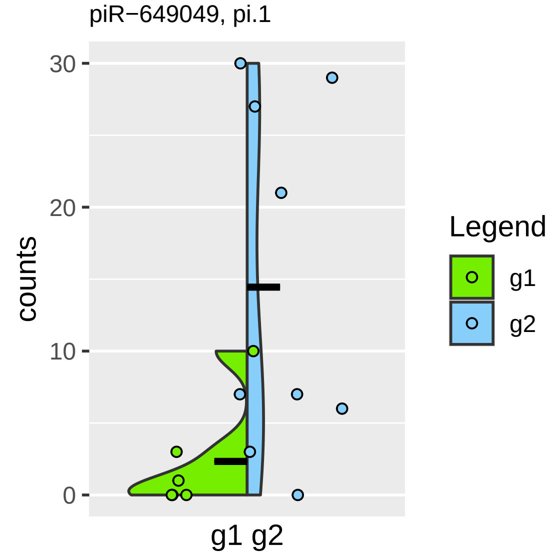

piR−649049, pi.1
30
20
10
0
counts
g1 g2
Legend
g1
g2

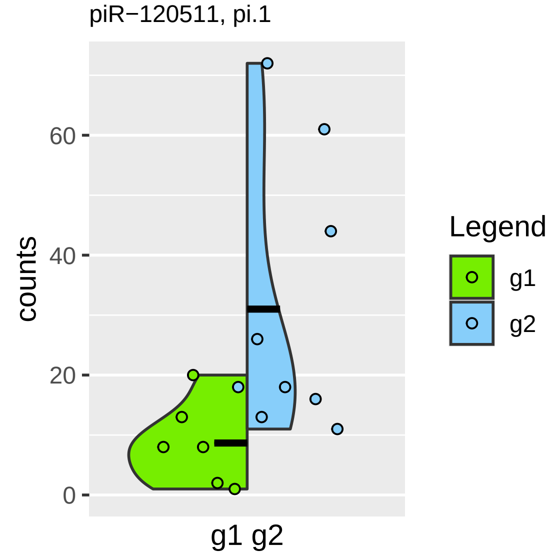
piR−120511, pi.1
counts
60
40
20
0
g1 g2
Legend
g1
g2

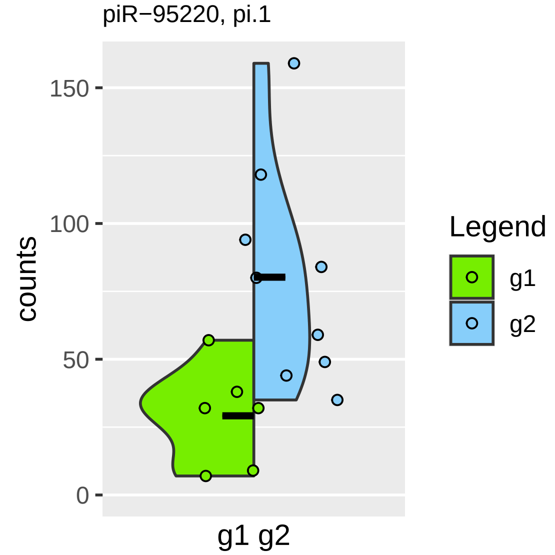

piR−95220, pi.1
150
100
50
0
counts
g1 g2
Legend
g1
g2

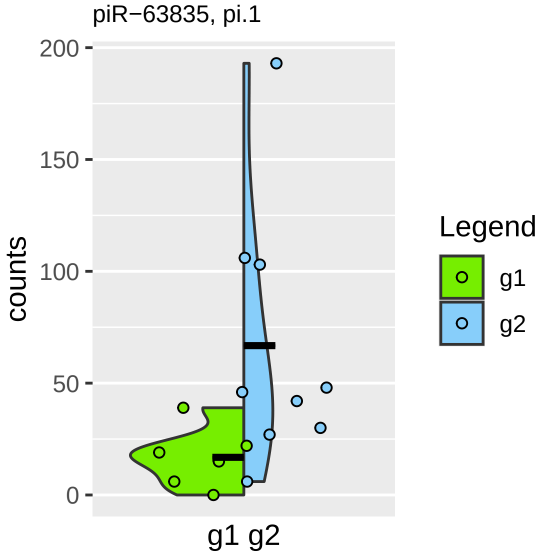

piR−63835, pi.1
counts
200
150
100
50
0
g1 g2
Legend
g1
g2

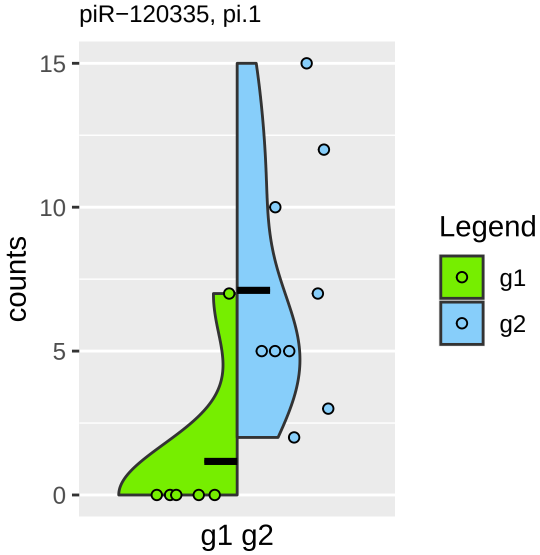

piR−120335, pi.1
counts
15
10
5
0
g1 g2
Legend
g1
g2

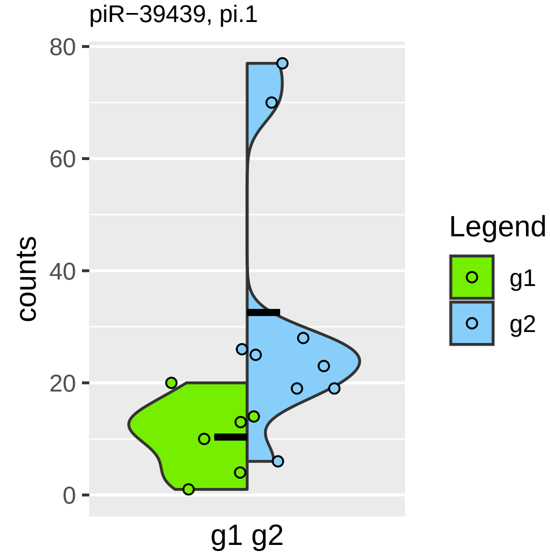

piR−39439, pi.1
80
60
40
20
0
counts
g1 g2
Legend
g1
g2

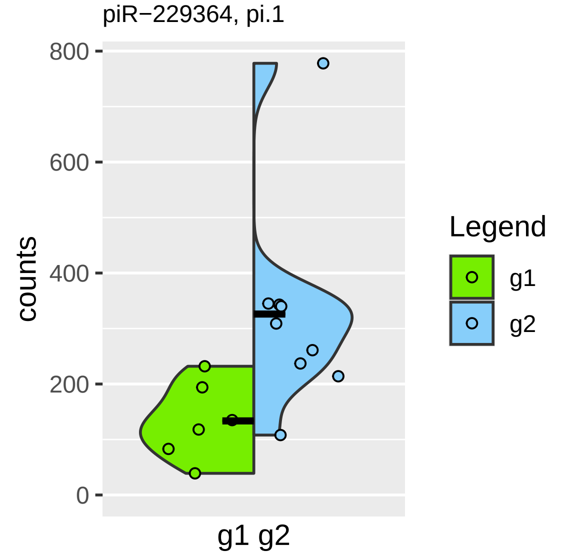

piR−229364, pi.1
800
600
400
200
0
counts
g1 g2
Legend
g1
g2

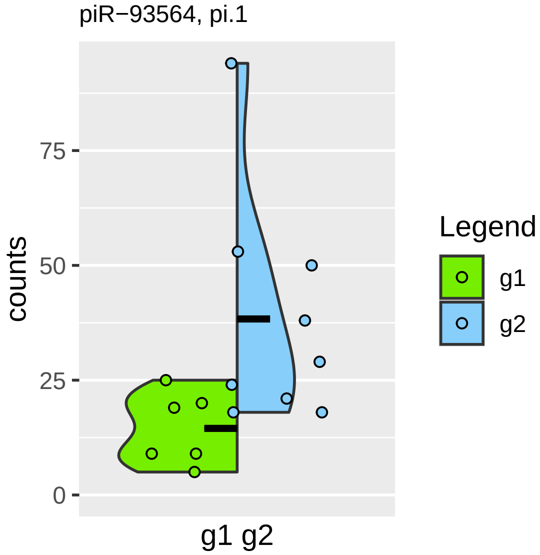

piR−93564, pi.1
counts
0
25
50
75
g1 g2
Legend
g1
g2

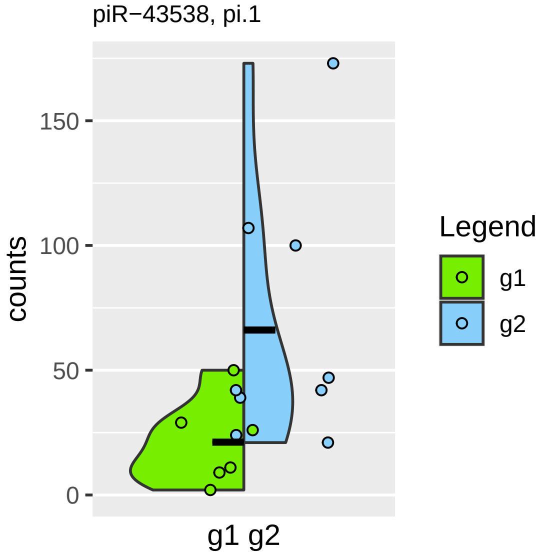
piR−43538, pi.1
counts
150
100
50
0
g1 g2
Legend
g1
g2

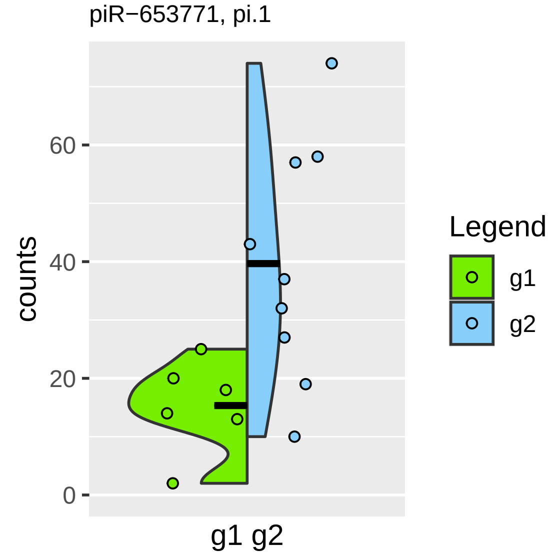

piR−653771, pi.1
counts
60
40
20
0
g1 g2
Legend
g1
g2

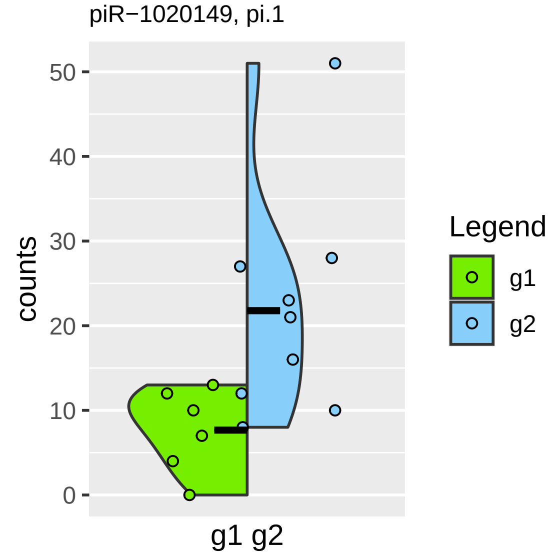
piR−1020149, pi.1
counts
50
40
30
20
10
0
g1 g2
Legend
g1
g2

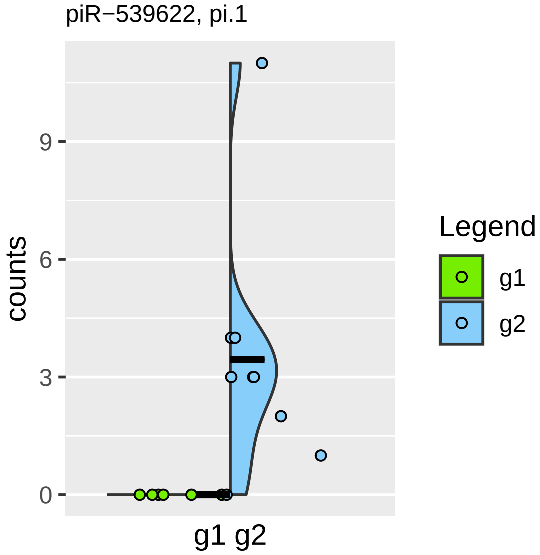
piR−539622, pi.1
counts
9
6
3
0
g1 g2
Legend
g1
g2

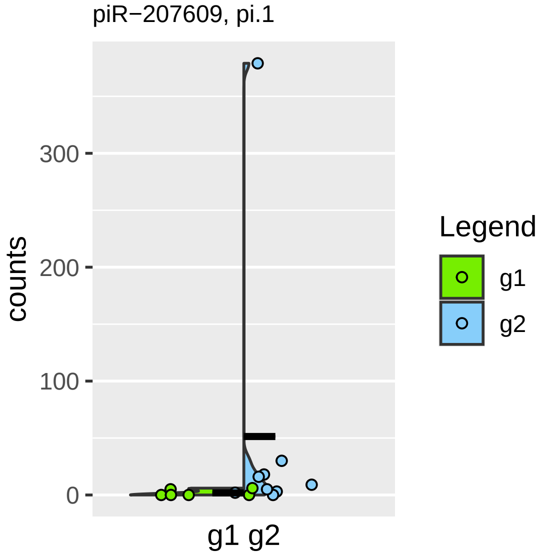

piR−207609, pi.1
counts
300
200
100
0
g1 g2
Legend
g1
g2

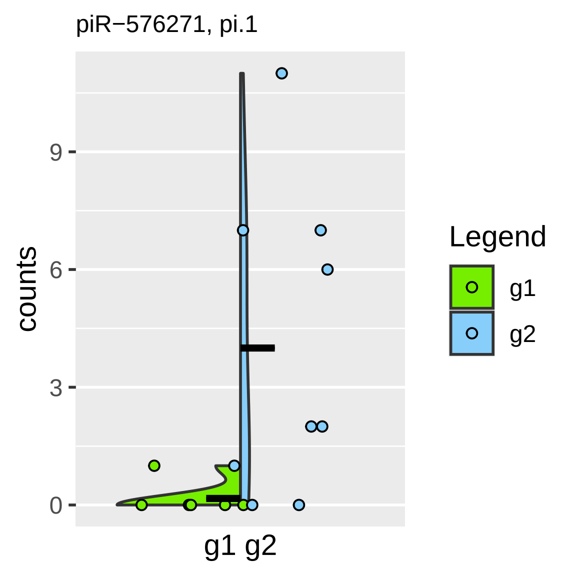

piR−576271, pi.1
counts
9
6
3
0
g1 g2
Legend
g1
g2

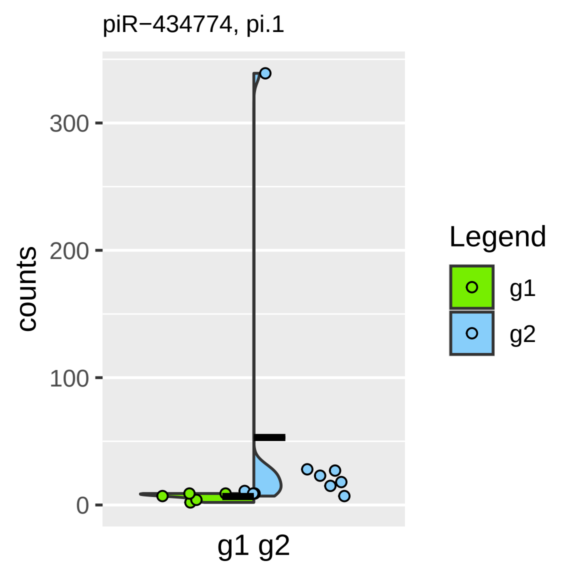

piR−434774, pi.1
counts
300
200
100
0
g1 g2
Legend
g1
g2

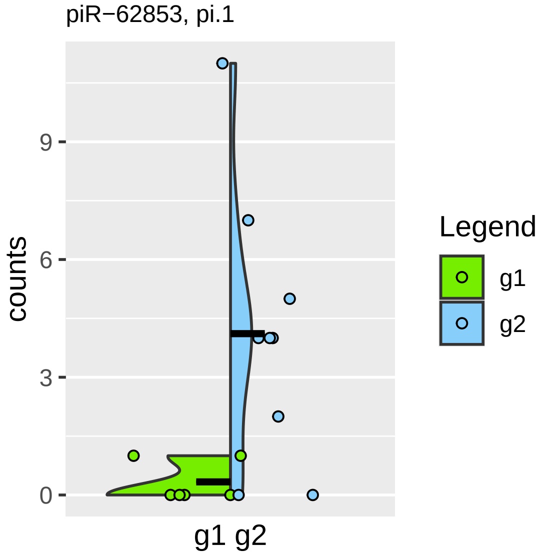
piR−62853, pi.1
counts
9
6
3
0
g1 g2
Legend
g1
g2

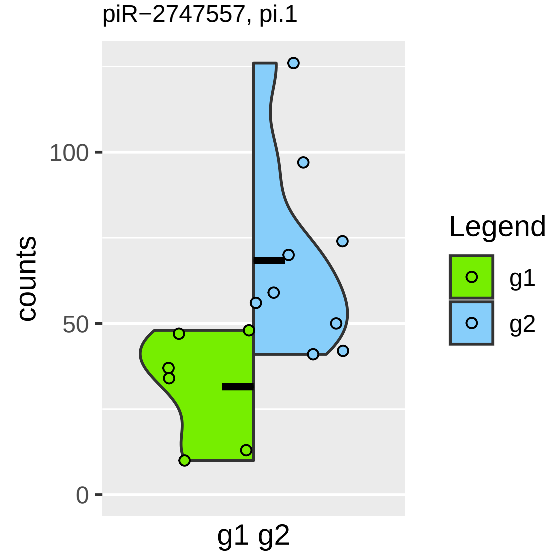
piR−2747557, pi.1
counts
100
50
0
g1 g2
Legend
g1
g2

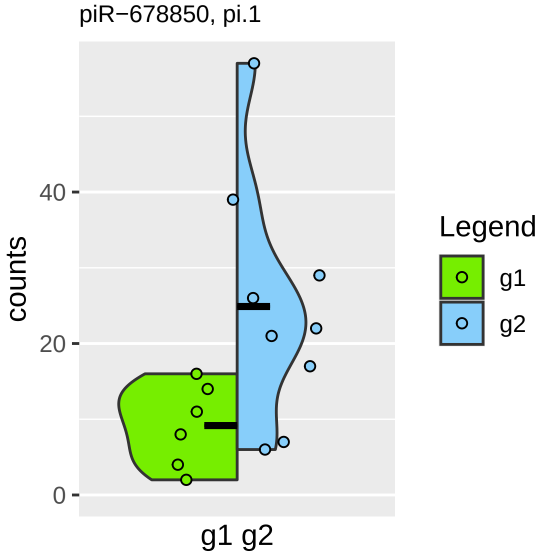

piR−678850, pi.1
counts
40
20
0
g1 g2
Legend
g1
g2

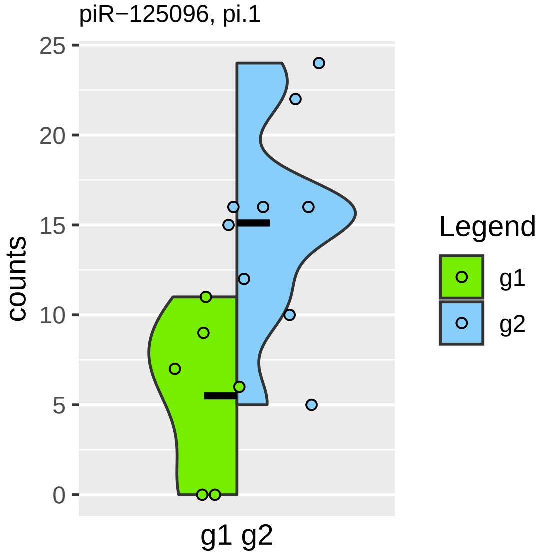

piR−125096, pi.1
25
20
15
10
5
0
counts
g1 g2
Legend
g1
g2

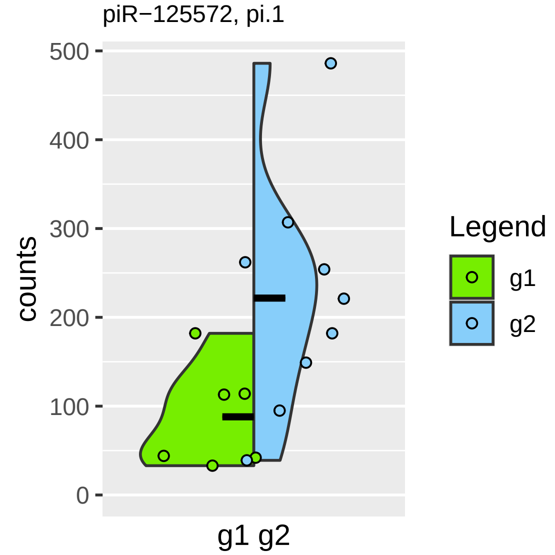
piR−125572, pi.1
counts
500
400
300
200
100
0
g1 g2
Legend
g1
g2

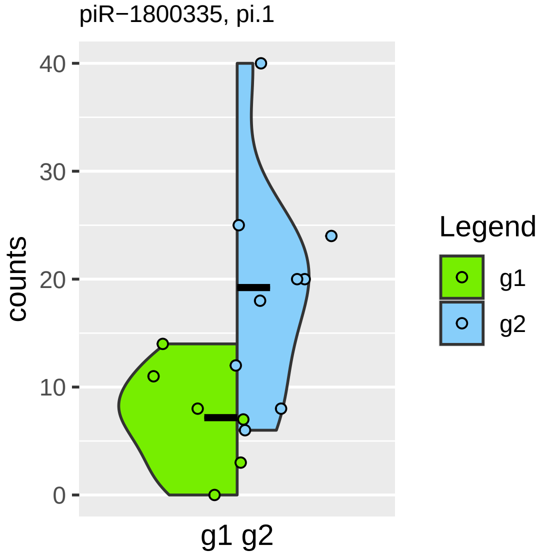
piR−1800335, pi.1
counts
40
30
20
10
0
g1 g2
Legend
g1
g2

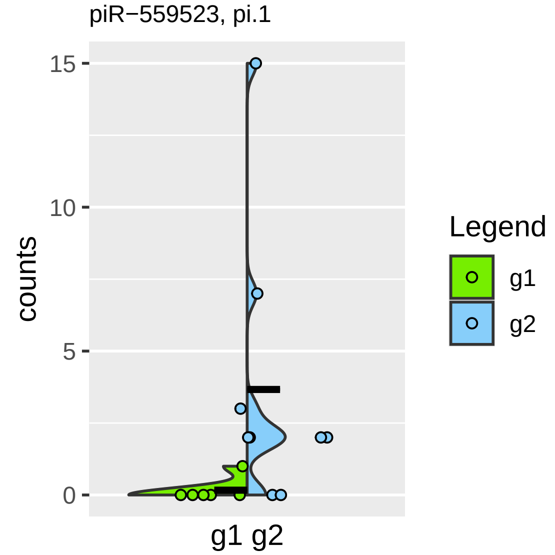

piR−559523, pi.1
counts
15
10
5
0
g1 g2
Legend
g1
g2

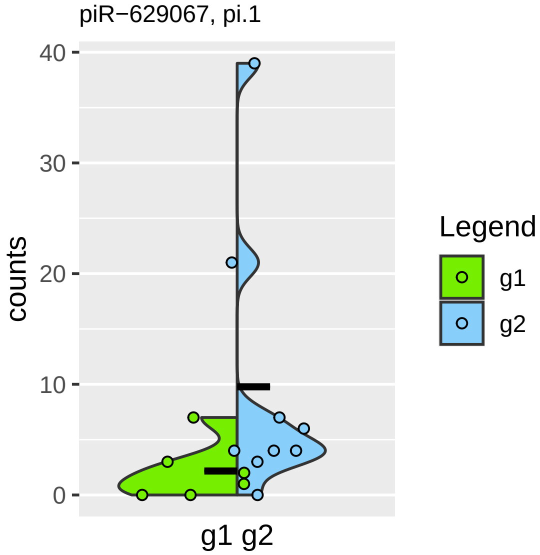
piR−629067, pi.1
40
30
20
10
0
counts
g1 g2
Legend
g1
g2

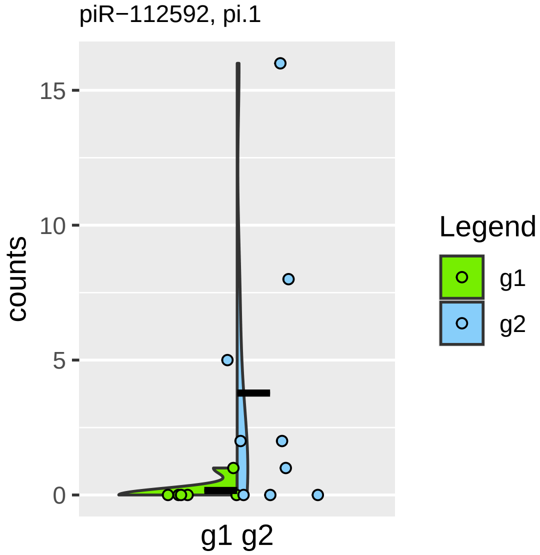

piR−112592, pi.1
counts
15
10
5
0
g1 g2
Legend
g1
g2

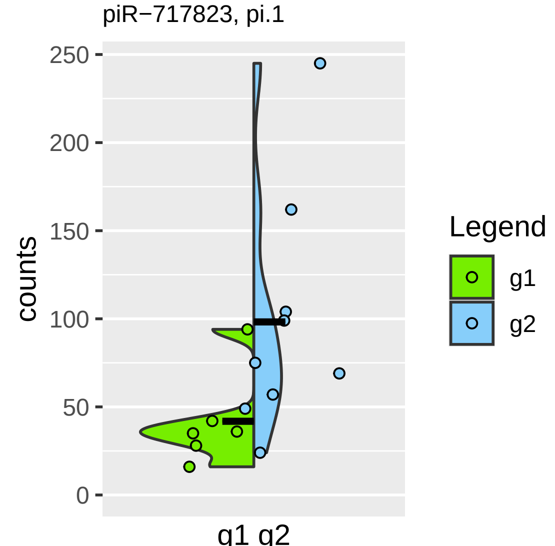

piR−717823, pi.1
counts
250
200
150
100
50
0
g1 g2
Legend
g1
g2

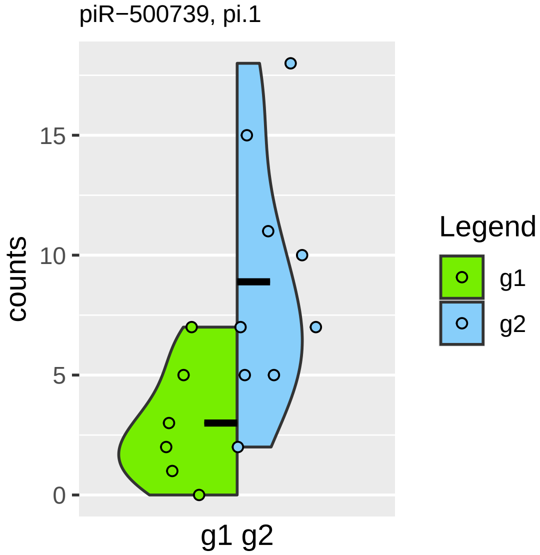

piR−500739, pi.1
counts
0
5
10
15
g1 g2
Legend
g1
g2

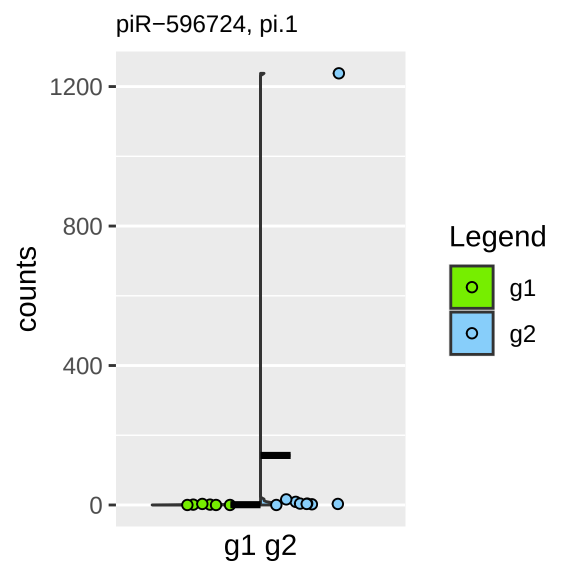

piR−596724, pi.1
counts
1200
800
400
0
g1 g2
Legend
g1
g2

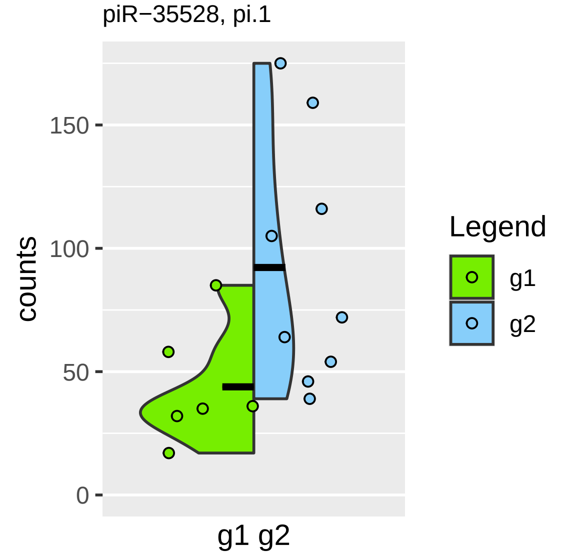

piR−35528, pi.1
counts
150
100
50
0
g1 g2
Legend
g1
g2

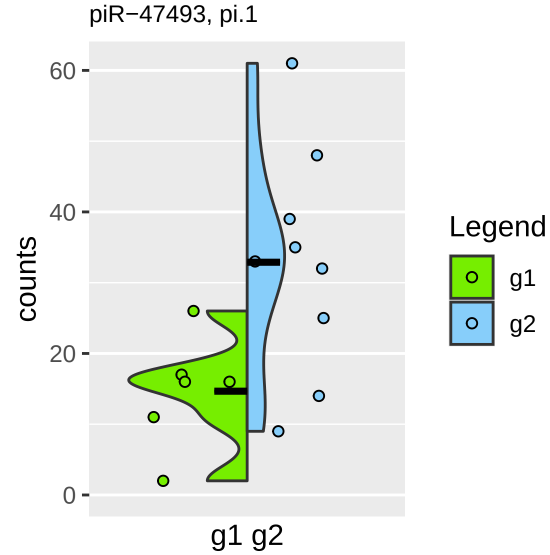
piR−47493, pi.1
counts
60
40
20
0
g1 g2
Legend
g1
g2

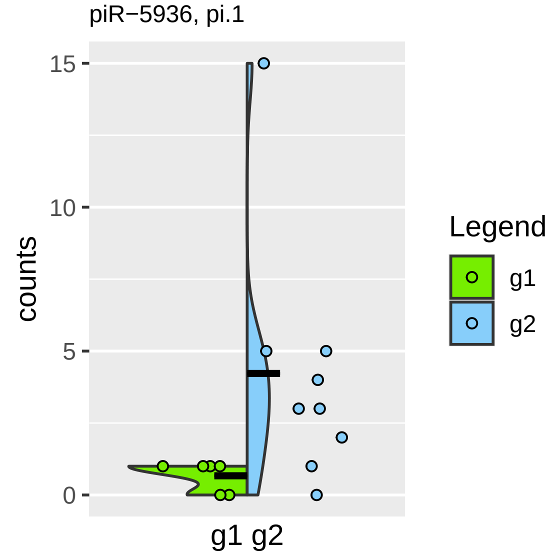
piR−5936, pi.1
15
10
5
0
counts
g1 g2
Legend
g1
g2

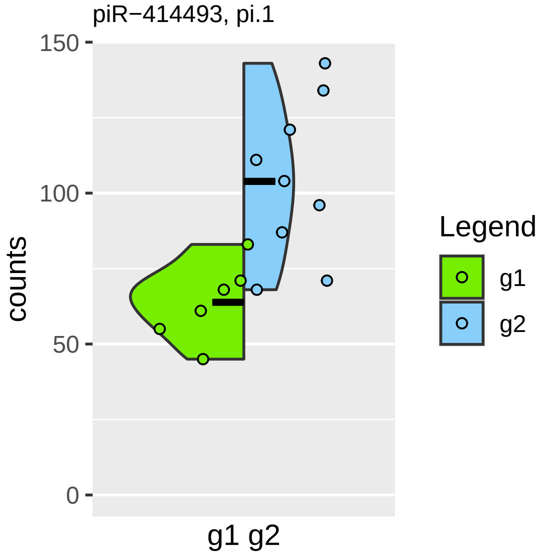
piR−414493, pi.1
150
100
50
0
counts
g1 g2
Legend
g1
g2

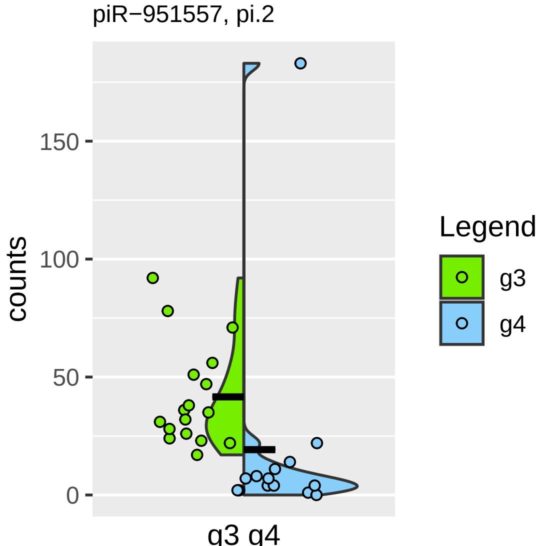
piR−951557, pi.2
counts
150
100
50
0
g3 g4
Legend
g3
g4

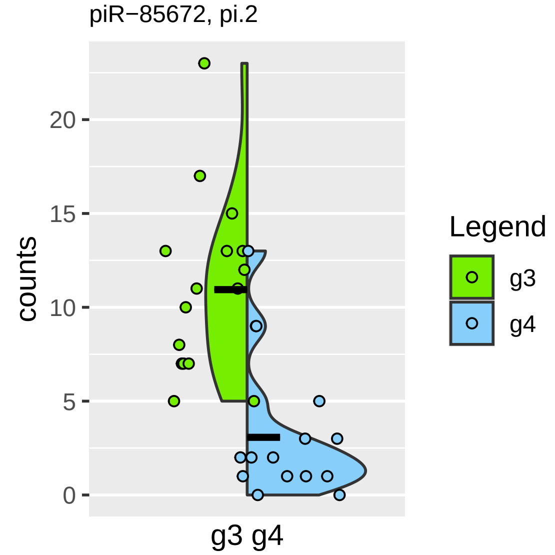

piR−85672, pi.2
counts
20
15
10
5
0
g3 g4
Legend
g3
g4

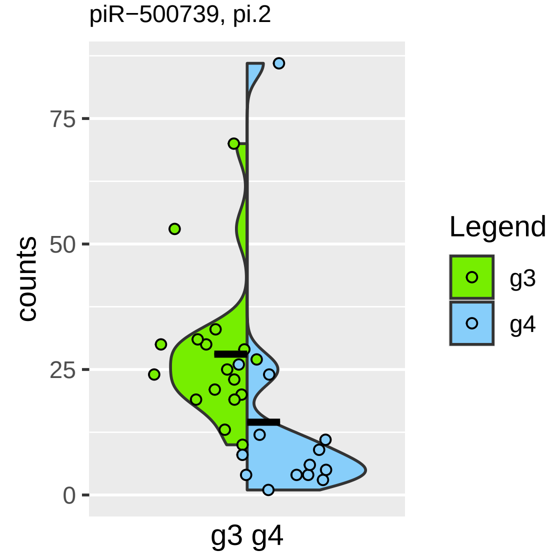

piR−500739, pi.2
counts
75
50
25
0
g3 g4
Legend
g3
g4

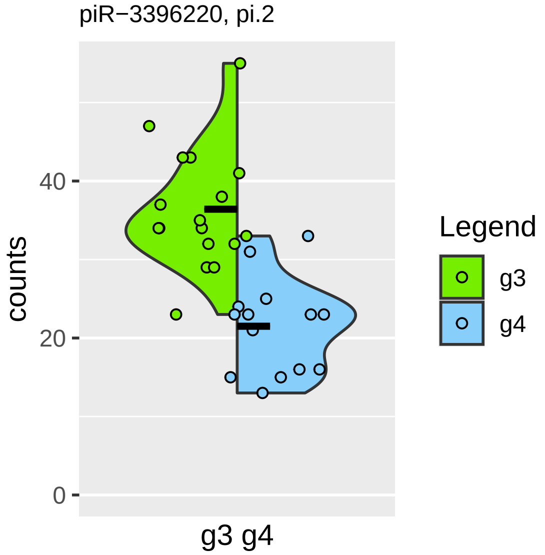
piR−3396220, pi.2
counts
40
20
0
g3 g4
Legend
g3
g4

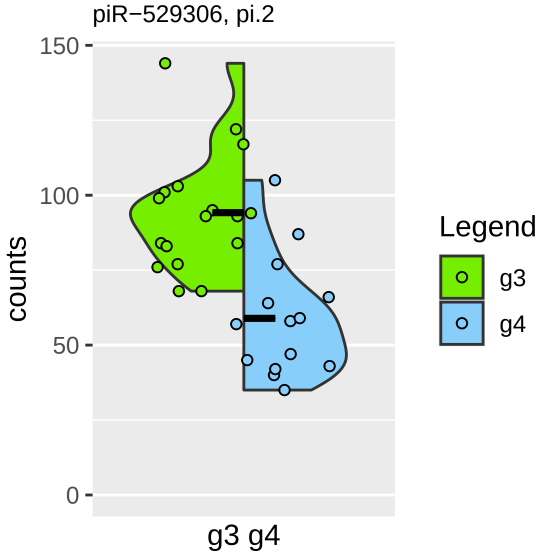
piR−529306, pi.2
150
100
50
0
counts
g3 g4
Legend
g3
g4

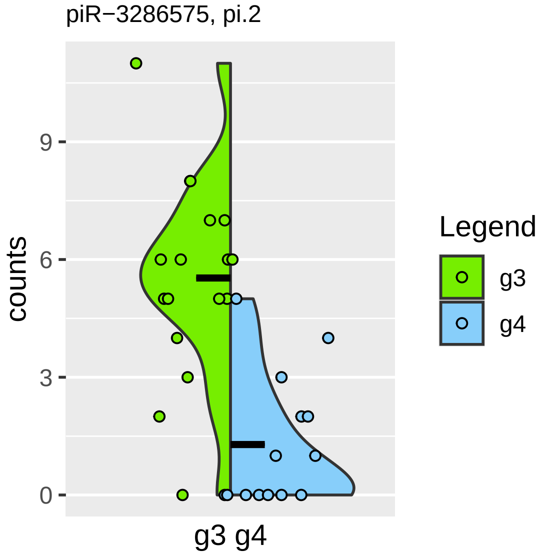
piR−3286575, pi.2
counts
9
6
3
0
g3 g4
Legend
g3
g4

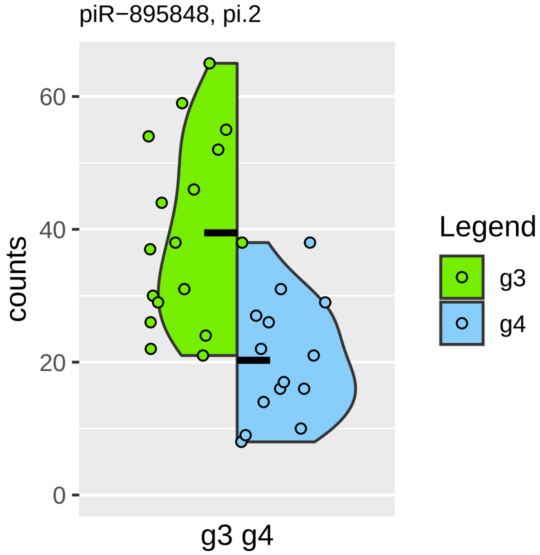
piR−895848, pi.2
counts
60
40
20
0
g3 g4
Legend
g3
g4

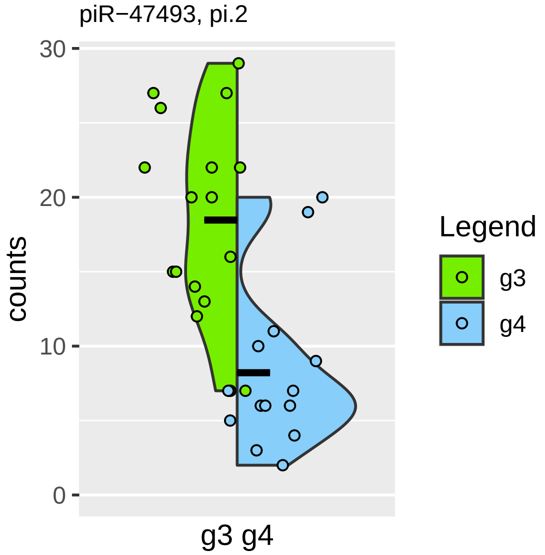

piR−47493, pi.2
counts
30
20
10
0
g3 g4
Legend
g3
g4

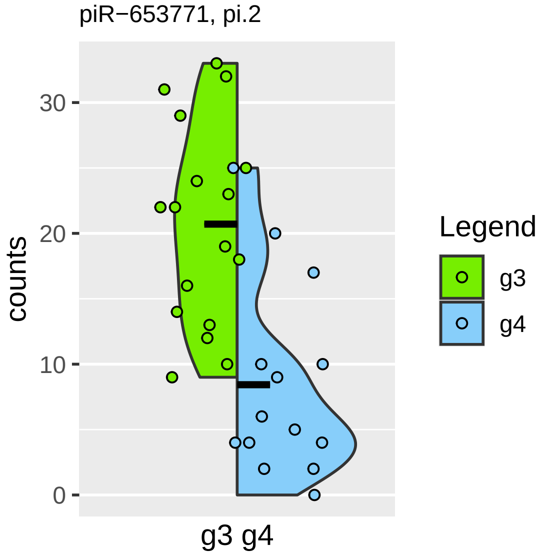
piR−653771, pi.2
counts
30
20
10
0
g3 g4
Legend
g3
g4

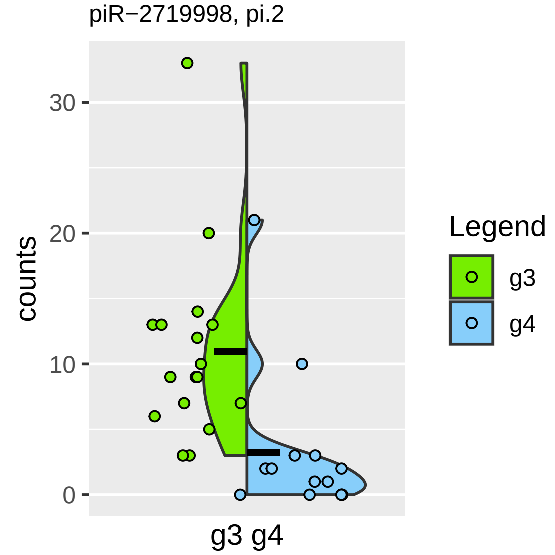

piR−2719998, pi.2
counts
30
20
10
0
g3 g4
Legend
g3
g4

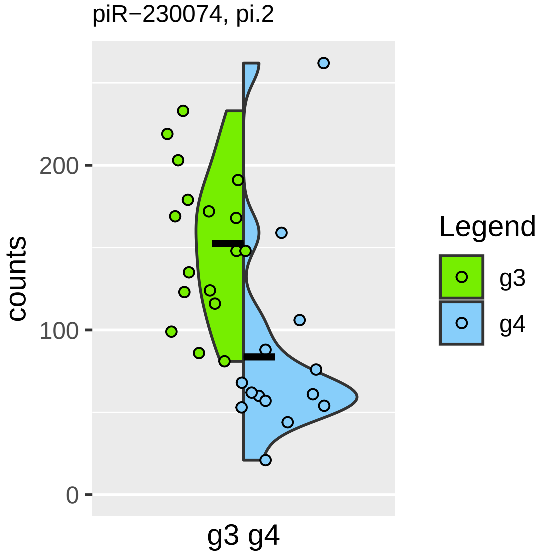

piR−230074, pi.2
200
100
0
counts
g3 g4
Legend
g3
g4

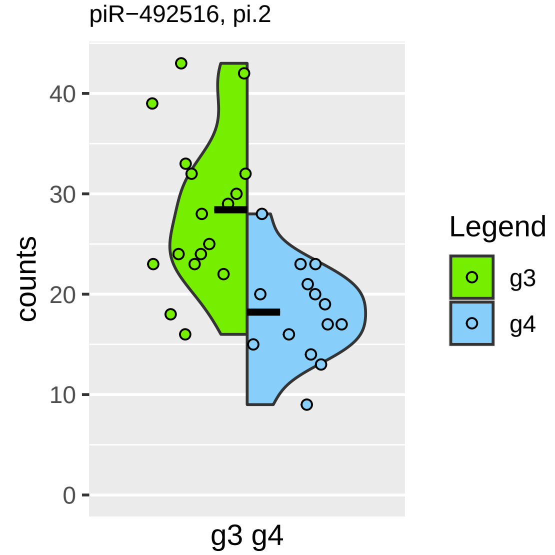

piR−492516, pi.2
counts
40
30
20
10
0
g3 g4
Legend
g3
g4

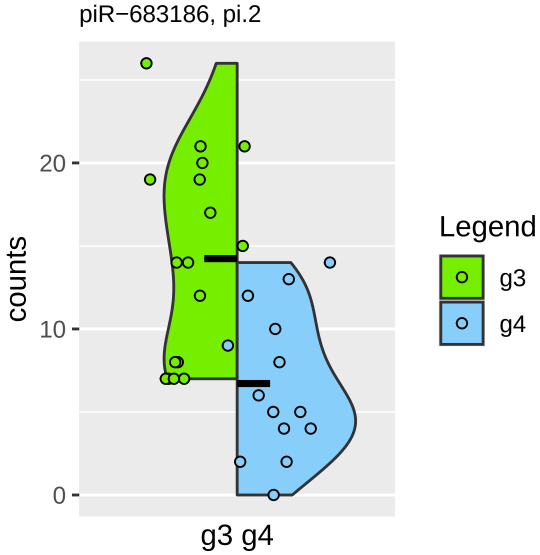

piR−683186, pi.2
counts
20
10
0
g3 g4
Legend
g3
g4

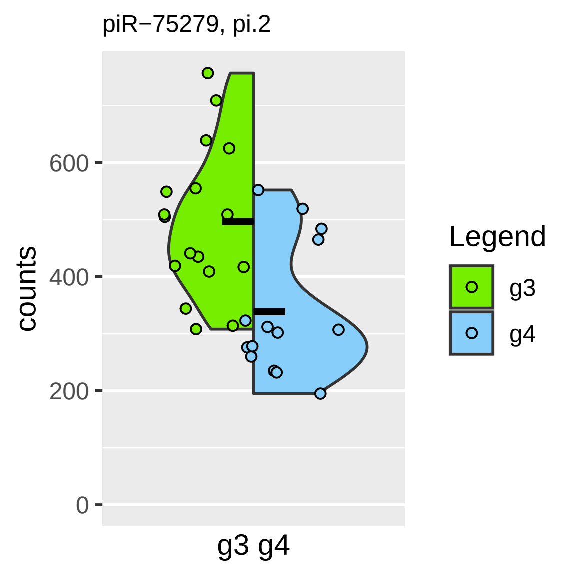
piR−75279, pi.2
counts
600
400
200
0
g3 g4
Legend
g3
g4

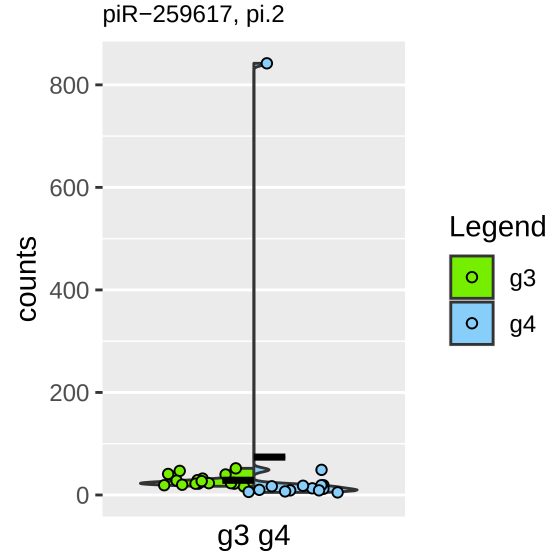

piR−259617, pi.2
800
600
400
200
0
counts
g3 g4
Legend
g3
g4

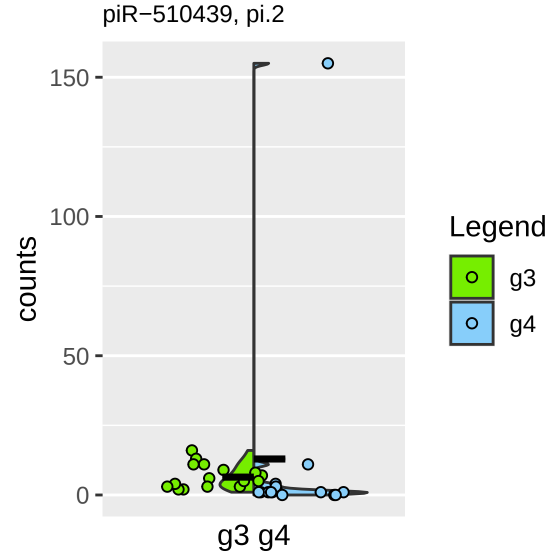

piR−510439, pi.2
150
100
50
0
counts
g3 g4
Legend
g3
g4

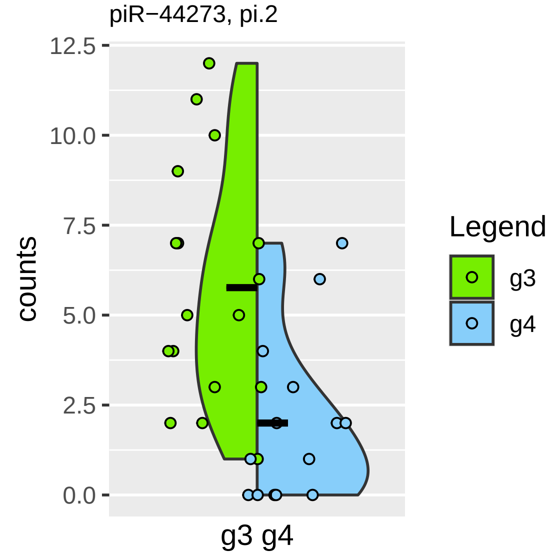

piR−44273, pi.2
12.5
10.0
7.5
5.0
2.5
0.0
counts
g3 g4
Legend
g3
g4

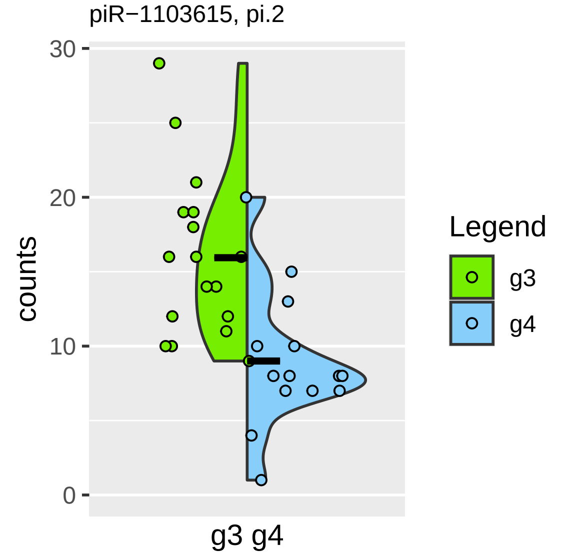

piR−1103615, pi.2
30
20
10
0
counts
g3 g4
Legend
g3
g4

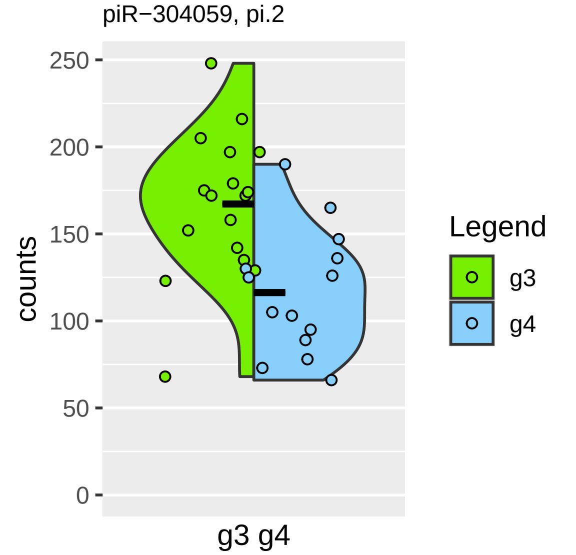
piR−304059, pi.2
counts
250
200
150
100
50
0
g3 g4
Legend
g3
g4

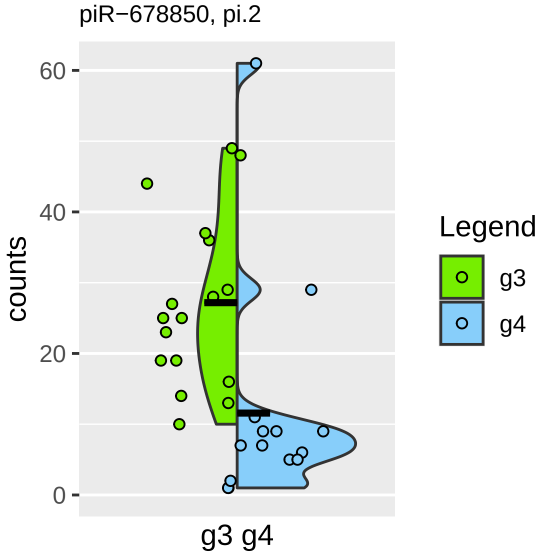

piR−678850, pi.2
counts
60
40
20
0
g3 g4
Legend
g3
g4

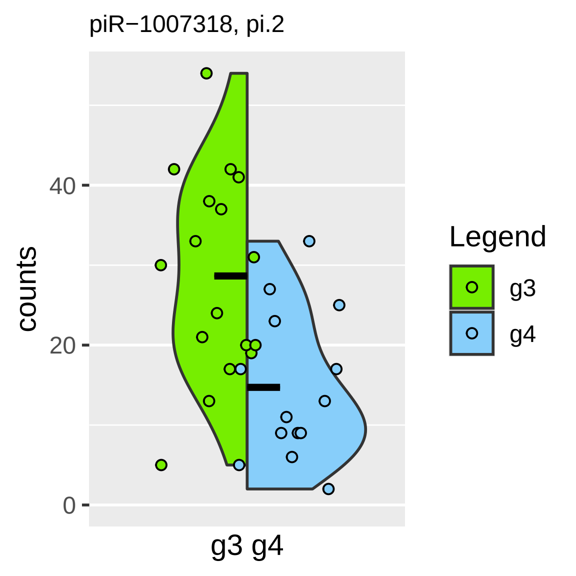
piR−1007318, pi.2
counts
0
20
40
g3 g4
Legend
g3
g4

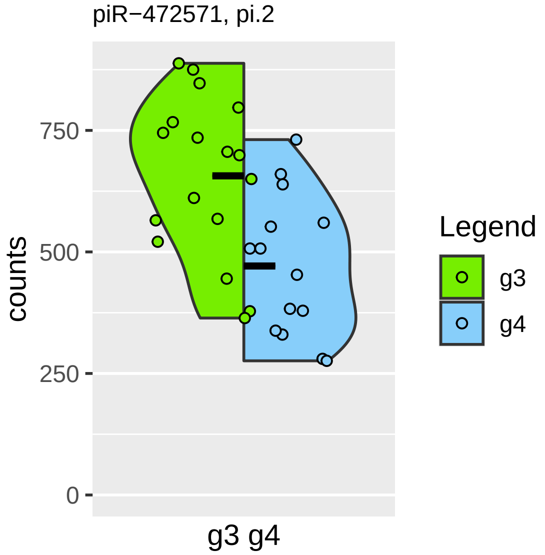

piR−472571, pi.2
counts
750
500
250
0
g3 g4
Legend
g3
g4

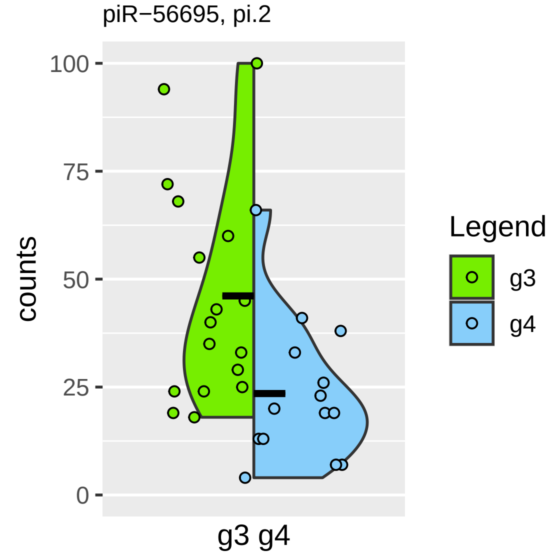
piR−56695, pi.2
counts
100
75
50
25
0
g3 g4
Legend
g3
g4

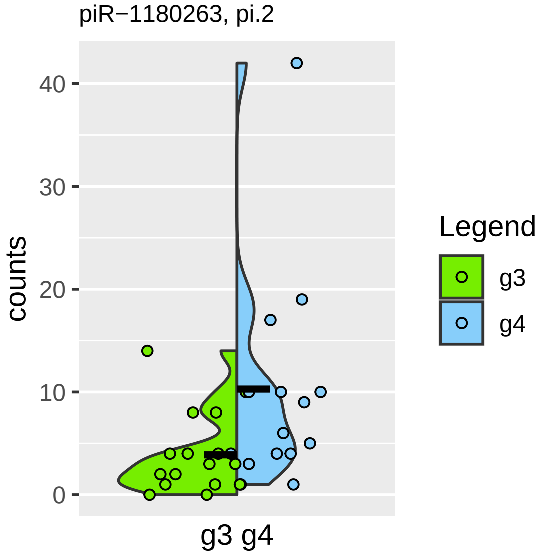
piR−1180263, pi.2
counts
0
10
20
30
40
g3 g4
Legend
g3
g4

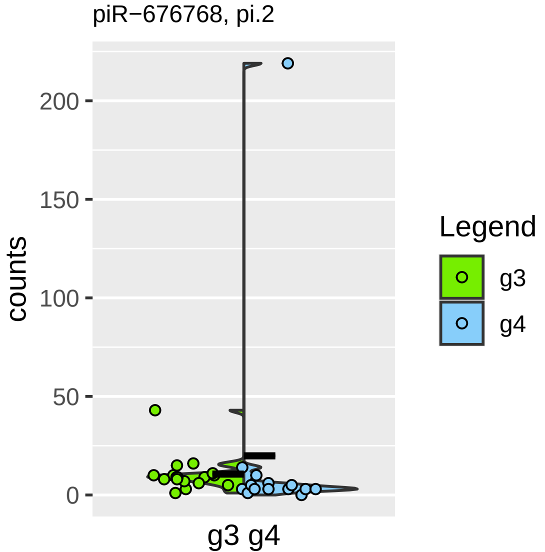

piR−676768, pi.2
counts
200
150
100
50
0
g3 g4
Legend
g3
g4

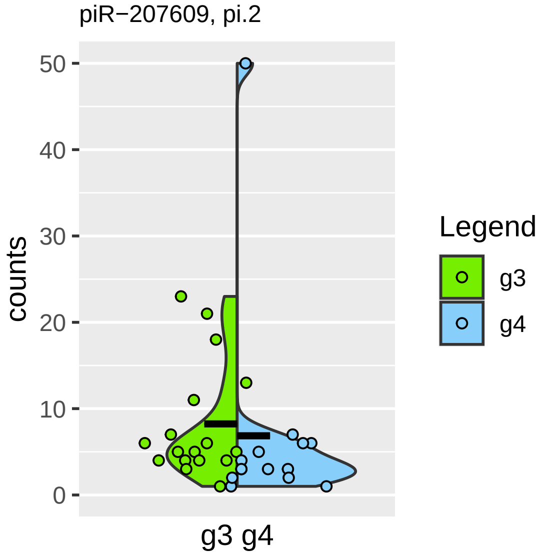

piR−207609, pi.2
counts
50
40
30
20
10
0
g3 g4
Legend
g3
g4

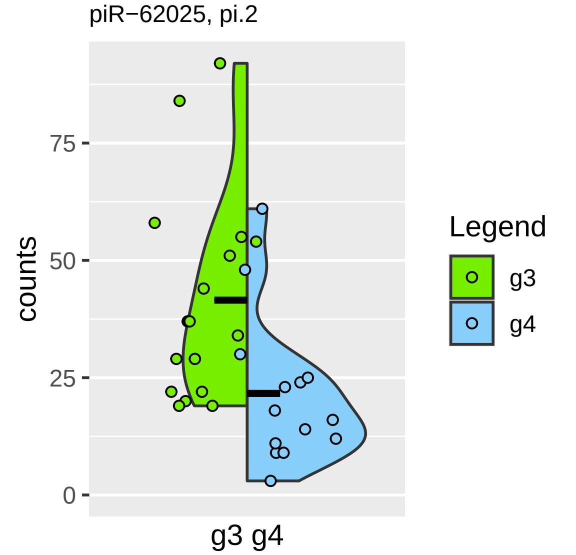
piR−62025, pi.2
counts
75
50
25
0
g3 g4
Legend
g3
g4

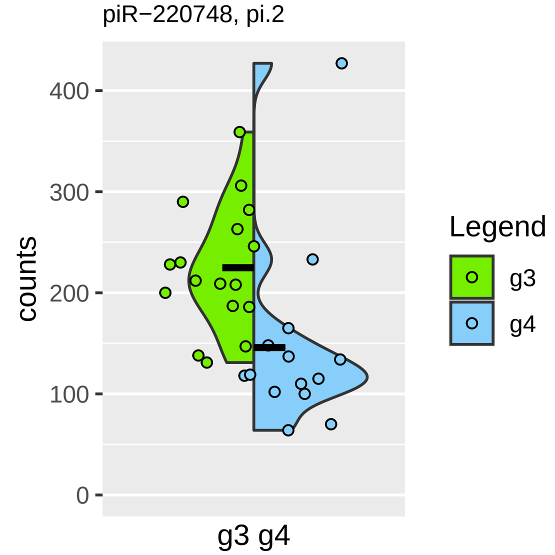
piR−220748, pi.2
counts
400
300
200
100
0
g3 g4
Legend
g3
g4

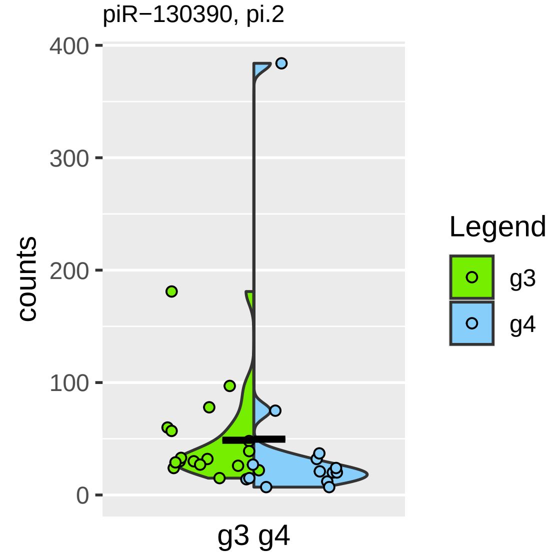

piR−130390, pi.2
counts
400
300
200
100
0
g3 g4
Legend
g3
g4

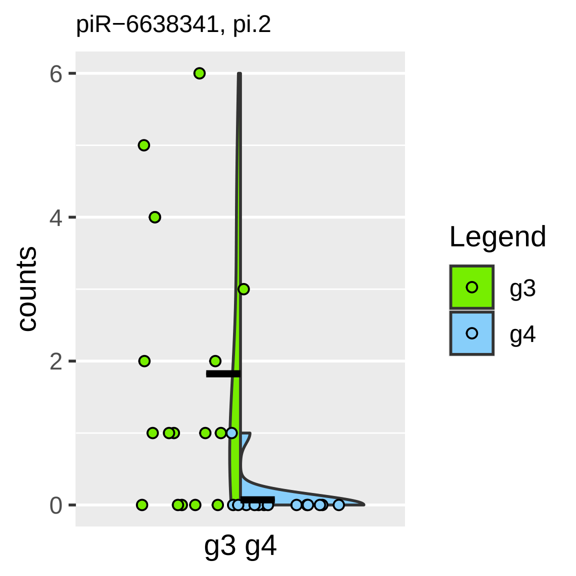
piR−6638341, pi.2
counts
6
4
2
0
g3 g4
Legend
g3
g4

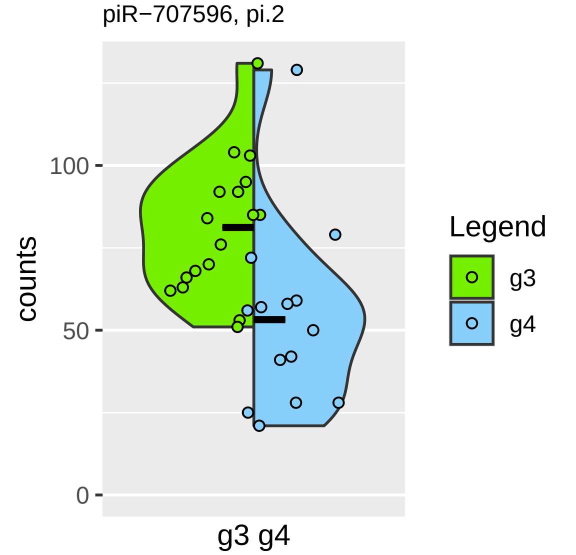

piR−707596, pi.2
counts
100
50
0
g3 g4
Legend
g3
g4

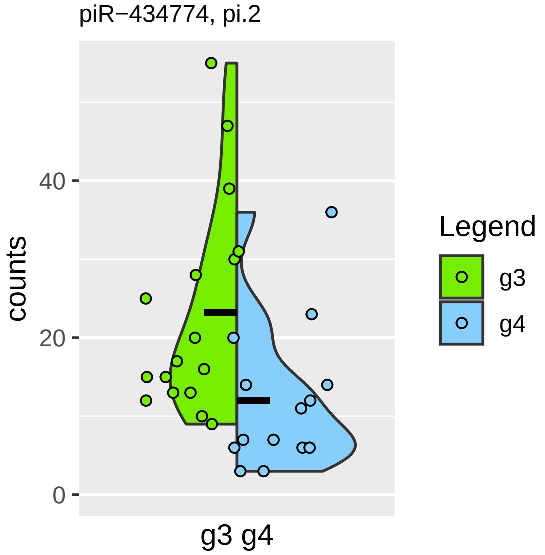

piR−434774, pi.2
counts
0
20
40
g3 g4
Legend
g3
g4

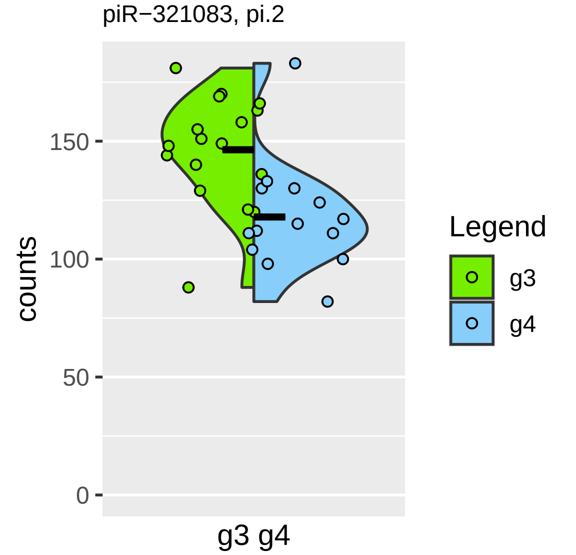

piR−321083, pi.2
counts
150
100
50
0
g3 g4
Legend
g3
g4

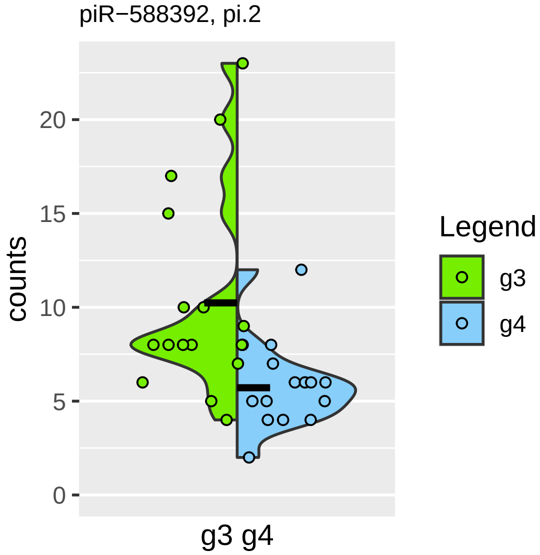
piR−588392, pi.2
counts
0
5
10
15
20
g3 g4
Legend
g3
g4

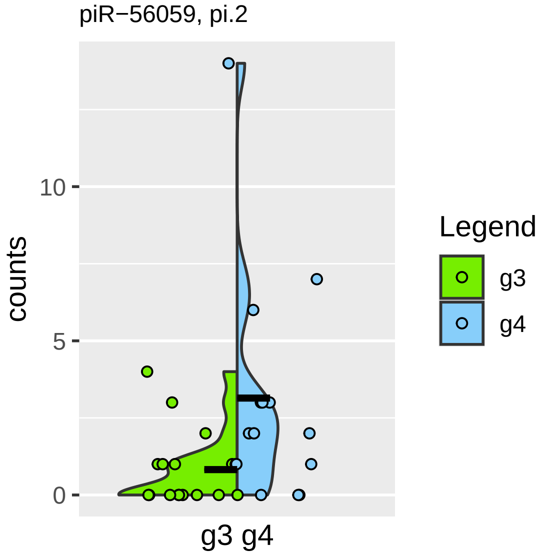

piR−56059, pi.2
counts
10
5
0
g3 g4
Legend
g3
g4

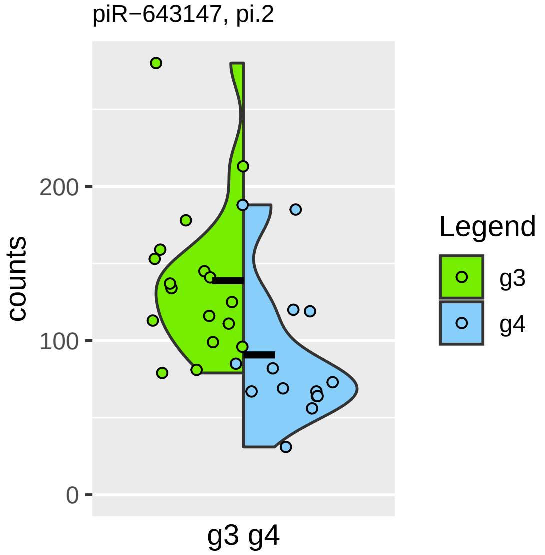
piR−643147, pi.2
counts
200
100
0
g3 g4
Legend
g3
g4

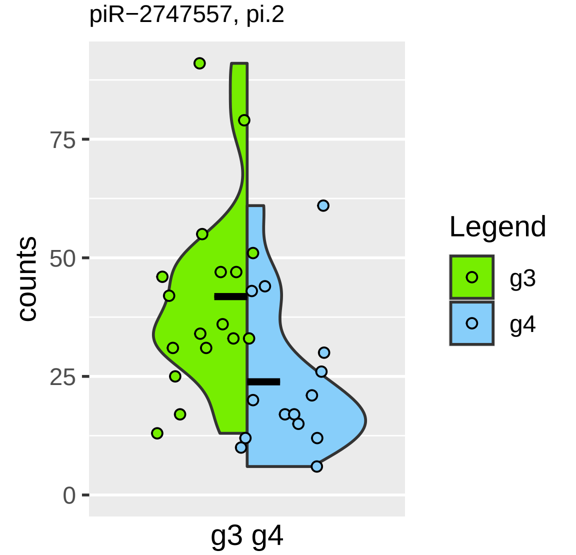

piR−2747557, pi.2
counts
75
50
25
0
g3 g4
Legend
g3
g4

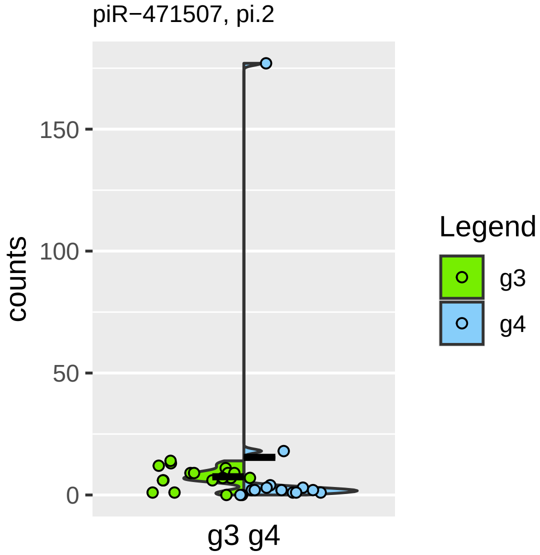
piR−471507, pi.2
counts
150
100
50
0
g3 g4
Legend
g3
g4

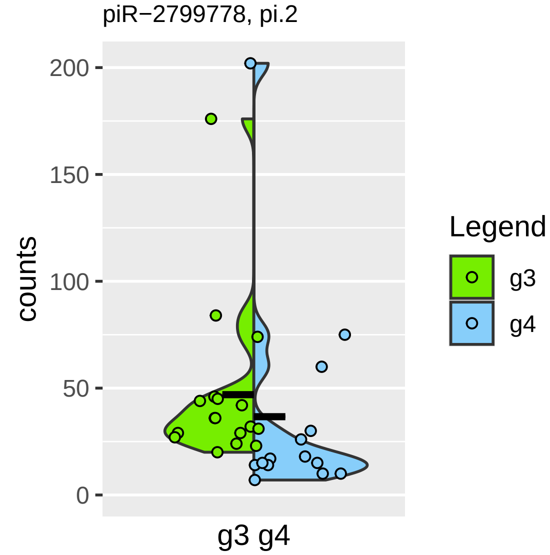

piR−2799778, pi.2
200
150
100
50
0
counts
g3 g4
Legend
g3
g4

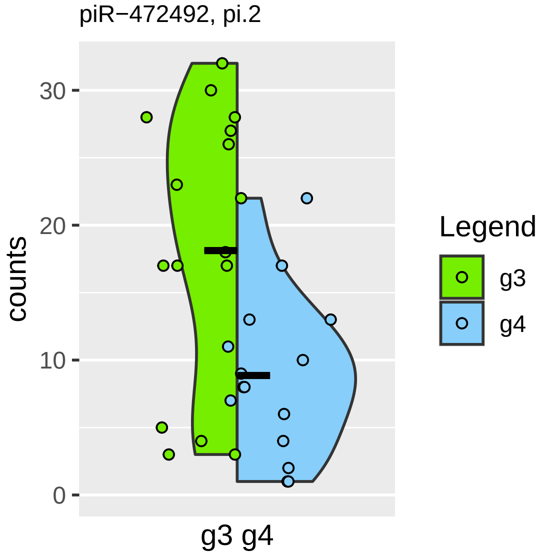
piR−472492, pi.2
counts
30
20
10
0
g3 g4
Legend
g3
g4

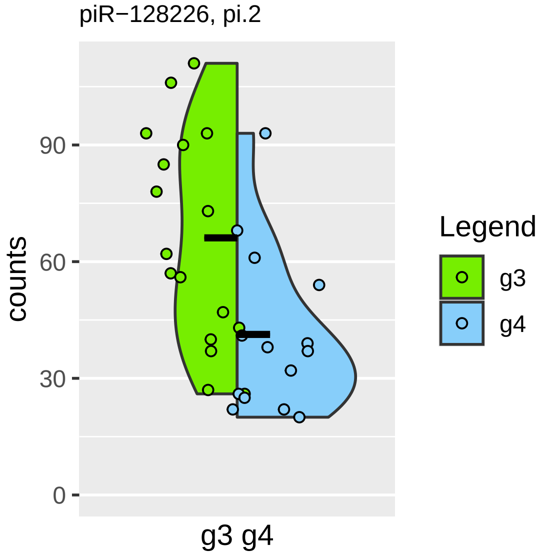

piR−128226, pi.2
counts
90
60
30
0
g3 g4
Legend
g3
g4

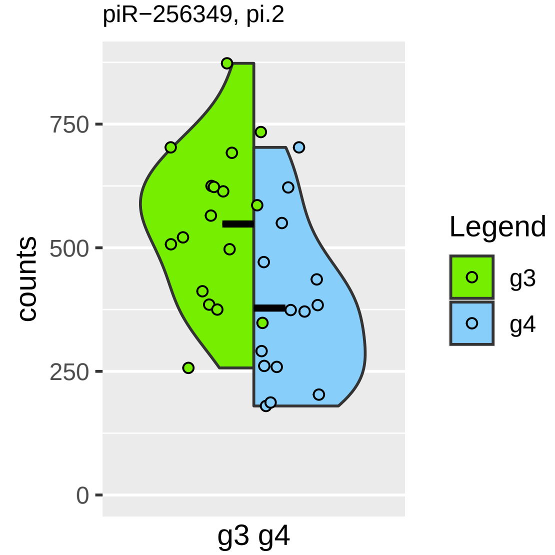

piR−256349, pi.2
counts
750
500
250
0
g3 g4
Legend
g3
g4

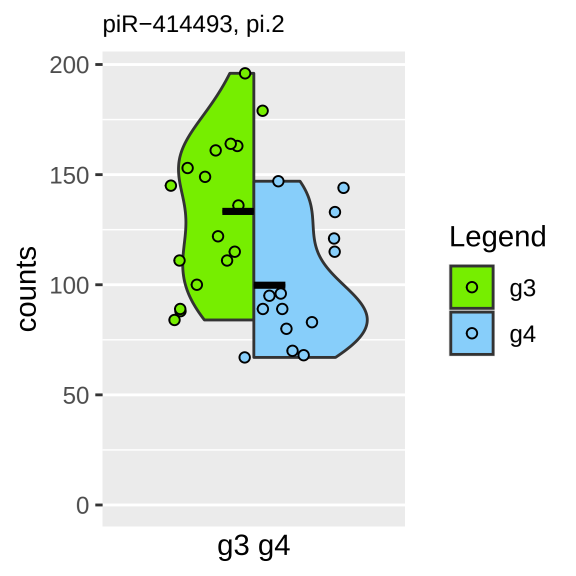
piR−414493, pi.2
200
150
100
50
0
counts
g3 g4
Legend
g3
g4

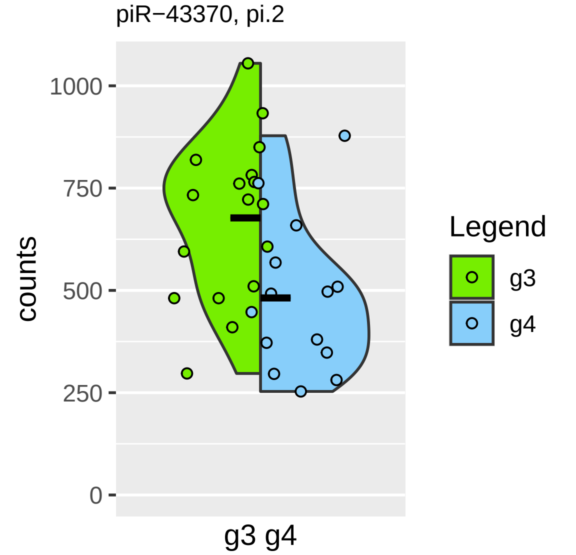

piR−43370, pi.2
counts
1000
750
500
250
0
g3 g4
Legend
g3
g4

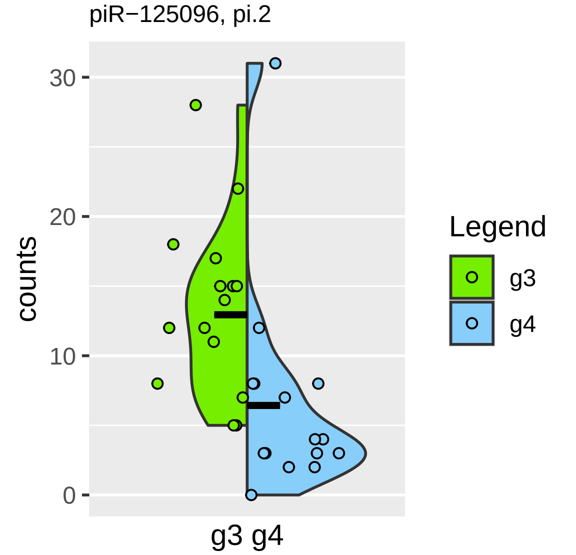
piR−125096, pi.2
counts
30
20
10
0
g3 g4
Legend
g3
g4

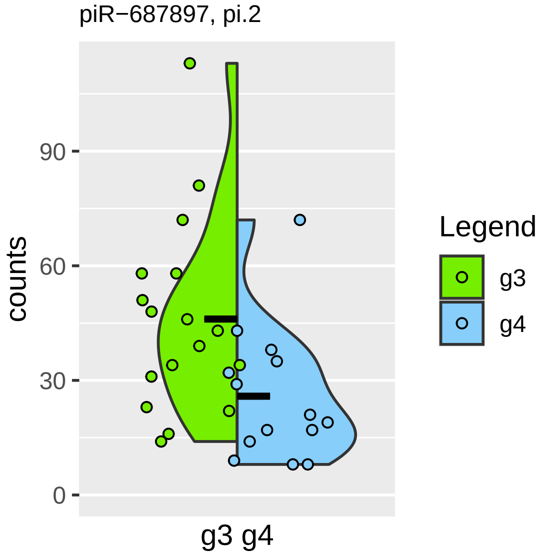

piR−687897, pi.2
counts
90
60
30
0
g3 g4
Legend
g3
g4

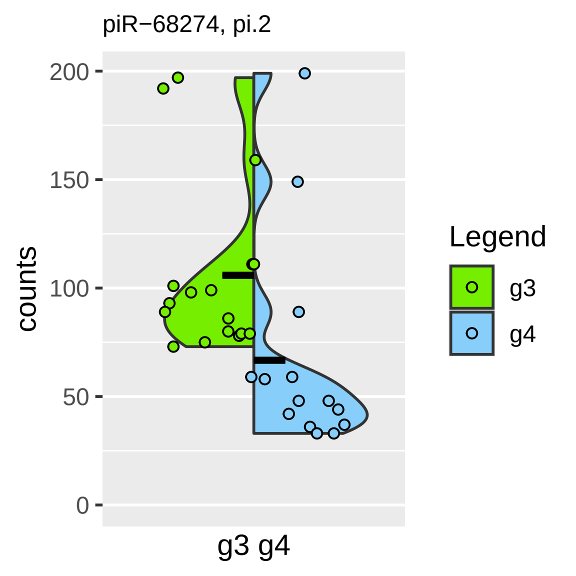
piR−68274, pi.2
counts
200
150
100
50
0
g3 g4
Legend
g3
g4

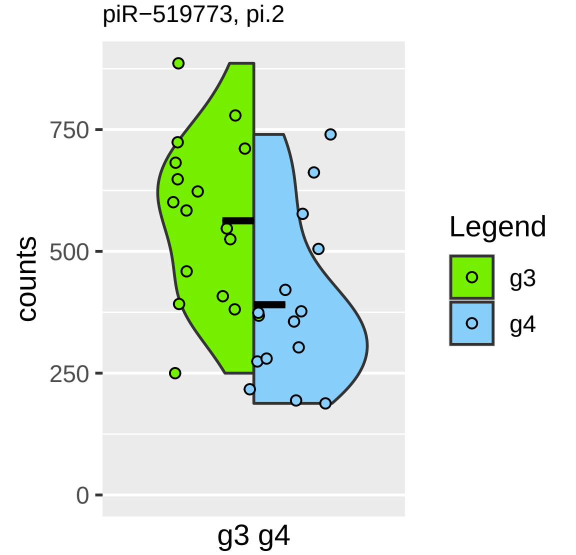
piR−519773, pi.2
counts
750
500
250
0
g3 g4
Legend
g3
g4

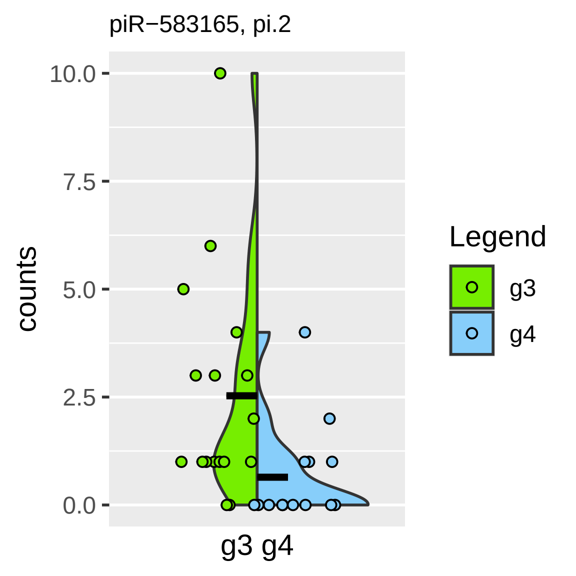
piR−583165, pi.2
10.0
7.5
5.0
2.5
0.0
counts
g3 g4
Legend
g3
g4

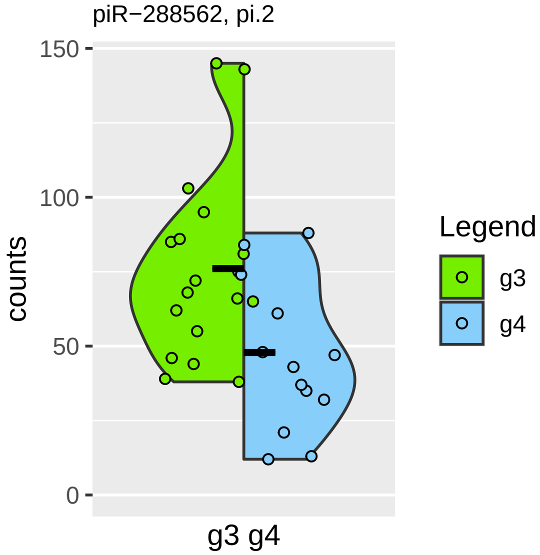

piR−288562, pi.2
150
100
50
0
counts
g3 g4
Legend
g3
g4

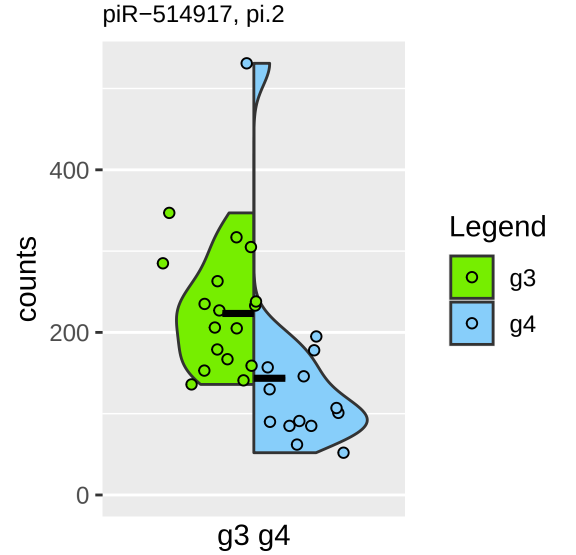

piR−514917, pi.2
counts
400
200
0
g3 g4
Legend
g3
g4

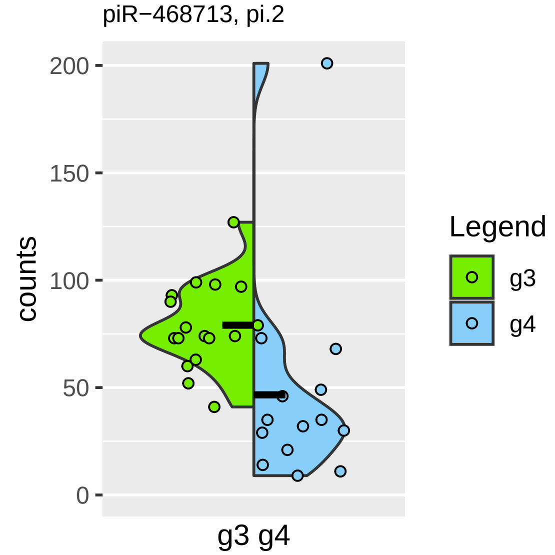

piR−468713, pi.2
200
150
100
50
0
counts
g3 g4
Legend
g3
g4

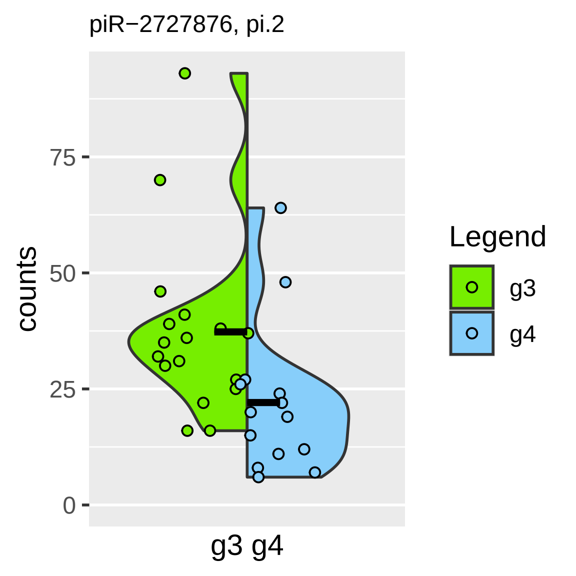
piR−2727876, pi.2
counts
75
50
25
0
g3 g4
Legend
g3
g4

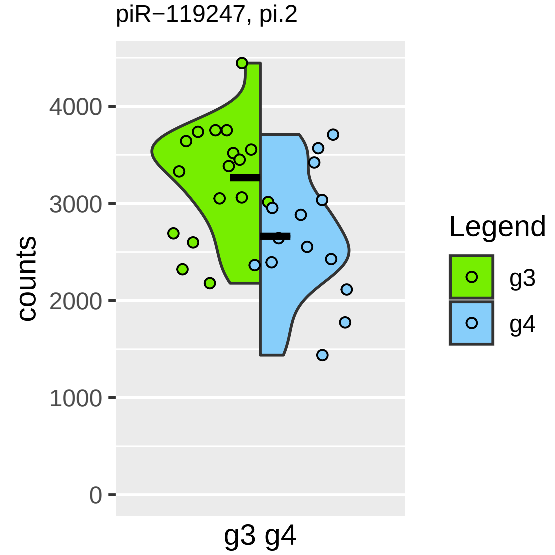

piR−119247, pi.2
4000
3000
2000
1000
0
counts
g3 g4
Legend
g3
g4

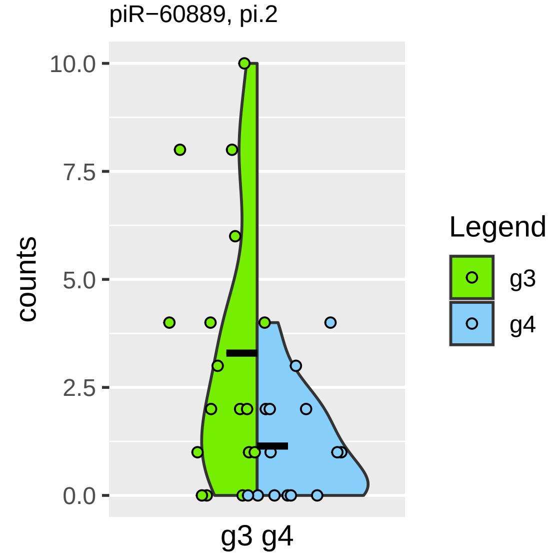

piR−60889, pi.2
10.0
7.5
5.0
2.5
0.0
counts
g3 g4
Legend
g3
g4

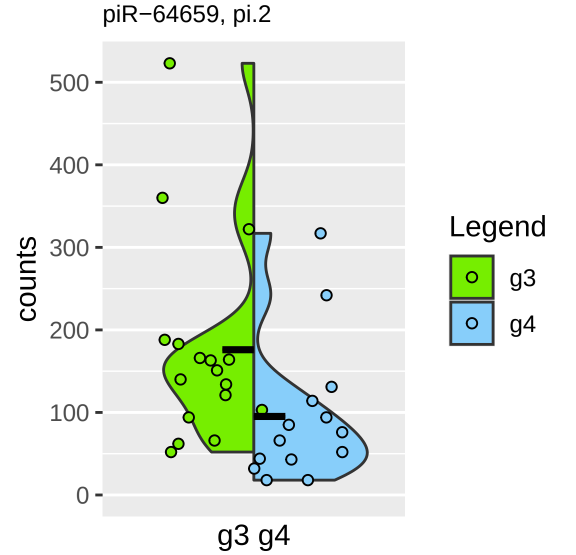
piR−64659, pi.2
500
400
300
200
100
0
counts
g3 g4
Legend
g3
g4

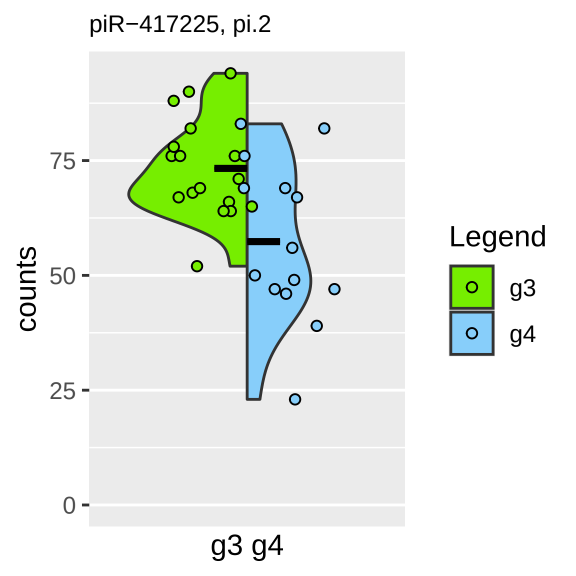

piR−417225, pi.2
counts
75
50
25
0
g3 g4
Legend
g3
g4

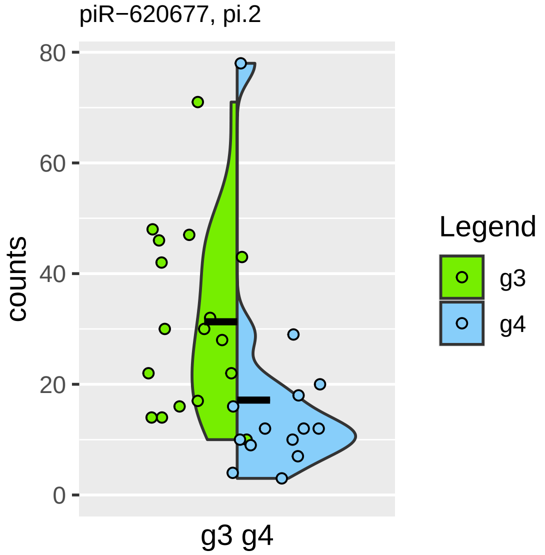

piR−620677, pi.2
counts
80
60
40
20
0
g3 g4
Legend
g3
g4

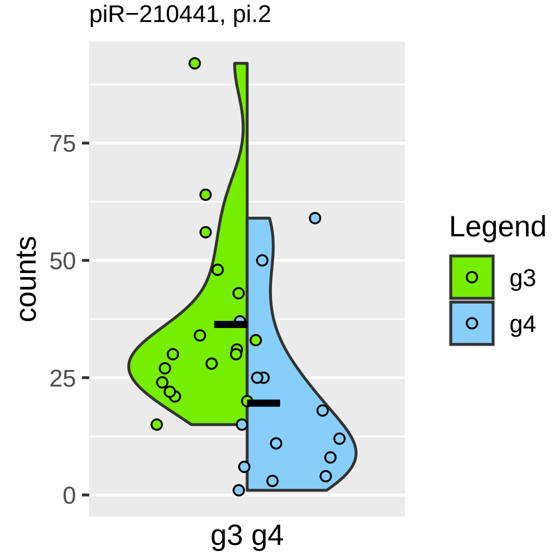

piR−210441, pi.2
counts
75
50
25
0
g3 g4
Legend
g3
g4

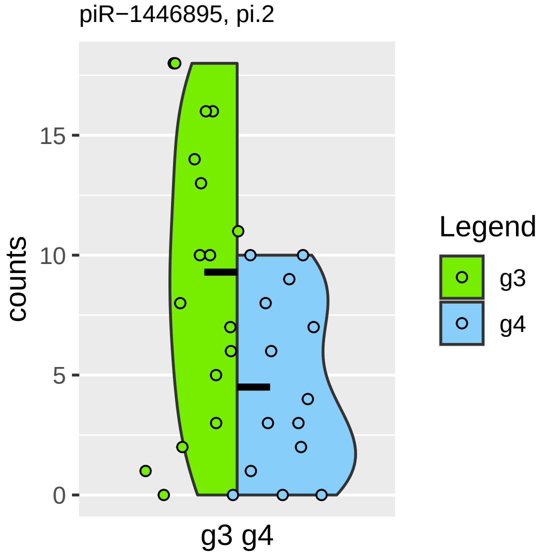

piR−1446895, pi.2
counts
15
10
5
0
g3 g4
Legend
g3
g4

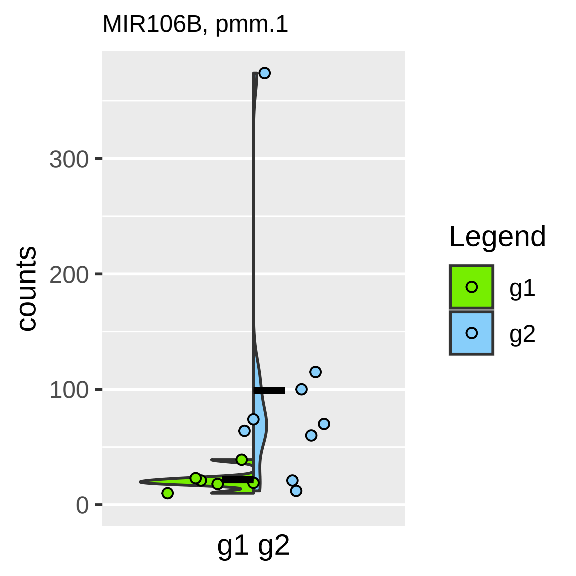

MIR106B, pmm.1
counts
300
200
100
0
g1 g2
Legend
g1
g2

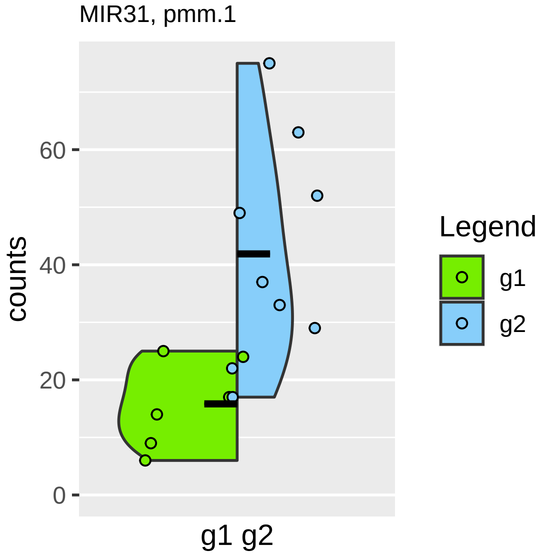

MIR31, pmm.1
counts
60
40
20
0
g1 g2
Legend
g1
g2

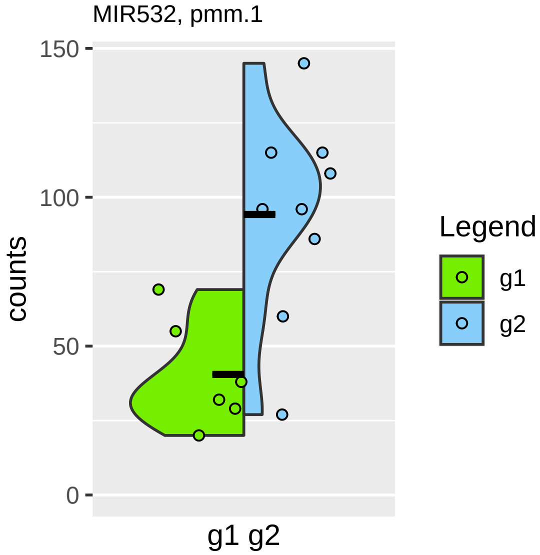

MIR532, pmm.1
150
100
50
0
counts
g1 g2
Legend
g1
g2

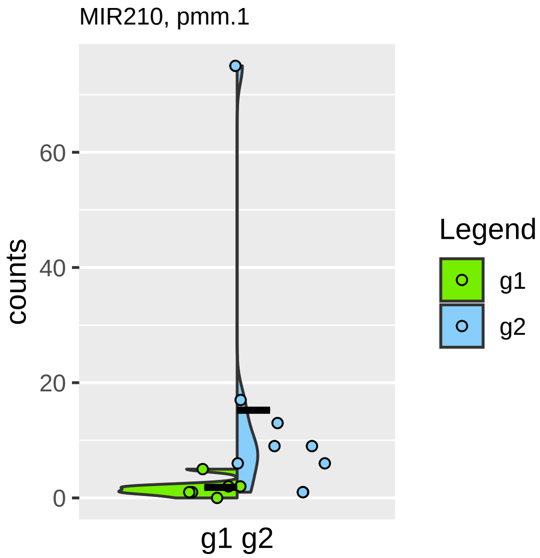
MIR210, pmm.1
counts
0
20
40
60
g1 g2
Legend
g1
g2

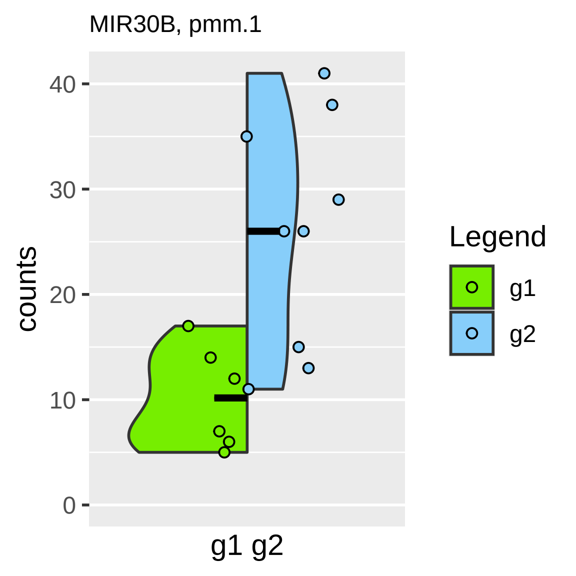

MIR30B, pmm.1
counts
0
10
20
30
40
g1 g2
Legend
g1
g2

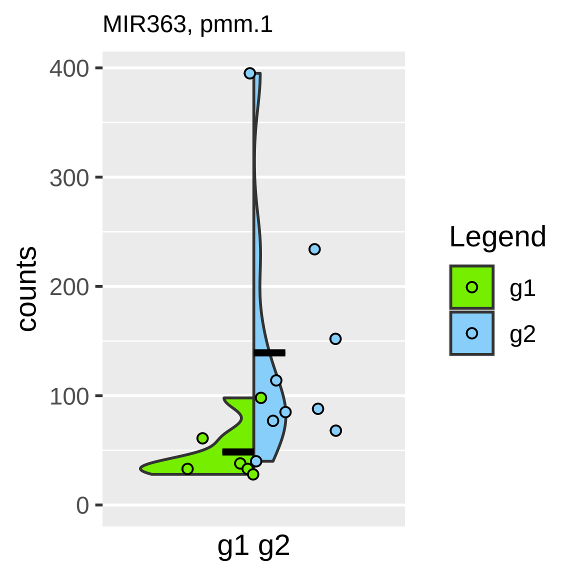

MIR363, pmm.1
counts
400
300
200
100
0
g1 g2
Legend
g1
g2

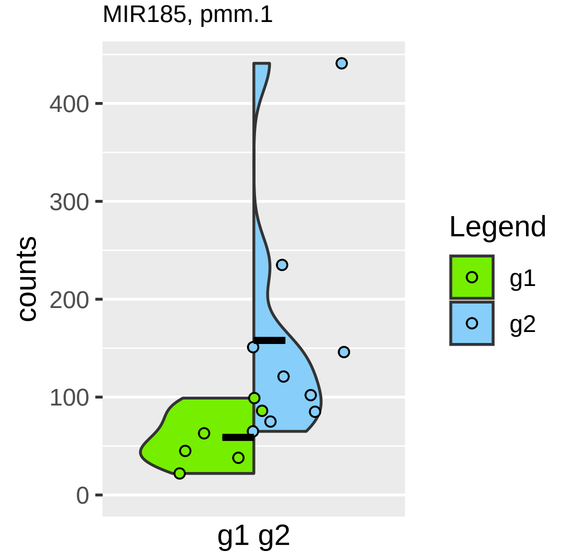

MIR185, pmm.1
counts
400
300
200
100
0
g1 g2
Legend
g1
g2

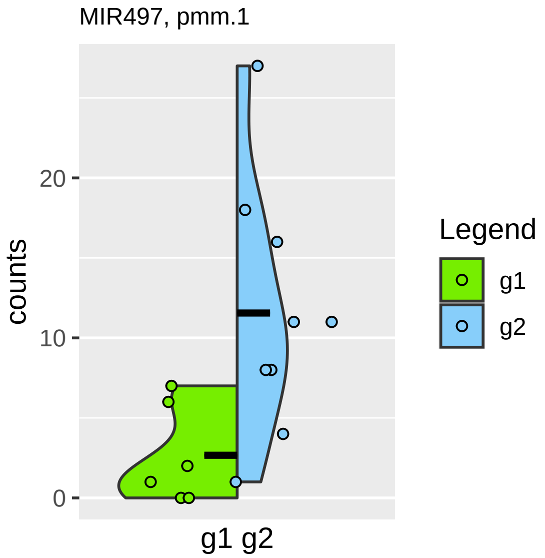

MIR497, pmm.1
counts
0
10
20
g1 g2
Legend
g1
g2

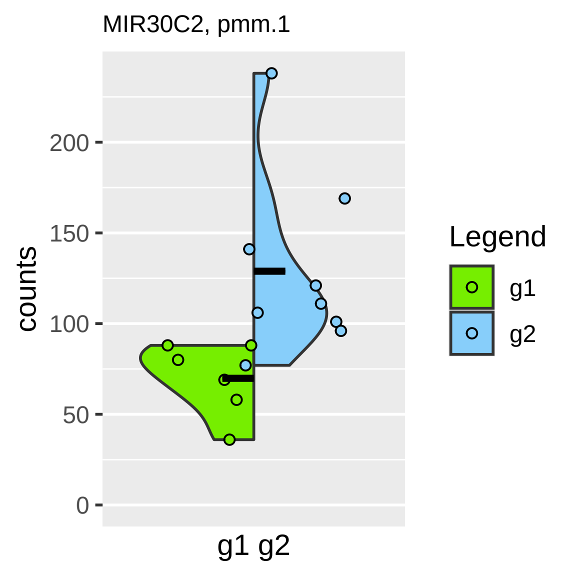

MIR30C2, pmm.1
counts
200
150
100
50
0
g1 g2
Legend
g1
g2

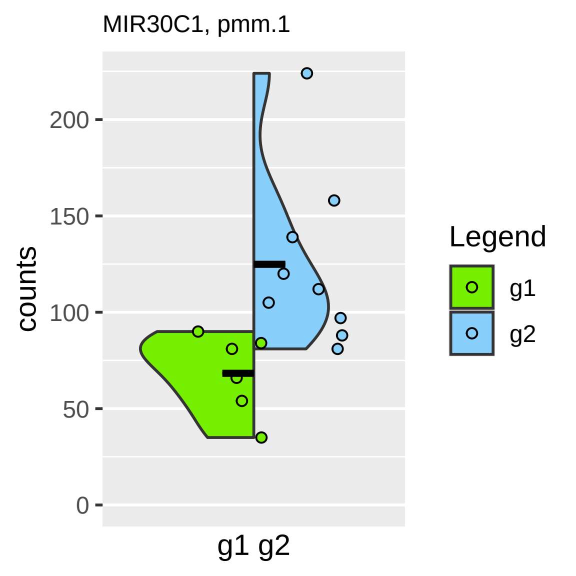
MIR30C1, pmm.1
counts
200
150
100
50
0
g1 g2
Legend
g1
g2

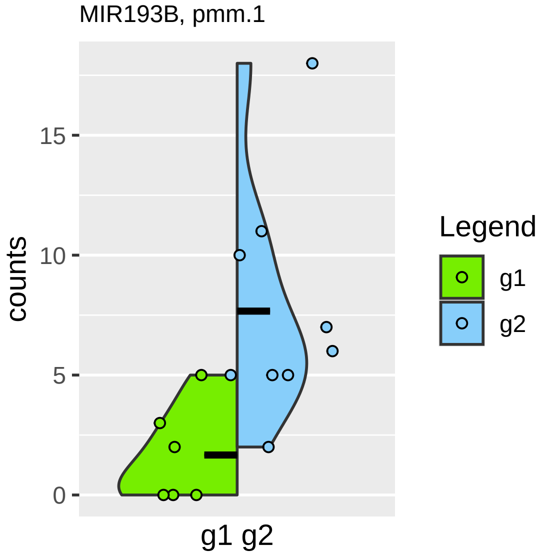
MIR193B, pmm.1
counts
0
5
10
15
g1 g2
Legend
g1
g2

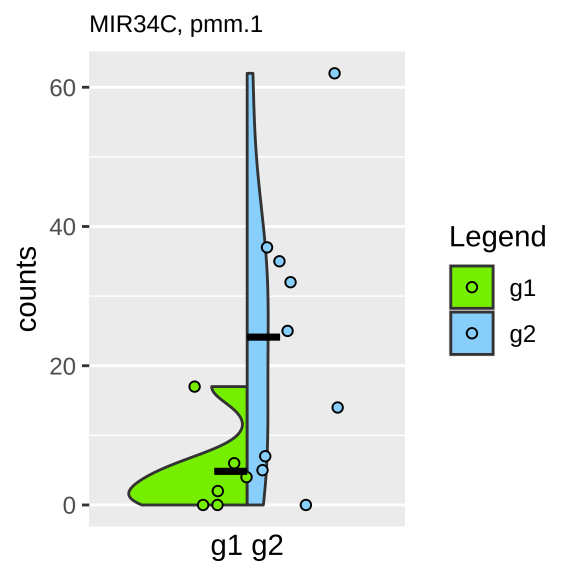

MIR34C, pmm.1
counts
60
40
20
0
g1 g2
Legend
g1
g2

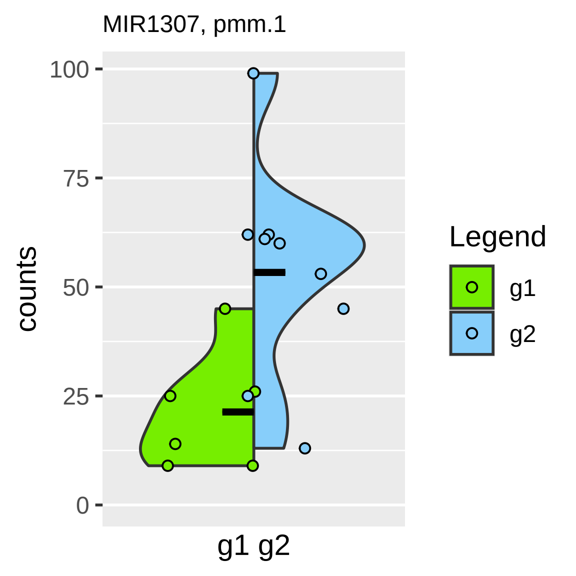
MIR1307, pmm.1
counts
100
75
50
25
0
g1 g2
Legend
g1
g2

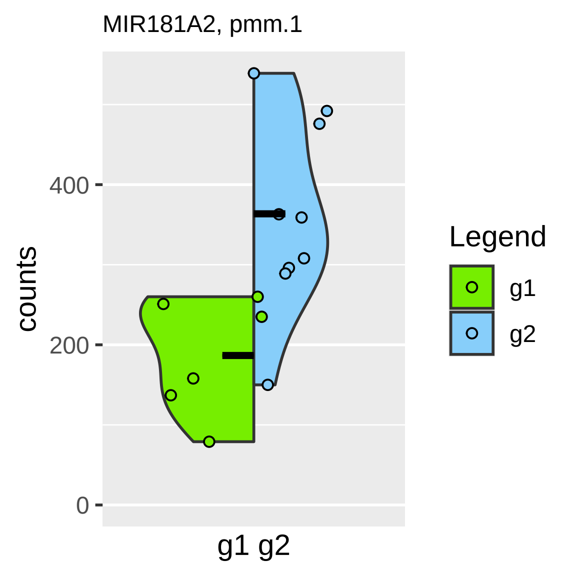
MIR181A2, pmm.1
counts
400
200
0
g1 g2
Legend
g1
g2

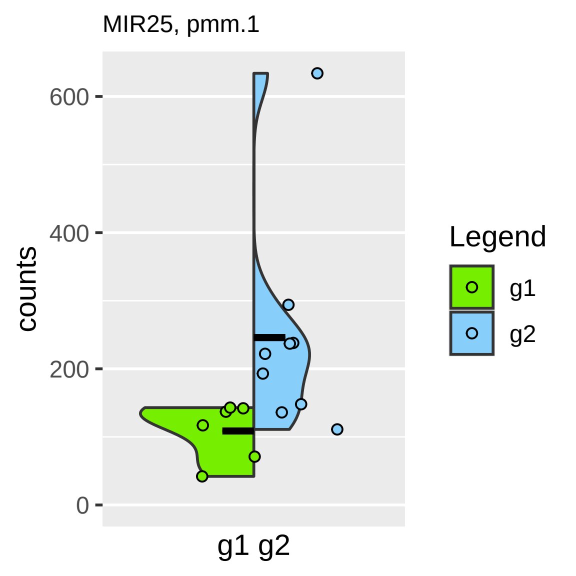

MIR25, pmm.1
counts
0
200
400
600
g1 g2
Legend
g1
g2

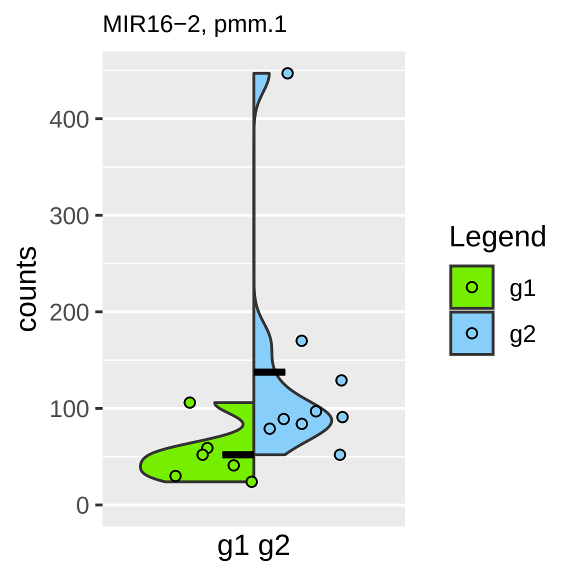

MIR16−2, pmm.1
counts
400
300
200
100
0
g1 g2
Legend
g1
g2

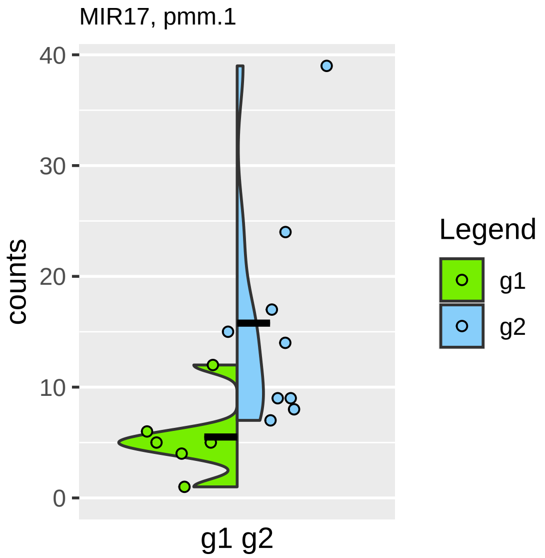

MIR17, pmm.1
counts
40
30
20
10
0
g1 g2
Legend
g1
g2

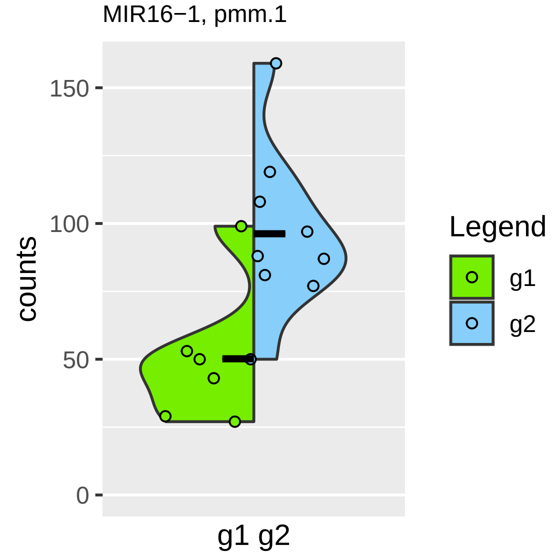

MIR16−1, pmm.1
counts
150
100
50
0
g1 g2
Legend
g1
g2

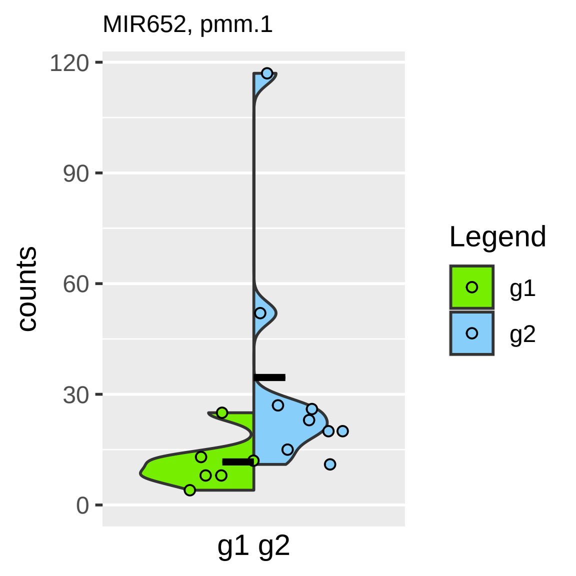
MIR652, pmm.1
120
90
60
30
0
counts
g1 g2
Legend
g1
g2

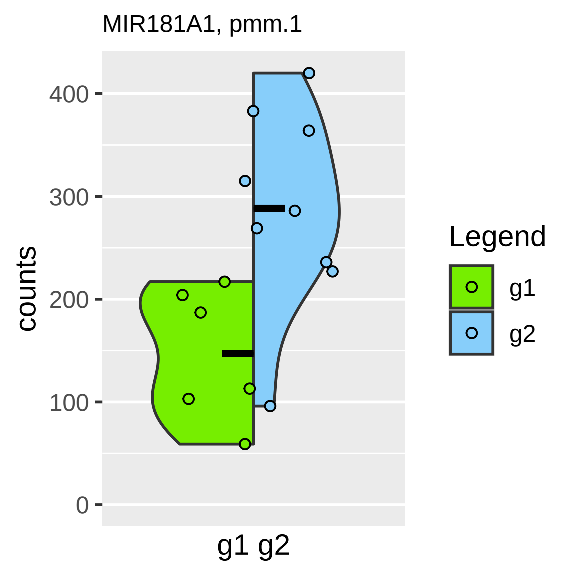

MIR181A1, pmm.1
counts
400
300
200
100
0
g1 g2
Legend
g1
g2

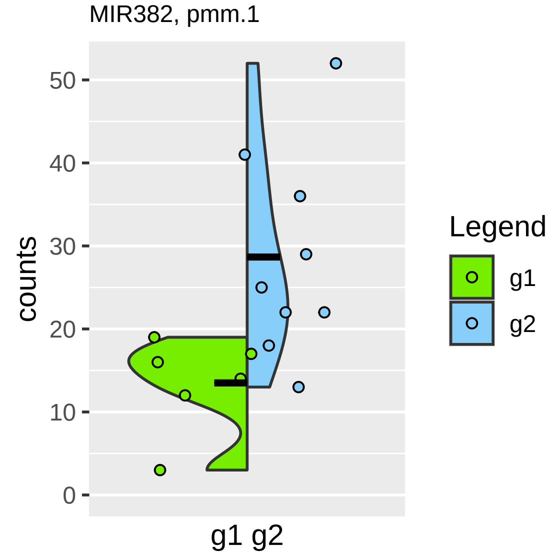
MIR382, pmm.1
50
40
30
20
10
0
counts
g1 g2
Legend
g1
g2

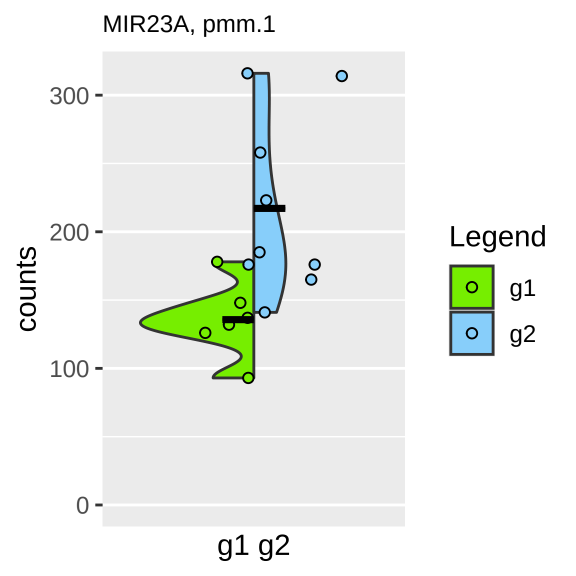
MIR23A, pmm.1
counts
300
200
100
0
g1 g2
Legend
g1
g2

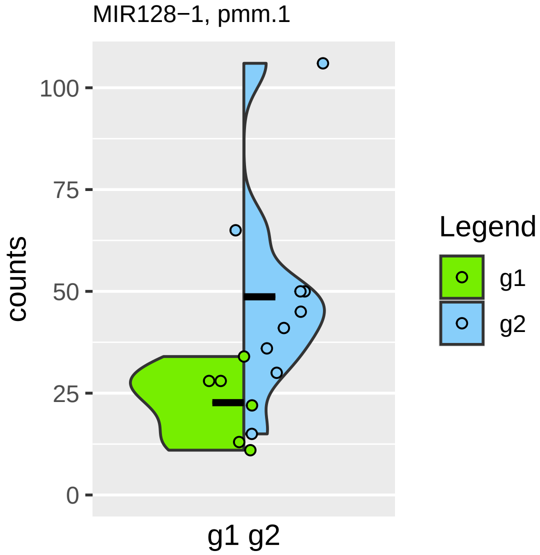

MIR128−1, pmm.1
counts
100
75
50
25
0
g1 g2
Legend
g1
g2

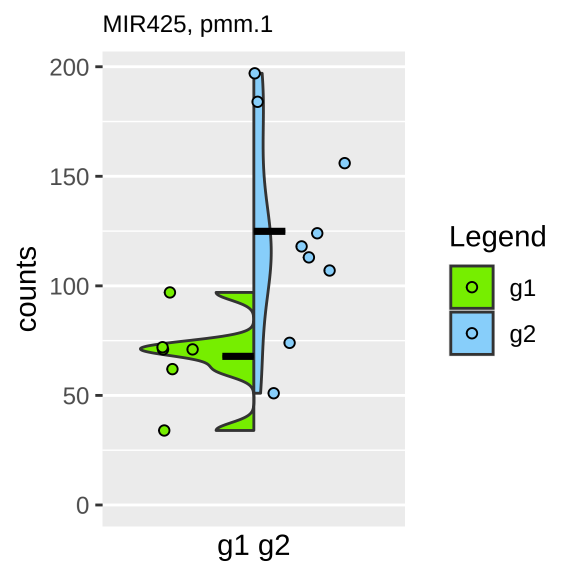

MIR425, pmm.1
200
150
100
50
0
counts
g1 g2
Legend
g1
g2

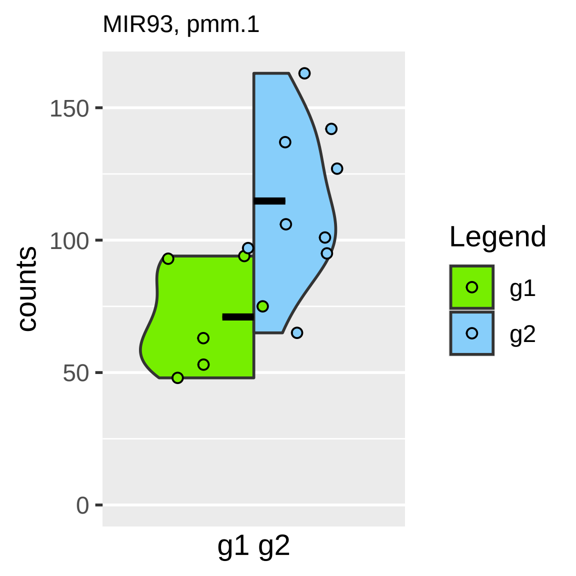

MIR93, pmm.1
150
100
50
0
counts
g1 g2
Legend
g1
g2

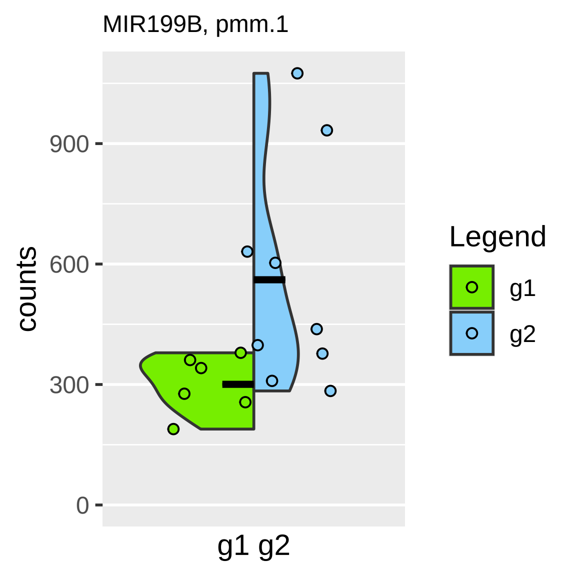
MIR199B, pmm.1
counts
900
600
300
0
g1 g2
Legend
g1
g2

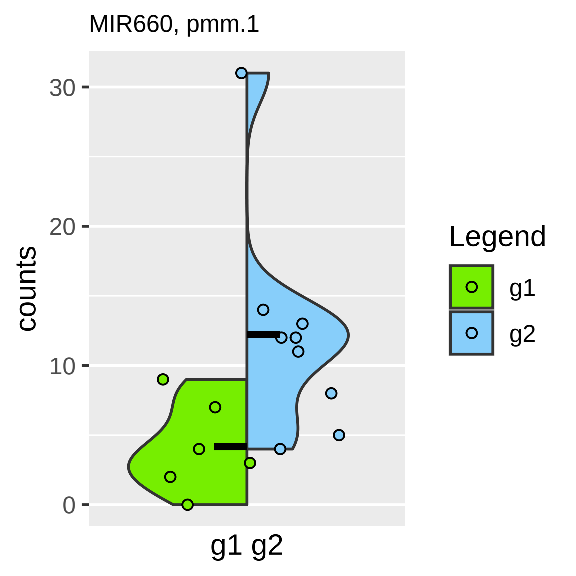
MIR660, pmm.1
30
20
10
0
counts
g1 g2
Legend
g1
g2

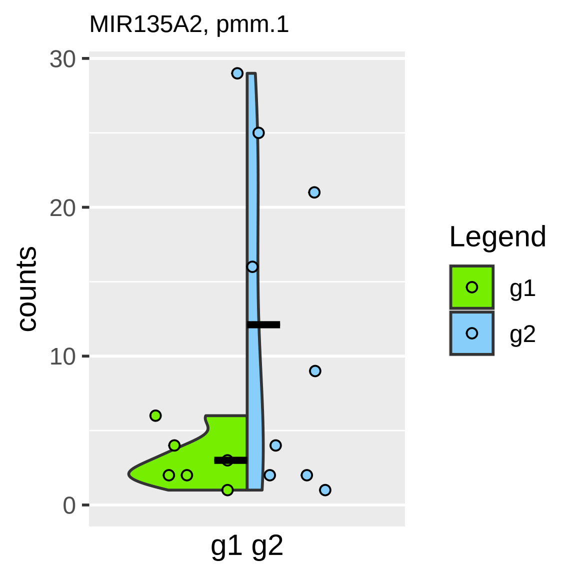

MIR135A2, pmm.1
counts
30
20
10
0
g1 g2
Legend
g1
g2

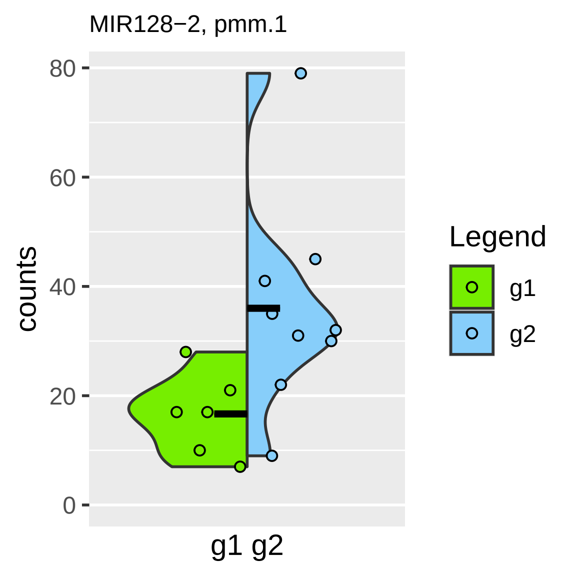
MIR128−2, pmm.1
counts
80
60
40
20
0
g1 g2
Legend
g1
g2

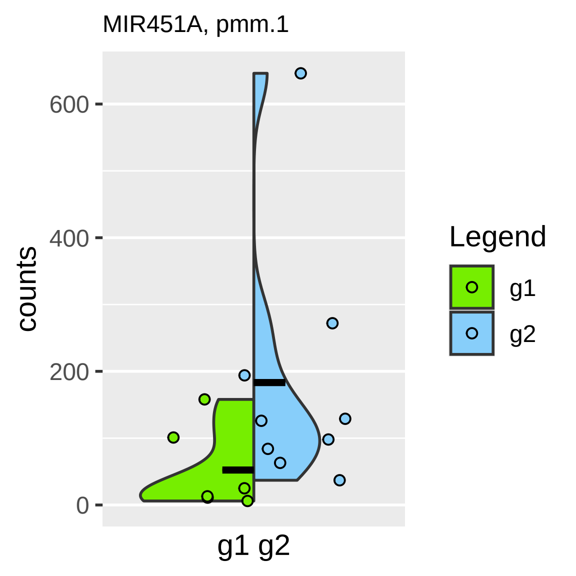

MIR451A, pmm.1
counts
600
400
200
0
g1 g2
Legend
g1
g2

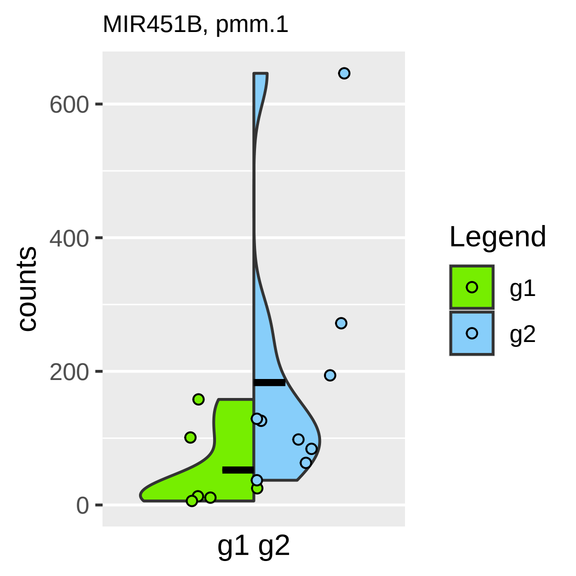
MIR451B, pmm.1
counts
600
400
200
0
g1 g2
Legend
g1
g2

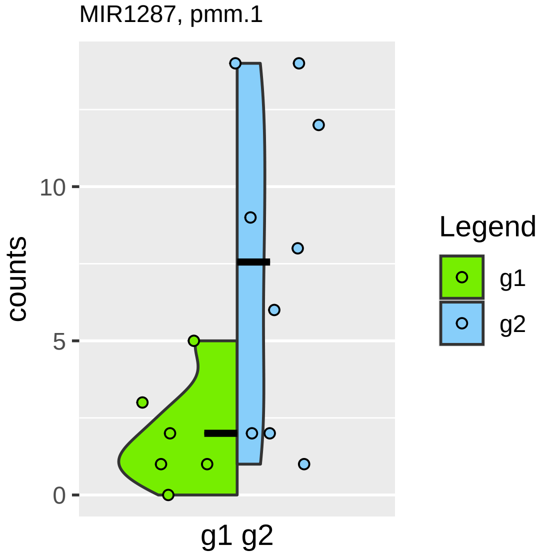
MIR1287, pmm.1
counts
10
5
0
g1 g2
Legend
g1
g2

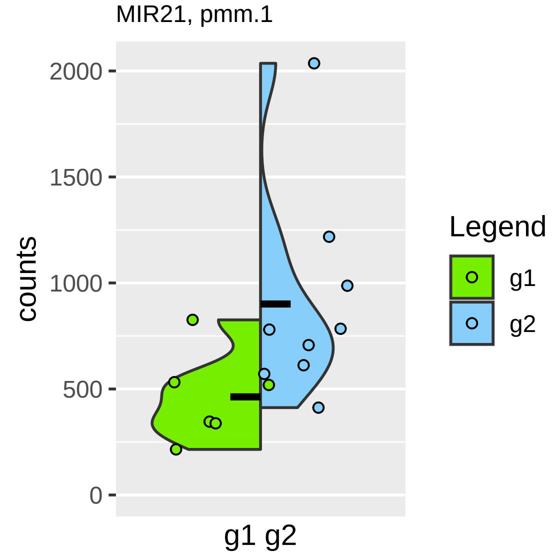

MIR21, pmm.1
counts
2000
1500
1000
500
0
g1 g2
Legend
g1
g2

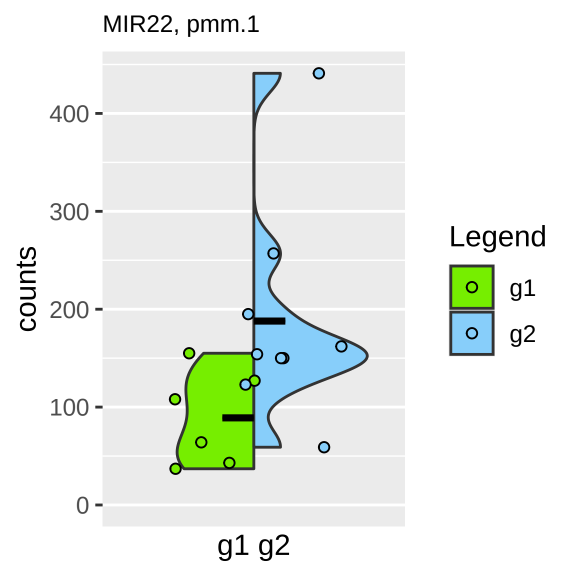

MIR22, pmm.1
counts
400
300
200
100
0
g1 g2
Legend
g1
g2

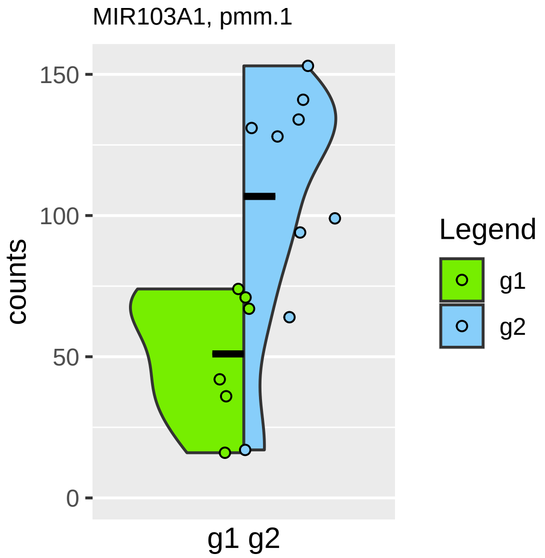

MIR103A1, pmm.1
150
100
50
0
counts
g1 g2
Legend
g1
g2

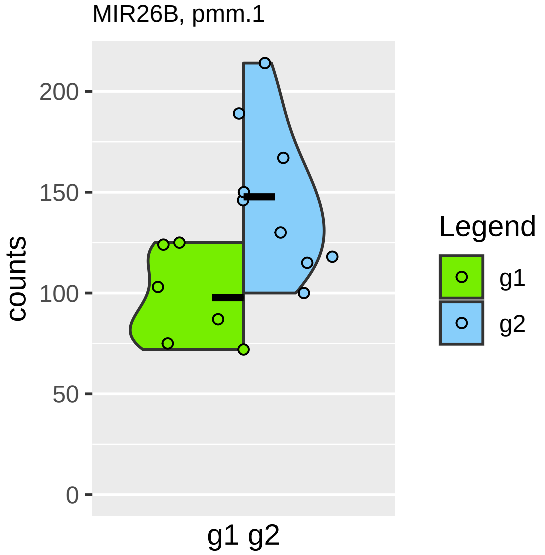

MIR26B, pmm.1
counts
0
50
100
150
200
g1 g2
Legend
g1
g2

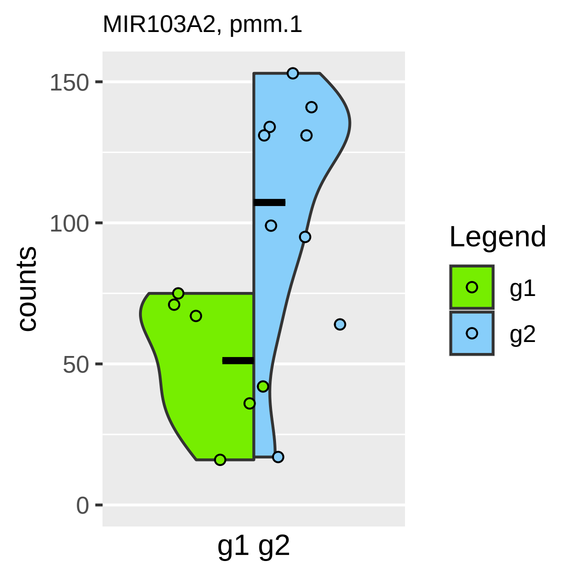

MIR103A2, pmm.1
150
100
50
0
counts
g1 g2
Legend
g1
g2

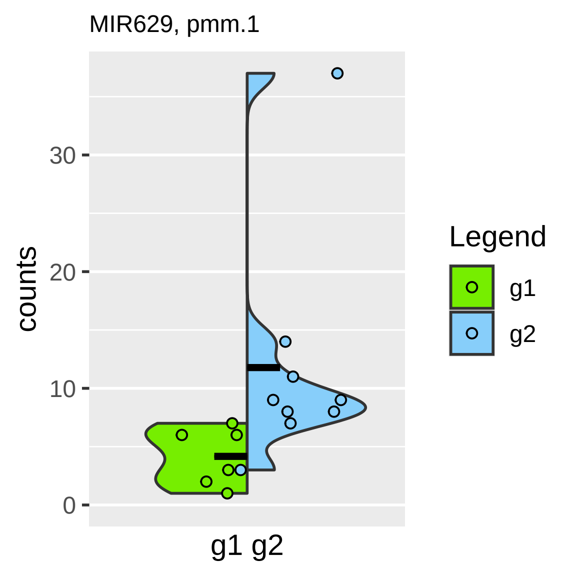
MIR629, pmm.1
counts
30
20
10
0
g1 g2
Legend
g1
g2

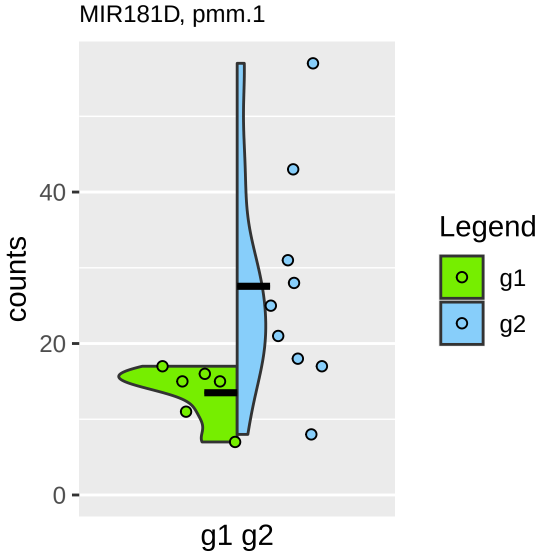

MIR181D, pmm.1
counts
40
20
0
g1 g2
Legend
g1
g2

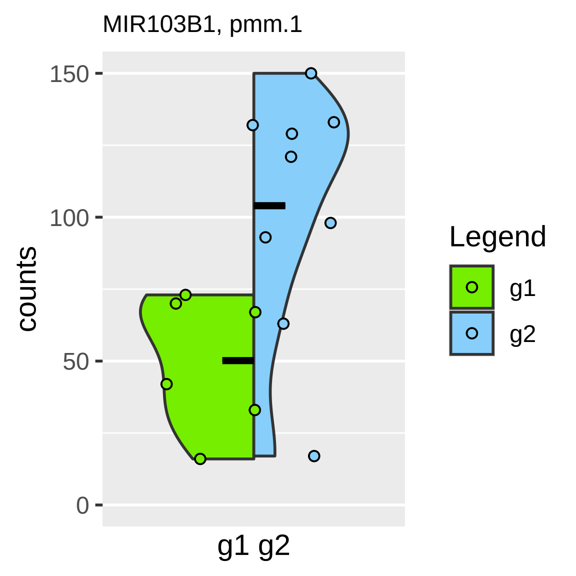

MIR103B1, pmm.1
counts
150
100
50
0
g1 g2
Legend
g1
g2

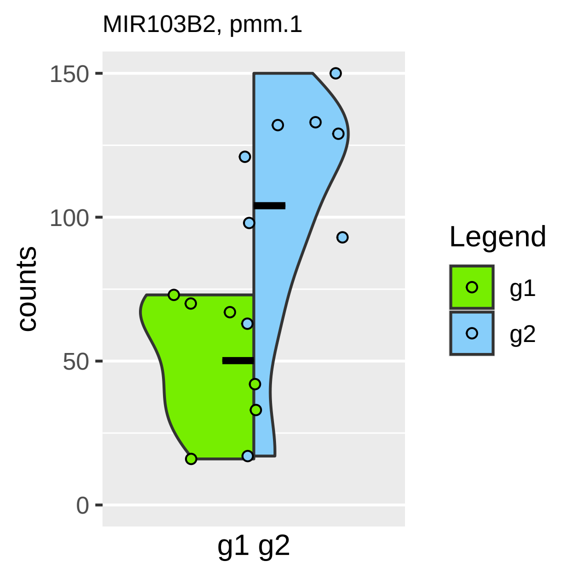
MIR103B2, pmm.1
150
100
50
0
counts
g1 g2
Legend
g1
g2

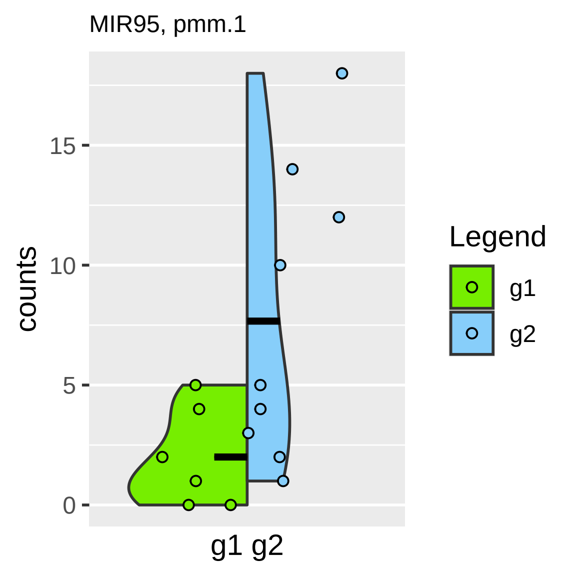

MIR95, pmm.1
counts
15
10
5
0
g1 g2
Legend
g1
g2

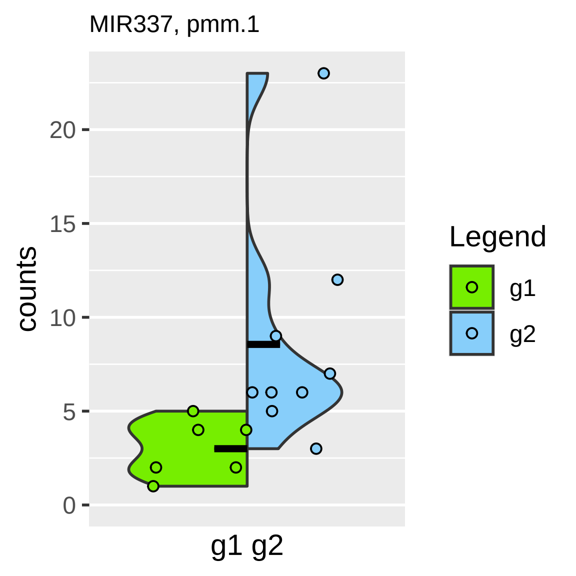
MIR337, pmm.1
counts
0
5
10
15
20
g1 g2
Legend
g1
g2

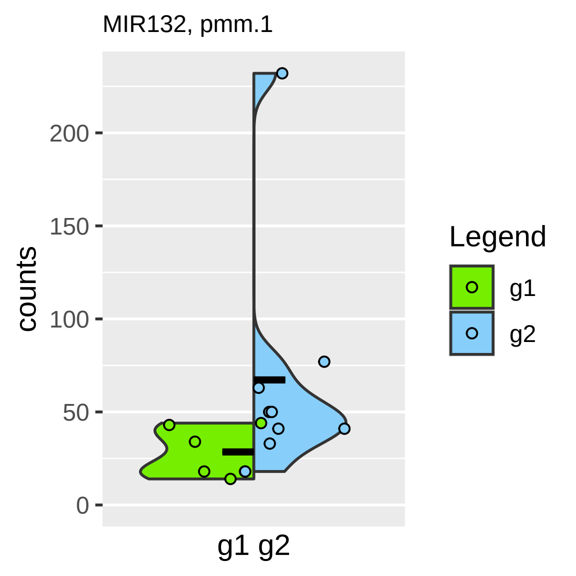

MIR132, pmm.1
counts
200
150
100
50
0
g1 g2
Legend
g1
g2

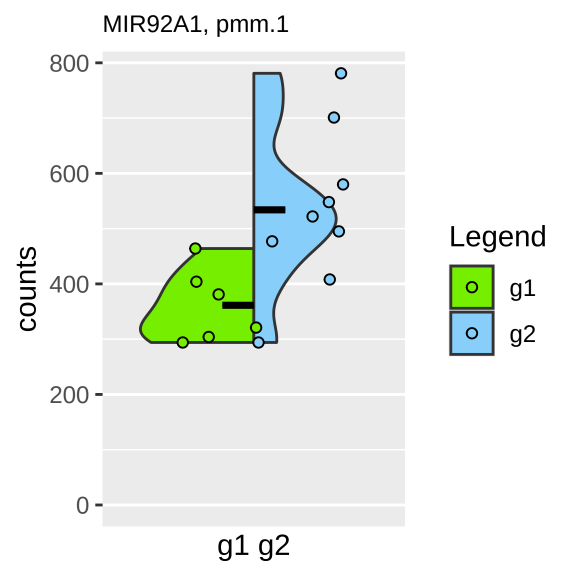

MIR92A1, pmm.1
counts
800
600
400
200
0
g1 g2
Legend
g1
g2

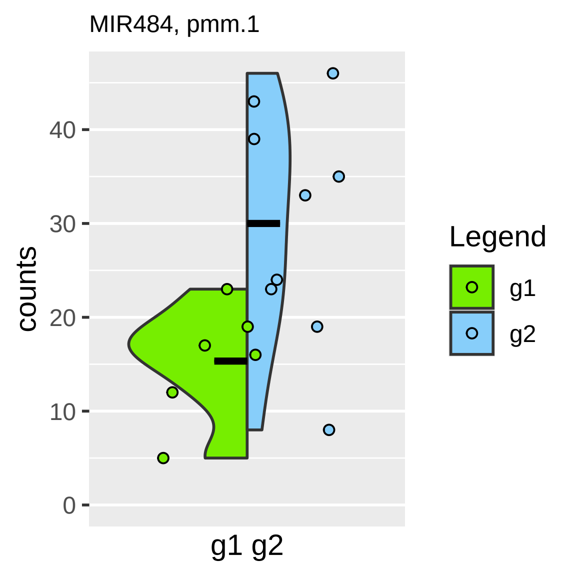

MIR484, pmm.1
counts
0
10
20
30
40
g1 g2
Legend
g1
g2

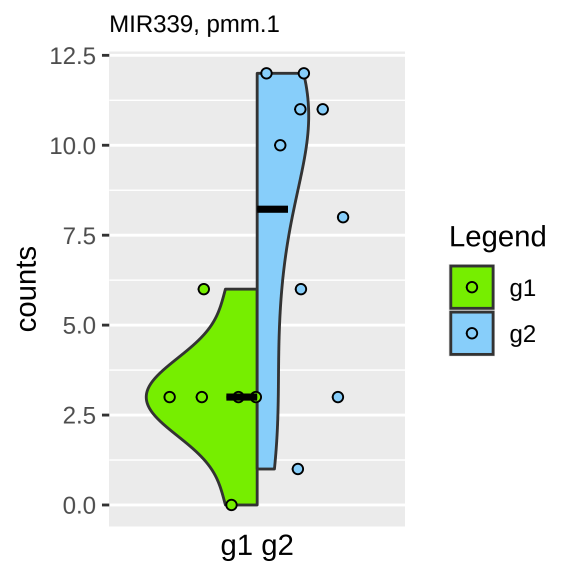

MIR339, pmm.1
12.5
10.0
7.5
5.0
2.5
0.0
counts
g1 g2
Legend
g1
g2

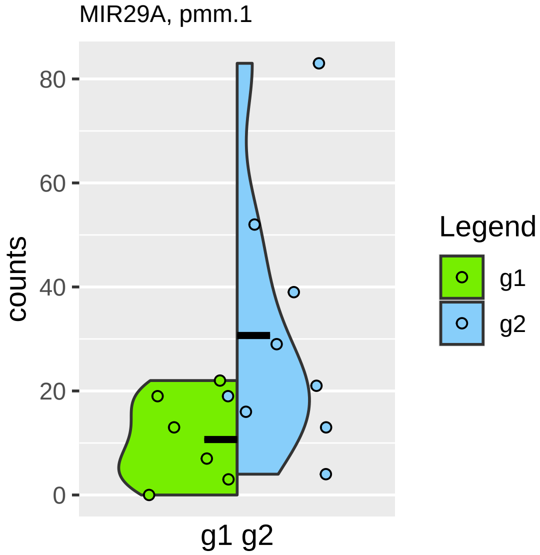

MIR29A, pmm.1
counts
80
60
40
20
0
g1 g2
Legend
g1
g2

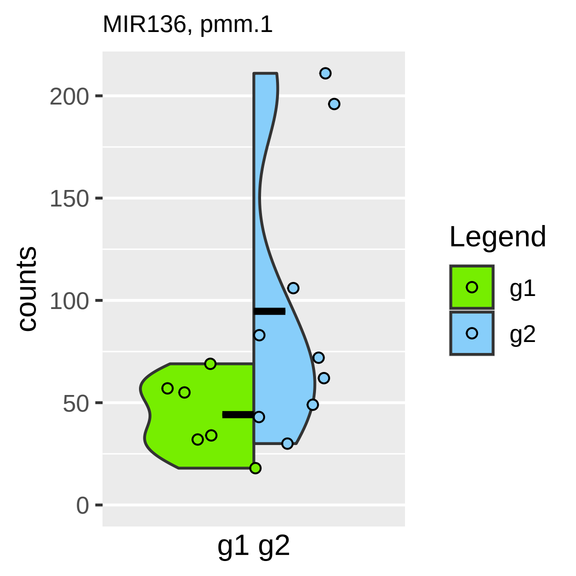
MIR136, pmm.1
counts
200
150
100
50
0
g1 g2
Legend
g1
g2

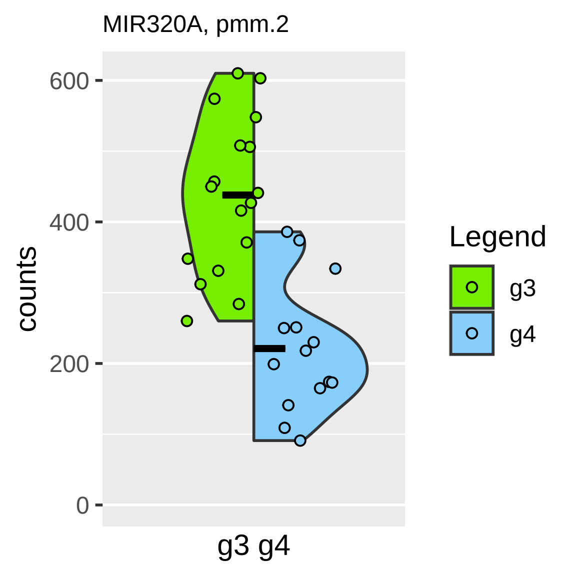
MIR320A, pmm.2
counts
600
400
200
0
g3 g4
Legend
g3
g4

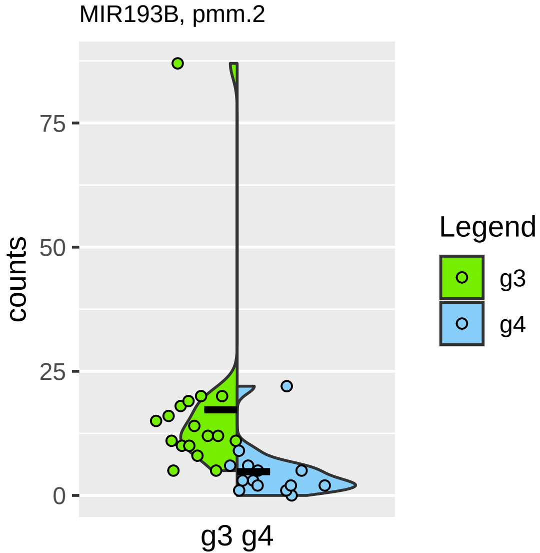

MIR193B, pmm.2
counts
75
50
25
0
g3 g4
Legend
g3
g4

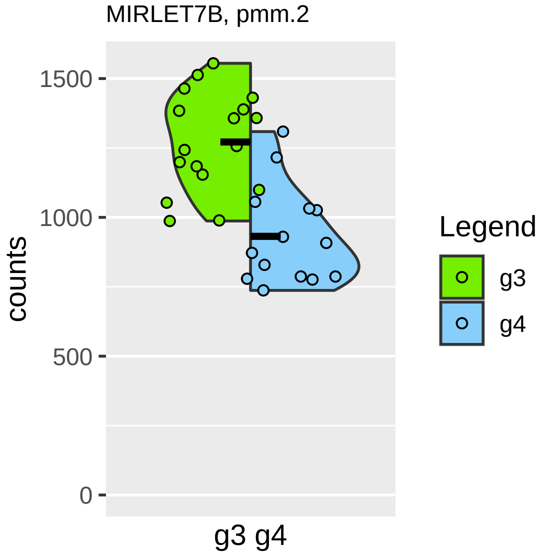
MIRLET7B, pmm.2
1500
1000
500
0
counts
g3 g4
Legend
g3
g4

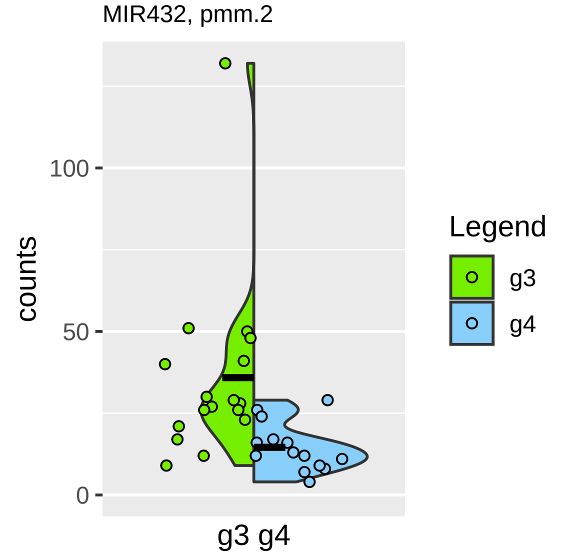

MIR432, pmm.2
counts
100
50
0
g3 g4
Legend
g3
g4

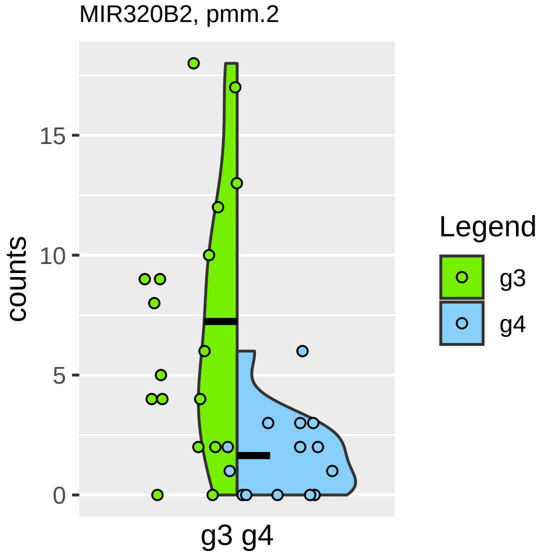

MIR320B2, pmm.2
counts
15
10
5
0
g3 g4
Legend
g3
g4

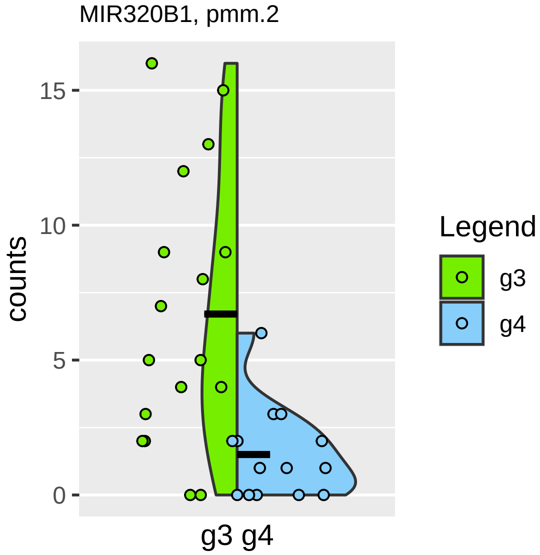

MIR320B1, pmm.2
counts
15
10
5
0
g3 g4
Legend
g3
g4

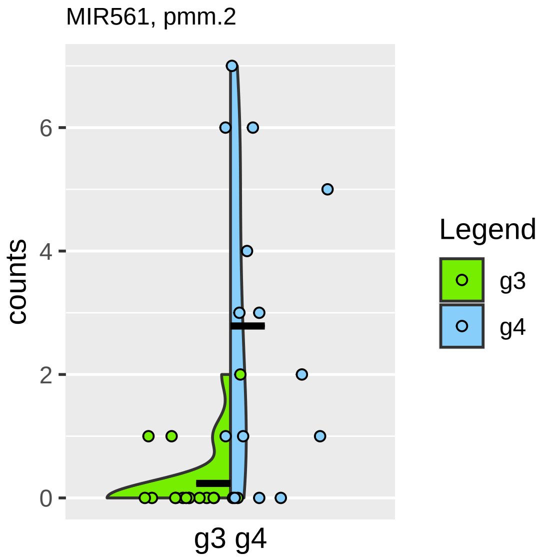
MIR561, pmm.2
counts
0
2
4
6
g3 g4
Legend
g3
g4

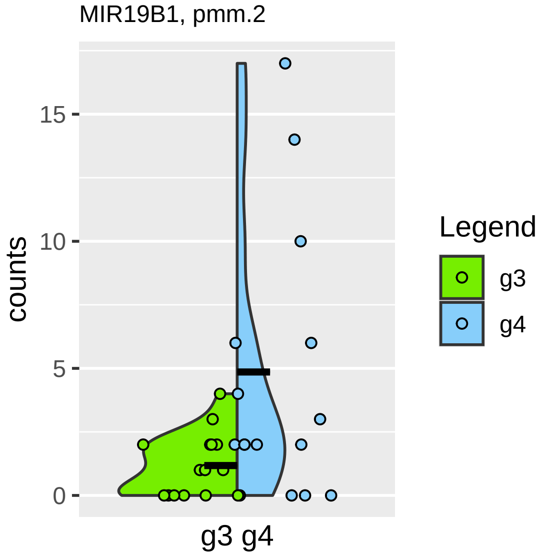

MIR19B1, pmm.2
counts
0
5
10
15
g3 g4
Legend
g3
g4

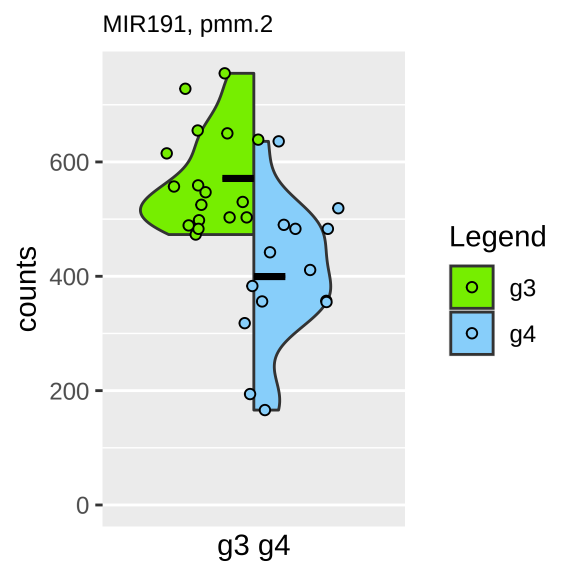
MIR191, pmm.2
counts
600
400
200
0
g3 g4
Legend
g3
g4

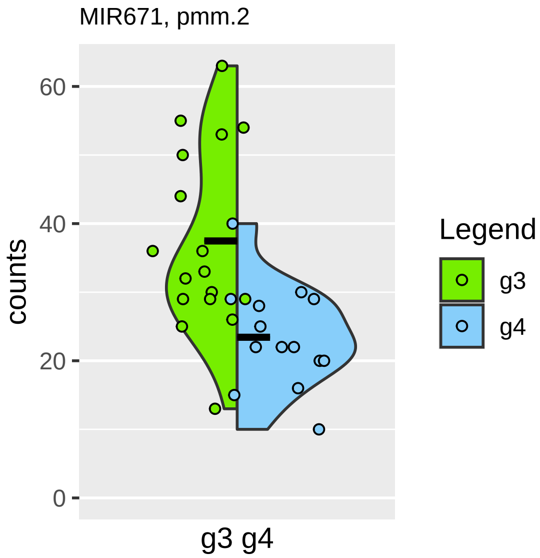
MIR671, pmm.2
counts
0
20
40
60
g3 g4
Legend
g3
g4

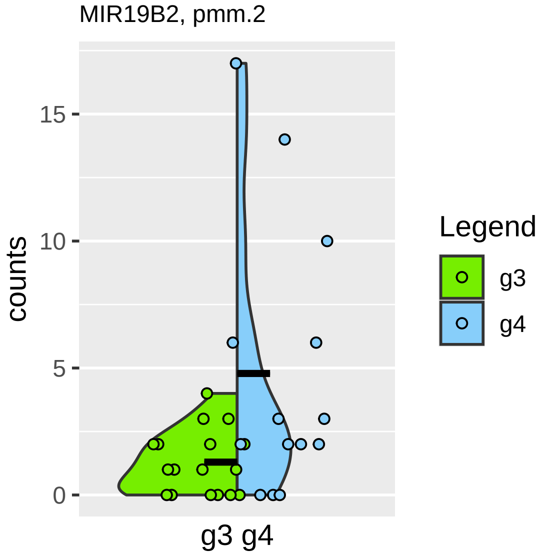
MIR19B2, pmm.2
counts
15
10
5
0
g3 g4
Legend
g3
g4

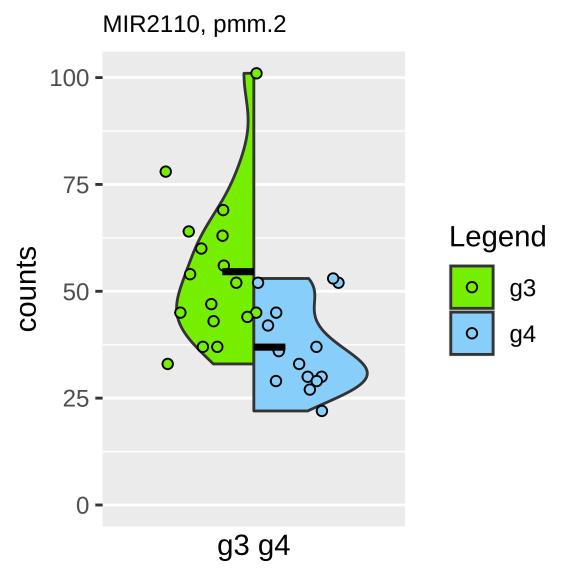

MIR2110, pmm.2
counts
100
75
50
25
0
g3 g4
Legend
g3
g4

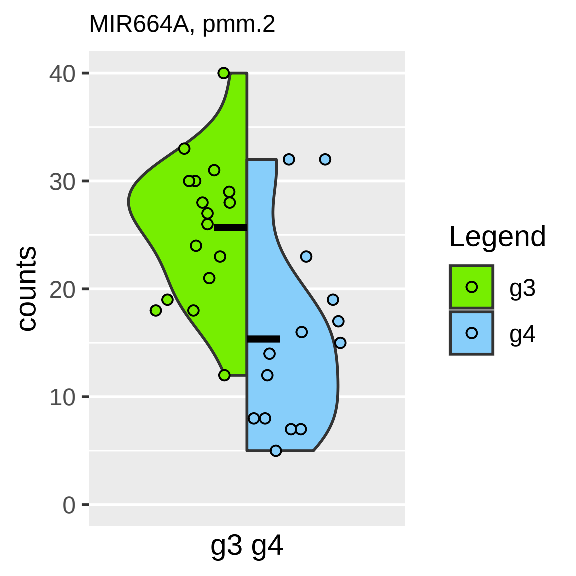

MIR664A, pmm.2
counts
40
30
20
10
0
g3 g4
Legend
g3
g4

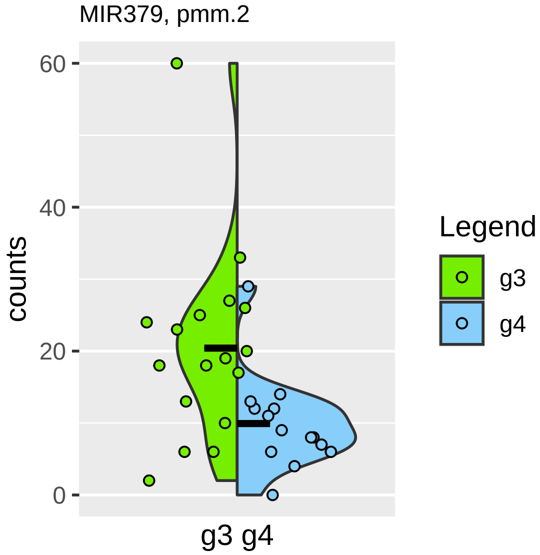
MIR379, pmm.2
counts
60
40
20
0
g3 g4
Legend
g3
g4

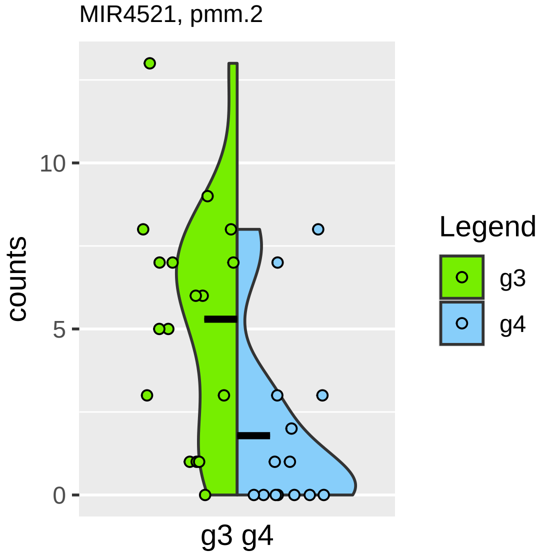
MIR4521, pmm.2
counts
0
5
10
g3 g4
Legend
g3
g4

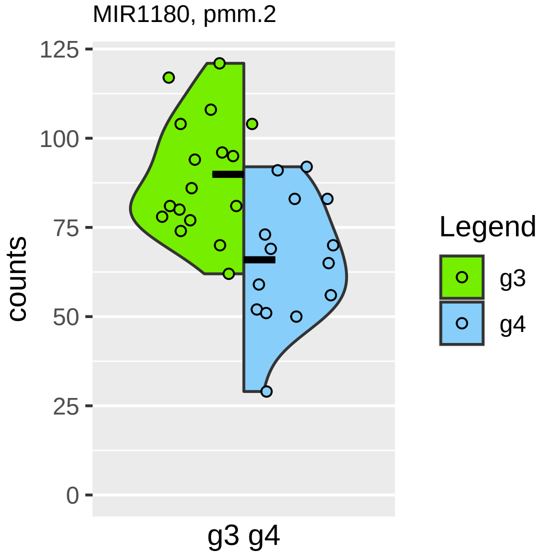

MIR1180, pmm.2
125
100
75
50
25
0
counts
g3 g4
Legend
g3
g4

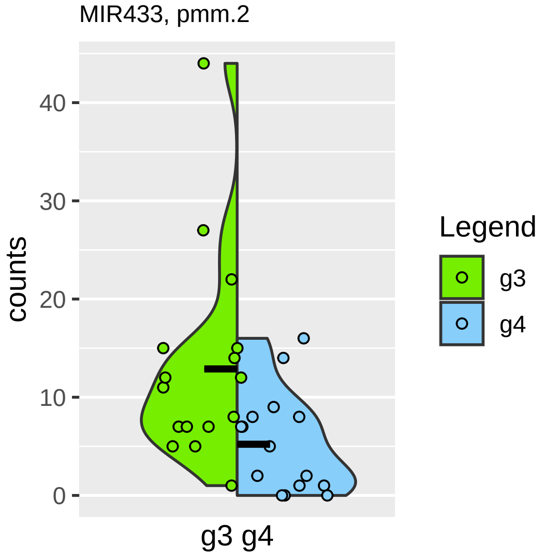

MIR433, pmm.2
counts
40
30
20
10
0
g3 g4
Legend
g3
g4
