## Supplementary material for "Evaluating the effectiveness of various small RNA alignment techniques in transcriptomic analysis by examining different sources of variability through a multi-alignment approach": Supplemetary MAF file: Statistic S1.pdf

### Supplementary information Statistic S1

This supplementary file is including additional information and R code for the Figure 4, 5, 6 and Table 2.

### Table 2 - differential candidates

The analysis was performed based on four basic DESeq2 test choices DESeq (standard) > DESeqMean ('fitType='mean') > nbW (negative binomial Wald test, despite Wald is also fueling standard) > LRT (Likelihood Ratio Test), for all sample groupings and for each of the six framework approaches. The significance level was FDR < 0.05. The DESeq test ranking above is given in a more relaxed test accuracy with 'standard' being the most stringent one. Some discussion on the used tests is given at bioconductor and code details are given below.
Nevertheless, the presented results in Table 2 are based on DESeq standard.

R code:

```
# DE
de.fn <- function(countdata, a1, wpath, wfile, sfile, anno=NULL, cond1=NULL, sign=F, test="DESeq", ret=F){
	# DESeq function
	# countdata: data matrix, samples in columns
	# a1: column data - conditions, vector e.g. c("A","N","N","N","A","N","A","C", ...)
	# wpath, wfile: path & file name results,  wpath, sfile: report file
	# anno: annotation: e.g. miR names - needs the row names of the countdata file
	# cond1: the pairwise condition which should become a result: contrast: e.g. c("condition","A","N")  fc=A/N
	# test: standard: DESeq, other: DESeqMean, nbW, LRT see below
	# ret: return results or not
	coldata <- data.frame(       # numbers with as.factor()
			row.names=dimnames(countdata)[[2]],
			condition=as.factor(a1),
			type=as.factor(a1)
	)
	cat("\n")
	#print(coldata)
	cat("\n nrow coldata ",nrow(coldata))

	## QC on filtered data
	cat("\n countdata<0  ",sum(countdata[countdata<0]))
	cat("\n countdata==0 ",sum(countdata[countdata==0]))
#	return(list(countdata=countdata,coldata=coldata))

	## create deseq data structure for differential analysis
	# countdata:  rows: transcript IDs  cols: sample names
	#          DESeqDataSetFromMatrix
	dds.mat <- DESeqDataSetFromMatrix(
			countData = countdata,
			colData = coldata,
			design = ~ condition)
	#		design = ~ batch + condition)        # if error here -> Please read the vignette section 'Model matrix not full rank':  seems to be no work around

	dds.mat <- estimateSizeFactors(dds.mat)

	# check properties
	cat("\n nrow dds.mat ",nrow(dds.mat),"\n")

	## differential
	if(test=="DESeq"){
		dds.mat <- DESeq(dds.mat)
	}else if(test=="DESeqMean"){
		dds.mat <- DESeq(dds.mat, fitType="mean")           # fitType mean for sparse miR data -- sometimes even this is crashing
	}else if(test=="nbW"){
		dds.mat <- estimateDispersionsGeneEst(dds.mat)
		dispersions(dds.mat) <- mcols(dds.mat)$dispGeneEst
		dds.mat <- nbinomWaldTest(dds.mat)
```

```
	}else if(test=="LRT"){
		dds.mat <- estimateDispersionsGeneEst(dds.mat)
		dispersions(dds.mat) <- mcols(dds.mat)$dispGeneEst
		dds.mat <- nbinomLRT(dds.mat, reduced= ~1)
	}

	## results 1
	# list names
	if(is.null(cond1)){
		cat("\n dds.mat obj ",resultsNames(dds.mat),"\n")
		cat("\n >>> choose a contrast argument and restart\n")
		return()
	}else{
		cat("\n chosen ",cond1,"\n")
	}

	# A-B
	res.A.N <- results(dds.mat, contrast=cond1)
	# order by adjusted p-value
	res.A.N <- as.data.frame(res.A.N)
	res.A.N[is.na(res.A.N$pvalue), "pvalue"] <- 1 	# technically substitute NA by 1 for 
intersection
	res.A.N[is.na(res.A.N$padj), "padj"] <- 1 	# technically substitute NA by 1 for 
intersection
	res.A.N <- res.A.N[order(res.A.N$pvalue), ]

	# number of significants
	yall <- nrow(res.A.N)
	y1true <- sum(res.A.N$pvalue<0.05)
	y2true <- sum(res.A.N$padj<0.05)
	cat("\n <0.05 :  FALSE  TRUE")
	cat("\n sign. p      ",yall-y1true," ",y1true)
	cat("\n sign. padj   ",yall-y2true," ",y2true)

	# write number of significants in a file
	fConn <- file(paste(wpath,"/",sfile,".tsv",sep=""), open="a")
	writeLines(paste(wfile, "pF", "pT", "padjF", "padjT", sep="\t"), fConn)
	writeLines(paste("", (yall-y1true), y1true, (yall-y2true), y2true, sep="\t"), fConn)
	writeLines("", fConn)
	close(fConn)

	# add additional annotations - needs row.names
	if(!is.null(anno)){
		res.A.N.df <- merge(as.data.frame(res.A.N), as.data.frame(anno), by="row.names", 
sort=F)
		row.names(res.A.N.df) <- res.A.N.df[,1]
		res.A.N.df <- res.A.N.df[,-1]
		res.A.N <- res.A.N.df
	}

	# create a  data+results  dataframe
	dcm <- as.data.frame(counts(dds.mat, normalized=T)-1)
	dcm <- round(dcm, 0)
	sti <- as.data.frame(res.A.N)
	#sti <- round(sti, 2)
	res.A.N.df <- merge(sti, dcm, by="row.names", sort=F)
	row.names(res.A.N.df) <- res.A.N.df[,1]
	res.A.N.df <- res.A.N.df[,-1]
	res.A.N.df <- res.A.N.df[order(res.A.N.df$padj), ]  # it is - but to assure
	# keep only sign.
	if(sign){
		res.A.N.df <- res.A.N.df[res.A.N.df$padj<0.05, ]
	}

	# write data DE used/adjusted counts
	write.table(x=res.A.N.df, file=paste(wpath,"/",wfile,".tsv",sep=""), sep="\t", dec=".", 
col.names=NA)		# NA means colnames with first blank
	cat("\n\n")
	if(ret){ return(res.A.N.df) }else{ return() }
}
```

### Figure 4 - Boxplots and ratios

Left column (blue)
All raw counts of all expressed RNA entities [total counts]
Middle column (green)
Only the counts for the subfraction of microRNAs [microRNA counts]
Right column (text)
The ratio : microRNA counts / total counts

The Pearson correlation coefficient is calculated between these two columns on the basis of the mean or median of all feature counts (RNA entities) per sample. Default parameters used. The coefficient of these coherently ordered columns is close to zero in the case of the mean and a magnitude higher and positive in the case of the median. This is indicating a noisy but coherent situation on the sample level.

R code:

```
### boxplot, sample
a <- c(d.groups$g1,d.groups$g2,d.groups$g3,d.groups$g4)

pdf("methX2025/boxplot01.pdf")
a0 <- aat.sam.chr[,a]
a0.1 <- apply(a0,2, function(x){x[x>=5]})
a1 <- sapply(a0.1, mean)
a1.md <- sapply(a0.1, median)
a1.1 <- sapply(a0.1, sum)

a0 <- aat.sam.mt$sts$mir[,a]
a0.2 <- apply(a0,2, function(x){x[x>=5]})
a2 <- sapply(a0.2, mean)
a2.md <- sapply(a0.2, median)
a2.1 <- sapply(a0.2, sum)

a4 <- length(a2.1)

par(mfrow=c(1,3))
boxplot(a0.1, outline=F, horizontal=T, las=1, cex=0.9, col="lightskyblue", axes=F, xlab="totalRNA\
nexpression [raw counts]", ylab="samples")
axis(side=1)
boxplot(a0.2, outline=F, horizontal=T, las=1, cex=0.9, col="chartreuse2", axes=F, xlab="microRNA\
nexpression [raw counts]", ylab="samples")
axis(side=1)
plot(1,1, type="n", xlim=c(0.1,1.9), ylim=c(0.5,a4+0.5), axes=F, xlab="proportion\nmicroRNA counts /
total counts", ylab="")
a5 <- a2.1/a1.1
a6 <- dimnames(a0)[[2]]
for(i in 1:a4){
       text(x=0.1, y=i, labels=paste(formatC(a5[i], , format="e", digits=2),sep=""), cex=0.7, adj=0)
       text(x=1.1, y=i, labels=a6[i], cex=0.7, adj=0)
}
a3 <- cor(a1,a2)
a3.md <- cor(a1.md,a2.md)
title(main=paste("cor mean ",round(a3,3)," cor median ",round(a3.md,3),sep=""))
par(mfrow=c(1,1))
dev.off()
```

**Figure 5 - Venn diagrams**

The diagrams indicate the overlap between all resulting molecule classes across the described 5 approaches (sts, stm, btm, btq, bbm see Table 2).
Every box color denote one approach. Dark red : overlap between all groups, yellow : specific for one group.

R code:

```
#  Venn diagram
# install.packages("BiocManager"); BiocManager::install(c("RBGL","graph"));
# install.packages("devtools"); library(devtools);
# install_github("js229/Vennerable")
library(Vennerable)
vignette("Venn")


## make an overlap view between pmm and sts / stm / btq / btm / bbm
# manual translation  miR... into MIR...
# names(a) :  "sts1" "sts2"  "stm1" "stm2"  "btq1" "btq2"  "btm1" "btm2"  "bbm1" "bbm2"  "pmm1"
"pmm2"
a1[[12]]

a1.1 <- NULL
a1.1[[1]] <-
c("MIR106B","MIR425","MIR363","MIR532","MIR210","MIRLET7D","MIR181A2","MIR185","MIR132","MIR221","MI
R25","MIR30C","MIR484","MIR17","MIR34C",
               "MIR30B","MIR106B","MIR1307","MIR21","MIR135A","MIR128","MIR16","MIR21","MIR652","MIR
455","MIR181A","MIR140","MIR199B","MIR424","MIR3184",
               "MIR31","MIR95","MIR423","MIR181A","MIR337","MIR144","MIR361","MIR30A","MIR22","MIR26
B","MIR200A","MIR30B","MIR103B","MIR103A","MIR486","MIR23A")
a1.1[[2]] <-
c("MIR193B","MIR320A","MIR483","MIR432","MIRLET7B","MIR320B","MIR561","MIR664A","MIR1307","MIR433","
MIR19B","MIR191","MIR342","MIR379")
a1.1[[3]] <- c("MIR106B","MIR425","MIR363","MIR532","MIR210","MIRLET7D","MIR181A-
2","MIR185","MIR221","MIR132","MIR25","MIR30C","MIR484","MIR17","MIR34C",
               "MIR30B","MIR106B","MIR1307","MIR21","MIR135A","MIR128","MIR16","MIR21","MIR652","MIR
455","MIR181A","MIR140","MIR199B","MIR424","MIR3184",
               "MIR31","MIR95","MIR423","MIR181A","MIR337","MIR144","MIR361","MIR30A","MIR22","MIR10
3A","MIR103B","MIR26B","MIR200A","MIR30B","MIR486","MIR23A")
a1.1[[4]] <-
c("MIR193B","MIR320A","MIR483","MIR432","MIRLET7B","MIR320B","MIR664A","MIR433","MIR19B","MIR191","M
IR342","MIR379")
a1.1[[5]] <-
c("MIR425","MIR200A","MIR136","MIR25","MIR363","MIR16","MIR181A2","MIR132","MIR221","MIR199A","MIR53
2","MIR185","MIRLET7D","MIR484","MIR30B",
               "MIR93","MIR361","MIR181A","MIR199B","MIR135A","MIR34C","MIR22")
a1.1[[6]] <-
c("MIR193B","MIR320A","MIR483","MIR432","MIR664A","MIR335","MIRLET7B","MIR561","MIR1180","MIR342","M
IR433")
a1.1[[7]] <-
c("MIR106B","MIR425","MIR532","MIR363","MIR199A","MIR210","MIR221","MIR185","MIR17","MIRLET7D","MIR1
03A","MIR25","MIR199B","MIR30C","MIR132",
               "MIR1307","MIR106B","MIR34C","MIR424","MIR181A2","MIR95","MIR21","MIR128","MIR484","M
IR140","MIR7","MIR1275","MIR21","MIR30A","MIR652",
               "MIR16","MIR107","MIR30B","MIR135A","MIR486","MIR409","MIR181A","MIR337","MIR574","MI
R30B","MIR629","MIR103B","MIR199B","MIR455","MIR361")
a1.1[[8]] <-
c("MIR193B","MIR320A","MIR483","MIRLET7B","MIR4454","MIR432","MIR664A","MIR433","MIR12136","MIR191",
"MIR342","MIR1260B","MIR379","MIR1260A")
a1.1[[10]] <- c("MIR4668","MIR193B")
a1.1[[11]] <- a1[[11]]
a1.1[[12]] <- a1[[12]]


pdf(file="methX2025/Venn_01_1.pdf", height=11, width=7)
av <- Venn( list(sts=unlist(a1[[1]]), stm=unlist(a1[[3]]), btq=unlist(a1[[5]]), btm=unlist(a1[[7]]),
bbm=unlist(a1[[9]])) )
av <- compute.Venn(av, doWeights=T, doEuler=F, type="battle")
av2 <- VennThemes(av, colourAlgorithm="signature")         # signature, binary, sequential
plot(av, gpList=av2)
dev.off()
pdf(file="methX2025/Venn_01_2.pdf", height=11, width=7)
```

```
av <- Venn( list(sts=unlist(a1[[2]]), stm=unlist(a1[[4]]), btq=unlist(a1[[6]]), btm=unlist(a1[[8]]),
bbm=unlist(a1[[10]])) )
av <- compute.Venn(av, doWeights=T, doEuler=F, type="battle")
av2 <- VennThemes(av, colourAlgorithm="signature")         # signature, binary, sequential
plot(av, gpList=av2)
dev.off()

# unique elements
length(unique(c(sts=unlist(a1[[1]]), stm=unlist(a1[[3]]), btq=unlist(a1[[5]]), btm=unlist(a1[[7]]),
bbm=unlist(a1[[9]]))))
length(c(sts=unlist(a1[[1]]), stm=unlist(a1[[3]]), btq=unlist(a1[[5]]), btm=unlist(a1[[7]]),
bbm=unlist(a1[[9]])))
# 57 unique elements of 159 input elements
length(unique(c(sts=unlist(a1[[2]]), stm=unlist(a1[[4]]), btq=unlist(a1[[6]]), btm=unlist(a1[[8]]),
bbm=unlist(a1[[10]]))))
length(c(sts=unlist(a1[[2]]), stm=unlist(a1[[4]]), btq=unlist(a1[[6]]), btm=unlist(a1[[8]]),
bbm=unlist(a1[[10]])))
# 21 unique elements of 53 input elements


pdf(file="methX2025/Venn_03_1.pdf", height=11, width=7)
av <- Venn( list(sts=unlist(a1.1[[1]]), pmm=unlist(a1.1[[11]])) )
av <- compute.Venn(av, doWeights=T, doEuler=F, type="circles")
av2 <- VennThemes(av, colourAlgorithm="signature")         # signature, binary, sequential
plot(av, gpList=av2)
dev.off()
pdf(file="methX2025/Venn_03_2.pdf", height=11, width=7)
av <- Venn( list(btq=unlist(a1.1[[5]]), pmm=unlist(a1.1[[11]])) )
av <- compute.Venn(av, doWeights=T, doEuler=F, type="circles")
av2 <- VennThemes(av, colourAlgorithm="signature")         # signature, binary, sequential
plot(av, gpList=av2)
dev.off()
pdf(file="methX2025/Venn_03_3.pdf", height=11, width=7)
av <- Venn( list(btm=unlist(a1.1[[7]]), pmm=unlist(a1.1[[11]])) )
av <- compute.Venn(av, doWeights=T, doEuler=F, type="circles")
av2 <- VennThemes(av, colourAlgorithm="signature")         # signature, binary, sequential
plot(av, gpList=av2)
dev.off()

pdf(file="methX2025/Venn_04_1.pdf", height=11, width=7)
av <- Venn( list(sts=unlist(a1.1[[2]]), pmm=unlist(a1.1[[12]])) )
av <- compute.Venn(av, doWeights=T, doEuler=F, type="circles")
av2 <- VennThemes(av, colourAlgorithm="signature")         # signature, binary, sequential
plot(av, gpList=av2)
dev.off()
pdf(file="methX2025/Venn_04_2.pdf", height=11, width=7)
av <- Venn( list(btq=unlist(a1.1[[6]]), pmm=unlist(a1.1[[12]])) )
av <- compute.Venn(av, doWeights=T, doEuler=F, type="circles")
av2 <- VennThemes(av, colourAlgorithm="signature")         # signature, binary, sequential
plot(av, gpList=av2)
dev.off()
pdf(file="methX2025/Venn_04_3.pdf", height=11, width=7)
av <- Venn( list(btm=unlist(a1.1[[8]]), pmm=unlist(a1.1[[12]])) )
av <- compute.Venn(av, doWeights=T, doEuler=F, type="circles")
av2 <- VennThemes(av, colourAlgorithm="signature")         # signature, binary, sequential
plot(av, gpList=av2)
dev.off()
```

### Figure 6 - Violin plots

Each diagram is created as a single graph per page, and specific graphs are manually assembled into one panel or automatically into a full group panel.
These violin plots display the density distribution and the actual count size per sample. The horizontal distribution of counts is introduced by the plot function to ensure that all points are visible (which is not always the case). The y-axis indicates the count number, while the x-axis serves as a measure for the density curve itself but is not explicitly plotted here.
This type of graph is well-suited for gaining a better understanding of the situation behind the p-values.
If dots are replaced with sample numbers, this could additionally indicate possible sample subgroup structures.

R code:

```
###  start importing DESeq counts values
a.deseq <- read.table("methX2025/DESeq/0res_de_overview_DESeq.tsv", header=T)
a.deseq <- a.deseq[c(1:22,25:26),]   #28-4=24
a <- row.names(a.deseq)
names(a.deseq)
k <- 1
a1 <- vector("list",0)       # creating supporting variable : a1
while(T){
        cat("\n",a[k])
        a1[[a[k]]] <- read.table(paste("methX2025/DESeq/",a[k],".tsv",sep=""), header=T)
        k <- k+1
        if(k>24){ break }     #24
}

sum(a.deseq[,4])      # 542 violin plot elements


# a.rag : max,  a1,  a2 feature names
i <- 1
k <- 1
l <- 1
a2 <- vector("character",0)          # creating supporting variable : a2
a.rag <- NULL
while(T){
        cat("\n",a[k])
        ade <- a.deseq[k, 4]
        if(ade>0){
                a.tmp2 <- d.coll[[i]]
                a.bl1 <- d.groups[[a.tmp2[3]]]       # names
                a.bl2 <- d.groups[[a.tmp2[4]]]
                a.tmp <- a1[[a[k]]]           # mat collection
                for(j in 1:ade){
                        a.feat <- a.tmp[j, 1]
                        a2[l] <- a.feat
                        one <- unlist(a.tmp[j, a.bl1])
                        two <- unlist(a.tmp[j, a.bl2])
                        a.rag[l] <- max(one,two)
                        l <- l+1
                }
        }
        a2[l] <- "-"; l <- l+1
        i <- i+1
        if(i>2){ i <- 1; a2[l] <- "---"; l <- l+1 }         # 2 (4) DE
        k <- k+1
        if(k>24){ break }     #24
}
# cf.  a.rag  with  a2  and select perfect  a.rag max  -- or -- let rag=NULL


###  creating all images at once,   based on  a1  and  a2 (feature names)
pdf("methX2025/test_violin_03.pdf",width=3,height=3)
i <- 1
k <- 1
l <- 1
a2 <- vector("character",0)
```

```
while(T){
        adn <- a[k]
        cat("\n",adn)
        for(sr in c("\\.DESeq","qm\\.","qt\\.","qp\\.")){
                adn <- sub(sr, "", adn)
        }
        ade <- a.deseq[k, 4]
        if(ade>0){
                a.tmp2 <- d.coll[[i]]
                a.bl1 <- d.groups[[a.tmp2[3]]]       # names
                a.bl2 <- d.groups[[a.tmp2[4]]]
                a.tmp <- a1[[a[k]]]           # mat collection
                for(j in 1:ade){
                        a.feat <- a.tmp[j, 1]
                        a2[l] <- a.feat
                        for(sr in c("Homo_sapiens_","hsa-")){
                                a.feat <- sub(sr, "", a.feat)
                        }
                        one <- unlist(a.tmp[j, a.bl1])
                        two <- unlist(a.tmp[j, a.bl2])
                        fa <- violin.split1(x=paste(a.tmp2[3]," ",a.tmp2[4],sep=""),
                                        y=list(one, two),
                                        z=c(a.tmp2[3],a.tmp2[4]), col2=c("chartreuse2","lightskyblue"),
main=paste(a.feat,adn,sep=", "), rag=NULL, cex=0.8)
                        plot(fa)
                        l <- l+1
                }
        }
        a2[l] <- "-"; l <- l+1
        i <- i+1
        if(i>2){ i <- 1; a2[l] <- "---"; l <- l+1 }         # 2 (4) DE
        k <- k+1
        if(k>24){ break }     #24
}
dev.off()


### three starter below for sts, pi, pmm  images,  resulting in -> three pdf files
# adjust names ...

a.sub <- c(row.names(qm.sts.1.DESeq)[1:10],
                row.names(qm.sts.2.DESeq)[1:10]
)
a.sub <- c(row.names(qp.pi.1.DESeq)[1:20],
                row.names(qp.pi.2.DESeq)[1:20]
)
a.sub <- c(row.names(qp.pmm.1.DESeq)[1:20],
                row.names(qp.pmm.2.DESeq)[1:20]
)

pdf("methX2025/test_violin_04_pmm.pdf",width=11,height=11)        # _mt  _pi  _pmm
i <- 1
k <- 1
l <- 1
m <- 1
a2 <- vector("character",0)
a3 <- NULL
while(T){               # experiment
        adn <- a[k]
        cat("\n",adn)
        ada <- sub("qp\\.", "", sub("\\.DESeq", "", adn))          # 1: qm\\. 2+3: qp\\.
        ade <- a.deseq[k, 4]
        if(ade>0){              # sign. exp.s
                a.tmp2 <- d.coll[[i]]
                a.bl1 <- d.groups[[a.tmp2[3]]]       # names
                a.bl2 <- d.groups[[a.tmp2[4]]]
                a.tmp <- a1[[a[k]]]           # mat collection
                for(j in 1:ade){
                        a.feat <- a.tmp[j, 1]
                        #cat("\n feat-mat",a.feat)
                        if(a.feat%in%a.sub){
                                a2[j] <- a.feat
                                for(sr in c("Homo_sapiens_","hsa-")){
                                        a.feat <- sub(sr, "", a.feat)
                                }
                                one <- unlist(a.tmp[j, a.bl1])
                                two <- unlist(a.tmp[j, a.bl2])
```

```
                                a3[[m]] <- violin.split1b(x=paste(a.tmp2[3]," ",a.tmp2[4],sep=""),
                                               y=list(one, two),
                                               z=c(a.tmp2[3],a.tmp2[4]),
col2=c("chartreuse2","lightskyblue"), main=paste(a.feat,ada,sep=", "), rag=NULL)
                                l <- l+1
                                m <- m+1
                                if(m>25){ m <- 1; ggarrange(plots=a3, ncol=5, nrow=5); a3 <- NULL }
                        }
                }
        }
        a2[l] <- "-"; l <- l+1
        i <- i+1
        if(i>2){ i <- 1; a2[l] <- "---"; l <- l+1 }         # 2 (4) DE
        k <- k+1
        if(k>24){ break }     #24
}
par(mfrow=c(1,1))
dev.off()


#####  basic violin function (could be called directly for a single plot)

violin.split1b <- function(x, y, z, col2, main="", rag=NULL){
        # split violin plots - package introdataviz
        #  design: 1-2
        # x: one character elements for the x position
        # y: two data vectors as a list
        # z: two character elements for all three legend entries as vector
        # col3: three colors as vector ["yellow" "#E69F00"]
        # rag: y scale max value
        require(ggplot2)
        require(ggtext)
        require(introdataviz)
        require(egg)
        if(length(x)!=1 | length(y)!=2 | length(z)!=2){ stop("x needs one and y,z needs two entries")
}
        lenA <- length(y[[1]])
        lenB <- length(y[[2]])
        raA <- range(y[[1]])
        raB <- range(y[[2]])
        if(is.null(rag)){ rag <- max(raA,raB) }
        # color order on page left to right - symbols inside x,m important for order
        aa <- data.frame(
                        x=c(rep("a",(lenA+lenB))),
                        y=c(y[[1]],y[[2]]),
                        m=c(rep("i",lenA),rep("j",lenB)) )
        fa <- ggplot(aa, aes(x=x, y=y, fill=m)) +
                        geom_split_violin() +
                        geom_point(pch=21, position=position_jitterdodge(jitter.width=NULL,
jitter.height=0, dodge.width=0.65, seed=NA)) +
                        scale_fill_manual(values=col2[c(1,2)], name="Legend", labels=c(z[1],z[2])) +
                        labs(x="", y="counts") + scale_x_discrete(name=x[1], labels=NULL, breaks=NULL)
+
                        scale_y_continuous(limits=c(0, rag)) +
                        ggtitle(paste("<span style='font-size: 10pt;'>",main,"</font>",sep="")) +
                        theme(plot.title=element_markdown()) +
                        stat_summary(fun=mean, fun.min=mean, fun.max=mean, geom="crossbar",
width=0.25, show.legend=F, position=position_dodge(width=.25))
                        )
        return(fa)     # ggarrange() in case -> next step
}
```
