## Supplementary material for "Evaluating the effectiveness of various small RNA alignment techniques in transcriptomic analysis by examining different sources of variability through a multi-alignment approach": Supplemetary MAF file: Statistic S2.pdf

**Supplementary information Statistic S2**

In addition to the main considerations outlined in the manuscript, here are some additional ideas on how quality ranking could be conducted for the microRNA class using the five basic approaches. Furthermore, premature RNAs are included, which can be technically considered closely related to the STS group.

Two simple modeling procedures are performed starting from the expressed microRNAs (2-4%), because otherwise the number of non expressed microRNAs would dominate.
The first assumes that all observed microRNAs might be true positives. This rationale is not completely unrealistic from the point that the huge majority of about 2000 annotated microRNAs is negative and with higher basic coverage very likely still more events will appear. The positives are marked by one while the negatives are denoted by 0. So the observed events are given in one vector while the model estimate in a second.
For DE-1 analysis the results for the key parameters accuracy, precision, and recall are given in here in Table 1. DE-2 is described in Table 2.
Table 1 and 2 is based on 75 respectively 28 unique observations. The differences between the data sets are moderate and not directly related to data size. The best performance is observed for the premature microRNAs, which is presumably due to the generally higher alignment count numbers. The bbm drop-out in Table 1 is directly associated with the absence of detected microRNAs. However, BBMap (bbm) appears to have the lowest performance in this case.

| | Table 1<br>DE-1<br>accuracy | precision | recall | Table 2<br>DE-2<br>accuracy | precision | recall |
|---|---|---|---|---|---|---|
| sts | 0.56 | 0.56 | 1 | 0.5 | 0.5 | 1 |
| stm | 0.56 | 0.56 | 1 | 0.42 | 0.42 | 1 |
| btq | 0.29 | 0.29 | 1 | 0.39 | 0.39 | 1 |
| btq | 0.54 | 0.54 | 1 | 0.5 | 0.5 | 1 |
| bbm | 0 | 0 | NA | 0.07 | 0.07 | 1 |
| pmm | 0.65 | 0.65 | 1 | 0.60 | 0.60 | 1 |

Accuracy: 0 to 1 (global not class). Towards 1 a better accuracy (1 – error_rate).
Precision: 0 to 1 (global). Towards 1 a better precision (ppv).
Recall: 0 to 1 (global). Towards 1 a better recall (sensitivity).

The second model approach is based on the consideration that the less often detected microRNAs might be false positives. Based on six approaches the threshold needs to be greater than three of six observations per molecule class and approach. This criteria is fulfilled in DE-1 in 26 of 75 cases and in DE-2 in 10 of 28. So the estimate vector is now not always 1 instead a mixture 0 and 1. Again accuracy, precision, and recall are calculated for all approaches.
Table 3 presents insight into DE-1 while Table 4 shows DE-2.

| | Table 3<br>DE-1<br>accuracy | precision | recall | Table 4<br>DE-2<br>accuracy | precision | recall |
|---|---|---|---|---|---|---|
| sts | 0.78 | 1 | 0.61 | 0.85 | 1 | 0.71 |
| stm | 0.76 | 0.96 | 0.59 | 0.92 | 1 | 0.83 |
| btq | 0.84 | 0.69 | 0.81 | 0.82 | 0.8 | 0.72 |
| btq | 0.77 | 0.96 | 0.60 | 0.85 | 1 | 0.71 |
| bbm | 0.65 | 0 | NA | 0.64 | 0.1 | 0.5 |
| pmm | 0.53 | 0.76 | 0.40 | 0.60 | 0.8 | 0.47 |

The quality in Table 3 and 4 has improved globally compared to Table 1 and 2. The artificial recall values in Table 1 and 2 have now been converted to normal scale values. Overall, this model appears to perform better.
To confirm if this truly reflects reality, further validation is required. It is understood that higher global read coverage for alignment, which involves preparing larger sample libraries, typically results in more expressed molecule classes on average. Despite the various considerations mentioned to identify reliable true positive candidates, this statistical approach may help address this challenge.

R code:

```
# install.packages("metrica")
library(metrica)

## we are not showing all the data customization procedures necessary to start this statistics
#  (several pages in this data design)
#  but
#  candidate intersections need to be prepared

## some exploratory data insight
#  sts 1,2 [46,14] / stm 1,2 [46,12] / btq 1,2 [22,11] / btm 1,2 [45,14] / bbm 1,2 [0,2] / pmm 1,2
[49,17]
#   1,2 : DE1 DE2
# name sets [[1]] to [[12]] already created

# number of entries
a1.1[[1]]
sum(unlist(lapply(a1.1, length)))    # 278
a <- a1.1[c(1,3,5,7,9,11)]
sum(unlist(lapply(a, length)))       # 208
a <- a1.1[c(2,4,6,8,10,12)]
sum(unlist(lapply(a, length)))       # 70

# unique entries
a <- unlist(a1.1)
a1 <- unique(a)               # 101
a <- unlist(a1.1[c(1,3,5,7,9,11)])
a1 <- unique(a)               # 75
a <- unlist(a1.1[c(2,4,6,8,10,12)])
a1 <- unique(a)               # 28


## simple modeling

## model 1 : estimate: all are expressed - upper extreme

a.stat1 <- matrix(0,6,3)
a.m <- rep(1,nrow(a.iuset1))
dimnames(a.stat1)[[1]] <- c("sts1","stm1","btq1","btm1","bbm1","pmm1")
dimnames(a.stat1)[[2]] <- c("accuracy","precision","recall")
for(i in 1:6){
       a.stat1[i,1] <- unlist( rm.attr( accuracy(obs=factor(a.iuset1[,i], levels=c("0","1")),
pred=factor(a.m, levels=c("0","1")), tidy=T)))
       a.stat1[i,2] <- unlist( rm.attr( precision(obs=factor(a.iuset1[,i], levels=c("0","1")),
pred=factor(a.m, levels=c("0","1")), tidy=T)))
       a.stat1[i,3] <- unlist( rm.attr( recall(obs=factor(a.iuset1[,i], levels=c("0","1")),
pred=factor(a.m, levels=c("0","1")), pos_level=2, tidy=T)))
}

confusion_matrix(obs=factor(a.iuset1[,1], levels=c("0","1")), pred=factor(a.m, levels=c("0","1")),
plot=F, unit="count")
# only for [1]
#          OBSERVED
# PREDICTED  0  1
#         0  0  0
#         1 33 42
write.table(a.stat1, file="a.stat1.txt", append=F, sep="\t", row.names=T, col.names=T)

a.stat2 <- matrix(0,6,3)
a.m <- rep(1,nrow(a.iuset2))
dimnames(a.stat2)[[1]] <- c("sts2","stm2","btq2","btm2","bbm2","pmm2")
dimnames(a.stat2)[[2]] <- c("accuracy","precision","recall")
```

```
for(i in 1:6){
	a.stat2[i,1] <- unlist( rm.attr( accuracy(obs=factor(a.iuset2[,i], levels=c("0","1")),
pred=factor(a.m, levels=c("0","1")), tidy=T)))
	a.stat2[i,2] <- unlist( rm.attr( precision(obs=factor(a.iuset2[,i], levels=c("0","1")),
pred=factor(a.m, levels=c("0","1")), tidy=T)))
	a.stat2[i,3] <- unlist( rm.attr( recall(obs=factor(a.iuset2[,i], levels=c("0","1")),
pred=factor(a.m, levels=c("0","1")), pos_level=2, tidy=T)))
}

confusion_matrix(obs=factor(a.iuset2[,1], levels=c("0","1")), pred=factor(a.m, levels=c("0","1")),
plot=F, unit="count")
# only for [1]
#           OBSERVED
# PREDICTED  0  1
#          0  0  0
#          1 14 14
write.table(a.stat2, file="a.stat2.txt", append=F, sep="\t", row.names=T, col.names=T)


## model 2 : estimate: some are expressed - criteria : observation frequency, range; threshold >3
(range) -> 1, rest -> 0

ncol(a.iuset1)        # 6
a <- rowSums(a.iuset1)
range(a)              # 1..5
a.m <- rep(0, nrow(a.iuset1)) # rows 75
a.m[a > 3] <- 1              # sum(a.m) 26

a.stat3 <- matrix(0,6,3)
dimnames(a.stat3)[[1]] <- c("sts1","stm1","btq1","btm1","bbm1","pmm1")
dimnames(a.stat3)[[2]] <- c("accuracy","precision","recall")
for(i in 1:6){
	a.stat3[i,1] <- unlist( rm.attr( accuracy(obs=factor(a.iuset1[,i], levels=c("0","1")),
pred=factor(a.m, levels=c("0","1")), tidy=T)))
	a.stat3[i,2] <- unlist( rm.attr( precision(obs=factor(a.iuset1[,i], levels=c("0","1")),
pred=factor(a.m, levels=c("0","1")), tidy=T)))
	a.stat3[i,3] <- unlist( rm.attr( recall(obs=factor(a.iuset1[,i], levels=c("0","1")),
pred=factor(a.m, levels=c("0","1")), pos_level=2, tidy=T)))
}

confusion_matrix(obs=factor(a.iuset1[,1], levels=c("0","1")), pred=factor(a.m, levels=c("0","1")),
plot=F, unit="count")
# only for [1]
#           OBSERVED
# PREDICTED  0  1
#          0 33 16
#          1  0 26
write.table(a.stat3, file="a.stat3.txt", append=F, sep="\t", row.names=T, col.names=T)

ncol(a.iuset2)        # 6
a <- rowSums(a.iuset2)
range(a)              # 1..5
a.m <- rep(0, nrow(a.iuset2)) # rows 28
a.m[a > 3] <- 1              # sum(a.m) 10

a.stat4 <- matrix(0,6,3)
dimnames(a.stat4)[[1]] <- c("sts1","stm1","btq1","btm1","bbm1","pmm1")
dimnames(a.stat4)[[2]] <- c("accuracy","precision","recall")
for(i in 1:6){
	a.stat4[i,1] <- unlist( rm.attr( accuracy(obs=factor(a.iuset2[,i], levels=c("0","1")),
pred=factor(a.m, levels=c("0","1")), tidy=T)))
	a.stat4[i,2] <- unlist( rm.attr( precision(obs=factor(a.iuset2[,i], levels=c("0","1")),
pred=factor(a.m, levels=c("0","1")), tidy=T)))
	a.stat4[i,3] <- unlist( rm.attr( recall(obs=factor(a.iuset2[,i], levels=c("0","1")),
pred=factor(a.m, levels=c("0","1")), pos_level=2, tidy=T)))
}

confusion_matrix(obs=factor(a.iuset2[,1], levels=c("0","1")), pred=factor(a.m, levels=c("0","1")),
plot=F, unit="count")
# only for [1]
#           OBSERVED
# PREDICTED  0  1
#          0 14  4
#          1  0 10
write.table(a.stat4, file="a.stat4.txt", append=F, sep="\t", row.names=T, col.names=T)
```
