## Supplementary material for "Evaluating the effectiveness of various small RNA alignment techniques in transcriptomic analysis by examining different sources of variability through a multi-alignment approach": Supplemetary MAF file: Table S1.pdf

### Table S1. Overview on resources, files and terms.

All references here are based on Homo sapiens.

| **Program for alignment** | **STAR** | **STAR** | **STAR** | **Bowtie2** | **Bowtie2** | **Bowtie2** | **BBMap** |
|---|---|---|---|---|---|---|---|
| Fasta file | mt.trans.hu.miRtRNA.<br>111.primary.fa | Homo_sapiens.<br>GRCh38.111.dna.<br>primary_assembly.fa | mte.trans.hu.GRCh38.<br>111.primary.fa | m2_miRtRNA.fasta | Homo_sapiens.<br>GRCh38.111.dna.<br>primary_assembly.fa | r_star_pirbase_gold<br>.fasta | m2_miRtRNA.fasta |
| Index folder | star2711x111HUmt | star2711x111HUpmm | star2711x111HUmte | bt2_mt | bt2_111HU | bt2_pir | bb_mt |
| GTF file | mt.Homo_sapiens.<br>miRtRNA.gtf | Homo_sapiens.<br>GRCh38.111.<br>mirbase.gtf | mte.Homo_sapiens.<br>GRCh38.111.gtf | no gtf | mt.Homo_sapiens.<br>miRtRNA.gtf<br>(for QualiMap) | no gtf | no gtf |
| Program parameter | outFilterMismatchNmax 0,<br>outFilterMultimapNmax 2,<br>outSAMmultNmax 1,<br>quantMode<br>TranscriptomeSAM. | outFilterMismatchNmax 0,<br>outFilterMultimapNmax 2,<br>outSAMmultNmax 1,<br>quantMode<br>TranscriptomeSAM. | outFilterMismatchNmax 0,<br>outFilterMultimapNmax 2,<br>outSAMmultNmax 1,<br>quantMode<br>TranscriptomeSAM. | --end-to-end | --local | --local | no special features used<br>(global) |
| Program to quantify | a) Salmon<br>b) Samtools idxstats | a) Samtools idxstats | a) Salmon<br>b) Samtools idxstats | a) Samtools idxstats | a) Samtools idxstats **<br>b) Qualimap count | a) Samtools idxstats | a) Samtools idxstats |
| Result acronyms | a) sts<br>b) stm | a) pmm | a) **<br>b) mte ** | a) btm | a) chr **<br>b) btq | a) pi | a) bbm |
| Result folder name | 03star1out | 03star2out | 03star3out | 04bt21out | 04bt22out | 04bt23out | 05bbm1out |
| Quantification output file(s) | a) quant.sf<br>b) mcounts.tsv | a) mcounts.tsv | a) mcounts.tsv | a) mcounts.tsv | a) mcounts.tsv **<br>b) qm.111HUmte.tsv | a) mcounts.tsv | a) mcounts.tsv |
| Reference features | 2653 mirbase microRNAs,<br>260 tRNAs | 1852 premature<br>microRNAs | 39 gene_biotype items<br>with now 4114450 (v113)<br>elements | 2653 mirbase microRNAs,<br>260 tRNAs | 39 gene_biotype items<br>with now 4114450 (v113)<br>elements | 19222 pirbase small RNAs | 2653 mirbase microRNAs,<br>260 tRNAs |
| Result id type | ENST - extra ENST<br>names for micro/tRNAs<br>(will be translated by R<br>functions to a microRNA<br>[mirbase] or tRNA name<br>[tRNA database]) | regular ENST name of<br>premature microRNAs | regular ENST name of all<br>features included in<br>standard Homo_sapiens.<br>GRCh38.111 | mirbase name | chromosome and contig<br>names<br>respectively<br>ENSG micro/tRNA name | pirbase name | mirbase name |
| Script function name | **staroutA** | **staroutB** | **staroutC** | **bt2outA** | **bt2outB** | **bt2outC** | **bbmoutA** |
| Support file for R import | namelist.txt | namelist.txt | namelist.txt | namelist.txt | namelist.txt | namelist.txt | namelist.txt |

**Note :**
1) Not every aligner is used in all three script files.
2)  **  : not used here.
