## Supplementary material for "Evaluating the effectiveness of various small RNA alignment techniques in transcriptomic analysis by examining different sources of variability through a multi-alignment approach": Supplemetary MAF file: Table S2.pdf

### Table S2. View on the replication of significant candidates between the different approaches.

Only two replicating candidates are shown in blue to exemplify the concept of the table.
Cf. also Figure 5.
Table sorted top to bottom by increasing p value.

**DE-1**

| STAR mt index, Salmon quantification | STAR mt index, Samtools quantification | Bowtie2 mt index, Samtools quantification | Bowtie2 Qualimap quantification, mt reference | BBMap mt index, Samtools quantification | STAR pmm index, Samtools quantification |
|---|---|---|---|---|---|
| sts | stm | btq | btm | bbm | pmm |
| hsa-miR-106b-3p | hsa-miR-106b-3p | hsa-miR-425-5p | hsa-miR-106b-3p | | MIR106B |
| hsa-miR-425-5p | hsa-miR-425-5p | hsa-miR-200a-3p | hsa-miR-425-5p | | MIR31 |
| hsa-miR-363-3p | hsa-miR-363-3p | hsa-miR-136-3p | hsa-miR-532-5p | | MIR532 |
| hsa-miR-532-5p | hsa-miR-532-5p | hsa-miR-25-3p | hsa-miR-363-3p | | MIR210 |
| hsa-miR-210-3p | hsa-miR-210-3p | hsa-miR-363-3p | hsa-miR-199a-3p | | MIR30B |
| hsa-let-7d-3p | hsa-let-7d-3p | hsa-miR-16-5p | hsa-miR-210-3p | | MIR363 |
| hsa-miR-181a-2-3p | hsa-miR-181a-2-3p | hsa-miR-181a-2-3p | hsa-miR-221-3p | | MIR185 |
| hsa-miR-185-5p | hsa-miR-185-5p | hsa-miR-132-3p | hsa-miR-185-5p | | MIR497 |
| hsa-miR-132-3p | hsa-miR-221-3p | hsa-miR-221-3p | hsa-miR-17-3p | | MIR30C2 |
| hsa-miR-221-3p | hsa-miR-132-3p | hsa-miR-199a-3p | hsa-let-7d-3p | | MIR30C1 |
| hsa-miR-25-3p | hsa-miR-25-3p | hsa-miR-532-5p | hsa-miR-103a-3p | | MIR193B |
| hsa-miR-30c-5p | hsa-miR-30c-5p | hsa-miR-185-5p | hsa-miR-25-3p | | MIR34C |
| hsa-miR-484 | hsa-miR-484 | hsa-let-7d-3p | hsa-miR-199b-3p | | MIR1307 |
| hsa-miR-17-3p | hsa-miR-17-3p | hsa-miR-484 | hsa-miR-30c-5p | | MIR181A2 |
| hsa-miR-34c-5p | hsa-miR-34c-5p | hsa-miR-30b-5p | hsa-miR-132-3p | | MIR25 |
| hsa-miR-30b-5p | hsa-miR-30b-5p | hsa-miR-93-5p | hsa-miR-1307-3p | | MIR16-2 |
| hsa-miR-106b-5p | hsa-miR-106b-5p | hsa-miR-361-3p | hsa-miR-106b-5p | | MIR17 |
| hsa-miR-1307-3p | hsa-miR-1307-3p | hsa-miR-181a-5p | hsa-miR-34c-5p | | MIR16-1 |
| hsa-miR-21-3p | hsa-miR-21-3p | hsa-miR-199b-5p | hsa-miR-424-5p | | MIR652 |
| hsa-miR-135a-5p | hsa-miR-135a-5p | hsa-miR-135a-5p | hsa-miR-181a-2-3p | | MIR181A1 |
| hsa-miR-128-3p | hsa-miR-128-3p | hsa-miR-34c-5p | hsa-miR-95-3p | | MIR382 |
| hsa-miR-16-5p | hsa-miR-16-5p | hsa-miR-22-3p | hsa-miR-21-3p | | MIR23A |
| hsa-miR-21-5p | hsa-miR-21-5p | | hsa-miR-128-3p | | MIR128-1 |
| hsa-miR-652-3p | hsa-miR-652-3p | | hsa-miR-484 | | MIR425 |
| hsa-miR-455-5p | hsa-miR-455-5p | | hsa-miR-140-5p | | MIR93 |
| hsa-miR-181a-5p | hsa-miR-181a-5p | | hsa-miR-7-5p | | MIR199B |
| hsa-miR-140-5p | hsa-miR-140-5p | | hsa-miR-1275 | | MIR660 |
| hsa-miR-199b-5p | hsa-miR-199b-5p | | hsa-miR-21-5p | | MIR135A2 |
| hsa-miR-424-5p | hsa-miR-424-5p | | hsa-miR-30a-3p | | MIR128-2 |
| hsa-miR-3184-5p | hsa-miR-3184-5p | | hsa-miR-652-3p | | MIR451A |
| hsa-miR-31-5p | hsa-miR-31-5p | | hsa-miR-16-5p | | MIR451B |
| hsa-miR-95-3p | hsa-miR-95-3p | | hsa-miR-107 | | MIR1287 |
| hsa-miR-423-3p | hsa-miR-423-3p | | hsa-miR-30b-5p | | MIR21 |
| hsa-miR-181a-3p | hsa-miR-181a-3p | | hsa-miR-135a-5p | | MIR22 |
| hsa-miR-337-3p | hsa-miR-337-3p | | hsa-miR-486-3p | | MIR103A1 |
| hsa-miR-144-3p | hsa-miR-144-3p | | hsa-miR-409-3p | | MIR26B |
| hsa-miR-361-3p | hsa-miR-361-3p | | hsa-miR-181a-5p | | MIR103A2 |
| hsa-miR-30a-3p | hsa-miR-30a-3p | | hsa-miR-337-3p | | MIR629 |
| hsa-miR-22-3p | hsa-miR-22-3p | | hsa-miR-574-3p | | MIR181D |
| hsa-miR-26b-5p | hsa-miR-103a-3p | | hsa-miR-30b-3p | | MIR103B1 |
| hsa-miR-200a-5p | hsa-miR-103b | | hsa-miR-629-5p | | MIR103B2 |
| hsa-miR-30b-3p | hsa-miR-26b-5p | | hsa-miR-103b | | MIR95 |
| hsa-miR-103b | hsa-miR-200a-5p | | hsa-miR-199b-5p | | MIR337 |
| hsa-miR-103a-3p | hsa-miR-30b-3p | | hsa-miR-455-5p | | MIR132 |
| hsa-miR-486-3p | hsa-miR-486-3p | | hsa-miR-361-3p | | MIR92A1 |
| hsa-miR-23a-5p | hsa-miR-23a-5p | | | | MIR484 |
| | | | | | MIR339 |
| | | | | | MIR29A |
| | | | | | MIR136 |

**DE-2**

| STAR mt index, Salmon quantification | STAR mt index, Samtools quantification | Bowtie2 mt index, Samtools quantification | Bowtie2 Qualimap quantification, mt reference | BBMap mt index, Samtools quantification | STAR pmm index, Samtools quantification |
|---|---|---|---|---|---|
| sts | stm | btq | btm | bbm | pmm |
| hsa-miR-193b-5p | hsa-miR-193b-5p | hsa-miR-193b-5p | hsa-miR-193b-5p | hsa-miR-4668-5p | MIR320A |
| hsa-miR-320a-3p | hsa-miR-320a-3p | hsa-miR-320a-3p | hsa-miR-320a-3p | hsa-miR-193b-5p | MIR193B |
| hsa-miR-483-5p | hsa-miR-483-5p | hsa-miR-483-5p | hsa-miR-483-5p | | MIRLET7B |
| hsa-miR-432-5p | hsa-miR-432-5p | hsa-miR-432-5p | hsa-let-7b-5p | | MIR432 |
| hsa-let-7b-5p | hsa-let-7b-5p | hsa-miR-664a-5p | hsa-miR-4454 | | MIR320B2 |
| hsa-miR-320b | hsa-miR-320b | hsa-miR-335-5p | hsa-miR-432-5p | | MIR320B1 |
| hsa-miR-561-5p | hsa-miR-664a-5p | hsa-let-7b-5p | hsa-miR-664a-5p | | MIR561 |
| hsa-miR-664a-5p | hsa-miR-433-3p | hsa-miR-561-5p | hsa-miR-433-3p | | MIR19B1 |
| hsa-miR-1307-5p | hsa-miR-19b-3p | hsa-miR-1180-3p | hsa-miR-12136 | | MIR191 |
| hsa-miR-433-3p | hsa-miR-191-5p | hsa-miR-342-3p | hsa-miR-191-5p | | MIR671 |
| hsa-miR-19b-3p | hsa-miR-342-3p | hsa-miR-433-3p | hsa-miR-342-3p | | MIR19B2 |
| hsa-miR-191-5p | hsa-miR-379-5p | | hsa-miR-1260b | | MIR2110 |
| hsa-miR-342-3p | | | hsa-miR-379-5p | | MIR664A |
| hsa-miR-379-5p | | | hsa-miR-1260a | | MIR379 |
| | | | | | MIR4521 |
| | | | | | MIR1180 |
| | | | | | MIR433 |
