## Supplementary material for "Evaluating the effectiveness of various small RNA alignment techniques in transcriptomic analysis by examining different sources of variability through a multi-alignment approach": Supplemetary MAF file: Table S3.pdf

### Table S3. Looking into the real counts of two selected microRNAs discussed in the 'Method Validation' section.

## DE-2

| bbm | baseMean | log2FC | lfcSE | stat | pvalue | padj | EH01 | EH02 | EH03 | EH04 | EH05 | EH06 | EH07 | EH08 | EH09 | EH10 | EH11 | EH12 | EH13 | EH14 | EH15 | EH16 | EH17 | E01 | E02 | E03 | E04 | E06 | E07 | E08 | E09 | E10 | E11 | E12 | E13 | E14 | E15 |
|---|---|---|---|---|---|---|---|---|---|---|---|---|---|---|---|---|---|---|---|---|---|---|---|---|---|---|---|---|---|---|---|---|---|---|---|---|---|
| hsa-miR-193b-5p | 3.484 | -1.606 | 0.361 | -4.451 | 8.54E-06 | 8.60E-03 | 1 | 3 | 4 | 4 | 4 | 3 | 1 | 6 | 7 | 7 | 3 | 2 | 5 | 4 | 5 | 6 | 3 | 3 | 2 | 0 | 0 | 0 | 0 | 0 | 1 | 0 | 1 | 1 | 0 | 0 | 1 |
| hsa-miR-4668-5p | 5.387 | -1.635 | 0.297 | -5.502 | 3.75E-08 | 7.55E-05 | 6 | 8 | 8 | 3 | 9 | 4 | 6 | 6 | 9 | 12 | 6 | 5 | 6 | 3 | 7 | 6 | 11 | 3 | 1 | 1 | 5 | 0 | 0 | 4 | 0 | 1 | 1 | 4 | 1 | 0 | 0 |

| sts | baseMean | log2FC | lfcSE | stat | pvalue | padj | EH01 | EH02 | EH03 | EH04 | EH05 | EH06 | EH07 | EH08 | EH09 | EH10 | EH11 | EH12 | EH13 | EH14 | EH15 | EH16 | EH17 | E01 | E02 | E03 | E04 | E06 | E07 | E08 | E09 | E10 | E11 | E12 | E13 | E14 | E15 |
|---|---|---|---|---|---|---|---|---|---|---|---|---|---|---|---|---|---|---|---|---|---|---|---|---|---|---|---|---|---|---|---|---|---|---|---|---|---|
| hsa-miR-193b-5p | 6.839 | -2.319 | 0.365 | -6.349 | 2.16E-10 | 7.05E-08 | 4 | 15 | 12 | 6 | 9 | 3 | 4 | 11 | 11 | 16 | 9 | 11 | 12 | 3 | 7 | 21 | 11 | 7 | 0 | 0 | 2 | 0 | 1 | 0 | 0 | 0 | 2 | 1 | 3 | 0 | 0 |
| hsa-miR-4668-5p | 1.001 | 0.002 | 0.566 | 0.004 | 9.97E-01 | 1.00E+00 | 0 | 0 | 0 | 0 | 0 | 0 | 0 | 0 | 0 | 0 | 0 | 0 | 0 | 0 | 0 | 0 | 0 | 0 | 0 | 0 | 0 | 0 | 0 | 0 | 0 | 0 | 0 | 0 | 0 | 0 | 0 |

| stm | baseMean | log2FC | lfcSE | stat | pvalue | padj | EH01 | EH02 | EH03 | EH04 | EH05 | EH06 | EH07 | EH08 | EH09 | EH10 | EH11 | EH12 | EH13 | EH14 | EH15 | EH16 | EH17 | E01 | E02 | E03 | E04 | E06 | E07 | E08 | E09 | E10 | E11 | E12 | E13 | E14 | E15 |
|---|---|---|---|---|---|---|---|---|---|---|---|---|---|---|---|---|---|---|---|---|---|---|---|---|---|---|---|---|---|---|---|---|---|---|---|---|---|
| hsa-miR-193b-5p | 6.839 | -2.319 | 0.365 | -6.361 | 2.01E-10 | 5.73E-08 | 4 | 15 | 12 | 6 | 9 | 3 | 4 | 11 | 11 | 16 | 9 | 11 | 12 | 3 | 7 | 21 | 11 | 7 | 0 | 0 | 2 | 0 | 1 | 0 | 0 | 0 | 2 | 1 | 3 | 0 | 0 |
| hsa-miR-4668-5p | 1.001 | 0.002 | 0.562 | 0.004 | 9.97E-01 | 1.00E+00 | 0 | 0 | 0 | 0 | 0 | 0 | 0 | 0 | 0 | 0 | 0 | 0 | 0 | 0 | 0 | 0 | 0 | 0 | 0 | 0 | 0 | 0 | 0 | 0 | 0 | 0 | 0 | 0 | 0 | 0 | 0 |

| btq | baseMean | log2FC | lfcSE | stat | pvalue | padj | EH01 | EH02 | EH03 | EH04 | EH05 | EH06 | EH07 | EH08 | EH09 | EH10 | EH11 | EH12 | EH13 | EH14 | EH15 | EH16 | EH17 | E01 | E02 | E03 | E04 | E06 | E07 | E08 | E09 | E10 | E11 | E12 | E13 | E14 | E15 |
|---|---|---|---|---|---|---|---|---|---|---|---|---|---|---|---|---|---|---|---|---|---|---|---|---|---|---|---|---|---|---|---|---|---|---|---|---|---|
| hsa-miR-193b-5p | 5.162 | -2.292 | 0.385 | -5.954 | 2.62E-09 | 7.83E-07 | 3 | 12 | 9 | 2 | 7 | 2 | 4 | 9 | 6 | 11 | 6 | 10 | 8 | 1 | 5 | 15 | 10 | 2 | 0 | 0 | 2 | 0 | 1 | 0 | 0 | 0 | 1 | 0 | 3 | 0 | 0 |
| hsa-miR-4668-5p | 1.001 | 0.002 | 0.564 | 0.004 | 9.97E-01 | 1.00E+00 | 0 | 0 | 0 | 0 | 0 | 0 | 0 | 0 | 0 | 0 | 0 | 0 | 0 | 0 | 0 | 0 | 0 | 0 | 0 | 0 | 0 | 0 | 0 | 0 | 0 | 0 | 0 | 0 | 0 | 0 | 0 |

| btm | baseMean | log2FC | lfcSE | stat | pvalue | padj | EH01 | EH02 | EH03 | EH04 | EH05 | EH06 | EH07 | EH08 | EH09 | EH10 | EH11 | EH12 | EH13 | EH14 | EH15 | EH16 | EH17 | E01 | E02 | E03 | E04 | E06 | E07 | E08 | E09 | E10 | E11 | E12 | E13 | E14 | E15 |
|---|---|---|---|---|---|---|---|---|---|---|---|---|---|---|---|---|---|---|---|---|---|---|---|---|---|---|---|---|---|---|---|---|---|---|---|---|---|
| hsa-miR-193b-5p | 9.258 | -2.072 | 0.325 | -6.379 | 1.79E-10 | 4.98E-08 | 6 | 18 | 13 | 8 | 10 | 11 | 5 | 13 | 16 | 24 | 11 | 13 | 16 | 5 | 9 | 28 | 17 | 8 | 2 | 0 | 4 | 0 | 2 | 1 | 0 | 3 | 5 | 2 | 4 | 0 | 2 |
| hsa-miR-4668-5p | 1.000 | 0.000 | 0.537 | 0.000 | 1.00E+00 | 1.00E+00 | 0 | 0 | 0 | 0 | 0 | 0 | 0 | 0 | 0 | 0 | 0 | 0 | 0 | 0 | 0 | 0 | 0 | 0 | 0 | 0 | 0 | 0 | 0 | 0 | 0 | 0 | 0 | 0 | 0 | 0 | 0 |

#### Mirbase

```
Accession      MIMAT0004767
Description    hsa-miR-193b-5p mature miRNA

Sequence

14 - CGGGGUUUUGAGGGCGAGAUGA - 35

Hairpins

            u               G  A  A  uua
guggucucagaa CGGGGUUUUGAGGGC AG UG gu   u
|||||||||||| ||||||||||||||| || || ||   g
uacuggguuuU GCCCUGAAACUCCCG UC Ac ua   u
            C               G  A  c  uuu

guggucucagaauCGGGGUUUUGAGGGCGAGAUGAguuuauguuuauccAACUGGCCCUCAAAGUCCCGCUuuuggggucau
((((((((((((.(((((((((((((((.((.((.((.........)).)).)).))))))))))))))).))))))))))))
```

```
Accession      MIMAT0019745
Description    hsa-miR-4668-5p mature miRNA

Sequence

1 - AGGGAAAAAAAAAAAGGAUUUGUC - 23

Hairpin

-A       A           G    gu  c   g
  GGGAAAA AAAAAGGAUUU UCuu  ag cag a
  ||||||| ||||||||||| ||||  || |||
  CCUUUUU UUUUUCCUAAA AGaa  uu guu u
GA       G           -    au  u   a

AGGGAAAAAAAAAAAGGAUUUGUCuuguagccaggauauuguuuuaaaGAAAAUCCUUUUUGUUUUUCCAG
.(((((((.(((((((((((.((((..((.(((....))).))..))))))))))))))))).)))))))..
```

**Distribution of reads**

Sample number : 85

| Name | Molecule counts | Negative samples | Range per sample |
|---|---|---|---|
| miR-193b-5p | 410 | 9 | 0-28 |
| miR-193b-3p | 339 | 18 | 0-63 |
| miR-4668-5p | 3 | 83 | 0-2 |
| miR-4668-3p | 0 | 85 | NA |

**Motif per read**

| Name | Dominant motif | Very rare |
|---|---|---|
| miR-193b-5p | CGGGGTTTTGAGGGCGAGATGA | TTTGAGGGCGAGATGA<br>CGGGGTTTTGAGGGCGAGAT |
| miR-193b-3p | AACTGGCCCTCAAAGTCCCGCT | ACTGGCCCTCAAAGTCCCGCT<br>AACTGGCCCTCAAAGTCCCGC |
| miR-4668-5p | AGGGAAAAAAAAAAAGGATTTGT | / |
| miR-4668-3p | / | / |
